# Discovery of 1,3,4-oxadiazoles with slow-action activity against *Plasmodium falciparum* malaria parasites

**DOI:** 10.1101/2023.03.09.531229

**Authors:** Katherine T. Andrews, Gillian M. Fisher, Meaghan Firmin, Andris J. Liepa, Tony Wilson, James Gardiner, Yacine Mohri, Anjana Rai, Andrew K. Davey, Antoine Masurier, Alix Delion, Alexandos A. Mouratidis, Oliver Hutt, Jeremy N. Burrows, John H. Ryan, Andrew G. Riches, Tina S. Skinner-Adams

## Abstract

To achieve malaria eradication, new preventative agents that act differently to front-line treatment drugs are needed. To identify potential chemoprevention starting points we screened a sub-set of the CSIRO Australia Compound Collection for compounds with slow-action *in vitro* activity against *Plasmodium falciparum*. This work identified *N*,*N*-dialkyl-5-alkylsulfonyl-1,3,4-oxadiazol-2-amines as a new antiplasmodial chemotype (e.g., **1** 96 h IC_50_ 550 nM) with a different action to delayed-death slow-action drugs. Structure activity relationship analysis of analogues identified multiple compounds with potent and selective *in vitro* activity against drug-sensitive and multi-drug resistant *Plasmodium* parasites (e.g., **31** and **32** 96 h IC_50_ <40 nM; SI >2,500). However subsequent studies in mice with lead compound **1**, which had the best microsomal stability of the compounds assessed, demonstrated rapid clearance (T_1/2_ ≤1.6 h) and poor oral *in vivo* efficacy. This indicates that improvements in the pharmacokinetic profile of *N*,*N*-dialkyl-5-alkylsulfonyl-1,3,4-oxadiazol-2-amines would be needed for the development of this chemotype for malaria chemoprophylaxis.

## INTRODUCTION

Almost half the world’s population is at risk of malaria, with 247 million cases and 619,000 deaths, mostly children under five years of age, attributed to this parasitic disease in 2021.^1^ While the World Health Organization recommends that RTS,S/AS01 (Mosquirix^TM^) be provided to all children in moderate to high malaria transmission areas to improve this position,^2^ this vaccine provides only ∼30% protection against severe malaria in this group,^3–6^ highlighting the need for multiple malaria control tools. Effective prevention and treatment drugs remain a critical part of malaria control, and more widely for the malaria eradication endgame strategy. New treatment drugs with novel modes of action are required to overcome the significant problem of malaria parasite drug resistance, including to the current gold-standard artemisinin-based combination treatment therapies (ACTs).^7–10^ New prophylactic chemotherapeutics are also needed to protect vulnerable populations in areas of high endemicity as we transition towards eradication and immunity begins to wane.

Malaria prophylaxis is currently dependent on a limited number of drugs that primarily target asexual blood-stage parasites, the life cycle stage responsible for the clinical symptoms of malaria. These drugs aim to kill parasites before the development of symptoms. However, the development of specific prophylactic drugs lags behind the development of treatment drugs and most available prophylactic agents are also used for treatment and are associated with resistant parasites or other limitations. Current prophylactic agents include: doxycycline, an antibiotic which requires daily dosing; mefloquine which is associated with neurological side-effects;^11, 12^ the atovaquone-proguanil combination (Malarone^®^)^13^, which is used for treatment and prophylaxis and; the 8-aminoquinoline tafenoquine, which is approved for prophylaxis and the radical cure of *P. vivax* malaria^14^. A limitation of tafenoquine is that it may cause severe hemolytic anemia in individuals with glucose-6-phosphate (G6PD) deficiency (∼8% prevalence rate in malaria-endemic regions{Baird, 2018 #51;Howes, 2012 #72}).

As prophylactic agents are vital to protect children and pregnant women in areas of seasonal and perennial transmission, as well as for malaria eradication^16^, and current agents have limitations, new, safe, effective, and affordable options are needed. Ideally these drugs should also have long *in vivo* half-lives (>100 h to potentially support monthly dosing) and novel modes of action to avoid cross-resistance with treatment drugs. Compounds with slow parasite killing or delayed death activity are potential prophylactic drug candidates as, unlike treatment drugs that should kill *Plasmodium* malaria parasites quickly to rapidly reduce parasite burden, this is not essential for prophylaxis drugs as subjects receiving the intervention are asymptomatic. The slow-action activity of these agents is also likely to reflect different modes of action and hence such agents are less likely to impact, or to be affected by, clinical resistance to treatment drugs. Here we present data describing the synthesis and activity assessment of a novel antiplasmodial compound series based on the 2-sulfonyl-1,3,4-oxadiazole scaffold with slow-action activity against asexual blood stage parasites.

## RESULTS AND DISCUSSION

### Antiplasmodial activity of hit compounds

Three *N,N*-dialkyl-5-alkylsulfonyl-1,3,4-oxadiazol-2-amines were identified with slow-action antiplasmodial activity following a sub-set screen of the Commonwealth Scientific and Industrial Research Organisation (CSIRO Australia) Compound Collection^17^ for *in vitro* growth inhibition activity against *P. falciparum* 3D7 (*Pf*3D7) asexual forms. Assays were carried out over 48 h (one developmental cycle) and 96 h (two developmental cycles). The criteria for a slow-action hit were 96 h 50% inhibitory concentration (*Pf*IC_50_) ≤1.0 µM (in line with Medicines for Malaria Venture product development partnership criteria^18^) and a 48 h/96 h IC_50_ ratio of >10. Slow action hit compounds **1**, **2** and **3** had 96 h IC_50_ values of 0.55 µM, 0.84 µM and 0.16 µM, and *Pf*3D7 48 h/96 h IC_50_ ratios of >91, 15, and 72, respectively (**Table 1**). As a comparator, the fast action malaria drug chloroquine has a *Pf*3D7 48 h/96 h IC_50_ ratio of 1 (**Table 1**). Three non-active analogues were synthesized providing initial structure activity relationship information. Compared with sulfone **2**, the sulfide **4** and sulfoxide **5** analogues demonstrated >60 and 16-fold weaker antiplasmodial activity at 96 h, respectively. Additionally, **6**, the 1,3,4-thiadiazole analog of **1**, was also inactive (**Table 1**), demonstrating the importance of a 1,3,4-oxadiazole core bearing 2-dialkylamino and 5-alkylsulfonyl groups to antiplasmodial activity.

**Table 1.**
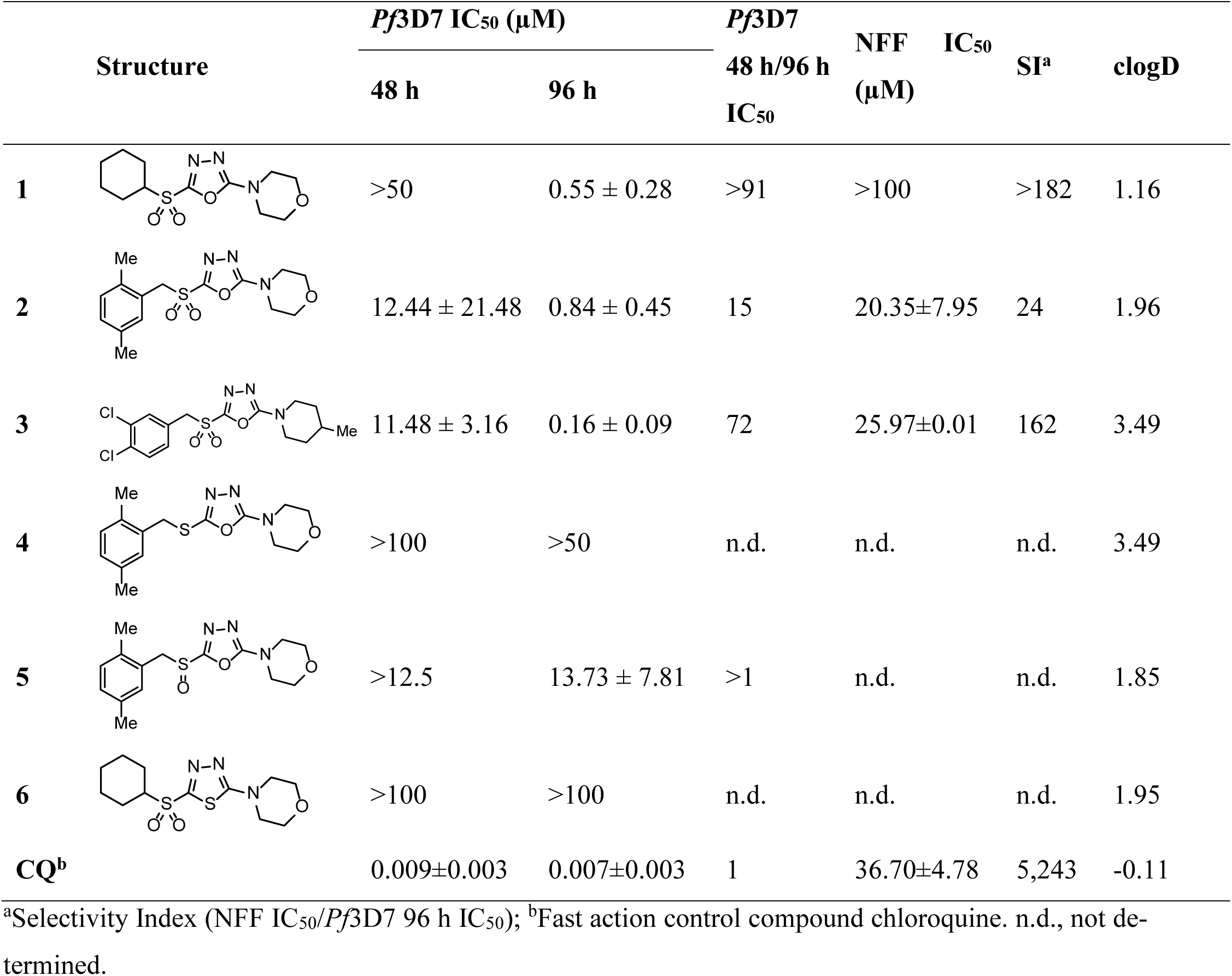
Antiplasmodial profile of slow-action N,N-Dialkyl-5-alkylsulfonyl-1,3,4-oxadiazol-2-amine screening hits 1-3 and related analogues 4-6.

Hit compounds **1**, **2** and **3** maintained similar activity towards the multi-drug resistant *P. falciparum* Dd2 line (*Pf*Dd2) with 96 h IC_50_’s of 0.50 µM, 0.57 µM and 0.07 µM, and *Pf*Dd2 48 h/96 h IC_50_ ratios of >50, 23 and 145, respectively (**Table 2**). When comparing the IC_50_ values of the drug resistant versus drug sensitive *P. falciparum* lines, all three compounds had a resistance index (RI; *Pf*Dd2/*Pf*3D7 IC_50_) ≤1, compared to RI 9 for the control drug chloroquine, to which Dd2 is resistant (**Table 2**). These data indicated that *P. falciparum* Dd2 parasites are not resistant to these slow-action hit compounds.

**Table 2.**
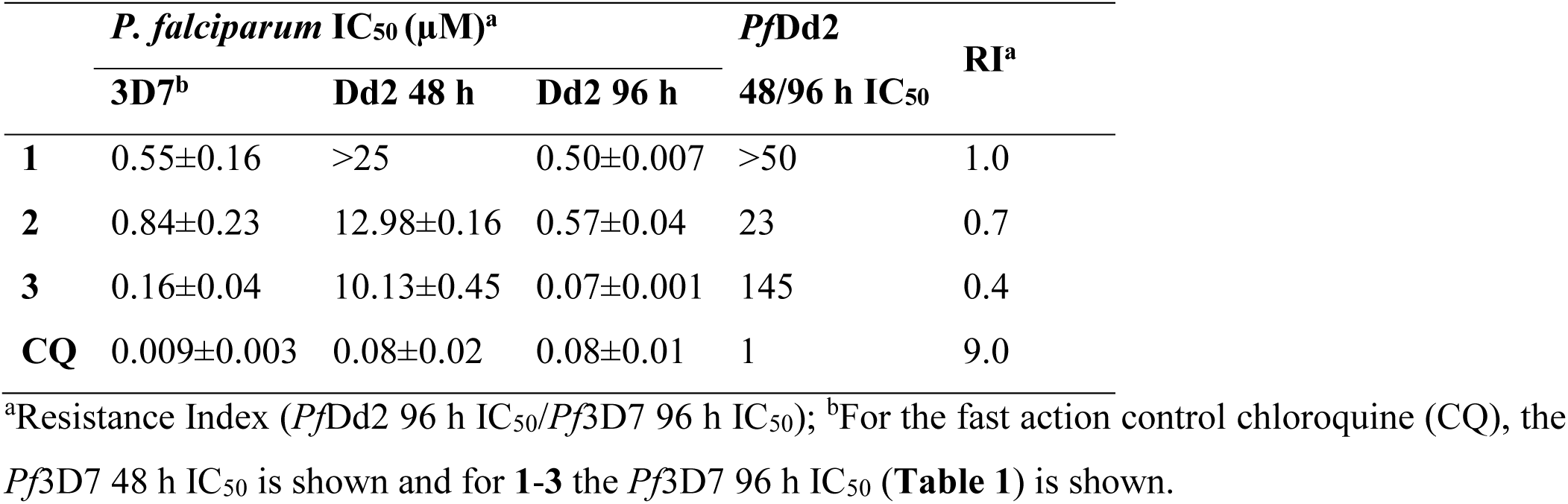
Antiplasmodial activity of 1-3 against multi-drug resistant P. falciparum Dd2 parasites.

To determine if the slow-action activity of **1**-**3** was like that observed for “delayed death” antimalarial drugs, isopentenyl pyrophosphate (IPP) rescue experiments were performed. Delayed-death inhibitors such as the antibiotic clindamycin target the essential *P. falciparum* apicoplast organelle with activity observed in the second cycle after exposure.^19, 20^ The only essential function of the apicoplast is the biosynthesis of isoprenoid precursors such as IPP, which can therefore be used to rescue the effects of “delayed death” inhibitors that target the apicoplast.^21^ The activity of **1**-**3** was not changed in the presence of IPP (P>0.05; **Table 3**), indicating a different action to delayed-death inhibitors. In contrast, the control drug clindamycin showed a 3,571-fold higher IC_50_ with IPP versus without, which confirmed the expected delayed death rescue of this compound’s action (**Table 3**).

**Table 3.** IPP rescue assays with *N,N*-Dialkyl-5-alkylsulfonyl-1,3,4-oxadiazol-2-amine screening hits 1-3 against *P. falciparum* 3D7 parasites.

|  | *P. falciparum* 3D7 IC<sub>50</sub> (μM)<sup>a</sup> |  | Fold Change | P value |
| --- | --- | --- | --- | --- |
|  | - IPP | + IPP |  |  |
| **CQ** | 0.005±0.001 | 0.006±0.003 | 1 | 0.68 |
| **CLIN** | 0.007±0.001 | >25 | 3,571 | n.d. |
| **1** | 0.55±0.28 | 1.79±0.118 | 3 | 0.14 |
| **2** | 0.84±0.45 | 0.84±0.70 | 1 | 1.00 |
| **3** | 0.16±0.09 | 0.20±0.12 | 1 | 0.60 |

<sup>a</sup>IC<sub>50</sub>’s determined in 96 h *in vitro* growth inhibition assays supplemented with and without 200 μM IPP; clindamycin (CLIN), a delayed death inhibitor, and chloroquine (CQ) were included as positive and negative controls, respectively. n.d., not determined.

Given these promising preliminary data suggesting the identification of a new anti-plasmodial structural series with a different mode of action to current drugs, and that SciFinder searches found no reports of *N,N*-dialkyl-5-alkylsulfonyl-1,3,4-oxadiazol-2-amines in the literature, structure-activity relationships were investigated in more detail. Compound **1** was chosen as the starting point for structure-activity relationship studies because, although it was less potent than **3**, it showed markedly better stability towards liver microsomes than both **3** and **2** and showed greater membrane permeability in a Caco-2 assay (see **Table 6**).

### Antiplasmodial structure activity relationships

The sulfonyl R^1^ and NR^2^R^3^ substituents of **1** were varied in turn (**Tables 4** and **5**, respectively). All compounds were assessed for fast-action (48 h assay) and slow-action (48 h assay/96 h assay IC_50_ ratio >10) antiplasmodial activity against *P. falciparum* 3D7 asexual forms. The effect of varying the sulfonyl substituent R^1^ on antiplasmodial activity of compounds with the RHS substituent fixed as morpholine was investigated (**Table 4**). Cyclopentyl **7** (96 h IC_50_ 5.82 µM) was 10-fold less active than cyclohexyl **1** (96 h IC_50_ 0.55 µM), while cycloheptyl **8** was 3-fold more active (96 h IC_50_ 0.18 µM) compared to **1**. Various cycloalkylmethyl groups (**9**, **10**, **11**) gave similar activity to **1**, while the neopentyl analogue (**12**; 96 h IC_50_ 1.34 µM) was >2-fold less potent. Relatively bulky lipophilic R^1^ groups such as adamantylmethyl (**14**; 96 h IC_50_ 0.40 µM) retained similar activity to **1**, whilst incorporation of CF_3_ groups (**15**, **16**; 96 h IC_50_ 3.19 µM and 4.30 µM, respectively) or heteroatoms (**17**, **18**, **19**; 96 h IC_50_ 5.05 µM, >50 µM, and 7.09 µM, respectively) into R^1^ resulted in decreased activity. The possibility that the benzylic methylene group of **2** and **3** (**Table 1**) was a metabolic liability, together with the poor microsomal stability of these compounds (see later) prompted us to examine compound **21**, which proved less potent (96 h IC_50_ 24.63 µM) than its benzyl analogue **20** (96 h IC_50_ 4.20 µM). Notably, analogue **20** (R^1^ = 3,4-dichlorobenzyl, 96 h IC_50_ 4.20 µM) was 26-fold less potent than hit compound **3** (96 h IC_50_ 0.16 µM) with the same R^1^ group but with NR^2^R^3^ = 4-methylpiperidine in place of morpholine.

**Table 4.**
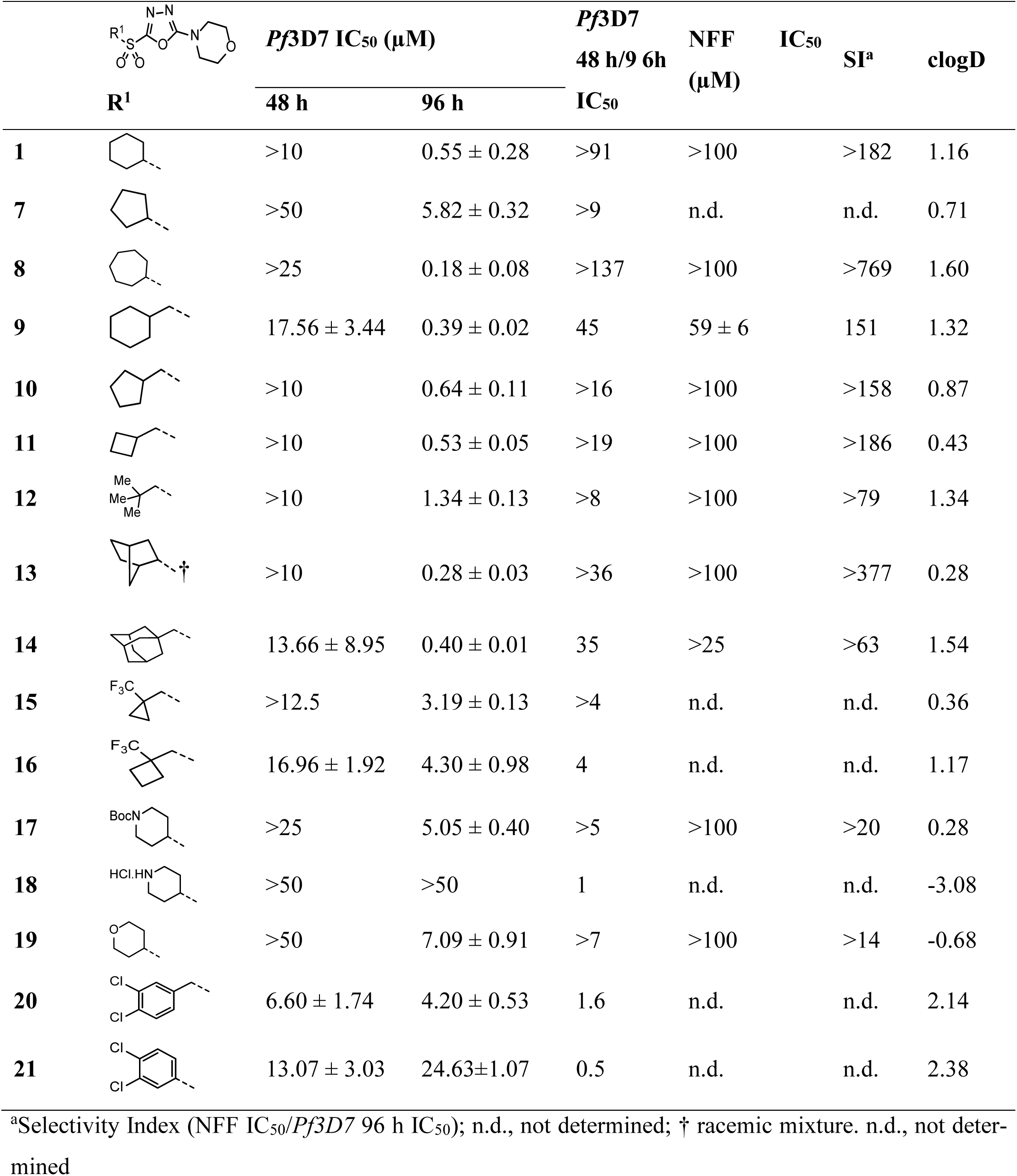
SAR data for N,N-dialkyl-5-alkylsulfonyl-1,3,4-oxadiazol-2-amines with different sulfonyl substituents.

Focus shifted to modifying the RHS amino substituent NR^2^R^3^, with the R^1^ group fixed as cyclohexyl (**Table 5**). Potency was retained when the morpholine ring of **1** (96 h IC_50_ 0.55 µM) was replaced with a piperidine (**23**; 96 h IC_50_ 0.40 µM). Incorporation of a 5-membered pyrrolidine ring (**22**; 96 h IC_50_ 3.53 µM) resulted in decreased activity vs **1**, whereas the 7-membered azepane ring (**24**; 96 h IC_50_ 0.09 µM) displayed increased potency. The oxazepane (**25**; 96 h IC_50_ 0.75 µM) was ∼8-fold less potent than **24**, while 1,2-oxazine **26** was relatively inactive (96 h IC_50_ 18 µM). Addition of a methyl substituent at various positions on the piperidinyl ring increased potency over **23**, with the 3-Me isomer (**28**; 96 h IC_50_ 0.10 µM) showing highest potency. Various isomeric dimethyl piperidines showed increased potency over **23**, with **32**, **33** and **31** (96 h IC_50_ 0.04 µM, 0.08 µM and 0.03 µM, respectively) being the best, albeit at the cost of considerably increased lipophilicity relative to the lead compound **1**. Morpholino analogues bearing either one methyl (**34**) or two (**35**, **36**) were more potent than the unsubstituted morpholino hit **1,** but less potent than their piperidino counterparts (96 h IC_50_ 0.38 µM, 0.24 µM and 0.29 µM, respectively). The bridged morpholine analogue **42** (96 h IC_50_ 0.10 µM, clogP 1.68) and spirocyclic^22^ analogues **37** (96 h IC_50_ 0.10 µM, clogP 1.41) and **39** (96 h IC_50_ 0.11 µM, clogP 1.41) all showed ∼5-fold improved activity relative to **1** (clogP 1.16) with minimal increase in predicted lipophilicity. Other bicyclic NR^2^R^3^ groups failed to show further improvements in potency. Less lipophilic compounds which incorporated either polar functionality (**55**; 96 h IC_50_ 6.75 µM) or ionisable functionality (**56**, **57**; 96 h IC_50_ 3.58 µM and 5.40 µM, respectively), showed decreased activity. However, cyano analogue **49** (96 h IC_50_ 0.49 µM) and methoxy analogue **48** (96 h IC_50_ 0.69 µM) were of similar potency to **1**. Benzo-fused analogues **61** and **62** (96 h IC_50_ 1.12 µM and 2.06 µM, respectively) were less potent than **1**.

**Table 5.**
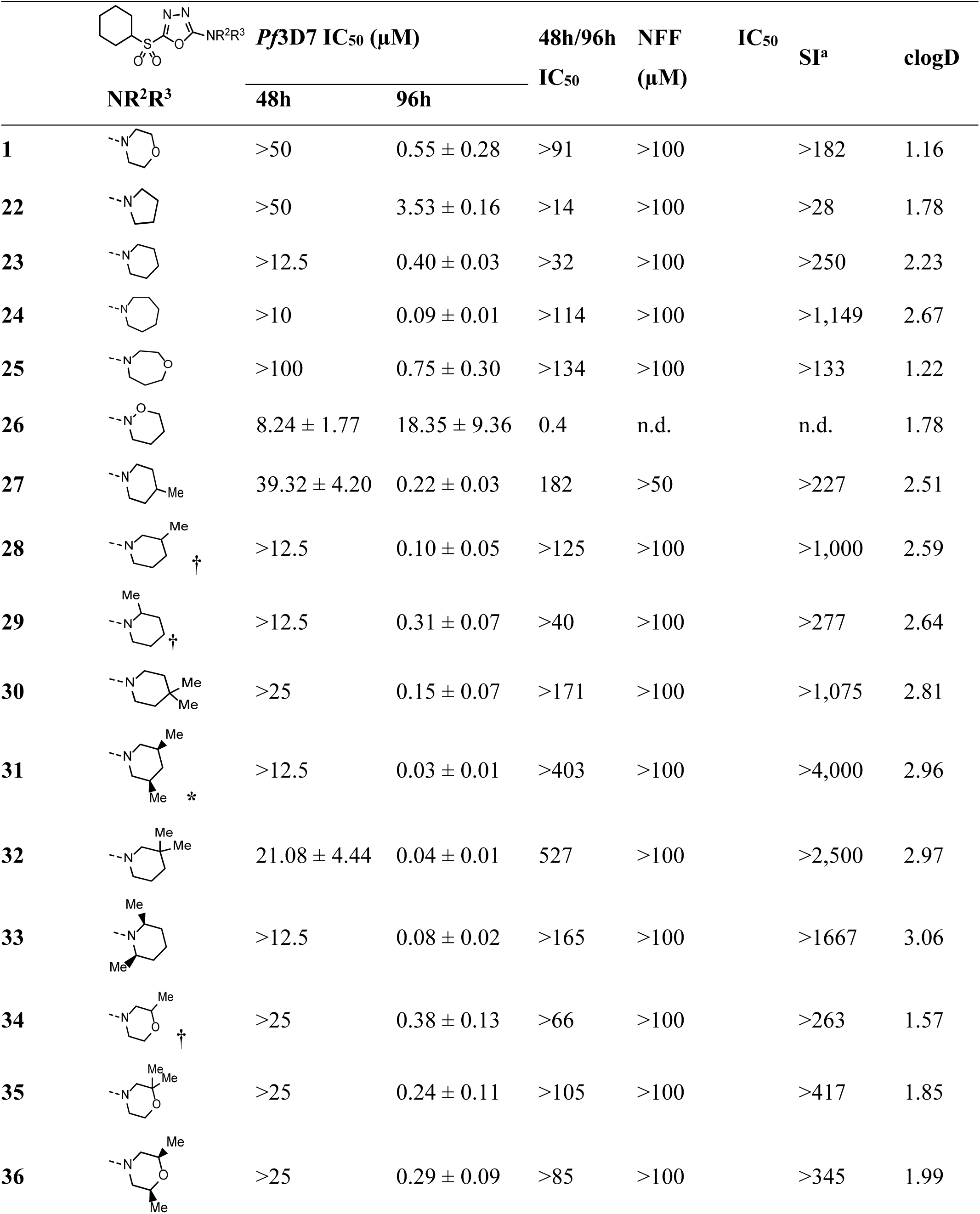
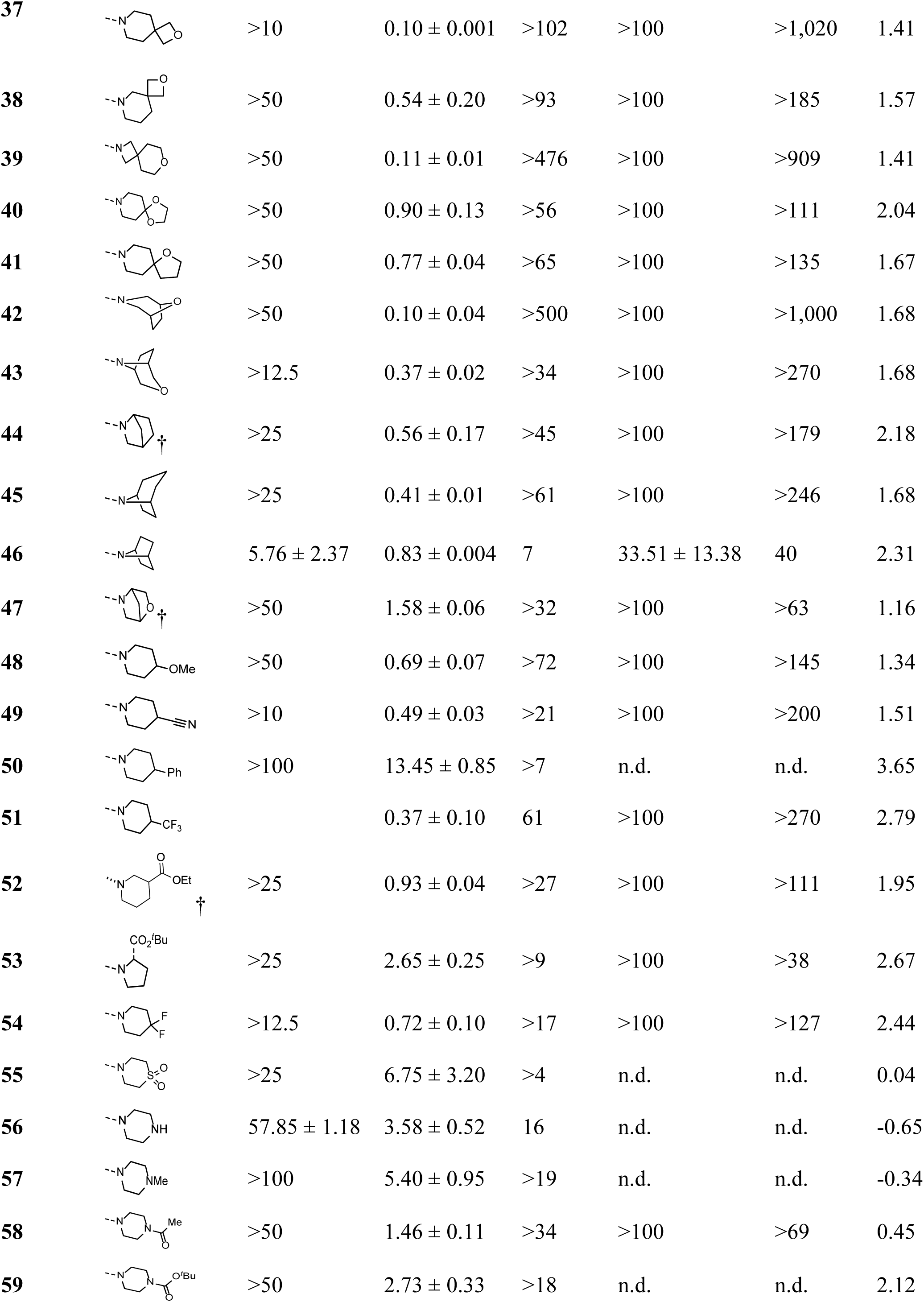
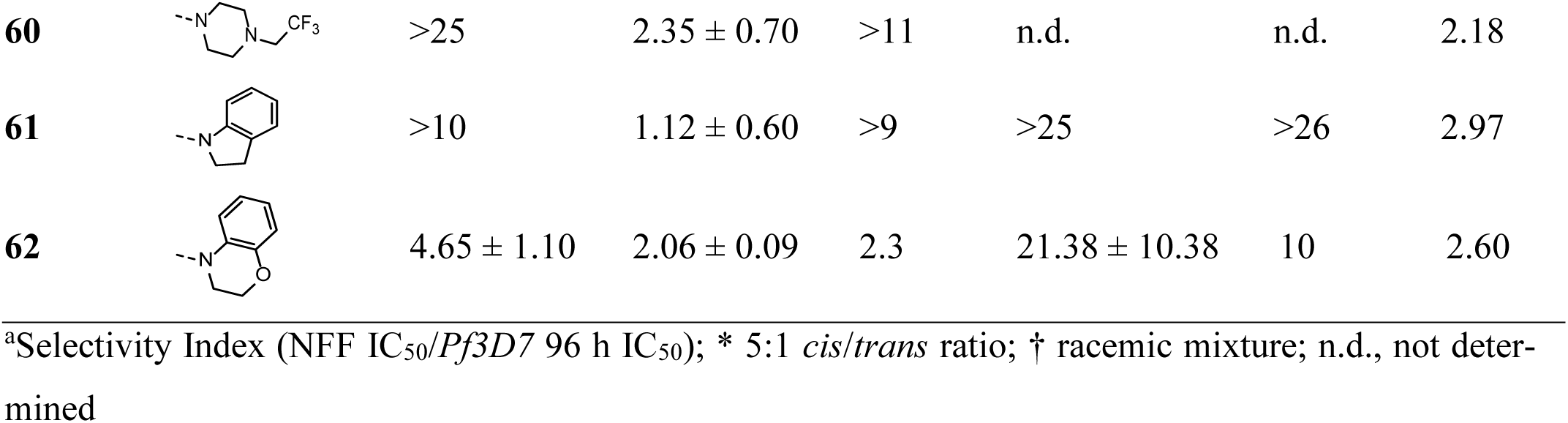
SAR data for *N,N*-dialkyl-5-cyclohexylsulfonyl-1,3,4-oxadiazol-2-amines with different 2-amino substituents.

Importantly, compounds with potency like, or better than, the screening hit **1** maintained a slow-action phenotype, as measured by their 48 h/96 h IC_50_ ratio (**Tables 4** and **5**). Likewise, when comparing the activity of compounds against *Pf*3D7 parasites versus human neonatal foreskin fibroblasts (NFF), Selectivity Indices (SI; NFF IC_50_/*Pf*3D7 96 h IC_50_) for compounds with a similar or better potency than lead compound **1** (SI >182) were generally similar or better (**Tables 4** and **5**).

### *In vitro* metabolism and permeability

The *in vitro* metabolic stability of the three initial slow-action hits (**1-3**; **Table 1**) and selected analogues was assessed using mouse and human liver microsomes. Compound **1** was found to possess greater stability towards human and mouse liver microsomes than **2** and **3** (**Table 6**). This could be attributed either to the lower lipophilicity of **1** and/or the presence of benzylic CH bonds in **2** and **3** as potential points for oxidative metabolism. Analogues **37** and **24** were selected for further studies based on their potency and selectivity for *P. falciparum* versus human cells (*Pf*3D7 96 h IC_50_ 0.10 µM and 0.09 µM, respectively; SI >1,000; **Table 5**) and relatively low clogD (1.41 and 2.67, respectively; **Table 5**). Both showed intermediate clearance in human liver microsomes and intermediate and high, respectively, clearance in mouse liver microsomes, suggesting that even small increases in lipophilicity relative to **1** (clogD 1.16) are detrimental to microsomal stability. The apparent permeability (P_app_) of **1**, **2**, **3** and **37** was assessed across confluent and differentiated Caco-2 cell monolayers in the apical to basolateral (A-B) direction. The apparent permeability (P_app_) values indicated high permeability across the Caco-2 cell monolayers for **1**, **2** and **37**. A P_app_ value for the most lipophilic compound **3** could not be determined due to low mass balance, presumably because of partitioning of the compound into the lipid bilayer.

### Chemical stability studies

5-Phenyl-2-sulfonyl-1,3,4-oxadiazoles are known to be susceptible to nucleophilic substitution by thiols.^23^ We therefore assessed the stability of lead compound **1** in the presence of methyl *N*-Boc cysteine in aqueous buffer/tetrahydrofuran (THF) and observed no degradation over 24 h. It was presumed that the electron donating 5-amino substituent in conjugation with the oxadiazolesulfonyl moiety would deactivate the sulfonyloxadiazole core towards nucleophilic attack. However, on storage at room temperature over several months, a solid sample of **1** partially degraded to unidentified, highly polar decomposition products. These lacked any UV chromophore, suggesting that hydrolysis of the oxadiazole core had occurred. Of note, it has recently been reported that related 2-amino-5-aryl-1,3,4-oxadiazoles can undergo hydrolysis to acyl hydrazides.^24^

### Pharmacokinetics

Due to the poor microsomal stability of **2**, **3**, **37** and **24**, we selected alternative compounds for subsequent pharmacokinetic assessment in mice alongside **1**. Compound **11** was chosen based on its low clogD, while **8** was chosen based on 3-fold improved potency (*Pf*3D7 96 h IC_50_ 0.18 µM; **Table 4**) and only slightly increased clogD relative to **1**. Compound **33** was selected based on ∼7-fold better potency than **1** (*Pf*3D7 96 h IC_50_ 0.08 µM; **Table 5**) and, offsetting its relatively high clogD, the possibility that the extra steric shielding around the oxadiazole core might provide greater hydrolytic and metabolic stability. Following oral administration at 100 mg/kg, maximum plasma concentrations of **1**, **11** and **8** were observed at 0.25 h and for **33** at 0.5 h suggesting rapid absorption (**Table 7**). Plasma concentrations remained measurable for up to 4 h post-dose for **11** and 7.5 h post-dose for **1**, **33** and **8** (**Table 7**). The elimination half-life of **1** and **8** could not be determined as the terminal phase of the profile was not well defined. Based on C_max_ and AUC values, **1** (**Figure 1A**) exhibited the highest exposure of the four compounds, followed by **11**, **8** and **33** (**Table 7**).

**Figure 1.**
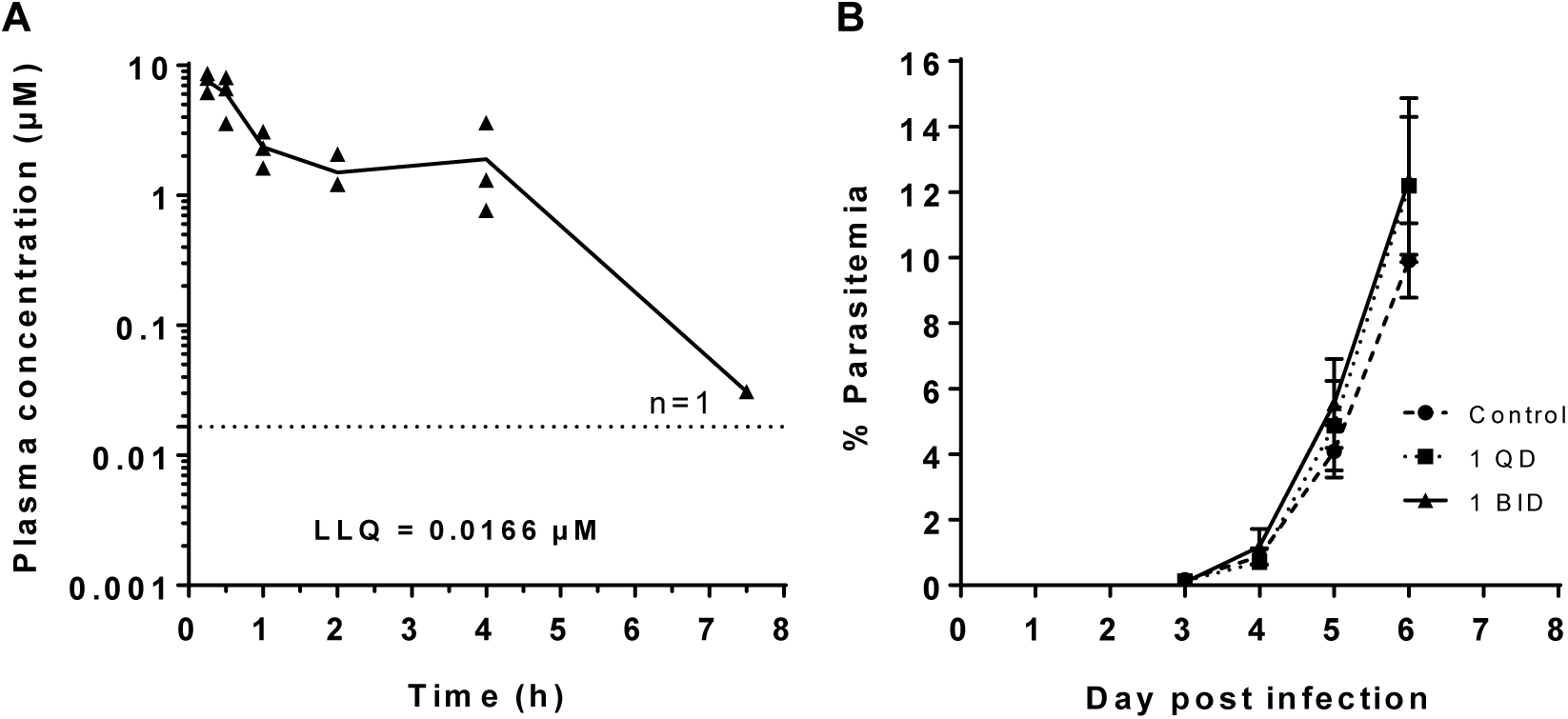
*In vivo* oral plasma concentrations of **1** and assessment in a murine malaria model. (**A**) Plasma concentrations of **1** were assessed in female BALB/c mice following oral administration of 100 mg/kg in 10% DMSO; 90% olive oil and sampling at 0.25, 0.5, 1, 2, 4, 7.5 and 24 h for three mice per timepoint. At 7.5 h, **1** was detected in only 1/3 mice (n=1) and was not detected at 24 h in all three mice. Lower limit of quantitation (LLQ) is the lowest acceptable calibration standard for which the back calculated concentration lies within ±20% of the nominal concentration. Calculated pharmacokinetic parameters are shown in Table 7. (**B**) *P. berghei* infected BALB/c mice (n=5 per group) were treated orally with 100 mg/kg **1** once daily (QD) or twice daily (BID) for three days starting 2 h post infection (p.i.). Groups of control mice were treated with vehicle (10% DMSO:90% olive oil) or once daily with 10 mg/kg chloroquine in PBS (not shown; no parasites detected). The mean % parasitemia (± SD) was monitored from day 3 post infection (p.i.) via microscopic examination of thin blood films prepared from peripheral blood.

**Table 6.** In vitro metabolism and permeability 1-3, 37 and 24.

| | clogD | Microsomal Stability | | | Caco-2 Permeability<br>A-B $P_{app}$ ($10^{-6}$ cm/s) |
| --- | --- | --- | --- | --- | --- |
| | | Species | $T_{1/2}$<br>(min) | Predicted $CL_{int, in vivo}$<br>(mL/min/kg) | |
| **1<sup>a</sup>** | 1.16 | Human | >255 | <6 | 86 |
|  |  | Mouse | 99 | 45 |  |
| **2** | 1.96 | Human | 5 | 260 | 39 |
|  |  | Mouse | 3 | 1,596 |  |
| **3<sup>b</sup>** | 3.49 | Human | 3 | 547 | c.n.d. |
|  |  | Mouse | <2 | >2,235 |  |
| **37** | 1.41 | Human | 112 | 13 | 44 |
|  |  | Mouse | 18 | 254 |  |
| **24** | 2.67 | Human | 68 | 21 | n.d. |
|  |  | Mouse | 7 | 658 |  |

c.n.d = could not be determined from data; n.d. = not determined

**Table 7.** Selected *N*,*N*-dialkyl-5-cyclohexylsulfonyl-1,3,4-oxadiazol-2-amines are rapidly absorbed and eliminated in mice.

| Parameter<sup>a</sup> | Compound |  |  |  |
| --- | --- | --- | --- | --- |
|  | **1** | **11** | **33** | **8** |
| Apparent $t_{1/2}$ (h) | c.n.c | 0.95<sup>c</sup> | 1.6 | c.n.c. |
| Plasma $C_{\max}$ ($\mu\text{M}$) | 7.61 | 2.14 | 0.31 | 0.65 |
| $T_{\max}$ (h) | 0.25 | 0.25 | 0.50 | 0.25 |
| Plasma $\text{AUC}_{0-t}$ ($\text{h} \cdot \mu\text{M}$)<sup>b</sup> | 11.5 | 3.61 | 0.39 | 1.16 |
| $\text{AUC}_{0-\text{inf}}$ ($\text{h} \cdot \mu\text{M}$) | c.n.c | 3.91<sup>c</sup> | 0.41 | c.n.c. |

<sup>a</sup>Pharmacokinetic studies were carried out in BALB/c mice following a single 100 mg/kg dose of **1**, **11**, **33** or **8** via gavage. Compounds were administered as uniform suspensions/emulsions. Plasma concentrations were assessed in samples collected at 0.25, 0.5, 1, 2, 4, 7.5 and 24 h (three mice per timepoint).
<sup>b</sup>AUC was calculated based on 0-t, where t is the last time point where measurable concentrations were detected (4 h for **11**; 7.5 h for **1** (one of three mice), **33** and **8**. No compounds were detected in any mice at 24 h. <sup>c</sup>Terminal elimination phase was estimated so value is an approximation; c.n.c. could not calculate.

### *In vivo* evaluation of compound toxicity and efficacy in mice

Compound **1**, with the best microsomal stability and pharmacokinetic profile of the compounds assessed, was selected for assessment of oral antiplasmodial efficacy *in vivo* in a *P. berghei* ANKA murine malaria model. First, the tolerability of **1** was assessed in uninfected mice (data not shown). No adverse clinical signs, except ruffled coats, were observed following single or twice daily oral doses for three days at concentrations up to 116 mg/kg. Liver and kidney pathology results were within normal parameters following two daily 116 mg/kg doses for three days. To assess the *in vivo* efficacy of **1** in the *P. berghei* ANKA murine malaria model, infected mice were treated orally with 100 mg/kg **1** either once daily or twice daily for three days. There was no significant difference in the peripheral blood parasitemia of treated mice when compared to the vehicle control group (**Figure 1B**). Chloroquine, included as antimalarial control drug, cured mice of infection (data not shown). It is likely that the poor *in vivo* antiplasmodial activity of **1** is due to its low compound exposure (**Figure 1A**; **Table 7**). While the calculated *in vitro* IC_90_ of **1** was 0.758 µM, this compound was only detected at this level (calculated average concentration (Cav) over 7.5 h of ∼1.5 µM) for 7.5 h in one of the three mice assessed. This would be well below the predicted exposure required for *in vivo* activity. Thus, significant optimization of potency and/or metabolism and other pharmacokinetic parameters for this chemotype would be required for improved *in vivo* efficacy. Determining the target(s) of these compounds may be helpful in identifying additional compounds with similar or better antiplasmodial potency and selectivity profiles but an improved pharmacokinetic profile. SciFinder searches found no reports of *N,N*-dialkyl-5-alkylsulfonyl-1,3,4-oxadiazol-2-amines in the literature, however compounds sharing the 1,3,4-oxadiazole core have been reported as inhibitors of *Plasmodium N*-myristoyltransferase (*Pf*NMT).^25^ An optimized compound (*Pf*NMT Ki = 8 nM) showed good activity against asexual stage *Plasmodium* parasites in a 48 h assay (EC_50_ = 302 nM), although slow action activity was not measured. Investigation of *Pf*NMT as a possible target could be a starting point for mode of action studies.

### Chemistry

The target compounds were prepared as shown in Schemes 1–5. The preparation of the intermediate thiotetrazoles **65a**–**p** is shown in Scheme 1. Alkyl halides or mesylates **63a**–**o** were converted to the respective alkylthiocyanates **64a**–**o** by reaction with potassium thiocyanate, while the aryldiazonium salt **63p** (formed by diazotization of 3,4-dichloroaniline **66**) was converted to aryl thiocyanate **64p** by copper-catalyzed reaction with potassium thiocyanate.^26^ Subsequent ZnBr_2_-catalyzed cycloaddition of sodium azide to thiocyanates **64a**–**p** gave the corresponding thiotetrazoles **65a**–**p**.^27^ The preparation of the target 1,3,4-oxadiazoles is summarized in Scheme 2. Reaction of selected tetrazole **65** with selected carbamoyl imidazole **68** (formed by reaction of secondary amine **67** with carbonyl diimidazole) afforded the corresponding 1,3,4-oxadiazoles **69** via tetrazole *N*-acylation and subsequent thermally-promoted loss of N_2_ and rearrangement (the Huisgen 1,3,4-oxadiazole synthesis).^28, 29^ Initially, experiments were performed using carbamoyl chlorides using reaction conditions developed by Detert for acid chlorides,^30^ however, the stable, crystalline carbamoyl imidazoles **68**^31^ proved to be more convenient reagents, although the reactions with the imidazoles required slightly higher temperatures/longer reaction times than the reactions with the chlorides. The target sulfones were obtained by *S*-oxidation of the respective thioethers **69** with trifluoroperoxyacetic acid, which was generated *in situ* using a modification of the method of Balicki.^32^ For those substrates containing acid sensitive functionality such as *N*-Boc groups, the oxidation reaction was buffered with sodium acetate. A number of target sulfones were prepared using *m*-chloroperbenzoic acid as oxidant. Sulfoxides were observed as intermediates during the oxidation reactions and in this way sulfoxide **5** and sulfone **2** were obtained from incomplete oxidation of thioether **4=69ba** (R^1^ = 2,5-dimethylphenylmethyl; R^2^ = morpholine) (Scheme 3). A number of targets were produced by synthetic modification of targets **17** and **59** (Scheme 4). Removal of the Boc-group of **17** and **59** using trifluoroacetic acid and triisopropylsilane afforded the respective amines **18** and **56**. Reductive methylation of **56** was achieved in one-pot with formaldehyde in formic acid^33^. Reductive alkylation of **56** with trifluoroacetic acid and phenylsilane^34^ afforded trifluoroethyl derivative **60**. The availability of a wide range of secondary amine and alkyl halide (or alcohol) starting materials enabled preparation of a broad variety of analogues (**Tables 4** and **5**). The 1,3,4-thiadiazole target **6** was prepared similarly as shown in Scheme 5. The reaction of tetrazole **65a** with 3-methyl-1-(morpholin-4-ylcarbonothioyl)-1H-imidazol-3-ium iodide **70** (prepared by reaction of morpholine with 1,1’-thiocarbonyldiimidazole and methyl iodide^31^) gave access to thio-1,3,4-thiadiazole **71**, which was then selectively oxidized to the corresponding sulfone **6** with *in situ* prepared trifluoroperacetic acid.

To our knowledge, this is the first report of *N,N*-dialkyl-5-alkylsulfonyl-1,3,4-oxadiazol-2-amines such as **1**–**3**, **7**–**17**, **19**–**55**, **58**–**59**, **61**–**62** and of the preparation of precursor 5-alkylthio-1,3,4-oxadiazol-2-amines such as **69** by this synthetic route. 5-Alkylthio-1,3,4-oxadiazol-2-amines have previously been prepared by *S*-alkylation^35^ or *S*-arylation^36^ of 1,3,4-oxadiazole-2(3*H*)-thiones.

**Scheme 1.**
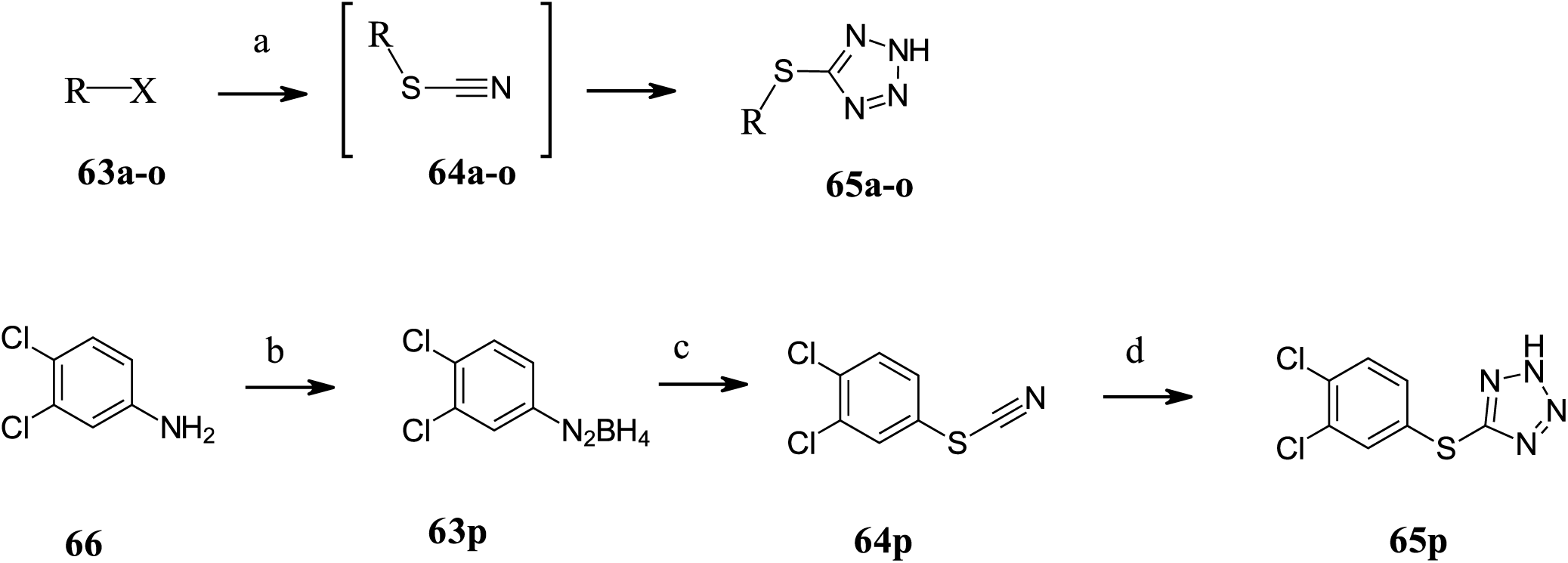
Synthesis of thiotetrazoles **65a**–**p**.*^a^* *^a^*Reagents and conditions: (a) **63a**–**o**, KSCN, DMF, 100 °C; NaN_3_, ZnBr_2_.2H_2_O, H_2_O/*t*-AmOH, reflux. (b) NaNO_2_, 25% HBF_4_, 0 °C. (c) KSCN, 0.1 equiv (MeCN)_4_CuBF_4_, Cu(BF_4_)_2_.H_2_O, TMEDA, MeCN, 0 °C. (d) NaN_3_, ZnBr_2_, H_2_O/*t*-AmOH, 100 °C.

**Scheme 2.**
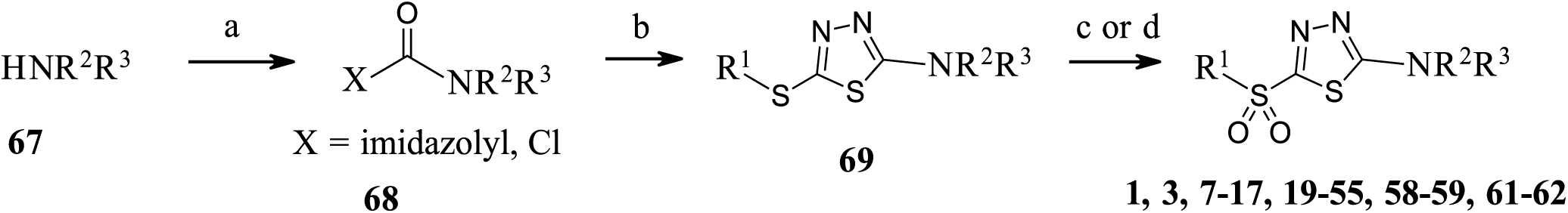
Synthesis of target 1,3,4-oxadiazoles. *^a^* *^a^*Reagents and conditions: (a) carbonyl diimidazole, THF (DMF for HCl salts), rt. (X = imidazolyl); triphosgene, DCM, sat’d NaHCO_3_, 0–rt. (X = Cl) (b) **65**, 2,4,6-collidine, PhOMe, 130 °C. (c) (CF_3_CO)_2_O, H_2_O_2_.urea, (NaOAc for acid sensitive substrates) MeCN, 0–rt. (d) *m*-CPBA, DCM, 0–rt.

**Scheme 3.**
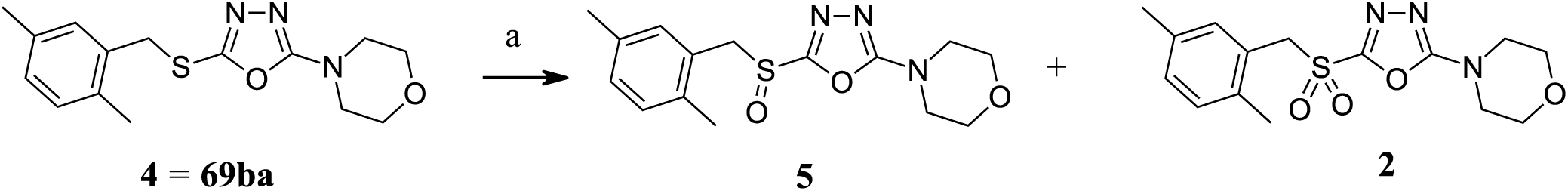
Preparation of target sulfoxide **5** and sulfone **2**.*^a^* *^a^*Reagents and conditions: (a) *m*-CPBA, DCM, rt.

**Scheme 4.**
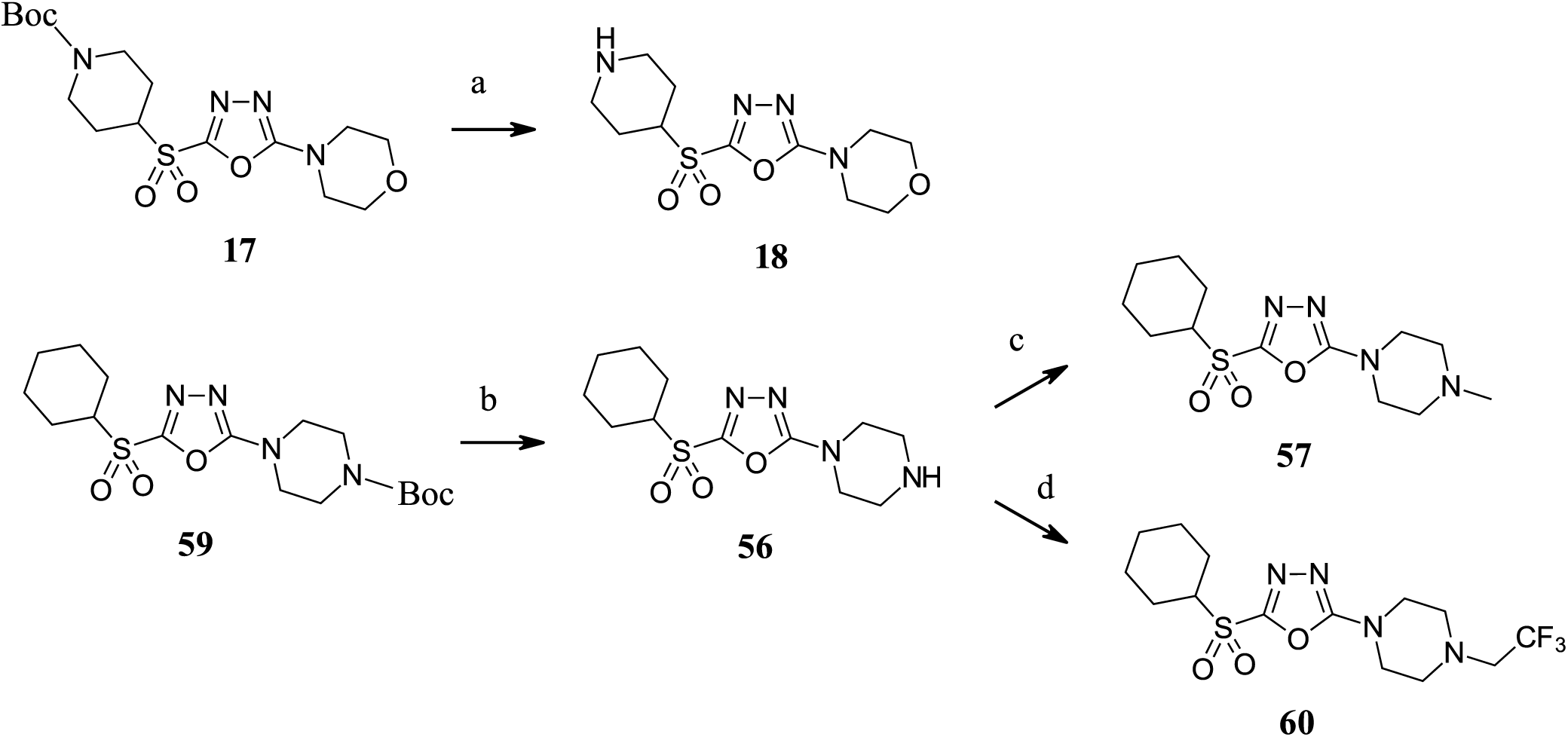
Preparation of targets **18**, **56**, **57** and **60**.*^a^* *^a^*Reagents and conditions: (a) CF_3_CO_2_H, i-Pr_3_SiH, H_2_O, rt. (b) 4M HCl in dioxane, DCM, rt. (c) 30% aq. CH_2_O, HCO_2_H, 80 °C. (d) PhSiH_3_, CF_3_CO_2_H, THF, 70 °C.

**Scheme 5:**
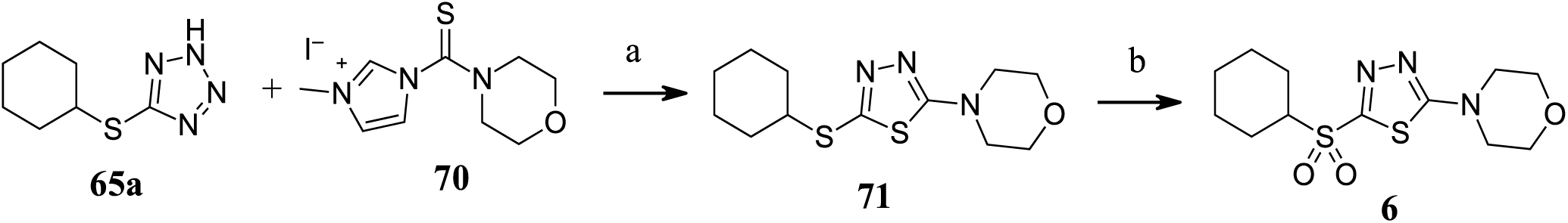
Synthesis of target 1,3,4-thiadiazole **6***^a^* *^a^*Reagents and conditions: (a) 2,4,6-collidine, PhOMe, 130 °C, 4 h. (b) CF_3_CO)_2_O, H_2_O_2_.urea, MeCN, 0– rt.

## CONCLUSIONS

This study identified *N*,*N*-dialkyl-5-alkylsulfonyl-1,3,4-oxadiazol-2-amines as a new series of antiplas-modial compounds with potent slow-action *in vitro* activity against drug sensitive and resistant *P. falciparum* malaria parasites with a different mode of action to current delayed-death antibiotics like clindamycin. SAR studies yielded interesting insights, with high potency and selectivity for *P. falciparum* versus human cells obtained for analogues such as **31** and **32** (96 h IC_50_ <40 nM; SI >2,500). While representative compound **1** demonstrated resistance to bionucleophile attack, it was found to degrade on storage under ambient conditions and representative compounds were rapidly eliminated from mice. This was reflected in the lack of activity observed in proof-of-concept *in vivo* efficacy studies with compound **1** in a murine model of malaria. The stability and pharmacokinetics of the compounds would need to be improved before further development. However, the mode-of-action of this new class of slow acting antiplasmodials is of significant interest and may aid in the development of new malaria chemoprophylactic agents.

## EXPERIMENTAL SECTION

### General Procedures

All solvents and chemicals were used as purchased without further purification. All anhydrous reactions were performed under a dry nitrogen atmosphere. All inorganic solutions are aqueous unless otherwise specified. The progress of all reactions was monitored using thin layer chromatography (TLC) performed on Merck pre-coated 0.25 mm silica F_254_ aluminium-backed plates (#5554) using standard solvent systems. Spots were visualized by irradiation with ultraviolet light (254 nm). Flash chromatography was performed using Merck (#9385, 230-400 mesh) silica gel 60 with the solvent system specified in the corresponding experiment. Melting points were determined on Büchi B-545 digital melting point apparatus and are uncorrected. Proton (^1^H) and carbon-13 (^13^C) NMR spectra were recorded on Bruker Avance 400 and 500 MHz NMR spectrometers with the sample held at 25±0.1°C using CDCl_3_, DMSO-*d*_6_, CD_3_CN or methanol-*d*_4_ as solvent. Chemical shifts are given in parts per million (ppm) and referenced using the Unified Scale relative to residual solvent (*δ*_H_ 7.24, 2.50, 1.94 and 3.31 for CDCl_3_, DMSO-*d*6, CD3CN and methanol-*d*_4_ respectively). ^13^C NMR spectra in CDCl_3_, DMSO-*d*_6_, CD_3_CN and methanol-*d*_4_ are referenced to the central resonance of the CDCl_3_ “triplet” (*δ*_C_ 77.23) and the DMSO-*d*6 (*δ*_C_ 39.51), CD_3_CN (*δ*C 1.39) and methanol-*d*_4_ (*δ*H 49.15) septets respectively. High resolution mass spectrometric (HRMS) analyses with electrospray ionization (ESI) were performed on a Thermo Scientific Q Exactive mass spectrometer fitted with a HESI-II ion source. Positive and/or negative ion electrospray mass spectra were recorded in an appropriate mass range set for 140,000 mass resolution. The probe was used with 0.3 ml/min flow of solvent. The nitrogen nebulizing/desolvation gas used for vaporization was heated to 350 °C in these experiments. The sheath gas flow rate was set to 35 and the auxiliary gas flow rate to 25 (both arbitrary units). The spray voltage was 3.0 kV and the capillary temperature was 300 °C.. Purity was determined by high performance liquid chromatography-mass spectrometry (LCMS). Purity of all final compounds was 95% or higher. The LCMS was performed on a Waters Acquity UPLC i-Class with QDa performance mass detector with adjustment-free atmospheric pressure ionisation (API) electrospray (ES) interface for reliability. Positive and negative ions were recorded simultaneously with full scan analysis in m/z range 50 to 1000. High purity nitrogen (>95%) nebulizing/desolvation gas is used for vaporization with the pressure regulated at 650 – 700 kPa. The probe temperature was set at 600 °C, the source temperature at 120 °C, the cone voltage was 15V whilst the capillary voltage was 1.5kV for positive and 0.8kV for negative ion. Spectral analysis was performed on a Waters PDA detector scanning from from 190 to 400 nm with chromatograms extracted using a wavelength of 254 nm. The chromatographic conditions are as follows: column: Acquity UPLC BEH C_18_ (50 x 2.1 mm, 1.7 µm particle size); mobile phase A: 100% Milli-Q water with 0.1% formic acid; mobile phase B: 100% acetonitrile with 0.1% formic acid; gradient: 95% A to 100% B over 4.50 min, 100% B for 1 min; flow rate: 0.400 mL/min; column temperature: 30 °C; sample injection volume: 1 µL.

### General Method for Synthesis of Thiotetrazoles 65

A stirred mixture of alkyl halide or mesylate **63** (1 equiv) and potassium thiocyanate (KSCN, 1.1 equiv) in dry dimethylformamide was heated at 100 °C for 18 h. The stirring was stopped, the mixture cooled and the precipitate was allowed to settle. The supernatant liquors were slowly transferred into a vigorously stirred mixture of sodium azide (1.1 equiv), zinc bromide dihydrate (1 equiv) in water. The retained precipitate from the first step was washed with *t*-amyl alcohol and the washings added to the sodium azide containing mixture, then the resultant mixture was heated at reflux for 7 h. After cooling, the reaction mixture was added to 2 M hydrochloric acid and extracted with diethyl ether. The organic layer was dried with anhydrous MgSO_4_, filtered, and evaporated under reduced pressure. The crude product was purified using flash chromatography with the solvent system and gradient determined using TLC.

*5-(Cyclohexylthio)-2H-tetrazole (**65a**)*. Prepared from bromocyclohexane (**63a**). Flash chromatography: ethyl acetate/dichloromethane/acetic acid (0:100:1→20:80:1). The product was further purified by trituration with diethyl ether/*heptane*. 3.06 g (46%) of a white solid; ^1^H NMR (400 MHz, CDCl_3_): *δ* = 3.86–3.78 (m, 1H), 2.16–2.03 (m, 2H), 1.80–1.70 (m, 2H), 1.65–1.46 (m, 3H), 1.45–1.20 (m, 3H); ^13^C NMR (100 MHz, CDCl_3_): *δ* = 154.8, 47.4, 33.6, 25.9, 25.6.

*5-(2,5-Xylyl)methylthio)-2H-tetrazole (**65b**).* Prepared from 2,5-xylylmethylchloride (**63b**). Flash chromatography: ethyl acetate/heptane (2:1). 0.163 g, 39% of a white solid. ^1^H NMR (400 MHz, CDCl_3_): *δ* =7.13–7.02 (m, 2H), 7.01–6.96 (m, 1H), 4.29 (s, 2H), 3.77–3.73 (m, 4H), 3.45–3.41 (m, 4H), 2.34 (s, 3H), 2.26 (s, 3H). ^13^C NMR (100 MHz, CDCl_3_): *δ* = 165.3, 156.8, 136.0, 134.1, 133.4, 131.0, 130.8, 129.3, 66.1, 46.3, 36.1, 21.0, 18.9.

*5-(3,4-Dichlorophenylmethylthio)-2H-tetrazole (**65c**).* Prepared from 1,2-dichloro-4-(chloromethyl)benzene (**63c**). The first operation was performed in refluxing *t*-amyl alcohol for 3 h. 3.33 g (89%) of a white solid; ^1^H NMR (400 MHz, DMSO-*d*_6_): *δ* = 7.69 (d, *J* = 2.1 Hz, 1H), 7.58 (d, *J* = 8.3 Hz, 1H), 7.39 (dd, *J* = 8.3, 2.1 Hz, 1H), 4.49 (s, 2H).

*5-(Cyclopentylthio)-2H-tetrazole (**65d**).* Prepared from cyclopentyl bromide (**63d**). Crude product purified by trituration with diethyl ether. 0.86 g (15%) of a white solid; ^1^H NMR (400 MHz, CDCl_3_): *δ* = 4.13–4.05 (m, 1H), 2.26–2.15 (m, 2H), 1.84–1.58 (m, 6H); ^13^C NMR (100 MHz): *δ* = 155.8, 46.4, 34.0, 24.8.

*5-(Cycloheptylthio)-2H-tetrazole (**65e**)*. Prepared from cycloheptylbromide (**63e**). Flash chromatography: ethyl acetate/heptane/acetic acid (20:80:1→40:60:1). 2.00 g (50%) of a white solid; ^1^H NMR (500 MHz, CDCl_3_): *δ* = 12.52 (brs, 1H), 4.03–3.97 (m, 1H), 2.19–2.12 (m, 2H), 1.81–1.50 (m, 10H).

*5-(Cyclohexylmethylthio)-2H-tetrazole (**65f**)*. Prepared from cyclohexylmethylbromide (**63f**). The first operation was performed in refluxing *t*-amyl alcohol for 16 h. Flash chromatography: ethyl acetate/dichloromethane/acetic acid (0:100:1→20:80:1). 1.86 g (65%) of a white solid; ^1^H NMR (400 MHz, CDCl_3_): *δ* = 8.00 (brs, 1H), 3.19 (d, *J* = 6.91 Hz, 2H), 1.86–1.78 (m, 2H), 1.74–1.58 (m, 5H), 1.24–0.88 (m, 6H).

*5-(Cyclopentylmethylthio)-2H-tetrazole (**65g**)*. Prepared from cyclopentylmethylmesylate (**63g**). Flash chromatography: ethyl acetate/heptane/acetic acid (20:80:1→40:60:1). 0.40 g (39%) of a white solid; ^1^H NMR (500 MHz, CDCl_3_): *δ* = 14.05 (brs, 1H), 3.33 (d, *J* = 7.4 Hz, 2H), 2.24 (apparent septet, *J* = 7.7 Hz, 1H), 1.88–1.80 (m, 2H), 1.68–1.50 (m, 4H), 1.31–1.22 (m, 2H).

*5-(Cyclobutylmethylthio)-2H-tetrazole (**65h**)*. Prepared from cyclobutylmesylate (**63h**).^37^ (Note: Cyclobutylmethylmesylate is an explosion hazard^38^) The crude product was purified by trituration with diethyl ether/heptane. 1.20 g, 100% of a white solid; ^1^H NMR (400 MHz, CDCl_3_): *δ* = 3.35 (d, *J* = 7.6 Hz, 1H), 2.70–2.59 (m, 1H), 2.14–2.05 (m, 2H), 1.90–1.65 (m, 4H).

*5-(Neopentylthio)-2H-tetrazole (**65i**)*. Prepared from 1-bromo-2,2-dimethylpropane (**63i**). The product was purified by trituration with diethyl ether/heptane. 5.13 g (91%) of white semi-solid; ^1^H NMR (400 MHz, CDCl_3_): *δ* = 3.26 (s, 2H), 1.00 (9H).

*exo-2-Norbornanylthio-2H-tetrazole (**65j**)*. A mixture of *endo*-2-(mesyloxy)norbornane (**63j**).^39^ (0.194 g, 1.02 mmol), KSCN (0.109 g, 1.12 mmol) and dimethylformamide (2 mL) was heated to 100 °C for 18 h. The mixture was cooled, then partitioned between diethyl ether and water. The organic phase was washed with brine, dried (MgSO_4_) and evaporated to give a mixture of the desired thiocyanate and an isothiocyanate side-product, which were separated by flash chromatography (dichloromethane/heptane, 25:75→50:50). The higher R*f* fraction gave *exo*-2-isothiocyanatonorbornane (0.036 g, 23%). The lower R*f* faction gave *exo-2-thiocyanatobicyclo[2.2.1]heptane* (**64j**).^40^ 0.061 g (39%); ^1^H NMR (400 MHz, CDCl_3_): *δ* = 3.39–3.34 (m, 1H), 2.49–2.46 (m, 1H), 2.39–2.35 (m, 1H), 1.85–1.78 (m, 1H), 1.74–1.61 (m, 2H), 1.58–1.48 (m, 1H), 1.41–1.33 (m, 1H), 1.32–1.13 (m, 3H). The e*xo-2-thiocyanatobicyclo[2.2.1]heptane* (**64j**) (102 mg, 0.666 mmol) was reacted with sodium azide according to the above General Synthetic Method. Flash chromatography: ethyl acetate/heptane/acetic acid, 20:80:1→40:60:2. 0.063 g (48%) as a white solid; ^1^H NMR (400 MHz, CDCl_3_): *δ* = 3.75–3.70 (m, 1H), 2.43–2.40 (m, 1H), 2.35–2.31 (m, 1H), 1.98–1.90 (m, 1H), 1.65–1.56 (m, 2H), 1.55–1.40 (m, 2H), 1.32–1.14 (m, 3H).

*5-[(1-Adamantanyl)methylthio]-2H-tetraazole (**65k**)*. Prepared from 1-[(mesyloxy)methyl]adamantane (**63k**).^41^ Flash chromatography: ethyl acetate/heptane/acetic acid, 20:80:1→40:60:1 0.250 mg (24%) of a white solid; ^1^H NMR (400 MHz, CDCl_3_): *δ* = 3.22 (s, 2H), 2.01–1.92 (m, 3H), 1.72–1.53 (m, 12H).

*5-(((1-(Trifluoromethyl)cyclopropyl)methyl)thio)-2H-tetrazole (**65l**)*. Prepared from (1-(trifluoromethyl)cyclopropyl)methyl methanesulfonate (**63l**).^42^ The crude product was purified by triturating with diethyl ether/heptane. 0.633 g (44%) of an off-white solid; ^1^H NMR (400 MHz, methanol-*d*_4_): *δ* = 3.67 (s, 2H), 1.16–1.11 (m, 2H), 1.03–0.98 (m, 2H).

*5-(((1-(Trifluoromethyl)cyclobutyl)methyl)thio)-2H-tetrazole (**65m**)*. Prepared from (1-(trifluoromethyl)cyclobutyl)methyl methanesulfonate (**63m**).^43^ Flash chromatography: ethyl acetate/dichloromethane/acetic acid 5:94:1→20:79:1. 0.350 g (26%) of a white solid; ^1^H NMR (400 MHz, methanol-*d*_4_): *δ* = 3.77 (s, 2H), 2.42–2.32 (m, 2H), 2.19–1.95 (m, 4H).

*tert-Butyl 4-((2H-tetrazol-5-yl)thio)piperidine-1-carboxylate (**65n**)*. Prepared from *tert*-Butyl 4-(mesyloxy)-1-piperidinecarboxylate (**63n**).^43^ Flash chromatography: ethyl acetate/heptane/acetic acid, 40:60:1→0:100:1. 86 mg (30%) of a colorless oil; ^1^H NMR (400 MHz, CDCl_3_): *δ* = 4.04–3.90 (m, 3H), 3.12–2.98 (m, 2H), 2.19–2.06 (m, 2H), 1.73–1.60 (m, 2H), 1.46 (s, 9H).

*5-((Tetrahydro-2H-pyran-4-yl)thio)-2H-tetrazole (**65o**)*. Prepared from 4-bromotetrahydrofuran (**63o**). Not further purified. 0.520 g (92%) of a white solid; ^1^H NMR (400 MHz, CDCl_3_): *δ* = 4.06–3.97 (m, 3H), 3.60–3.52 (m, 2H), 2.18–2.10 (m, 2H), 1.89–1.77 (m, 2H).

### General Method for Synthesis of Carbamoyl Imidazoles 68

The secondary amine or amine hydrochloride salt (1 equiv) was dissolved in anhydrous tetrahydrofuran (THF) (2 mL per mmol amine) (dimethylformamide (DMF) for hydrochloride or hemi-oxalate salts), 1,1’-carbonyldiimidazole (1.2 equiv) was added and then the resulting mixture was stirred at room temperature for 18 h. After evaporation under reduced pressure, the residue was dissolved in dichloromethane and the organic layer was washed with water, brine, dried over MgSO_4_, filtered and evaporated. Unless otherwise noted, the product was used without further purification in the next step.

*(1-Imidazolyl)morpholinomethanone (**68a**)*. 0.83 g (87%) of a white solid; ^1^H NMR (400 MHz, CDCl_3_): *δ* = 7.86–7.84 (m, 1H), 7.18–7.17 (m, 1H), 7.09–7.08 (m, 1H), 3.76–3.72 (m, 4H), 3.63–3.59 (m, 4H).

*(1-Imidazolyl)(4-methyl-1-piperidyl)methanone (**68b**)*. Dimethylformamide was used as solvent in place of tetrahydrofuran. 0.791 g (97%) of a colorless oil; ^1^H NMR (400 MHz, CDCl_3_): *δ* = 7.83–7.82 (m, 1H), 7.17–7.16 (m, 1H), 7.06-7.05 (m, 1H), 4.10–4.00 (m, 2H), 3.03–2.95 (m, 2H), 1.79–1.59 (m, 3H), 1.27–1.15 (m, 2H), 0.98 (d, *J* = 6.5 Hz, 3H).

*(1-Imidazolyl)(1-pyrrolidinyl)methanone (**68c**)*. 0.460 g (81%) of a white solid; ^1^H NMR (400 MHz, CDCl_3_): *δ* = 7.99–7.98 (m, 1H), 7.33–7.31 (m, 1H), 7.04–7.03 (m, 1H), 3.64–3.55 (m, 4H), 2.01–1.90 (m, 4H).

*(1H-Imidazol-1-yl)(piperidin-1-yl)methanone (**68d**)*. 0.460 g (70%) of white crystals; ^1^H NMR (500 MHz, CDCl_3_): *δ* = 7.83–7.82 (m, 1H), 7.17–7.16 (m, 1H), 7.07–7.06 (m, 1H), 3.54–3.50 (m, 4H), 1.73–1.61 (m, 6H).

*Azepan-1-yl(1H-imidazol-1-yl)methanone (**68e**)*. 0.324 g (74%) of pale yellow crystals; ^1^H NMR (400 MHz, CDCl_3_): *δ* = 7.85 (s, 1H), 7.21 (t, *J* = 1.3 Hz, 1H), 7.05 (s, 1H), 3.55–3.49 (m, 4H), 1.83–1.74 (m, 4H), 1.64–1.56 (m, 4H).

*(1-Imidazolyl)(1,4-oxazepan-4-yl)methanone (**68f**)*. 0.327 g (80%) of a colorless oil; ^1^H NMR (400 MHz, CDCl_3_): *δ* = 7.88–7.86 (m, 1H), 7.22–7.20 (m, 1H), 7.08–7.07 (m, 1H), 3.84–3.76 (m, 4H), 3.72–3.66 (m, 4H), 2.04–1.97 (m, 2H).

*(1-Imidazolyl)(1,2-oxazinan-2-yl)methanone (**68g**)*. 0.364 g (77%) of a colorless oil which solidified on standing; ^1^H NMR (400 MHz, CDCl_3_): *δ* = 8.23–8.22 (m, 1H), 7.54–7.52 (m, 1H), 7.02–7.01 (m, 1H), 4.00–3.96 (m, 2H), 3.89–3.84 (m 2H), 1.92–1.81 (m, 4H).

*(1H-Imidazol-1-yl)(3-methylpiperidin-1-yl)methanone (**68h**)*. 0.300 g (77%) of a yellow oil; ^1^H NMR (400 MHz, CDCl_3_): *δ* = 7.84–7.82 (m, 1H), 7.19–7.15 (m, 1H), 7.08–7.05 (m, 1H), 4.02–3.90 (m, 2H), 3.01–2.93 (m, 1H), 2.65 (dd, *J* = 13.1, 10.9 Hz, 1H), 1.92–1.84 (m, 1H), 1.80–1.64 (2H), 1.63–1.50 (m, 1H), 1.24–1.12 (m, 1H), 0.91 (d, *J* = 6.6 Hz, 3H).

*(1H-Imidazol-1-yl)(4,4-dimethylpiperidin-1-yl)methanone (**68j**)*. Prepared from 4,4-dimethylpiperidine hydrochloride. 0.140 g (40%) of a colorless oil; ^1^H NMR (500 MHz, CDCl_3_): *δ* = 7.84–7.82 (m, 1H), 7.18–7.16 (m, 1H), 7.07–7.06 (m, 1H), 3.55–3.51 (m, 4H), 1.46–1.42 (m, 4H), 1.01 (s, 6H).

*(3,3-Dimethyl-1-piperidyl)(1-imidazolyl)methanone (**68l**)*. 0.200 g (42%) of a colorless oil; ^1^H NMR (400 MHz, CDCl_3_): *δ* = 7.83–7.82 (m, 1H), 7.19–7.16 (m, 1H), 7.08–7.06 (m, 1H), 3.50–3.45 (m, 2H), 3.21 (s, 2H), 1.73–1.65 (m, 2H), 1.48–1.44 (m, 2H), 0.93 (s, 6H).

*(±)-(1-Imidazolyl)(3-methyl-1-piperidyl)methanone (**68n**)*. 0.367 g (83%) as a yellow oil; ^1^H NMR (400 MHz, CDCl_3_): *δ* = 7.85 (s, 1H), 7.18–7.17 (m, 1H), 7.10–7.08 (m, 1H), 3.94–3.89 (m. 3H), 3.65–3.57 (m, 2H), 3.27–3.20 (m, 1H), 2.88 (dd, *J* = 13.3, 10.5 Hz, 1H), 1.19 (d, *J* = 6.2 Hz, 3H).

*(2,2-Dimethyl-4-morpholinyl)(1-imidazolyl)methanone (**68o**)*. 0.352 g (93%) of a colorless oil; ^1^H NMR (400 MHz, CDCl_3_): *δ* = 7.86–7.85 (m, 1H), 7.19–7.17 (m, 1H), 7.11–7.09 (m, 1H), 3.81–3.78 (m, 2H), 3.59–3.55 (m, 2H), 3.38 (br s, 2H), 1.24 (s, 6H).

cis*-2,6-Dimethyl-4-morpholinyl](1-imidazolyl)methanone (**68p**)*. 0.450 g (90%) of a white solid; ^1^H NMR (400 MHz, CDCl_3_): *δ* = 7.86–7.84 (m, 1H), 7.19–7.16 (m, 1H), 7.11–7.08 (m, 1H), 3.81–3.77 (m, 2H), 3.59–3.55 (m, 2H), 3.58 (s, 2H), 1.24 (d, *J* = 6.2 Hz, 6H).

(1H*-Imidazol-1-yl)(2-oxa-7-azaspiro[3.5]nonan-7-yl)methanone (**68q**)*. Prepared from 2-oxa-7-azaspiro[3.5]nonane hemi-oxalate. 0.385 g (98%) of a colorless oil that solidified on standing; ^1^H NMR (400 MHz, CDCl_3_): *δ* = 7.83–7.81 (m, 1H), 7.16–7.14 (m, 1H), 7.09–7.06 (m, 1H), 4.46 (s, 4H), 3.51–3.46 (m, 4H), 1.98–1.93 (m, 4H).

*(1-Imidazolyl)(2-oxa-6-aza-6-spiro[3.5]nonyl)methanone (**68r**)*. Prepared from 2-oxa-6-azaspiro[3.5]nonane hydrochloride. 0.290 g (85%) as a yellow oil; ^1^H NMR (400 MHz, CDCl_3_): *δ* = 7.84–7.83 (m, 1H), 7.18–7.16 (m, 1H), 7.10–7.09 (m, 1H), 4.40–4.35 (m, 4H), 3.79 (br s, 2H), 3.48–3.44 (m, 2H), 1.98–1.93 (m, 2H), 1.65–1.58 (m, 2H).

*(1-Imidazolyl)(7-oxa-2-aza-2-spiro[3.5]nonyl)methanone (**68s**)*. 0.180 g (41%) of a colorless oil; ^1^H NMR (500 MHz, CDCl_3_): *δ* = 7.98–7.97 (m, 1H), 7.29–7.27 (m, 1H), 7.07–7.06 (m, 1H), 4.03 (s, 4H), 3.64–3.60 (m, 4H), 1.84–1.80 (m, 4H).

*(1,4-Dioxa-8λ^4^-aza-8-spiro[4.5]decyl)(1-imidazolyl)methanone (**68t**)*. 0.750 g (90%) of a white solid; ^1^H NMR (500 MHz, CDCl_3_): *δ* = 7.84–7.83 (m, 1H), 7.18–7.16 (m, 1H), 7.08–7.07 (m, 1H), 3.97 (s, 4H), 3.68–3.64 (m, 4H), 1.79–1.75 (m, 4H).

*(1-Imidazolyl)(1-oxa-8λ^4^-aza-8-spiro[4.5]decyl)methanone (**68u**)*. Colorless oil; ^1^H NMR (500 MHz, CDCl_3_): *δ* = 7.84–7.82 (m, 1H), 7.18–7.16 (m, 1H), 7.07–7.06 (m, 1H), 3.86–3.77 (m, 4H), 3.50–3.43 (m, 2H), 1.98–1.91 (m, 2H), 1.75–1.60 (m, 6H).

*(1-Imidazolyl)(8-oxa-3-azabicyclo[3.2.1]oct-3-yl)methanone (**68v**)*. 0.263 g (67%) of colorless crystals; ^1^H NMR (400 MHz, CDCl_3_): *δ* = 7.85-7.84 (m, 1H), 7.17-7.16 (m, 1H), 7.08-7.07 (m, 1H), 4.40-4.36 (m, 2H), 3.85-3.70 (m, 2H), 3.42 (dd, *J* = 12.8, 1.7 Hz, 2H), 2.04-1.95 (m, 2H), 1.86-1.78 (m, 2H).

*(1-Imidazolyl)(3-oxa-8-azabicyclo[3.2.1]oct-8-yl)methanone (**68w**)*. Prepared from 3-oxa-8-azabicyclo[3.2.1]octane hydrochloride. 0.164 g (79%) of a white solid; ^1^H NMR (400 MHz, CDCl_3_): *δ* = 7.93–7.89 (m, 1H), 7.24–7.21 (m, 1H), 7.09–7.06 (m, 1H), 4.32–4.27 (m, 2H), 3.80 (br d, *J* = 11.3 Hz, 2H), 3.67 (br d, *J* = 11.3 Hz, 2H), 2.10–1.95 (m, 4H).

*(2-Azabicyclo[2.2.1]hept-2-yl)(1-imidazolyl)methanone (**68x**)*. 0.250 g (49%) of colorless oil; ^1^H NMR (400 MHz, CDCl_3_): *δ* = 7.97–7.95 (m, 1H), 7.30–7.28 (m, 1H), 7.06–7.04 (m, 1H), 4.42 (br s, 1H), 3.64–3.59 (m, 1H), 3.20–3.15 (m, 1H), 2.69–2.65 (m, 1H), 1.95–1.68 (m, 4H), 1.60–1.42 (m, 2H).

*(8-Azabicyclo[3.2.1]oct-8-yl)(1-imidazolyl)methanone (**68y**)*. 0.170 g (81%) as a light brown oil; ^1^H NMR (400 MHz, CDCl_3_): *δ* = 7.92–7.89 (m, 1H), 7.25–7.23 (m, 1H), 7.06–7.04 (m, 1H), 4.42 (br s, 2H), 2.07–1.96 (m, 2H), 1.90–1.55 (m, 8H).

*(7-Azabicyclo[2.2.1]hept-7-yl)(1-imidazolyl)methanone (**68z**)*. Prepared from 7-azabicyclo[2.2.1]heptane hydrochloride. 0.200 g (70%) of a colorless solid; ^1^H NMR (400 MHz, CDCl_3_): *δ* = 7.94 (s, 1H), 7.27–7.25 (m, 1H), 7.06–7.04 (m, 1H), 4.43–4.39 (m, 2H), 1.94–1.87 (m, 4H), 1.58–1.50 (m, 4H).

*(±)-(1-Imidazolyl)(2-oxa-5-azabicyclo[2.2.1]hept-5-yl)methanone (**68a’**)*. Prepared from (±)-2-oxa-5-azabicyclo[2.2.1]heptane hydrochloride. 0.128 g (60%) of a white solid; ^1^H NMR (400 MHz, CDCl_3_): *δ* = 7.94 (s, 1H), 7.26–7.23 (m, 1H), 7.04–7.02 (m, 1H), 4.73 (br s, 1H), 4.64 (br s, 1H), 4.03 (d, *J* = 8.0 Hz, 1H), 3.83 (dd, *J* = 8.0, 1.6 Hz, 1H), 3.62–3.53 (m, 2H), 1.98–1.90 (m, 2H).

(1H*-Imidazol-1-yl)(4-methoxypiperidin-1-yl)methanone (**68b’**)*. 0.280 g (∼100%) as a colorless oil; ^1^H NMR (400 MHz, CDCl_3_): *δ* = 7.82 (s, 1H), 7.15 (t, *J* = 1.3 Hz, 1H), 7.05 (s, 1H), 3.74–3.67 (m, 2H), –3.41 (m, 3H), 3.34 (s, 3H), 1.91–1.82 (m, 2H), 1.74–1.65 (m, 2H).

*1-(1H-Imidazole-1-carbonyl)piperidine-4-carbonitrile (**68c’**)*. 0.338 g (93%) as a colorless oil; ^1^H NMR (500 MHz, CDCl_3_): *δ* = 7.83–7.81 (m, 1H), 7.15–7.14 (m, 1H), 7.08–7.06 (m, 1H), 3.76–3.69 (m, 2H), 3.62–3.56 (m, 2H), 2.98–2.93 (m, 1H), 2.01–1.89 (m, 4H).

*(1-Imidazolyl)[4-(trifluoromethyl)-1-piperidyl]methanone (**68e’**)*. 0.360 g (87%) as a pale brown oil that solidified on standing; ^1^H NMR (400 MHz, CDCl_3_): *δ* = 7.86–7.84 (m, 1H), 7.19–7.16 (m, 1H), 7.10–7.09 (m, 1H), 4.25–4.17 (m, 2H), 3.08–2.99 (m, 2H), 2.40–2.26 (m, 1H), 2.02–1.95 (m, 2H), 1.70–1.58 (m, 2H).

tert*-Butyl 1-[(1-imidazolyl)carbonyl]-2-pyrrolidinecarboxylate (**68g’**)*. 0.249 g (80%) as a pale yellow oil; ^1^H NMR (500 MHz, CDCl_3_): *δ* = 8.01–7.95 (m, 1H), 7.34–7.29 (m, 1H), 7.05–7.04 (m, 1H), 4.50–4.45 (m, 1H), 3.78–3.66 (m, 2H), 2.36–2.28 (m, 1H), 2.10–1.90 (m, 3H), 1.43 (s, 9H).

*(4,4-Difluoro-1-piperidyl)(1-imidazolyl)methanone (**68h’**)*. 0.200 g (45%) as a colorless solid; ^1^H NMR (400 MHz, CDCl_3_): *δ* = 7.87–7.85 (m, 1H), 7.19–7.17 (m, 1H), 7.11–7.09 (m, 1H), 3.73–3.69 (m, 4H), 2.13–2.02 (m, 4H).

*4-[(1-Imidazolyl)carbonyl]-1λ^6^,4-thiazinane-1,1-dione (**68i’**)*. The white precipitate which appeared when the crude product was partitioned between dichloromethane and water was collected by filtration.

0.106 g (41%) as a white solid; ^1^H NMR (400 MHz, (DMSO-*d*_6_): *δ* = 8.07–8.04 (m, 1H), 7.52–7.50 (m, 1H), 7.06–7.05 (m, 1H), 3.90–3.83 (m, 4H), 3.37–3.33 (m, 4H).

*1-(4-(1H-Imidazole-1-carbonyl)piperazin-1-yl)ethan-1-one (**68j’**)*. Due to the water solubility of product, the aqueous workup was omitted. Evaporation of the tetrahydrofuran afforded a mixture of imidazole and title compound. 0.352 g (∼100%) as a colorless oil; ^1^H NMR (400 MHz, CDCl_3_): *δ* = 7.87–7.86 (m, 1H), 7.18–7.17 (m, 1H), 7.09–7.08 (m, 1H), 3.67–3.45 (m, 8H), 2.11 (s, 3H).

tert*-Butyl 4-(1*H*-imidazole-1-carbonyl)piperazine-1-carboxylate (**68k’**)*. 1.36 g (90%) as white crystals; ^1^H NMR (400 MHz, CDCl_3_): *δ* = 7.86 (s, 1H), 7.18 (t, *J* = 1.4 Hz, 1H), 7.09 (s, 1H), 3.59–3.55 (s, 4H), –3.49 (s, 4H), 1.46 (s, 9H).

*(1*H*-Imidazol-1-yl)(indolin-1-yl)methanone (**68l’**)*. 0.358 g (94%) of pale pink solid; ^1^H NMR (400 MHz, CDCl_3_): *δ* = 8.00 (s, 1H), 7.34–7.32 (m, 1H), 7.25–7.16 (m, 3H), 7.13–7.11 (m, 1H), 7.08 (dd, *J* = 7.5 Hz, *J* = 1.0 Hz), 4.21–4.16 (m, 2H), 3.18 (t, *J* = 8.1 Hz, 2H).

*(2,3-Dihydro-4*H*-benzo[*b*][1,4]oxazin-4-yl)(1*H*-imidazol-1-yl)methanone (**68m’**)*. The crude product was purified by flash chromatography (ethyl acetate). 0.201 g (64%) as a white solid; ^1^H NMR (400 MHz, CDCl_3_): *δ* = 7.86 (s, 1H), 7.15 (t, *J* = 1.4 Hz, 1H), 7.06–6.99 (m, 2H), 6.92 (d, *J* = 8.1 Hz, 2H), 6.78–6.72 (m, 2H), 4.41–4.37 (m, 2H), 3.97–3.94 (m, 2H).

### General Method for Synthesis of Carbamoyl Chlorides 68

Triphosgene (0.40 equiv) was added to an ice-cooled, rapidly stirred mixture of secondary amine (1 equiv), dichloromethane (DCM) (5 mL per mmol amine) and sat’d NaHCO_3_ (5 mL per mmol amin). Stirring was continued at 0 °C for 30 min then the ice bath was removed and stirring was continued at room temperature for 2 h. The organic layer was separated, washed with brine, dried (MgSO_4_), filtered and evaporated to give the carbamoyl chloride. The carbamoyl chlorides thus generated were used immediately in subsequent reactions.

*2-Methyl-1-piperidinecarbonyl chloride (**68i**)*. 0.086 g (53%) of a pale yellow solid; ^1^H NMR (400 MHz, CDCl_3_): *δ* = 4.63–4.55 (m, 1H), 4.18–4.10 (m, 1H), 3.26–2.78 (m, 1H), 1.76–1.36 (m, 6H), 1.22 (d, *J* = 7.0 Hz, 3H).

cis*-3,5-Dimethyl-1-piperidinecarbonyl chloride (**68k**)*. 0.102 g (66%) of a colorless oil; ^1^H NMR (400 MHz, CDCl_3_): *δ* = 4.27–4.16 (m, 2H), 2.54–2.45 (m, 1H), 2.35–2.26 (m, 1H), 1.88–1.78 m, 1H), 1.73–1.58 (m, 2H), 0.92–0.86 (m, 6H), 0.80–0.70 (m, 1H) (major (*cis*) isomer).

cis*-2,6-Dimethyl-1-piperidinecarbonyl chloride (**68m**)*. 0.288 g (25%) of a pale yellow oil; ^1^H NMR (400 MHz, CDCl_3_): *δ* = 4.52–4.44 (m, 2H), 1.82–1.47 (m, 6H), 1.28 (d, *J* = 7.1 Hz, 6H).

*4-Phenyl-1-piperidinecarbonyl chloride (**68d’**)*. 0.118 g (85%) of a pale yellow solid; ^1^H NMR (400 MHz, CDCl_3_): *δ* = 7.34–7.28 (m, 2H), 7.24–7.16 (m, 3H), 4.51–4.41 (m, 2H), 3.22–3.11 (m, 1H), 3.02–2.92 (m, 1H), 2.74 (tt, *J* = 12.2, 3.7 Hz, 1H), 1.96–1.88 (m, 2H), 1.79–1.66 (m, 2H).

*(±)-ethyl 1-(chloroformyl)-3-piperidinecarboxylate (**68f’**)*. 0.110 g (78%) of a pale yellow oil containing *ca.* 25% symmetrical urea side product; ^1^H NMR (400 MHz, CDCl_3_): *δ* = 4.30–4.10 (m, 3H), 4.02-3.95 (m, 1H), 3.46 (dd, *J* = 13.7, 9.7 Hz, 1H), 3.24–3.08 (m, 1H), 2.57–2.49 (m, 1H), 2.12–2.01 (m, 1H), 1.84–1.63 (m, 2H), 1.62–1.49 (m, 1H), 1.28–1.20 (m, 3H).

### General Method for Synthesis of Target Alkylsulfonyl Oxadiazoles

A mixture of tetrazole **65** (1 equiv), carbamoyl imidazole or chloride **68** (1 equiv), 2,4,6-collidine (1.2 equiv) and anisole (4 mL per mmol tetrazole) was heated to 130 °C until the evolution of nitrogen had ceased and TLC analysis indicated complete consumption of tetrazole **65**, typically 4 to 18 h. After cooling to ambient temperature, the mixture was diluted with ethyl acetate (10 mL per mmol tetrazole) and washed with 1M hydrochloric acid. The aqueous phase was extracted with ethyl acetate. The combined ethyl acetate extracts were washed with saturated sodium bicarbonate and brine, dried (MgSO_4_), filtered and evaporated to dryness. After purification with flash chromatography, the alkylthio-oxadiazoles **69** were used directly in the next step.

The alkylthio-oxadiazole **69** (1 equiv, 0.13 mmol) was dissolved in acetonitrile (6 mL) and urea-hydrogen peroxide complex (0.315 g, 3.36 mmol, 4 equiv) was added and the mixture cooled to 0 °C. Tri-fluoroacetic anhydride (0.350 mL, 2.52 mmol, 3 equiv) was added dropwise over 2 min. The reaction mixture was stirred at 0 °C for 10 min and then allowed to warm to room temperature and stirred until TLC analysis indicated complete conversion (1 h). Next, 2M sodium bisulfite (5 mL) was added and the mixture was stirred vigorously for 30 min. The mixture was concentrated under vacuum then the residue was diluted with 1M hydrochloric acid (15 mL) and extracted with ethyl acetate (3 x 15 mL). The combined organic layers were washed with saturated sodium bicarbonate (30 mL), brine (30 mL), then dried over MgSO_4_, filtered and evaporated to dryness.

Where noted, alternatively alkylthio-oxadiazoles were oxidised with *m*-chloroperbenzoic acid as follows: A solution of the alkylthio-oxadiazoles (0.36 g, 1.00 mmol, 1 equiv) in dichloromethane (20 mL) was cooled to 0 °C and *m*-chloroperbenzoic acid (70 wt% peracid, 0.42 g, 2.60 mmol, 2.6 equiv) was added. The mixture was allowed to warm to room temperature and stirred for 18 h. Further *m*-chloroperbenzoic acid (70 wt % peracid, 0.42 g, 2.60 mmol, 2.6 equiv) was added and stirring continued for a further 6 h. The mixture was washed with 10% aqueous Na_2_SO_3_ (15 mL) followed by sat aq. NaHCO_3_ (20 mL) then dried over MgSO_4_, filtered and evaporated to dryness. Purification was achieved with flash chromatography//trituration.

*2-(Cyclohexylthio)-5-morpholino-1,3,4-oxadiazole* (**69aa**). Flash chromatography: ethyl acetate/*n*-hexane (50:50 → 100:0). 1.03 g (61%) of a white solid; ^1^H NMR (400 MHz, CDCl_3_): *δ* = 3.79–3.75 (m, 4H), 3.50–3.42 (m, 5H), 2.10–2.04 (m, 2H), 1.79–1.73 (m, 1H), 1.63–1.46 (m, 2H), 1.45–1.23 (m, 3H). ^13^C NMR (100 MHz, CDCl_3_): *δ* = 165.4, 156.6, 66.1, 47.4, 46.3, 33.6, 26.0, 25.6.

*2-(Cyclohexylsulfonyl)-5-morpholino-1,3,4-oxadiazole (**1**).* Flash chromatography: ethyl acetate/dichloromethane (20:80) followed by trituration of the isolated product with dichloromethane. 0.220 g (55%) of white solid; ^1^H NMR (400 MHz, CDCl_3_): *δ =* 3.84–3.79 (m, 4H), 3.65–3.60 (m, 4H), 3.35–3.25 (m, 1H), 2.26–2.18 (m, 2H), 1.99–1.90 (m, 2H), 1.78–1.70 (m, 1H), 1.66–1.53 (m, 2H), 1.39–1.17 (m, 3H). ^13^C NMR (100 MHz, CDCl_3_): *δ =* 164.9, 155.2, 66.0, 63.8, 46.1, 25.1, 25.04, 24.98. HRMS (ESI): m/z calculated for C_12_H_20_O_4_N_3_S + H^+^ [M + H]^+^: 302.1169. Found: 302.1166. LCMS (ESI): *^t^*R = 2.40 min; peak area = 99.7%, m/z: 302 [M+H]^+^.

*5-[(3,4-Dichlorophenyl)mesyl]-2-(4-methyl-1-piperidyl)-1,3,4-oxadiazole (**3**).* Prepared by oxidation with *m*-chloroperbenzoic acid. The product was recrystallized from ethyl acetate/heptane. 0.25 g (67%) of colorless solid; ^1^H NMR (400 MHz, CDCl_3_): *δ* = 7.45 (d, *J* = 8.3 Hz, 1H), 7.42 (d, *J* = 2.1 Hz, 1H), 7.21 (dd, *J* = 8.3, 2.1 Hz, 1H), 4.59 (s, 2H), 3.98–3.93 (m, 2H), 3.07–2.99 (m, 2H), 1.75–1.69 (m, 2H), 1.66–1.56 (m, 1H), 1.26–1.14 (m, 2H), 0.97 (d, *J* = 6.6 Hz, 3H). LCMS (ESI): *^t^*R = 3.41 min; peak area = 98.4%; m/z = 390 [M+H]^+^.

*2-(Cyclopentylsulfonyl)-5-morpholino-1,3,4-oxadiazole (**7**).* Flash chromatography: ethyl acetate/heptane (60:40). 0.024 g (42%) of a white solid; ^1^H NMR (400 MHz, CDCl_3_): *δ* = 3.89–3.76 (m, 5H), 3.62–3.57 (m, 4H), 2.24–2.14 (m, 2H), 2.11–2.01 (m, 2H), 1.87–1.76 (m, 2H), 1.73–1.63 (m, 2H). LCMS (ESI): *^t^*R = 1.96 min; peak area = 94.4%, m/z: 288 [M+H]^+^.

*2-(Cycloheptylsulfonyl)-5-morpholino-1,3,4-oxadiazole (**8**).* Flash chromatography: ethyl acetate/heptane (40:60→60:40). 0.074 g (47%) of a white solid; ^1^H NMR (500 MHz, CDCl_3_): *δ* = 3.81–3.78 (m, 4H), 3.62–3.59 (m, 4H), 3.44–3.37 (m, 1H), 2.30–2.22 (m, 2H), 1.90–1.79 (m, 4H), 1.61–1.48 (m, 6H). LCMS(ESI): *^t^*R = 2.47 min; peak area = 99.7%; m/z = 316 [M+H]^+^.

*2-(Cyclohexylmesyl)-5-morpholino-1,3,4-oxadiazole (**9**).* Flash chromatography: ethyl acetate/heptane (40:60→60:40). 0.015 g (27%) of a colorless oil; ^1^H NMR (400 MHz, CDCl_3_): *δ* = 3.82–3.77 (m, 4H), 3.62–3.58 (m, 4H), 3.34 (d, *J* = 6.4 Hz, 2H), 2.20–2.08 (m, 1H), 1.97–1.89 (m, 2H), 1.75–1.60 (m, 2H), 1.36–1.07 (m, 6H). LCMS (ESI): *^t^*R = 2.61 min; peak area = 95.9%, m/z: 316 [M+H]^+^.

*2-(Cyclopentylmesyl)-5-morpholino-1,3,4-oxadiazole (**10**).* Flash chromatography: ethyl acetate/heptane (40:60→80:20). 0.069 g (72%) of colorless oil; ^1^H NMR (400 MHz, CDCl_3_): *δ* = 3.81–3.77 (m, 4H), 3.62–3.58 (m, 4H), 3.49 (d, *J* = 7.0 Hz, 2H), 2.50–2.38 (m, 1H), 2.03–1.93 (m, 2H), 1.72–1.51 (m, 4H), 1.35–1.22 (m, 2H). LCMS (ESI): *^t^*R = 2.36 min; peak area = 97.7%; m/z = 302 [M+H]^+^.

*2-(Cyclobutylmesyl)-5-morpholino-1,3,4-oxadiazole (**11**).* Flash chromatography: ethyl acetate/heptane (30:70→50:50). 0.065 g (77%) of white solid; ^1^H NMR (400 MHz, CDCl_3_): *δ* = 3.81–3.78 (m, 4H), 3.62-3.58 (m, 4H), 3.51 (d, *J* = 7.3 Hz, 1H), 2.99–2.86 (m, 1H), 2.25–2.13 (m, 2H), 2.03–1.80 (m, 4H).

^13^C NMR (100 MHz, CDCl_3_): *δ* = 164.8, 156.2, 66.0, 61.1, 46.2, 29.2, 28.5, 19.5. HRMS (ESI): m/z calculated for C₁₁H₁₈O₄N₃³²S + H^+^ [M+H]^+^: 288.1013. Found: 288.1011. LCMS (ESI): *^t^*R = 2.07 min; peak area = 99.4%; m/z = 288 [M+H]^+^.

*2-Morpholino-5-(neopentylsulfonyl)-1,3,4-oxadiazole (**12**).* Flash chromatography: ethyl acetate/heptane (30:70→50:50). 0.051 g (67%) of white solid; ^1^H NMR (400 MHz, CDCl_3_): *δ* = 3.81–3.77 (m, 4H), 3.62–3.58 (m, 4H), 3.41 (s, 2H), 1.22 (s, 9H). LCMS (ESI): *^t^*R = 2.25 min; peak area = 99.6%; m/z = 290 [M+H]^+^.

*(±)-5-exo-5-Morpholino-2-(2-norbornanylsulfonyl)-1,3,4-oxadiazole (**13**).* Flash chromatography: ethyl acetate/heptane (30:70→50:50). 0.031 g (66%) of white solid; ^1^H NMR (400 MHz, CDCl_3_): *δ* = 3.81–3.77 (m, 4H), 3.62–3.58 (m, 4H), 3.44–3.37 (m, 1H), 2.90–2.86 (m, 1H), 2.48–2.43 (m, 1H), 2.16–2.08 (m, 1H), 1.84–1.78 (m, 1H), 1.75–1.55 (m, 3H), 1.34–1.19 (m, 3H). LCMS (ESI): *^t^*R = 2.35 min; peak area = 99.7%; m/z = 314 [M+H]^+^.

*2-(1-Adamantanylmesyl)-5-morpholino-1,3,4-oxadiazole (**14**).* Flash chromatography: ethyl acetate/heptane (20:80→60:40). 0.016 g (47%) of white solid; ^1^H NMR (500 MHz, CDCl_3_): *δ* = 3.81–3.78 (m, 4H), 3.61–3.59 (m, 4H), 3.28 (s, 2H), 2.01–1.94 (m, 3H), 1.72–1.65 (m, 6H), 1.31–1.22 (m, 6H). LCMS (ESI): *^t^*R = 3.11 min; peak area = 99.7%; m/z = 368 [M+H]^+^.

*5-Morpholino-2-{[1-(trifluoromethyl)cyclopropyl]mesyl}-1,3,4-oxadiazole (**15**).* Flash chromatography: ethyl acetate/dichloromethane (20:80). Product obtained after freeze-drying from acetonitrile. 0.020 g (45%) of white solid; ^1^H NMR (400 MHz, CDCl_3_): *δ* = 3.82–3.78 (m, 4H), 3.78 (s, 2H), 3.63–3.60 (m, 4H), 1.31–1.27 (m, 2H), 1.24–1.21 (m, 2H). LCMS (ESI): *^t^*R = 2.23 min; peak area = 98.8%; m/z = 342 [M+H]^+^.

*5-Morpholino-2-{[1-(trifluoromethyl)cyclobutyl]mesyl}-1,3,4-oxadiazole (**16**).* Flash chromatography: ethyl acetate/dichloromethane (15:85). White solid, yield 30% (0.020 g, 0.056 mmol). ^1^H NMR (400 MHz, CDCl_3_): *δ* = 3.82–3.78 (m, 4H), 3.78 (s, 2H), 3.64–3.60 (m, 4H), 2.66–2.58 (m, 2H), 2.50–2.41 (m, 2H), 2.17–2.06 (m, 2H). LCMS (ESI): *^t^*R = 2.54 min; peak area = 98.6%; m/z = 356 [M+H]^+^.

*tert-Butyl 4-(5-morpholino-1,3,4-oxadiazol-2-ylsulfonyl)-1-piperidinecarboxylate (**17**).* Flash chromatography: ethyl acetate/dichloromethane (40:60→100:0). 0.036 g (69%) of colorless oil that solidified on standing; ^1^H NMR (400 MHz, CDCl_3_): *δ* = 4.36–4.20 (br s, 2H), 3.83–3.78 (m, 4H), 3.64–3.59 (m, 4H), 3.49–3.40 (m, 1H), 2.78–2.72 (m, 2H), 2.17–2.12 (m, 2H), 1.83–1.71 (m, 2H), 1.44 (s, 9H). LCMS (ESI): *^t^*R = 2.46 min; peak area = 99.3%; m/z = 347 [M-C_4_H_8_+H]^+^.

*2-Morpholino-5-(tetrahydro-2H-pyran-4-ylsulfonyl)-1,3,4-oxadiazole (**19**).* Flash chromatography: ethyl acetate. 0.021 g (47%) of white solid; ^1^H NMR (400 MHz, CDCl_3_): *δ* = 4.14–4.08 (m, 2H), 3.82–3.78 (m, 4H), 3.63–3.52 (m, 5H), 3.45–3.36 (m, 2H), 2.11–2.03 (m, 2H), 2.01–1.89 (m, 2H). LCMS (ESI): *^t^*R = 1.48 min; peak area = 97.2%; m/z = 304 [M+H]^+^.

*2-[(3,4-Dichlorophenyl)mesyl]-5-morpholino-1,3,4-oxadiazole (**20**).* Flash chromatography: ethyl acetate/heptane (40:60→60:40). 0.027 g (32%) of white solid; ^1^H NMR (400 MHz, CDCl_3_): *δ* = 7.46 (d, *J* = 8.3 Hz, 1H), 7.43 (d, *J* = 2.1 Hz, 1H), 7.22 (dd, *J* = 8.3, 2.1 Hz, 1H), 4.61 (s, 2H), 3.78–3.74 (m, 4H), 3.54–3.50 (m, 4H). LCMS (ESI): *^t^*R = 2.69 min; peak area = 95.0%; m/z: 378 [M+H]^+^.

*5-(Cyclohexylsulfonyl)-2-(1-pyrrolidinyl)-1,3,4-oxadiazole (**22**).* Flash chromatography: ethyl acetate/heptane (60:40). 0.071 g (46%) of white solid; ^1^H NMR (400 MHz, CDCl_3_): *δ* = 3.60–3.56 (m, 4H), 3.23 (tt, *J* = 12.2, 3.5 Hz, 1H), 2.21–2.16 (m, 2H), 2.06–2.01 (m, 4H), 1.93–1.89 (m, 2H), 1.72–1.68 (m, 1H), 1.62–1.50 (m, 2H), 1.34–1.16 (m, 3H). LCMS (ESI): *^t^*R = 2.48 min; peak area = 96.0%; m/z: 286 [M+H]^+^.

*2-(Cyclohexylsulfonyl)-5-piperidino-1,3,4-oxadiazole (**23**).* Flash chromatography: ethyl acetate/heptane (20:80→40:60). 0.042 g (14%) of white solid; ^1^H NMR (500 MHz, CDCl_3_): *δ* = 3.59–3.54 (m, 4H), 3.30–3.22 (m, 1H), 2.22–2.16 (m, 2H), 1.95–1.89 (m, 2H), 1.74–1.63 (m, 7H), 1.64–1.55 (m, 2H), 1.36–1.12 (m, 3H). LCMS (ESI): *^t^*R = 2.79 min; peak area = 97.5%; m/z: 300 [M+H]^+^.

*2-(1-Azepanyl)-5-(cyclohexylsulfonyl)-1,3,4-oxadiazole (**24**).* Flash chromatography: ethyl acetate/heptane (40:60). 0.135 g (49%) of white solid; ^1^H NMR (400 MHz, CDCl_3_): *δ* = 3.65–3.60 (m, 4H), 3.30–3.21 (m, 1H), 2.23–2.16 (m, 2H), 1.96–1.88 (m, 2H), 1.84–1.76 (m, 4H), 1.75–1.68 (m, 1H), 1.64–1.51 (m, 6H), 1.36–1.14 (m, 3H). ^13^C NMR (100 MHz, CDCl_3_): *δ* = 165.3, 154.4, 63.8, 48.9, 28.1, 27.7, 25.1, 25.1, 25.0. HRMS (ESI): m/z calculated for C14H24O3N3S + H^+^ [M+H]^+^: 314.1533. Found: 314.1533. LCMS (ESI): *^t^*R = 2.92 min; peak area = 99.3%; m/z: 314 [M+H]^+^.

*5-(Cyclohexylsulfonyl)-2-(1,4-oxazepan-4-yl)-1,3,4-oxadiazole (**25**).* Flash chromatography: ethyl acetate/heptane (40:60). 0.022 g (46%) of colorless oil; ^1^H NMR (400 MHz, CDCl_3_): *δ* = 3.85–3.82 (m, 2H), 3.81–3.75 (m, 6H), 3.32–3.22 (m, 1H), 2.23–2.15 (m, 2H), 2.06–1.99 (m, 2H), 1.96–1.88 (m, 2H), 1.75–1.67 (m, 1H), 1.63–1.51 (m, 2H), 1.36–1.14 (m, 3H). LCMS (ESI): *^t^*R = 2.24 min; peak area = 99.9%; m/z = 316 [M+H]^+^.

*5-(Cyclohexylsulfonyl)-2-(1,2-oxazinan-2-yl)-1,3,4-oxadiazole (**26**).* Flash chromatography: ethyl acetate/heptane (40:60→60:40). 0.018 g (34%) of colorless oil; ^1^H NMR (400 MHz, CDCl_3_): *δ* = 4.14–4.10 (m, 2H), 3.80–3.76 (m, 2H), 3.29 (tt, *J* = 12.2, 3.5 Hz, 1H), 2.22–2.14 (m, 2H), 1.95–1.79 (m, 6H), 1.75–1.67 (m, 1H), 1.65–1.51 (m, 2H), 1.36–1.14 (m, 3H). LCMS (ESI): *^t^*R = 2.70 min; peak area = 95.8%; m/z = 302 [M+H]^+^.

*5-(Cyclohexylsulfonyl)-2-(4-methyl-1-piperidyl)-1,3,4-oxadiazole (**27**).* Flash chromatography: ethyl acetate/heptane (20:80→50:50). 0.022 g (20%) of white solid; ^1^H NMR (400 MHz, CDCl_3_): *δ* = 4.09–4.04 (m, 2H), 3.29–3.22 (m, 1H), 3.12–3.08 (m, 2H), 2.21–2.16 (m, 2H), 1.95–1.90 (m, 2H), 1.78–1.69 (m, 2H), 1.62–1.54 (m, 3H), 1.33–1.18 (m, 6H), (d, *J* = 6.6 Hz, 3H). LCMS (ESI): *^t^*R = 3.08 min; peak area = 99.7%; m/z = 314 [M+H]^+^.

*(±)-5-(Cyclohexylsulfonyl)-2-(3-methyl-1-piperidyl)-1,3,4-oxadiazole (**28**).* Flash chromatography: ethyl acetate/dichloromethane (0:100→5:95). 0.060 g (49%) of yellow oil; ^1^H NMR (400 MHz, CDCl_3_): *δ* = 3.99–3.94 (m, 2H), 3.29–3.22 (m, 1H), 3.05–3.00 (m, 1H), 2.74–2.68 (m, 1H), 2.22–2.17 (m, 2H), 1.95–1.89 (m, 3H), 1.77–1.68 (m, 3H), 1.63–1.53 (m, 3H), 1.33–1.14 (m, 4H), 0.94 (d, *J* = 6.6 Hz, 3H). LCMS (ESI): *^t^*R = 3.06 min; peak area = 98.6%; m/z = 314 [M+H]^+^.

*(±)-5-(Cyclohexylsulfonyl)-2-(2-methyl-1-piperidyl)-1,3,4-oxadiazole (**29**).* Prepared by oxidation with *m*-chloroperbenzoic acid. Flash chromatography: ethyl acetate/dichloromethane (0:100→10:90).

0.030 g (40%) of white solid; ^1^H NMR (500 MHz, CDCl_3_): *δ* = 4.42–4.36 (m, 1H); 3.92–3.86 (m, 1H); 3.30–3.18 (m, 2H); 2.23–2.16 (m, 2H); 1.96–1.88 (m, 2H); 1.84–1.50 (m, 10H); 1.35–1.15 (m, 3H); 1.26 (d, *J* = 6.9 Hz, 3H). LCMS (ESI): *^t^*R = 3.00 min; peak area = 99.3%; m/z = 314 [M+H]^+^.

*5-(Cyclohexylsulfonyl)-2-(4,4-dimethyl-1-piperidyl)-1,3,4-oxadiazole (**30**).* Flash chromatography: ethyl acetate/heptane (20:80→40:60). 0.018 g (10%) of white solid; ^1^H NMR (400 MHz, CDCl_3_): *δ* = 3.59–3.55 (m, 4H), 3.25 (m, 1H), 2.22–2.15 (m, 2H), 1.96–1.88 (m 2H), 1.73–1.72 (m, 1H), 1.70–1.51 (m, 2H), 1.49–1.44 (m, 4H), 1.36–1.15 (m, 3H), 1.00 (s, 6H). LCMS (ESI): *^t^*R = 3.20 min; peak area = 97.5%; m/z = 328 [M+H]^+^.

Cis*/*trans*-5-(Cyclohexylsulfonyl)-2-(3,5-dimethyl-1-piperidyl)-1,3,4-oxadiazole (**31**).* Prepared by oxidation with *m*-chloroperbenzoic acid. Flash chromatography: ethyl acetate/dichloromethane (0:100→10:90). 0.042 g (32%) of white solid; 93:7 mixture of diastereoisomers determined by ^1^H NMR and LCMS analyses. ^1^H NMR (500 MHz, CDCl_3_): *δ* = 4.04–3.99 (m, 2H), 3.29–3.22 (m, 1H), 2.57–2.51 (m, 2H), 2.21–2.17 (m, 2H), 1.94–1.90 (m, 2H), 1.87–1.82 (m, 1H), 1.78–1.69 (m, 3H), 1.62–1.52 (m, 2H), 1.34–1.17 (m, 4H), 0.93 (d, *J* = 6.8 Hz, 6H) (major diastereoisomer). LCMS (ESI): *^t^*R = 3.25 min; peak area = 7.0% (minor diastereoisomer); m/z = 328 [M+H]^+^, *^t^*R = 3.35 min; peak area = 93.0% (major diastereoisomer); m/z = 328 [M+H]^+^.

*5-(Cyclohexylsulfonyl)-2-(3,3-dimethyl-1-piperidyl)-1,3,4-oxadiazole (**32**).* Flash chromatography: ethyl acetate/dichloromethane (0:100→5:95). Freeze dried from acetonitrile. 0.045 g (55%) of white solid; ^1^H NMR (400 MHz, CDCl_3_): *δ* = 3.55–3.51 (m, 2H), 3.29–3.19 (m, 3H), 2.20–2.16 (m, 2H), 1.94–1.89 (m, 2H), 1.74–1.67 (m, 3H), 1.64–1.50 (m, 2H), 1.48–1.43 (m, 2H), 1.37–1.10 (m, 3H), 0.97 (s, 6H). LCMS (ESI): *^t^*R = 3.20 min; peak area = 97.5%; m/z = 328 [M+H]^+^.

Cis*-5-(Cyclohexylsulfonyl)-2-(2,6-dimethyl-1-piperidyl)-1,3,4-oxadiazole (**33**).* Flash chromatography: ethyl acetate/heptane (20:80→40:60). 0.425 g (86%) of white solid; ^1^H NMR (400 MHz, CDCl_3_): *δ* = 4.33–4.29 (m, 2H), 3.26 (tt, *J* = 12.2, 3.5 Hz, m, 1H), 2.23–2.18 (m, 2H), 1.96–1.90 (m, 2H), 1.87–1.55 (m, 10H), 1.36–1.14 (m, 2H), 1.31 (d, *J* = 7.0 Hz, 6H). ^13^C NMR (100 MHz, CDCl_3_): *δ* = 164.7, 154.3, 63.8, 49.1, 29.8, 25.2, 25.1, 25.0, 20.3, 13.4. HRMS (ESI) m/z calculated for C15H26O3N3S + H^+^ [M+H]^+^: 328.1689. Found: 328.1687. LCMS (ESI): *^t^*R = 3.19 min; peak area = 98.0%; m/z: 328 [M+H]^+^.

*(±)-5-(Cyclohexylsulfonyl)-2-(2-methyl-4-morpholinyl)-1,3,4-oxadiazole (**34**).* Flash chromatography: ethyl acetate/heptane (50:50). Freeze dried from acetonitrile. 0.074 g (68%) of yellow oil that solidified on standing; ^1^H NMR (400 MHz, CDCl_3_): *δ* = 3.98 (dd, *J* = 11.0, 2.8 Hz, 1H), 3.93–3.83 (m, 2H), 3.75–3.64 (m, 2H), 3.35–3.25 (m, 2H), 2.95 (dd, *J* = 12.8, 10.4 Hz, 1H), 2.26–2.18 (m, 2H), 1.99–1.90 (m, 2H), 1.78–1.70 (m, 1H), 1.66–1.52 (m, 3H), 1.39–1.19 (m, 3H), 1.22 (d, *J* = 3.8 Hz, 3H). LCMS (ESI): *^t^*R = 2.49 min; peak area = 99.0%; m/z: 316 [M+H]^+^.

*5-(Cyclohexylsulfonyl)-2-(2,2-dimethyl-4-morpholinyl)-1,3,4-oxadiazole (**35**).* Flash chromatography: ethyl acetate/heptane (50:50). Freeze dried from acetonitrile. 0.062 g (44%) of yellow oil that solidified on standing; ^1^H NMR (400 MHz, CDCl_3_): *δ* = 3.86–3.82 (m, 2H), 3.58–3.55 (m, 2H), 3.38 (s, 2H), 3.33–3.24 (m, 1H), 2.25–2.18 (m, 2H), 1.99–1.91 (m, 2H), 1.78–1.70 (m, 1H), 1.66–1.53 (m, 3H), 1.39–1.20 (m, 3H), 1.29 (s, 6H). LCMS (ESI): *^t^*R = 2.61 min; peak area = 98.3%; m/z: 330 [M+H]^+^.

cis*-5-(Cyclohexylsulfonyl)-2-(2,6-dimethyl-4-morpholinyl)-1,3,4-oxadiazole (**36**).* Flash chromatography: ethyl acetate/dichloromethane (10:90). 0.189 g (60%) of white solid; ^1^H NMR (400 MHz, CDCl_3_): *δ* = 3.89–3.84 (m, 2H), 3.76–3.66 (m, 2H), 3.32–3.23 (m, 1H), 2.84 (dd, *J* = 12.7, 10.4 Hz, 2H), 2.23–2.16 (m, 2H), 1.97–1.89 (m, 2H), 1.75–1.69 (m, 1H), 1.64–1.51 (m, 2H), 1.36–1.16 (m, 3H), 1.25 (d, *J* = 6.3 Hz, 6H). LCMS (ESI): *^t^*R = 2.70 min; peak area = 99.6%; m/z: 330 [M+H]^+^.

*5-(Cyclohexylsulfonyl)-2-(2-oxa-7-aza-7-spiro[3.5]nonyl)-1,3,4-oxadiazole (**37**)*. Flash chromatography: ethyl acetate. 0.110 g (83%) of colorless oil; ^1^H NMR (400 MHz, CDCl_3_): *δ* = 4.46 (s, 4H), 3.56–3.51 (m, 4H), 3.30–3.22 (m, 1H), 2.22–2.14 (m, 2H), 2.00–1.95 (m, 4H), 1.95–1.88 (m, 2H), 1.74–1.67 (m, 1H), 1.62–1.50 (m, 2H), 1.36–1.13 (m, 3H). LCMS (ESI): *^t^*R = 2.44 min; peak area = 98.9%; m/z: 342 [M+H]^+^.

*5-(Cyclohexylsulfonyl)-2-(2-oxa-6-aza-6-spiro[3.5]nonyl)-1,3,4-oxadiazole (**38**)*. Flash chromatography: ethyl acetate/dichloromethane (20:90). Freeze dried from acetonitrile. 0.143 g (63%) of low melting white solid; ^1^H NMR (400 MHz, CDCl_3_): *δ* = 4.41 (s, 4H), 3.78 (s, 2H), 3.50 (t, *J* = 5.7 Hz, 2H), 3.35–3.26 (m, 1H), 2.26–2.18 (m, 2H), 1.99–1.91 (m, 4H), 1.78–1.57 (m, 5H), 1.39–1.17 (m, 3H). LCMS (ESI):*^t^*R = 2.37 min; peak area = 98.9%; m/z: 342 [M+H]^+^.

*5-(Cyclohexylsulfonyl)-2-(7-oxa-2-aza-2-spiro[3.5]nonyl)-1,3,4-oxadiazole (**39**)*. Flash chromatography: ethyl acetate/dichloromethane (20:90). 0.007 g (32%) of white solid; ^1^H NMR (400 MHz, CDCl_3_): *δ* = 4.00 (s, 4H), 3.65–3.61 (m, 4H), 3.30–3.21 (m, 1H), 2.22–2.14 (m, 2H), 1.96–1.88 (m, 2H), 1.87–1.82 (m, 4H), 1.75–1.50 (m, 3H), 1.36–1.14 (m, 3H). LCMS (ESI): *^t^*R = 2.50 min; peak area = 97.6%; m/z: 342 [M+H]^+^.

*5-(Cyclohexylsulfonyl)-2-(1,4-dioxa-8-aza-8-spiro[4.5]decyl)-1,3,4-oxadiazole (**40**)*. Flash chromatography: ethyl acetate/heptane (40:80→60:40). 0.050 g (27%) of white solid; ^1^H NMR (400 MHz, CDCl_3_): *δ* = 3.98 (s, 4H), 3.74–3.70 (m, 4H), 3.30–3.22 (m, 1H), 2.22–2.16 (m, 2H), 1.96–1.88 (m, 2H), 1.82–1.77 (m, 4H), 1.74–1.68 (m, 1H), 1.62–1.52 (m, 2H), 1.35–1.15 (m, 3H). LCMS (ESI): *^t^*R = 2.52 min; peak area = 98.3%; m/z: 358 [M+H]^+^.

*5-(Cyclohexylsulfonyl)-2-(1-oxa-8-aza-8-spiro[4.5]decyl)-1,3,4-oxadiazole (**41**)*. Flash chromatography: ethyl acetate/heptane (40:60→60:40). 0.018 g (16%) of white solid; ^1^H NMR (400 MHz, CDCl_3_): *δ* = 3.87–3.77 (m, 4H), 3.56–3.49 (m, 2H), 3.36–3.33 (m, 1H), 2.19–2.12 (m, 1H), 1.98–1.90 (m, 3H), 1.84–1.80 (m, 2H); 1.74–1.59 (m, 8H); 1.46–1.23 (m, 4H). LCMS (ESI): *^t^*R = 2.77 min; peak area = 99.8%; m/z = 356 [M+H]^+^.

*5-(Cyclohexylsulfonyl)-2-(8-oxa-3-azabicyclo[3.2.1]oct-3-yl)-1,3,4-oxadiazole (**42**)*. Flash chromatography: ethyl acetate/heptane (50:50→100:0). 0.053 g (50%) of colorless oil; ^1^H NMR (400 MHz, CDCl_3_): *δ* = 4.47–4.44 (m, 2H), 3.65–3.60 (m, 2H), 3.45 (dd, *J* = 12.5, 2.5 Hz, 2H), 3.28 (tt, *J* = 12.2, 3.5 Hz, m, 1H), 2.23–2.15 (m, 2H), 2.09–1.99 (m, 2H), 1.98–1.83 (m, 4H), 1.75–1.69 (m, 1H), 1.63–1.51 (m, 2H), 1.36–1.14 (m, 3H). LCMS (ESI): *^t^*R = 2.37 min; peak area = 96.0%; m/z 328 [M+H]^+^.

*5-(Cyclohexylsulfonyl)-2-(3-oxa-8-azabicyclo[3.2.1]oct-8-yl)-1,3,4-oxadiazole (**43**)*. Flash chromatography: ethyl acetate/heptane (30:70→100:0). 0.030 g (32%) of white solid; ^1^H NMR (400 MHz, CDCl_3_): *δ* = 4.31–4.28 (m, 2H), 3.81 (d, *J* = 11.4 Hz, 2H), 3.64 (broad d, *J* = 11.4 Hz, 2H), 3.34–3.26 (m, 1H), 2.24–2.06 (m, 6H), 1.96–1.91 (m, 2H), 1.75–1.69 (m, 1H), 1.64–1.52 (m, 2H), 1.37–1.15 (m, 3H). LCMS (ESI): *^t^*R = 2.48 min; peak area = 96.2%; m/z 328 [M+H]^+^.

*(±)-2-(2-Azabicyclo[2.2.1]heptan-2-yl)-5-(cyclohexylsulfonyl)-1,3,4-oxadiazole (**44**)*. Flash chromatography: ethyl acetate/heptane (40:60→60:40). Freeze-dried from acetonitrile. 0.022 g (45%) of white solid; ^1^H NMR (400 MHz, methanol-*d*_4_): *δ* = 4.38 (s, 1H), 3.57–3.52 (m, 1H), 3.41–3.32 (m, 1H), 3.25 (dd, *J* = 9.1, 1.3 Hz, 1H), 2.73 (m, 1H), 2.18–2.10 (m, 2H), 1.96–1.88 (m, 2H), 1.87–1.68 (m, 5H), 1.67–1.62 (m, 1H), 1.57–1.46 (m, 3H), 1.44–1.17 (m, 3H). LCMS (ESI): *^t^*R = 2.77 min; peak area = 97.3%; m/z 312 [M+H]^+^.

*2-(8-Azabicyclo[3.2.1]octan-8-yl)-5-(cyclohexylsulfonyl)-1,3,4-oxadiazole (**45**).* Flash chromatography: ethyl acetate/heptane (30:70→50:50). Freeze-dried from acetonitrile. 0.009 g (13%) as white solid; ^1^H NMR (400 MHz, methanol-*d*_4_): *δ* = 4.39–4.33 (m, 2H), 3.44–3.35 (m, 1H), 2.19–2.10 (m, 4H), 1.98–1.80 (m, 6H), 1.76–1.68 (m, 1H), 1.65–1.19 (m, 9H). LCMS (ESI): *^t^*R = 3.00 min; peak area = 98.5%; m/z 326 [M+H]^+^.

*2-(7-Azabicyclo[2.2.1]heptan-7-yl)-5-(cyclohexylsulfonyl)-1,3,4-oxadiazole (**46**).* Flash chromatography: ethyl acetate/heptane (35:65). Freeze-dried from acetonitrile. 0.004 g (5%) of white solid; ^1^H NMR (400 MHz, methanol-*d*_4_): *δ* = 4.47–4.44 (m, 2H), 3.41 (m, 1H), 2.17–2.09 (m, 2H), 1.96–1.85 (m, 5H), 1.76–1.68 (m, 1H), 1.67–1.60 (m, 4H), 1.57–1.46 (m, 2H), 1.45–1.19 (m, 4H). LCMS (ESI): *^t^*R = 2.94 min; peak area = 99.0%; m/z 312 [M+H]^+^.

*(±)-5-(Cyclohexylsulfonyl)-2-(2-oxa-5-azabicyclo[2.2.1]hept-5-yl)-1,3,4-oxadiazole (**47**).* Flash chromatography: ethyl acetate/heptane (50:50→100:0). 0.012 g (54%) of white solid; ^1^H NMR (400 MHz, methanol-*d*_4_): *δ* = 4.76–4.70 (m, 2H), 3.94 (d, *J* = 7.8 Hz, 1H), 3.87 (dd, *J* = 7.9, 1.4 Hz, 1H), 3.63 (dd, *J* = 9.9, 1.4 Hz, 1H), 3.56 (d, *J* = 9.9 Hz, 1H), 3.43–3.36 (m, 1H), 2.20–2.11 (m, 2H), 2.11–2.02 (m 2H), 1.96–1.88 (m, 2H), 1.76–1.68 (m, 1H), 1.58–1.46 (m, 2H), 1.44–1.18 (m, 3H). LCMS (ESI): *^t^*R = 2.17 min; peak area = 97.2%; m/z 314 [M+H]^+^.

*5-(Cyclohexylsulfonyl)-2-(4-methoxy-1-piperidyl)-1,3,4-oxadiazole (**48**).* Flash chromatography: ethyl acetate/heptane (60:40). 0.099 g (47%) of white solid; ^1^H NMR (400 MHz, CDCl_3_): *δ* = 3.79–3.71 (m, 2H), 3.53–3.45 (m, 3H), 3.35 (s, 3H), 3.28–3.19 (m, 1H), 2.17 (dd, *J* = 13.3, 1.9 Hz, 2H), 1.94–1.86 (m, 4H), 1.77–1.67 (m, 3H), 1.54 (qd, *J* = 12.5, 3.4Hz), 1.34–1.16 (m, 3H). LCMS (ESI): *^t^*R = 2.50 min; peak area = 96.8%; m/z 330 [M+H]^+^.

*1-(5-(Cyclohexylsulfonyl)-1,3,4-oxadiazol-2-yl)-4-piperidinecarbonitrile (**49**).* Flash chromatography: ethyl acetate/heptane (60:40). 0.053 g (42%) of pale yellow solid; ^1^H NMR (400 MHz, CDCl_3_): *δ* = 3.82–3.74 (m, 2H), 3.70–3.62 (m, 2H), 3.27 (tt, *J* = 12.2, 3.5 Hz, 1H), 2.99–2.92 (m, 1H), 2.18 (dd, *J* =13.3, 1.8 Hz, 2H), 2.08–1.88 (m, 6H), 1.71 (d, *J* = 12.3 Hz, 1H), 1.62–1.50 (m, 2H), 1.36–1.14 (m, 3H). LCMS (ESI): *^t^*R = 2.45 min; peak area = 98.7%; m/z 325 [M+H]^+^.

*5-(Cyclohexylsulfonyl)-2-(4-phenyl-1-piperidyl)-1,3,4-oxadiazole (**50**).* Flash chromatography: ethyl acetate/heptane (20:80→40:60). 0.022 g (23%) as white solid; ^1^H NMR (400 MHz, CDCl_3_): *δ* = 7.34– 7.29 (m, 2H), 7.24–7.17 (m, 3H), 4.28–4.21 (m, 2H), 3.32–3.17 (m, 3H), 2.75 (tt, *J* = 12.1, 3.4 Hz, 1H), 2.24–2.16 (m, 2H), 2.00–1.90 (m, 4H), 1.87–1.68 (m, 3H), 1.64–1.52 (m, 2H), 1.37–1.16 (m, 3H). LCMS (ESI): *^t^*R = 3.41 min; peak area = 96.8%; m/z 376 [M+H]^+^.

*5-(Cyclohexylsulfonyl)-2-[4-(trifluoromethyl)-1-piperidyl]-1,3,4-oxadiazole (**51**).* Flash chromatography: ethyl acetate/dichloromethane (10:90). 0.025 g (20%) as yellow solid; ^1^H NMR (400 MHz, CDCl_3_): *δ* = 4.27–4.20 (m, 2H), 3.34–3.24 (m, 1H), 3.18–3.09 (m, 2H), 2.38–2.25 (m, 1H), 2.25–2.17 (m, 2H), 2.07–1.99 (m, 2H), 1.99–1.90 (m, 2H), 1.78–1.67 (m, 2H), 1.67–1.56 (m, 3H), 1.39–1.19 (m, 3H). LCMS (ESI): *^t^*R = 3.07 min; peak area = 99.8%; m/z 368 [M+H]^+^.

*(±)-Ethyl 1-(5-(cyclohexylsulfonyl)-1,3,4-oxadiazol-2-yl)-3-piperidinecarboxylate (**52**).* Flash chromatography: ethyl acetate/dichloromethane, 0:100→10:90. 0.014 g (9%) as pale yellow solid; ^1^H NMR (500 MHz, CDCl_3_): *δ* = 4.14 (q, *J* = 7.1 Hz, 2H), 4.07 (dd, *J* = 13.3, 4.0 Hz, 1H), 3.89–3.87 (m, 1H), 3.44 (dd, *J* = 13.3, 9.6 Hz, 1H), 3.29–3.24 (m, 2H), 2.61 (m, 1H), 2.20–2.18 (m, 2H), 2.11–2.08 (m, 1H), 1.94–1.90 (m, 2H), 1.85–1.83 (m, 1H), 1.78–1.50 (m, 4H), 1.33–1.18 (m, 6H). LCMS (ESI): *^t^*R = 3.17 min; peak area = 99.8%; m/z 372 [M+H]^+^.

tert*-Butyl (*S*)-1-[5-(cyclohexylsulfonyl)-1,3,4-oxadiazol-2-yl]-2-pyrrolidinecarboxylate (**53**).* Flash chromatography: ethyl acetate/heptane/acetic acid (59:40:1). 0.007 g (11%) as orange crystals; ^1^H NMR (400 MHz, CDCl_3_): *δ* = 4.44–4.39 (m, 2H), 3.79–3.73 (m, 2H), 3.69–3.62 (m, 2H), 3.38–3.32 (m, 1H), 3.28–3.22 (m, 1H), 2.39–2.29 (m, 2H), 2.20–2.13 (m, 2H), 2.09–1.20 (m, 6H), 1.42 (s, 9H). *^t^*R = 2.80 min; peak area = 95.1%; m/z: 330 [M-(C_4_H_8_)+H]^+^.

*5-(Cyclohexylsulfonyl)-2-(4,4-difluoro-1-piperidyl)-1,3,4-oxadiazole (**54**).* Purified by trituration with dichloromethane/heptane. 0.048 g (39%) of white solid; ^1^H NMR (400 MHz, CDCl_3_): *δ* = 3.79–3.74 (m, 4H), 3.29 (tt, *J* = 12.2, 3.5 Hz, 1H), 2.21–2.06 (m, 6H), 1.99–1.91(m, 2H), 1.74–1.70 (m, 1H), 1.63–1.52 (m, 2H), 1.37–1.16 (m, 3H). LCMS (ESI) *^t^*R = 2.80 min; peak area = 98.3%; m/z: 336 [M+H]^+^.

*4-[5-(Cyclohexylsulfonyl)-1,3,4-oxadiazol-2-yl]-1λ^6^,4-thiazinane-1,1-dione (**55**).* Flash chromatography: ethyl acetate/heptane (60:40→100:0). 0.103 g (∼100%) of white solid; ^1^H NMR (400 MHz, CDCl_3_): *δ* = 4.19–4.14 (m, 4H), 3.31 (tt, *J* = 12.1, 3.5 Hz, 1H), 3.19–3.16 (m, 4H), 2.22–2.14 (m, 2H), 1.98–1.90 (m, 2H), 1.76–1.69 (m, 1H), 1.63–1.52 (m, 2H), 1.37–1.16 (m, 3H). LCMS (ESI): *^t^*R = 2.13 min; peak area = 99.1%; m/z = 350 [M+H]^+^.

*1-{4-[5-(Cyclohexylsulfonyl)-1,3,4-oxadiazol-2-yl]-1-piperazinyl}-1-ethanone (**58**).* Flash chromatography: ethyl acetate/methanol (97:3). 0.041 g (15%) of pale yellow solid; ^1^H NMR (400 MHz, CDCl_3_): *δ* = 3.77–3.73 (m, 2H), 3.66–3.58 (m, 6H), 3.29 (tt, *J* = 12.2, 6.9 Hz, 1H), 2.22–2.16 (m, 2H), 2.14 (s, 3H), 1.97–1.90 (m, 2H), 1.73 (d, *J* = 12.2 Hz, 1H) 1.63–1.52 (m, 2H), 1.37–1.17 (m, 3H). LCMS (ESI) (ESI): *^t^*R = 2.63 min; peak area = 99.0%; m/z = 358 [M+H]^+^.

*tert-Butyl 4-[5-(cyclohexylsulfonyl)-1,3,4-oxadiazol-2-yl]-1-piperazinecarboxylate (**59**).* Flash chromatography: ethyl acetate/heptane (20:80→40:60). 0.180 g (82%) of white solid; ^1^H NMR (400 MHz, CDCl_3_): *δ* = 3.60–3.52 (m, 8H), 3.27 (tt, *J* = 12.1, 3.5 Hz, 1H), 2.23–2.15 (m, 2H), 1.97–1.88 (m, 2H), 1.75–1.68 (m, 1H), 1.62–1.51 (m, 2H), 1.46 (s, 9H), 1.37–1.15 (m, 3H). LCMS (ESI): *^t^*R = 3.03 min; peak area = 99.6%; m/z = 345 [(M-C_4_H_8_)+H]^+^.

*5-(Cyclohexylsulfonyl)-2-(1-indolinyl)-1,3,4-oxadiazole (**61**).* Prepared by oxidation with *m*-chloroperbenzoic acid. Flash chromatography: ethyl acetate/heptane (40:60→60:40). 0.008 g (8%) of white solid; ^1^H NMR (500 MHz, CDCl_3_): *δ* = 7.75 (d, *J* = 8.0 Hz, 1H), 7.29–7.24 (m, 2H), 7.07–7.02 (m, 1H), 4.25 (m, 2H), 3.36–3.29 (m, 3H), 2.25–2.20 (m, 2H), 1.96–1.90 (m, 2H), 1.75–1.71 (m, 1H), 1.64–1.55 (m, 2H), 1.34–1.15 (m, 3H). LCMS (ESI): *^t^*R = 3.18 min; peak area = 95.0%; m/z: 334 [M+H]^+^.

*5-(Cyclohexylsulfonyl)-2-(3,4-dihydro-2*H*-1,4-benzoxazin-4-yl)-1,3,4-oxadiazole (**62**).* Flash chromatography: ethyl acetate/heptane (40:60→60:40). 0.170 g (∼100%) of white solid; ^1^H NMR (400 MHz, CDCl_3_): *δ* = 7.98 (dd, *J* = 8.2, 1.5 Hz, 1H), 7.09–7.03 (m, 1H), 7.02–6.97 (m, 1H), 6.97–6.94 (dd, *J* = 8.0, 1.7 Hz, 1H), 4.37–4.34 (m, 2H), 4.17–4.12 (m, 2H), 3.33 (tt, *J* = 12.2, 3.5 Hz, 1H), 2.26–2.19 (m, 2H), 1.98–1.90 (m, 2H), 1.76–1.69 (m, 1H), 1.67–1.55 (m, 2H), 1.38–1.16 (m, 3H). LCMS (ESI): *^t^*R = 3.12 min; peak area = 96.4%; m/z = 350 [M+H]^+^.

*5-Morpholino-2-[(2,5-xylyl)methylthio]-1,3,4-oxadiazole (**4**).* Flash chromatography: ethyl acetate/heptane (2:1). 0.163 g (39%) of white solid; ^1^H NMR (400 MHz, CDCl_3_): *δ* = 7.13–7.02 (m, 2H), 7.01–6.96 (m, 1H), 4.29 (s, 2H), 3.77–3.73 (m, 4H), 3.45–3.41 (m, 4H), 2.34 (s, 3H), 2.26 (s, 3H). ^13^C NMR (125 MHz, CDCl_3_): *δ* = 165.3, 156.8, 136.0, 134.1, 133.4, 131.0, 130.8, 129.3, 66.1, 46.3, 36.1, 21.0, 18.9. LCMS (ESI): *^t^*R = 2.85 min; peak area = 98.5%; m/z = 306 [M+H]^+^.

*5-Morpholino-2-[(2,5-xylyl)mesyl]-1,3,4-oxadiazole (**2**)* and *5-morpholino-2-[(2,5-xylyl)methylsulfinyl]-1,3,4-oxadiazole (**5**)*. The sulfide **4** (0.400 g, 1.31 mmol) was dissolved in dichloromethane (15 mL) then *m*-chloroperbenzoic acid (77 wt%, 0.763 g, 3.40 mmol) was added and the mixture was stirred at room temperature for 18 h. The mixture was quenched by addition of 10% aqueous NaHSO_3_. The organic phase was separated, and the aqueous phase extracted with dichloromethane (2x). The combined organic phase was washed with brine, dried over MgSO_4_ and evaporated. Flash chromatography of the residue, eluting with ethyl acetate/heptane (40:60→60:40) gave two fractions. The faster eluting fraction gave *title compound* **2**. 0.26 g (60%) of white solid; ^1^H NMR (400 MHz, CDCl_3_): *δ* = 7.13–7.07 (m, 2H), 6.98 (m, 1H), 4.68 (s, 2H), 3.77–3.73 (m, 4H), 3.52–3.48 (m, 4H), 2.36 (s, 3H), 2.27 (s, 3H). ^13^C NMR (100 MHz, CDCl_3_): *δ* = 164.8, 156.1, 136.2, 136.1, 132.8, 131.3, 130.7, 123.8, 66.0, 59.9, 46.1, 21.0, 19.5. LCMS (ESI): *^t^*R = 2.60 min; peak area = 98.8%; m/z = 338 [M+H]^+^. The slower eluting fraction gave *title compound* **5.** 0.042 g (10%) of colorless oil; ^1^H NMR (400 MHz, CDCl_3_): *δ* = 7.10–7.02 (m, 2H), 6.98 (m, 1H), 4.61 (d, *J* = 13.0 Hz, 1H), 4.52 (d, *J* = 13.0 Hz, 1H), 3.80–3.76 (m, 4H), 3.58–3.54 (m, 4H), 2.33 (s, 3H), 2.26 (s, 3H). ^13^C NMR (125 MHz, CDCl_3_): *δ* = 165.4, 159.5, 136.4, 134.8, 132.2, 131.1, 130.2, 126.8, 66.1, 58.1, 46.2, 21.0, 19.4. LCMS (ESI): *^t^*R = 2.23 min; peak area = 99.0%; m/z = 322 [M+H]^+^.

*5-Morpholino-2-(4-piperidylsulfonyl)-1,3,4-oxadiazole (**18**). N*-Boc-protected amine **17** (0.032 g, 0.080 mmol) was dissolved in a mixture of trifluoroacetic acid (1.80 mL), triisopropyl silane (0.100 mL) and water (0.100 mL). The mixture was stirred at room temperature for 2 h then evaporated. The residue was partitioned between ethyl acetate (20 mL) and 2M hydrochloric acid (20 mL). The aqueous phase was separated and evaporated. 0.028 g (100%) of colorless foam; ^1^H NMR (400 MHz, methanol-*d*_4_): *δ* = 3.93–3.85 (m, 1H), 3.83–3.80 (m, 4H), 3.64–3.60 (m, 4H), 3.60–3.54 (m, 2H), 3.16–3.07 (m, 2H), 2.44–2.36 (m, 2H), 2.14–2.02 (m, 2H). LCMS (ESI): *^t^*R = 0.87 min; peak area = 99.6%; m/z = 303 [M+H]^+^.

*2-(3,4-Dichlorophenylsulfonyl)-5-morpholino-1,3,4-oxadiazole (**21**).* A flask was charged with 3,4-dichloroaniline (**66**) (0.65 g, 4.0 mmol), then a solution of sodium nitrite (0.305 g, 4.4 mmol) in 25% tetrafluoroboric acid (5 mL) was added dropwise at 0 °C. The mixture was stirred at 0 °C for 2 h. Filtration afforded *1-(3,4-dichlorophenyl)-2-(tetrafluoro-l5-boraneyl)diazene* (**63p**) which was refrigerated until required. 0.582 g (56%) of white solid; ^1^H NMR (400 MHz, CD_3_CN): *δ* = 8.63 (d, *J* = 2.4 Hz, 1H), 8.41 (dd, *J* = 2.4, 9.0 Hz, 1H), 8.07 (d, *J* = 9.0 Hz, 1H).

A flask was charged with potassium thiocyanate (0.26 g, 2.7 mmol) and dry acetonitrile (1 mL). Tetrakis(acetonitrile) copper (I) tetrafluoroborate (0.042 g, 0.13 mmol), copper (II) tetrafluoroborate hydrate (0.032 g, 0.13 mmol) followed by tetramethylethylenediamine (TMEDA) (20 μL, 0.13 mmol) in acetonitrile (1 mL) were added. After stirring for 30 min, the reaction mixture was cooled to 0 °C and a solution of *1-(3,4-dichlorophenyl)-2-(tetrafluoro-l5-boraneyl)diazene* (**63p**) (0.35 g, 1.34 mmol) in acetonitrile (5 mL) was slowly added. The mixture was stirred for an hour at 0 °C then allowed to warm to room temperature and poured into diethyl ether (50 mL), filtered through Celite™ to remove the precipitated inorganic salts and then evaporated. Flash chromatography eluting with ethyl acetate/heptane (2:98) gave *1,2-dichloro-4-thiocyanatobenzene* (**64p**). 0.12 g (44%) of colorless oil; ^1^H NMR (500 MHz, CDCl_3_): *δ* = 7.64 (d, *J* = 2.3 Hz, 1H), 7.53 (d, *J* = 8.5 Hz, 1H), 7.37 (dd, *J* = 2.3, 8.5 Hz, 1H).

A microwave tube was charged with *1,2-dichloro-4-thiocyanatobenzene* (**64p**) (0.040 g, 0.20 mmol) and *t*-amyl alcohol (1 mL). A solution of sodium azide (0.014 g, 0.22 mmol) and zinc bromide dihydrate (0.048 g, 0.18 mmol) in water (2 mL) was added. The tube was sealed and heated at 100 °C for 20 h. The mixture was cooled to room temperature then 1M hydrochloric acid (2 mL) and ethyl acetate (2 mL) were added with stirring until the solids dissolved and the aqueous phase was adjusted to pH 1. The organic phase was separated and dried over MgSO_4_, filtered, and evaporated. Flash chromatography eluting with ethyl acetate/acetic acid (99:1) to give *5-(3,4-Dichlorophenylthio)-2*H*-tetrazole* (**65p**). 0.053 g (55%) of colorless solid; ^1^H NMR (400 MHz, CDCl_3_): *δ* = 7.72 (d, *J* = 2.0 Hz, 1H), 7.49 (d, *J* = 8.4 Hz, 1H), 7.45 (dd, *J* = 8.4, 2.0 Hz, 1H).

The reaction of *5-(3,4-Dichlorophenylthio)-2*H*-tetrazole* (**65p**) with carbamoyl imidazole **68a** was carried out according to the General Procedure. Flash chromatography eluting with ethyl acetate/heptane (80:20) gave *2-(3,4-dichlorophenylthio)-5-morpholino-1,3,4-oxadiazole*. 0.025 g (37%) of a white solid; ^1^H NMR (400 MHz, CDCl_3_): *δ* = 7.62 (d, *J* = 2.2 Hz, 1H), 7.44 (d, *J* = 8.44 Hz, 1H), 7.35 (dd, *J* = 8.4, 2.2 Hz, 1H), 3.76–3.79 (m, 4H), 3.48–3.51 (m, 4H).

The oxidation of *2-(3,4-dichlorophenylthio)-5-morpholino-1,3,4-oxadiazole* was carried out according to the General Procedure. Flash chromatography eluant: ethyl acetate/heptane (60:40→100:0) afforded the *title compound*. 0.010 g (37%) of white solid; ^1^H NMR (400 MHz, CDCl_3_): *δ* = 8.13 (d, *J* = 2.2 Hz, 1H), 7.89 (dd, *J* = 2.2, 8.5 Hz, 1H), 7.68 (d, *J* = 8.5 Hz, 1H), 3.77 (t, *J* = 4.9 Hz, 4H), 3.57 (t, *J* = 4.9 Hz, 4H). ^13^C NMR (100 MHz, CDCl_3_): *δ* = 164.9, 156.4, 140.9, 137.5, 134.8, 132.0, 130.9, 128.0, 65.9, 46.1. LCMS (ESI): *^t^*R = 2.75 min; peak area = 94.9%; m/z = 364 [M+H]^+^.

*5-(Cyclohexylsulfonyl)-2-(1-piperazinyl)-1,3,4-oxadiazole hydrochloride (**56**).* The *N*-Boc piperazine **59** (0.283 g, 0.707 mmol) was dissolved in dichloromethane (2 mL) and 4M hydrochloric acid in dioxane (0.265 μL, 1.06 mmol) was added. The mixture was stirred at room temperature for 4 h then evaporated. The residue was dissolved in water and freeze-dried. 0.137 g (58%) of pale yellow powder; ^1^H NMR (free base) (400 MHz, CDCl_3_): *δ* = 4.10 (broad s, 1H, N<u>H</u>), 3.67–3.62 (m, 4H), 3.27 (tt, *J* = 12.2, 3.4 Hz, 1H), 3.05–3.01 (m, 4H), 2.23–2.15 (m, 2H), 1.97–1.88 (m, 2H), 1.76–1.69 (m, 1H), 1.62–1.50 (m, 2H), 1.36–1.14 (m, 3H). LCMS (ESI): *^t^*R = 1.41 min; peak area = 99.2%; m/z: 301 [M+H]^+^.

*5-(Cyclohexylsulfonyl)-2-(4-methyl-1-piperazinyl)-1,3,4-oxadiazole (**57**)*. The amine **56** (0.044 g, 0.15 mmol) (free base) was dissolved in formic acid (40 μL, 1.1 mmol) and formaldehyde (30% aq, 136 μL, 1.50 mmol) was added. The mixture was heated to 80 °C for 2 h (complete conversion by LCMS analysis). After cooling, the mixture was diluted with water (5 mL) and adjusted to pH 11 by the addition of concentrated aqueous ammonia. The mixture was extracted with ethyl acetate (2 x 5 mL). The combined organic phase was washed with brine, dried over Na_2_SO_4_, filtered, and evaporated. The crude product was purified by flash chromatography eluting with methanol/ethyl acetate (10:90). 0.019 g (41%) of colorless oil; ^1^H NMR (400 MHz, CDCl_3_): *δ* = 3.66–3.61 (m, 4H), 3.31–3.22 (m, 1H), 2.53–2.48 (m, 4H), 2.34 (s, 3H), 2.23–2.15 (m, 2H), 1.97–1.88 (m, 2H), 1.75–1.67 (m, 1H), 1.63–1.50 (m, 2H), 1.36–1.14 (m, 3H). LCMS (ESI): *^t^*R = 1.45 min; peak area = 97.3%; m/z: 315 [M+H]^+^.

*5-(Cyclohexylsulfonyl)-2-[4-(2,2,2-trifluoroethyl)-1-piperazinyl]-1,3,4-oxadiazole (**60**)*. A dry microwave vial was charged with the secondary amine **56** (0.033 g, 0.11 mmol) and tetrahydrofuran (0.2 mL). The vial was sealed and immersed in an oil bath pre-heated to 70 °C and phenylsilane (27 μL, 0.22 mmol) was injected through the septum, followed by trifluoroacetic acid (15 μL, 0.19 mmol). The vial was maintained at 70 °C for 2 h. After cooling, the mixture was partitioned between ethyl acetate and sat’d NaHCO_3._ The organic phase was dried over Na_2_SO_4_, filtered, and evaporated to give a crude product which consisted of a 3:1 mixture of the desired amine and the intermediate trifluoroacetamide, which were difficult to separate chromatographically. The mixture was dissolved in methyl *t*-butyl ether and extracted twice with 2M hydrochloric acid. The aqueous phase was carefully neutralized with solid NaHCO_3_ and extracted with ethyl acetate. The combined organic phase was washed with brine, dried over Na_2_SO_4_, filtered and evaporated. 0.012 g (29%) of colorless oil; ^1^H NMR (400 MHz, CDCl_3_): *δ* = 3.66–3.62 (m, 4H), 3.31–3.23 (m, 1H), 3.04 (q, *J* = 9.4 Hz, 2H), 2.82–2.77 (m, 4H), 2.23–2.15 (m, 2H), 1.97–1.88 (m 2H), 1.76–1.67 (m, 1H), 1.63–1.51 (m, 2H), 1.38–1.14 (m, 3H). LCMS (ESI): *^t^*R = 2.99 min; peak area = 97.8%; m/z = 383 [M+H]^+^.

*2-(Cyclohexylsulfonyl)-5-morpholino-1,3,4-thiadiazole (**6**)*. A mixture of 3-methyl-1-(morpholin-4-ylcarbonothioyl)-1H-imidazol-3-ium iodide (**70**) (0.392 g, 1.11 mmol) (cross-ref 32), tetrazole **65a** (0.170 g, 0.922 mmol), anisole (4 mL) and collidine (147 µL, 0.914 mmol) was heated to 130 °C for 4 h. The mixture was allowed to cool to room temperature, diluted with ethyl acetate (15 mL) and washed successively with 2M hydrochloric acid, saturated NaHCO_3_ and brine. The organic layer was dried over anhydrous MgSO_4_ filtered and evaporated. The product was purified by flash chromatography eluting with ethyl acetate /heptane (20:80→30:70) affording *2-(cyclohexylthio)-5-morpholino-1,3,4-thiadiazole* (**71**). 0.17 g (65%) of colorless oil; ^1^H NMR (400 MHz, CDCl_3_): *δ* = 3.83–3.79 (m, 4H), 3.51–3.47 (m, 5H), 2.12–2.05 (m, 2H), 1.81–1.73 (m, 2H), 1.65–1.58 (m, 1H), 1.54–1.42 (m, 2H), 1.41–1.23 (m, 3H). LCMS (ESI) analysis: *^t^*R = 2.92 min; peak area = 98.5%; m/z: 286 [M+H]^+^. Oxidation of sulfide **71** (160 mg, 0.50 mmol) was performed according to the General Method. Flash chromatography eluting with ethyl acetate/heptane (30:70 → 50:50) and ethyl acetate/dichloromethane (10:90) gave the *title compound* (**6**). 0.14 g (78%) of white solid; ^1^H NMR (400 MHz, CDCl_3_): *δ =* 3.85–3.82 (m, 4H), 3.62–3.59 (m, 4H), 3.39–3.31 (m, 1H), 2.21–2.16 (m, 2H), 1.93–1.88 (m, 2H), 1.72–1.68 (m, 1H), 1.63–1.55 (m, 2H), 1.35–1.14 (m, 3H). ^13^C NMR (100 MHz, CDCl_3_): *δ* = 186.5, 166.8, 66.0, 61.4, 49.3, 25.4, 25.2, 24.9. LCMS (ESI): *^t^*R = 2.40 min; peak area = 98.7%; m/z = 318 [M+H]^+^.

### *P. falciparum* growth inhibition assays

*P. falciparum* parasites were cultured *in vitro* in ORh+ human erythrocytes in RPMI 1640 (cat # R8758; Sigma) supplemented with 10% heat inactivated human serum and 5 mg/L gentamycin at 37°C in 5% O_2_ and 5% CO_2_ in N_2_, essentially as described previously.^44^ *In vitro* growth inhibition was assessed in 48 h (one developmental cycle) and 96 h assays (two developmental cycles) against *P. falciparum* parasitized erythrocytes using [^3^H]-hypoxanthine incorporation, as previously described.^45^ Assays were performed with synchronous early ring cultures at a 1% parasitemia and 1% haematocrit for 48 h assays and 0.1% parasitemia and 2% haematocrit for 96 h assays. [^3^H]-hypoxanthine incorporation was determined using a Perkin Elmer/Wallac Trilux 1450 MicroBeta scintillation counter following the harvesting of cultures onto glass fibre filter mats. Percentage growth inhibition was calculated relative to matched vehicle controls included in each assay and IC_50_ values were determined via log-linear interpolation of inhibition curves^46^. In each case, at least two independent assays were performed, each in triplicate, and data shown as mean (± SD).

IPP rescue experiments performed using 96 h assays, as described above. In each case two parallel assays were performed, with one replica assay supplemented with 200 µM IPP^21^ and the other without. Clindamycin was included as a positive delayed death control compound and chloroquine as a negative control. Two independent assays were performed, each in triplicate wells and percentage growth inhibition and IC_50_ values determines as described above.

### *In vitro* cytotoxicity assays

Cytotoxicity was assessed against human neonatal foreskin fibroblasts (NFFs), as described previously,^47^ in at least two independent experiments, with each test carried out in triplicate. Percentage growth inhibitions were calculated relative to matched vehicle controls included in each assay (taken as 100% growth) and IC_50_ values calculated using log linear interpolation of inhibition curves.^46^

### *In vitro* stability and permeability

The permeability of the test compounds across confluent and differentiated Caco-2 cell monolayers was assessed in the apical to basolateral (A-B) direction only using aqueous transport buffer (pH7.4), as previously described.^48^

### Pharmacokinetics

Pharmacokinetic animal studies were conducted at the Centre for Drug Candidate Optimisation, Monash University, using established procedures in accordance with the Australian Code of Practice for the Care and Use of Animals for Scientific Purposes, and the study protocols were reviewed and approved by the Monash Institute of Pharmaceutical Sciences Animal Ethics Committee. The systemic exposure of test compounds was carried out in non-fasted female BALB/c mice (15-21 g; n=3 per timepoint) with access to food and water ad libitum. Compounds were suspended in 10% (v/v) DMSO in olive oil and administered orally via gavage (100 mg/kg; 100 µL/dose). No adverse events were observed. Blood samples were collected via submandibular bleed (∼120 μL) at 15 min and 30 min and 1, 2, 4, 7.5 and 24 h post dose into polypropylene Eppendorf tubes containing heparin as anticoagulant and a stabilisation cocktail (containing Complete^®^ (a protease inhibitor cocktail) and potassium fluoride) to minimise the potential for *ex vivo* compound degradation in blood/plasma samples. Once collected, blood samples were centrifuged immediately, supernatant plasma was removed and stored at -80°C until analysis by liquid chromatography-mass spectrometry (LC–MS). The plasma concentration versus time profile was defined by the average plasma concentration at each sample time, and PK parameters were calculated using non-compartmental methods (PKSolver Version 2.0).

### *In vivo* evaluation of compound tolerability

*In vivo* tolerability was assessed in female BALB/c mice (18-20 g; food and water *ad libitum*) using an escalating dose protocol. Mice were given a single dose of **1** per oral using a gavage needle (100 µL/dose in 10% DMSO: 90% olive oil) beginning at 1.6 mg/kg, then escalating to 16 mg/kg and 116 mg/kg (at this concentration as a suspension). Three mice were assessed for each dose alongside a control mouse that received vehicle only (10% DMSO: 90% olive oil). After each dose, mice were monitored for seven days (clinical signs and weight) and liver and kidney pathology assessed on day seven following euthanasia (Cerberus Sciences, Australia). Following this, tolerability was carried out in single and twice daily dosing at 116 mg/kg for 3 days and clinical signs and pathology assessed as above. Protocols were performed according to a Griffith University Animal Ethics Committee approved project (#ESK/01/17/AEC) and were a requirement before proceeding to *in vivo* efficacy studies in a murine malaria model.

### Murine model of malaria

*In vivo* efficacy was evaluated in female BALB/c mice (18-20 g; food and water *ad libitum*) infected with 1x 10^5^ *P. berghei* ANKA infected erythrocytes taken from a passage mouse. Mice were dosed orally via gavage (100 µL/dose) with vehicle control (10% DMSO: 90% olive oil) or 116 mg/kg/dose **1** suspended in vehicle. Mice were dosed for three days, beginning 2 h post infection (p.i.), either once daily (qd) or twice daily (bid). Chloroquine (10 mg/kg in PBS; single daily doses) was included as positive control. Mice were monitored daily for clinical signs of infection and peripheral blood parasitemia was assessed from day 3-4 p.i. via microscopic analysis of DiffQuik^®^ (POCD Healthcare, Australia) stained thin blood smears prepared via tail snip. Parasitemia was calculated as the mean number of parasites/100 erythrocytes for ∼1,000 (or greater) erythrocytes for each mouse and timepoint. Mice were euthanised using a scorecard of clinical signs, or at ∼15–25% parasitemia, or when weight loss compared to infection day exceeded ∼15%. Protocols were performed according to a Griffith University Animal Ethics Committee approved project (#ESK/02/17/AEC).

### Supporting Information

1. LCMS traces for key target compounds (with in vivo data). S2–S5
2. NMR spectra for example intermediates and all tested compounds. S6–S81

Authors will release the atomic coordinates and experimental data upon article publication.

**Corresponding Author Information:** Katherine T. Andrews-; John H. Ryan -; Andrew G. Riches -; Tina S. Skinner-Adams -

**Present/Current Author Addresses:** Antoine Masurier - Laboratoire Gulliver (CNRS UMR 7083), ESPCI, Bât. F/G, 10 rue Vauquelin, 75005 Paris, France; Alexandros A. Mouratidis - Akesa Pharma, Level 6/141 Flinders Lane, Melbourne, Victoria 3000, Australia; Meaghan Firmin - Metro North Health Service, 2 Hospital Rd, Biloela, Queensland 4715, Australia.

### Author Contributions

KTA, TSA, JHR, AR and GMF contributed equally to this work. The manuscript was written through contributions of all authors. All authors have given approval to the final version of the manuscript.

## Supporting information

LCMS traces and NMR spertra

## Acknowledgements

We thank the National Health and Medical Research Council of Australia for funding (APP1102365) to KTA, TSA, OH, JHR and AD. AR was supported by Griffith University GUIPRS and GUPRS scholarships. We thank the Australian Red Cross Lifeblood for the provision of human blood and sera for culture of *Plasmodium* parasites and the Centre for Drug Candidate Optimisation (CDCO), Monash University, for ADME/PK work, which is partially supported by the Monash University Technology Research Platform network and Therapeutic Innovation Australia (TIA) through the Australian Government National Collaborative Research Infrastructure Strategy (NCRIS) program. We are grateful for the provision of compounds by the Griffith Institute for Drug Discovery (GRIDD) Compounds Australia facility, which is partially supported by TIA through the Australian Government NCRIS program.

## Notes

### Competing Interest Statement

The authors have declared no competing interest.

