## Supplementary material for "Discovery of 1,3,4-oxadiazoles with slow-action activity against *Plasmodium falciparum* malaria parasites": LCMS traces and NMR spertra

[d]Medicines for Malaria Venture, Geneva, Switzerland.

[*] Equal contributions

### Co-corresponding Authors

**Table of Contents**

1. LCMS traces for key target compounds (with in vivo data). S2–S5

2. NMR spectra for example intermediates and all tested compounds. S6–S81

**1.** LCMS traces for key target compounds (with in vivo data).

**Figure S1**. LCMS trace and report for compound **1**

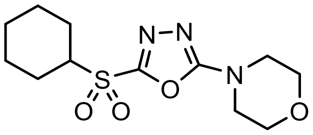

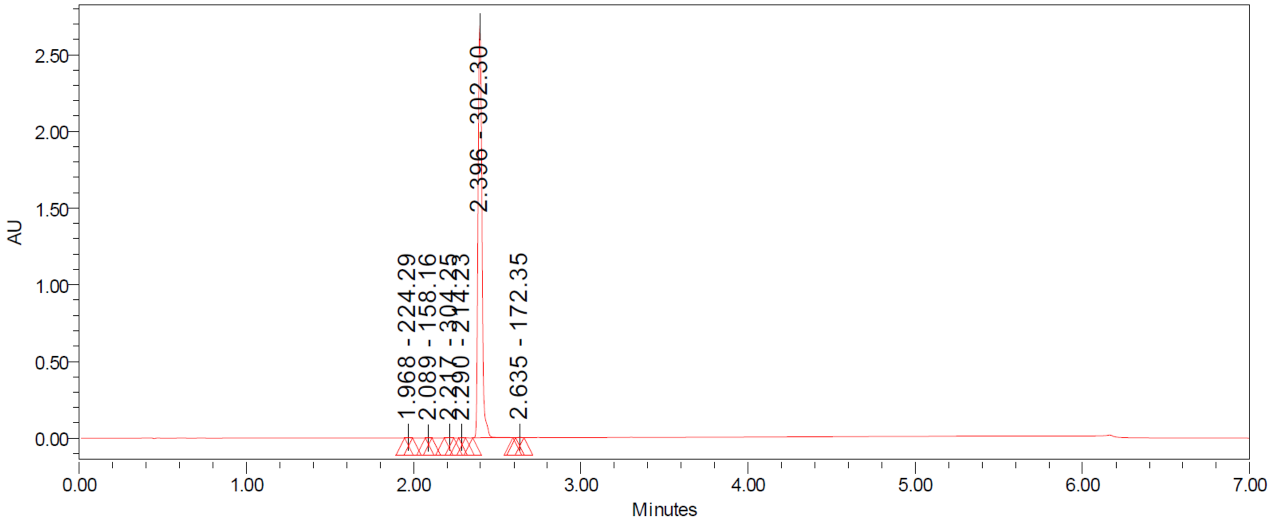


| | RT | Area | % Area | Height | Base Peak (m/z) |
|---|---|---|---|---|---|
| 1 | 1.968 | 2833 | 0.06 | 2287 | 224.29 |
| 2 | 2.089 | 907 | 0.02 | 767 | 158.16 |
| 3 | 2.217 | 3193 | 0.07 | 2407 | 304.25 |
| 4 | 2.290 | 2386 | 0.05 | 2102 | 214.23 |
| 5 | 2.396 | 4378029 | 99.72 | 2691065 | 302.30 |
| 6 | 2.635 | 2809 | 0.06 | 2107 | 172.35 |

**Figure S2**. LCMS trace and report for compound **8**

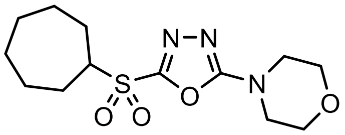

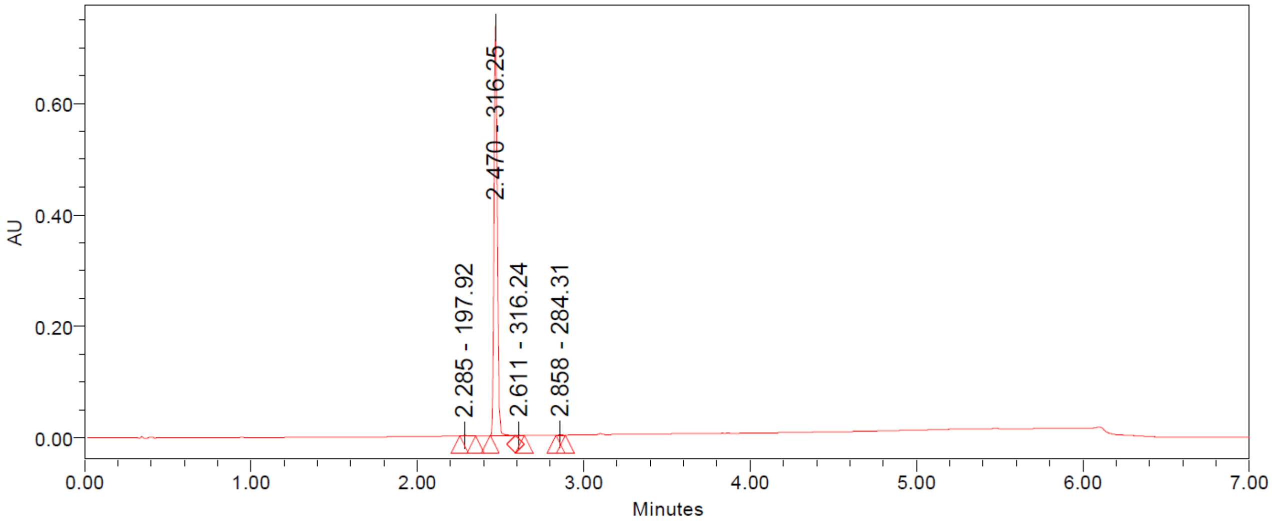


| | RT | Area | % Area | Height | Base Peak (m/z) |
|---|---|---|---|---|---|
| 1 | 2.285 | 1420 | 0.15 | 690 | 197.92 |
| 2 | 2.470 | 972519 | 99.67 | 733087 | 316.25 |
| 3 | 2.611 | 892 | 0.09 | 670 | 316.24 |
| 4 | 2.858 | 935 | 0.10 | 700 | 284.31 |

**Figure S3**. LCMS trace and report for compound **11**

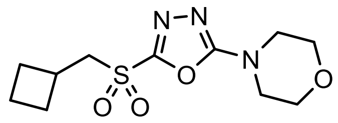

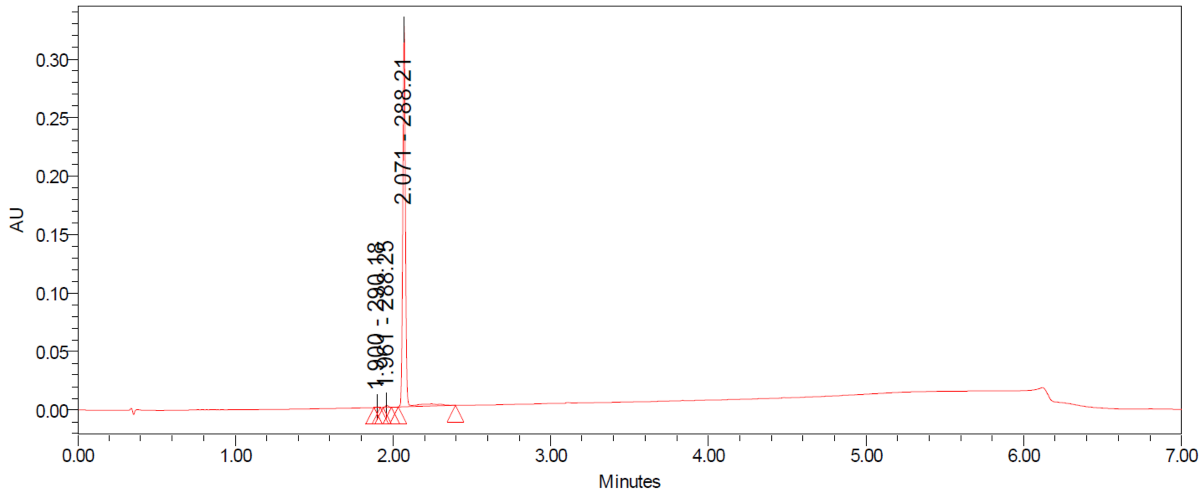


| | RT | Area | % Area | Height | Base Peak (m/z) |
|---|---|---|---|---|---|
| 1 | 1.900 | 511 | 0.13 | 465 | 290.18 |
| 2 | 1.961 | 2067 | 0.51 | 1791 | 288.25 |
| 3 | 2.071 | 400760 | 99.36 | 324793 | 288.21 |

**Figure S4**. LCMS trace and report for compound **33**

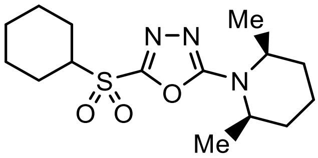


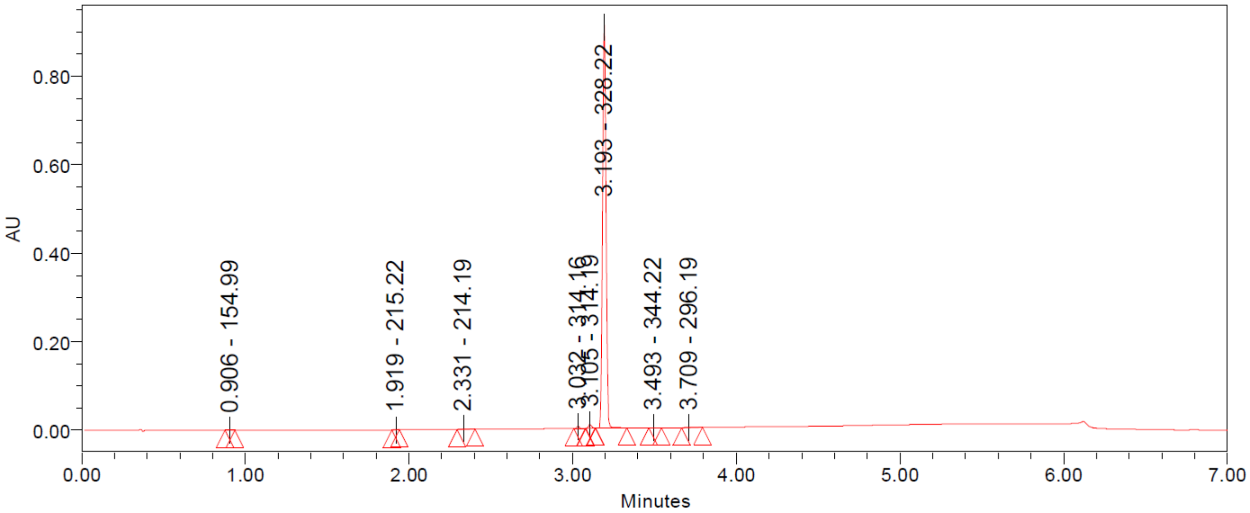


| | RT | Area | % Area | Height | Base Peak (m/z) |
|---|---|---|---|---|---|
| 1 | 0.906 | 1582 | 0.12 | 862 | 154.99 |
| 2 | 1.919 | 359 | 0.03 | 290 | 215.22 |
| 3 | 2.331 | 3286 | 0.25 | 1091 | 214.19 |
| 4 | 3.032 | 5904 | 0.45 | 4262 | 314.16 |
| 5 | 3.105 | 11856 | 0.91 | 8324 | 314.19 |
| 6 | 3.193 | 1282203 | 98.03 | 907583 | 328.22 |
| 7 | 3.493 | 1196 | 0.09 | 548 | 344.22 |
| 8 | 3.709 | 1624 | 0.12 | 622 | 296.19 |

**2.** NMR spectra for example intermediates and all tested compounds.

**Figure S5**. [1]H NMR spectrum (400 MHz, $CDCl_3$) of **65a**

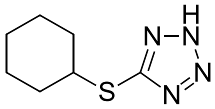

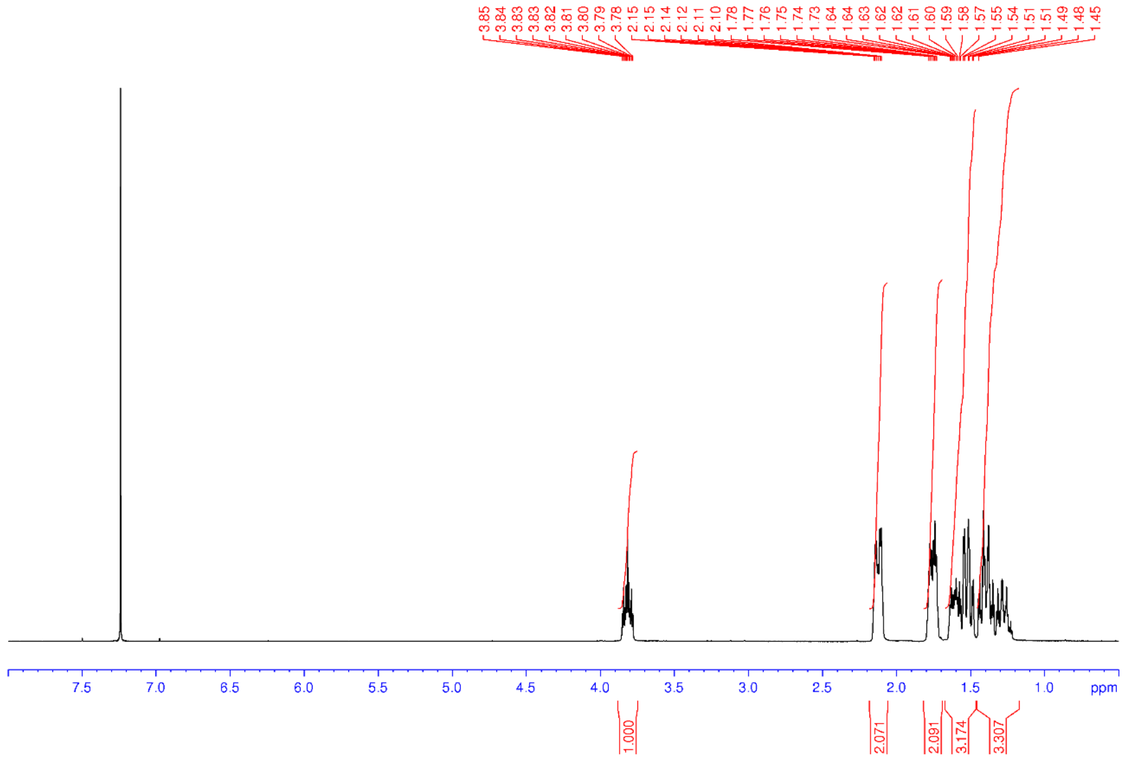

**Figure S6**. $^{13}$c NMR spectrum (100 MHz, $CDCl_3$) of **65a**

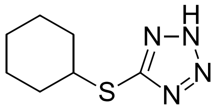


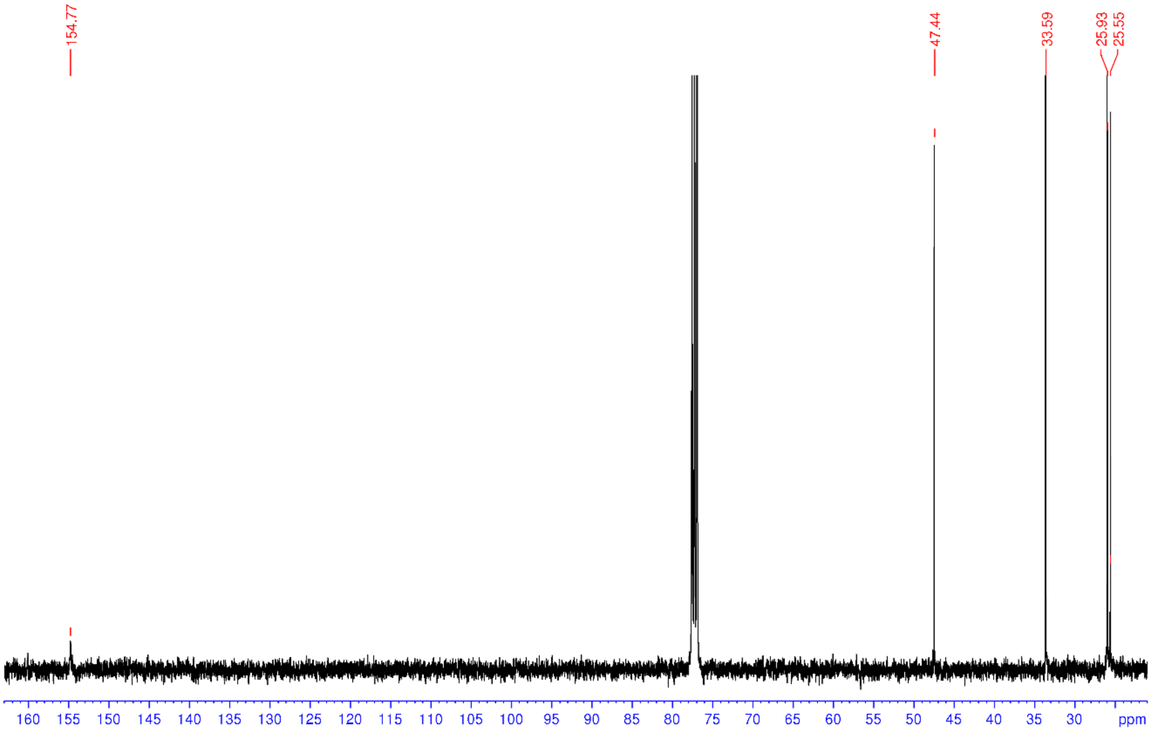

**Figure S7**. $^1$H NMR spectrum (400 MHz, $CDCl_3$) of **68a**

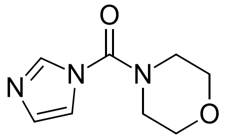


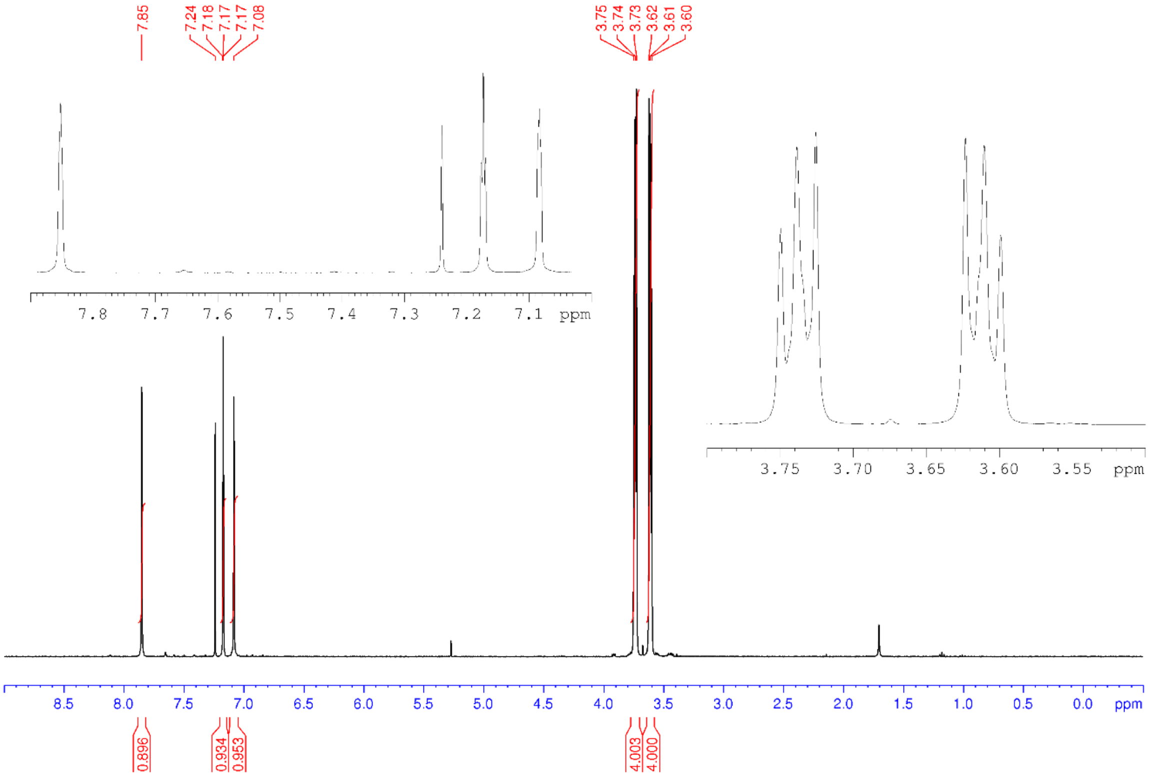

**Figure S8**. $^{1}$H NMR spectrum (400 MHz, $CDCl_3$) of **69aa**

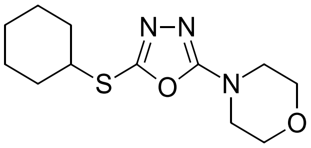


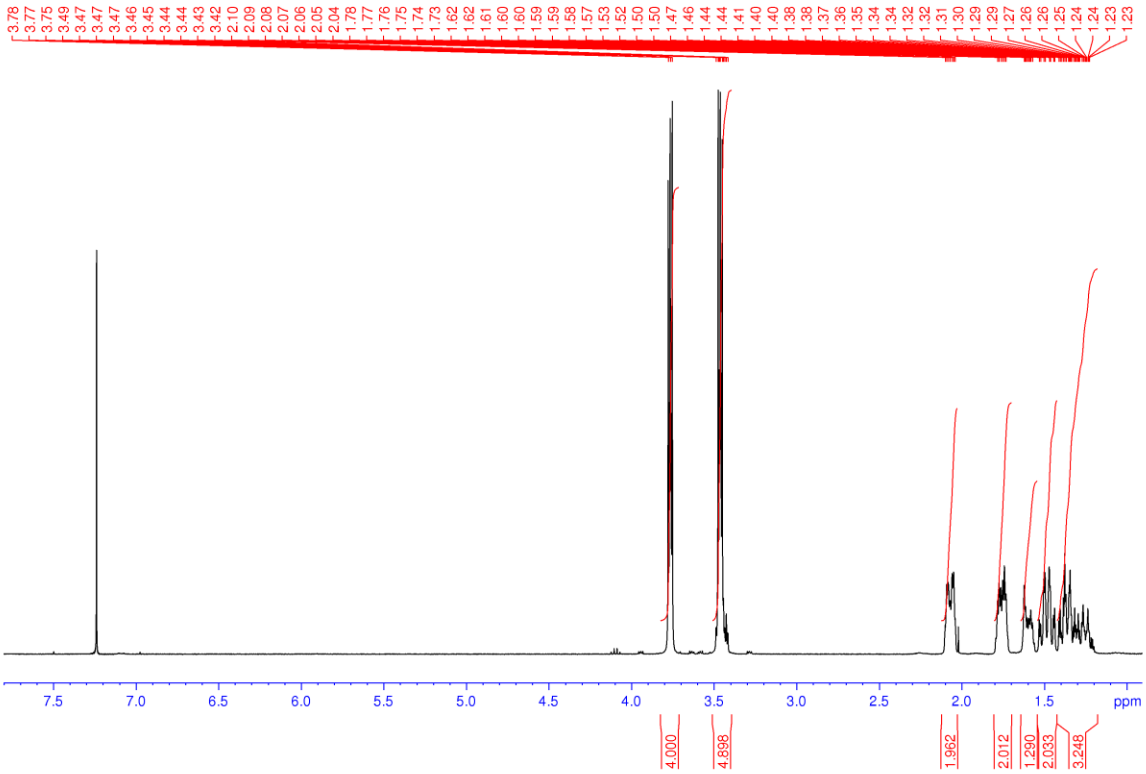

**Figure S9**. $^{13}C$ NMR spectrum (100 MHz, $CDCl_3$) of **69aa**

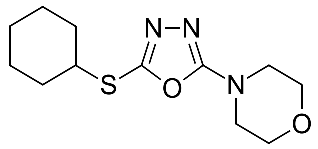

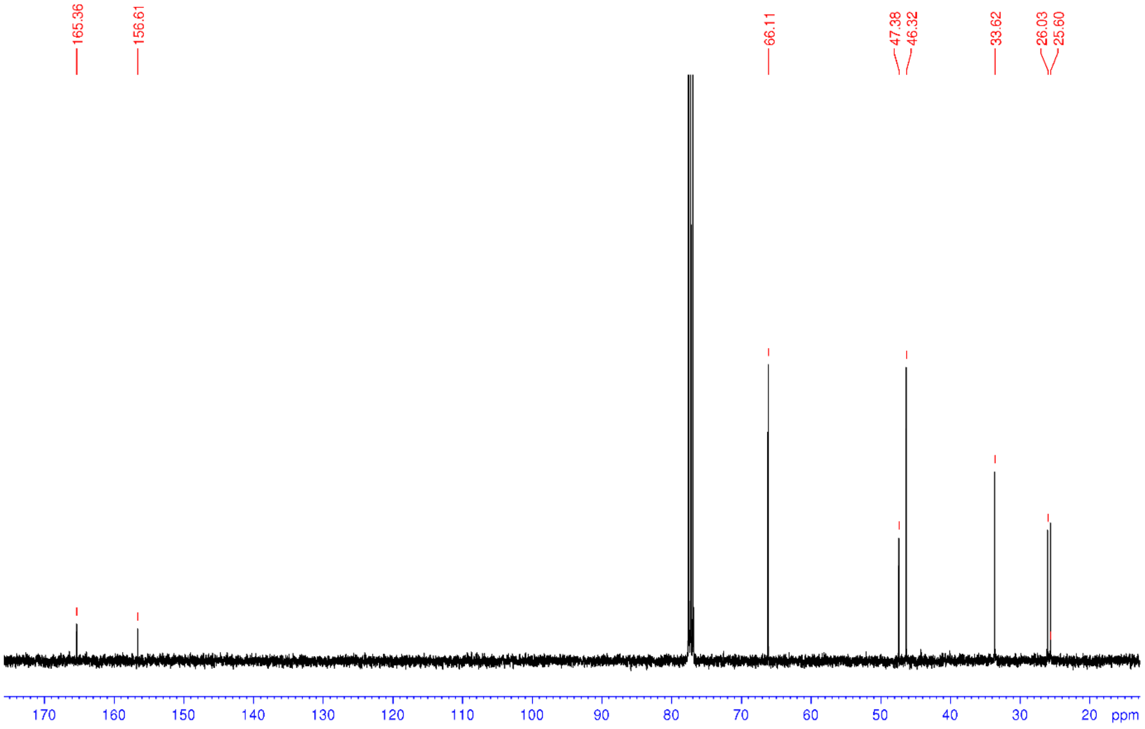

**Figure S10**. $^{1}H$ NMR spectrum (400 MHz, $CDCl_3$) of **1**

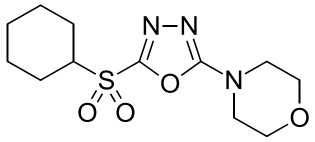

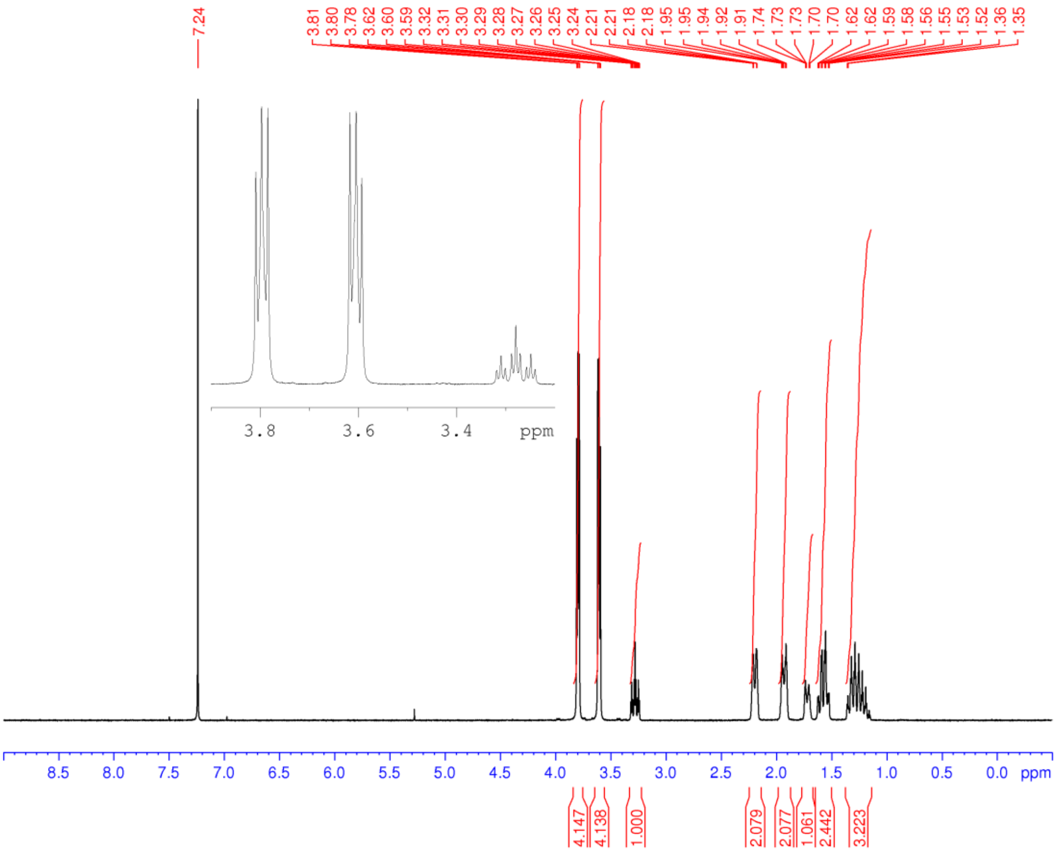

**Figure S11**. $^{13}$C NMR spectrum (100 MHz, $CDCl_3$) of **1**

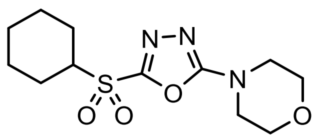


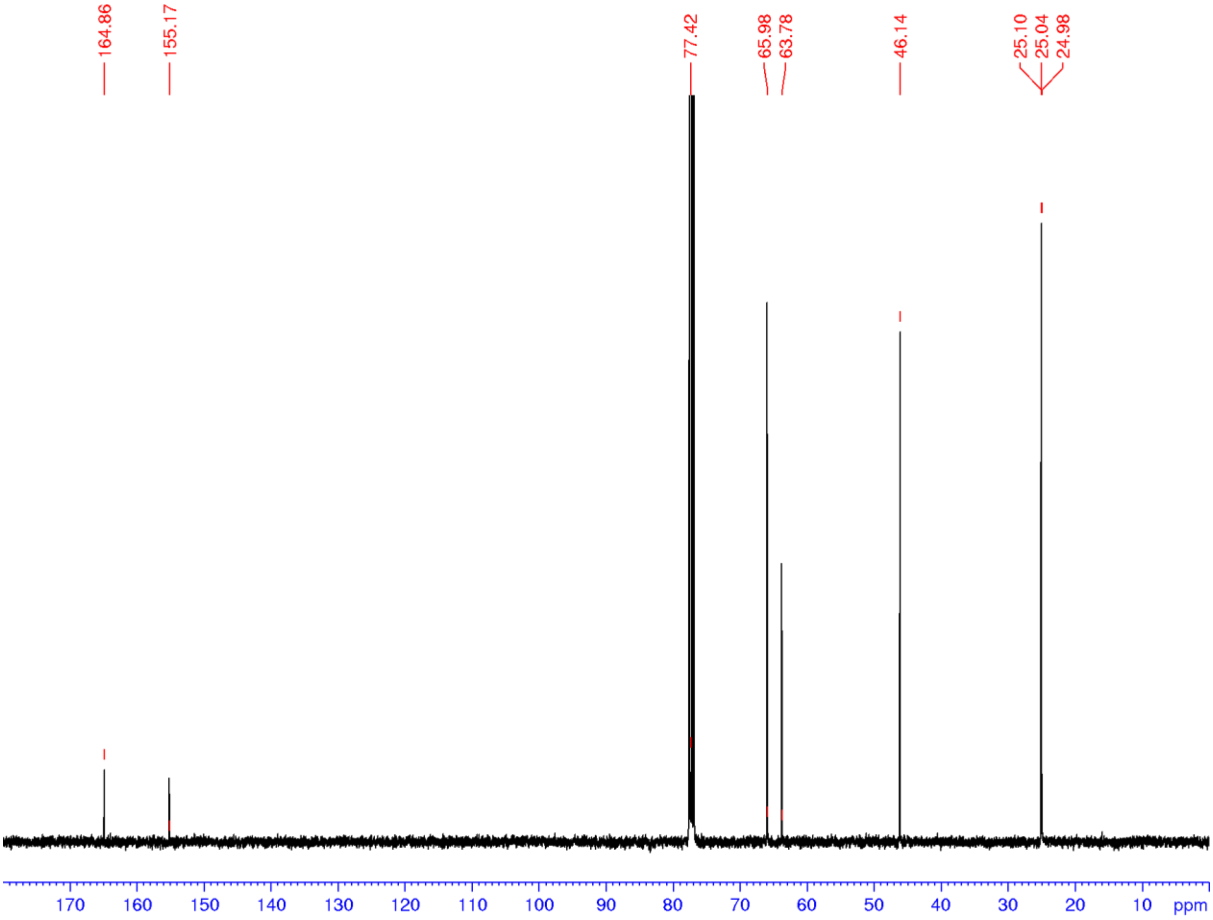

**Figure S12**. $^{1}H$ NMR spectrum (400 MHz, $CDCl_3$) of **2**

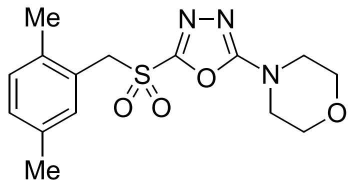

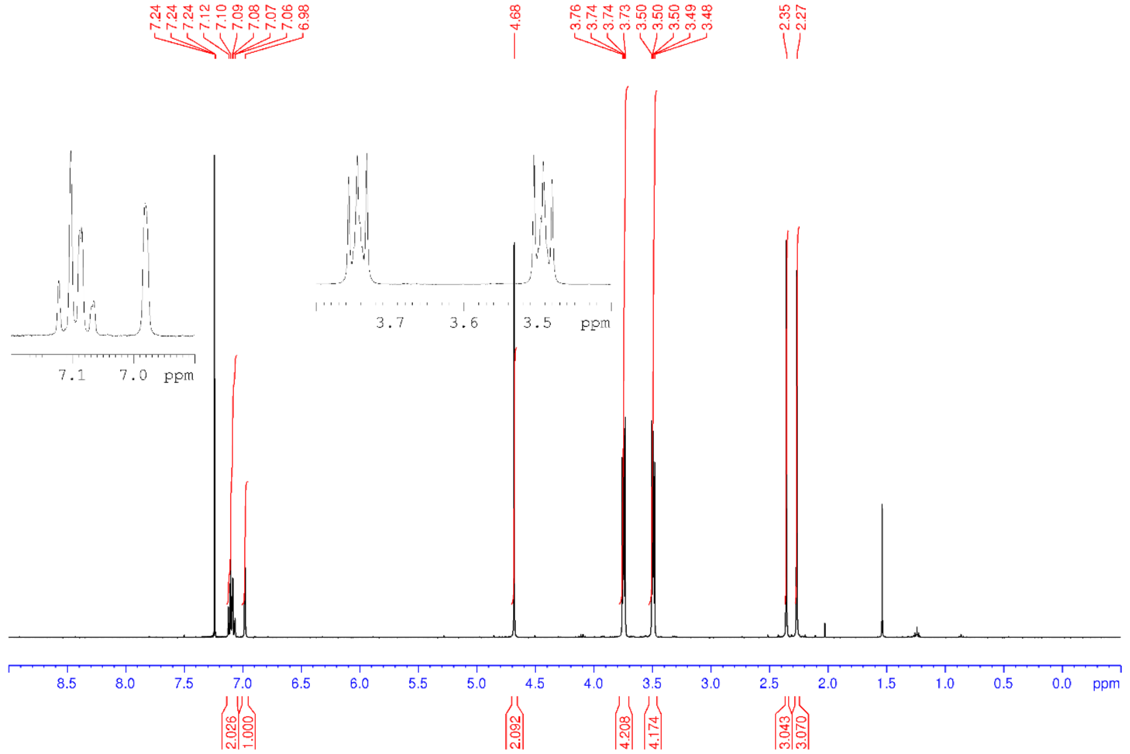

**Figure S13**. $^{13}$C NMR spectrum (100 MHz, $CDCl_3$) of **2**

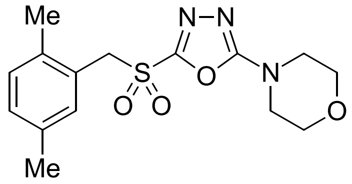

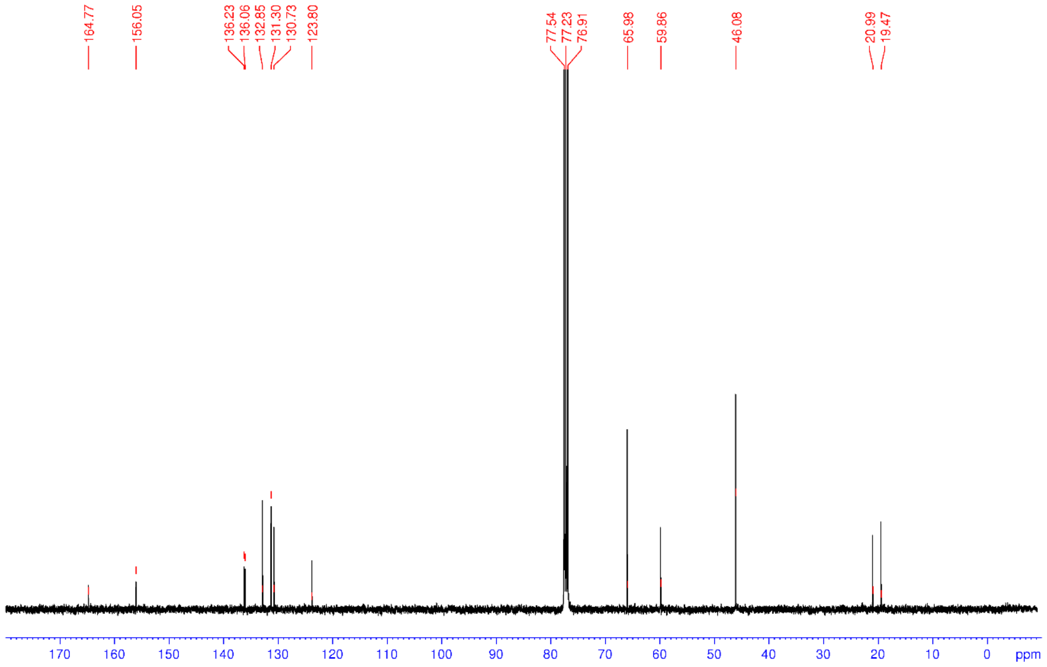

**Figure S14.** $^{1}$H NMR spectrum (400 MHz, $CDCl_3$) of **3**

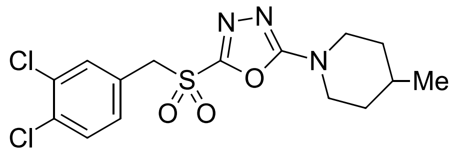


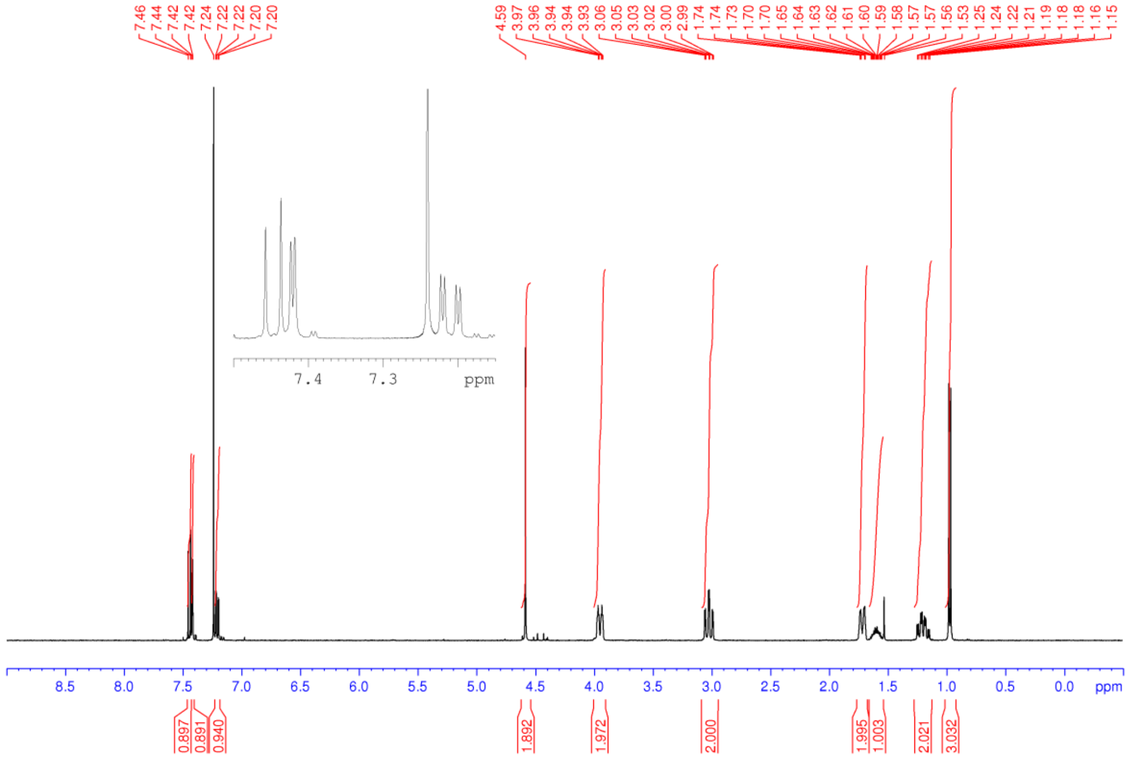

**Figure S15**. $^1$H NMR spectrum (400 MHz, $CDCl_3$) of **4**

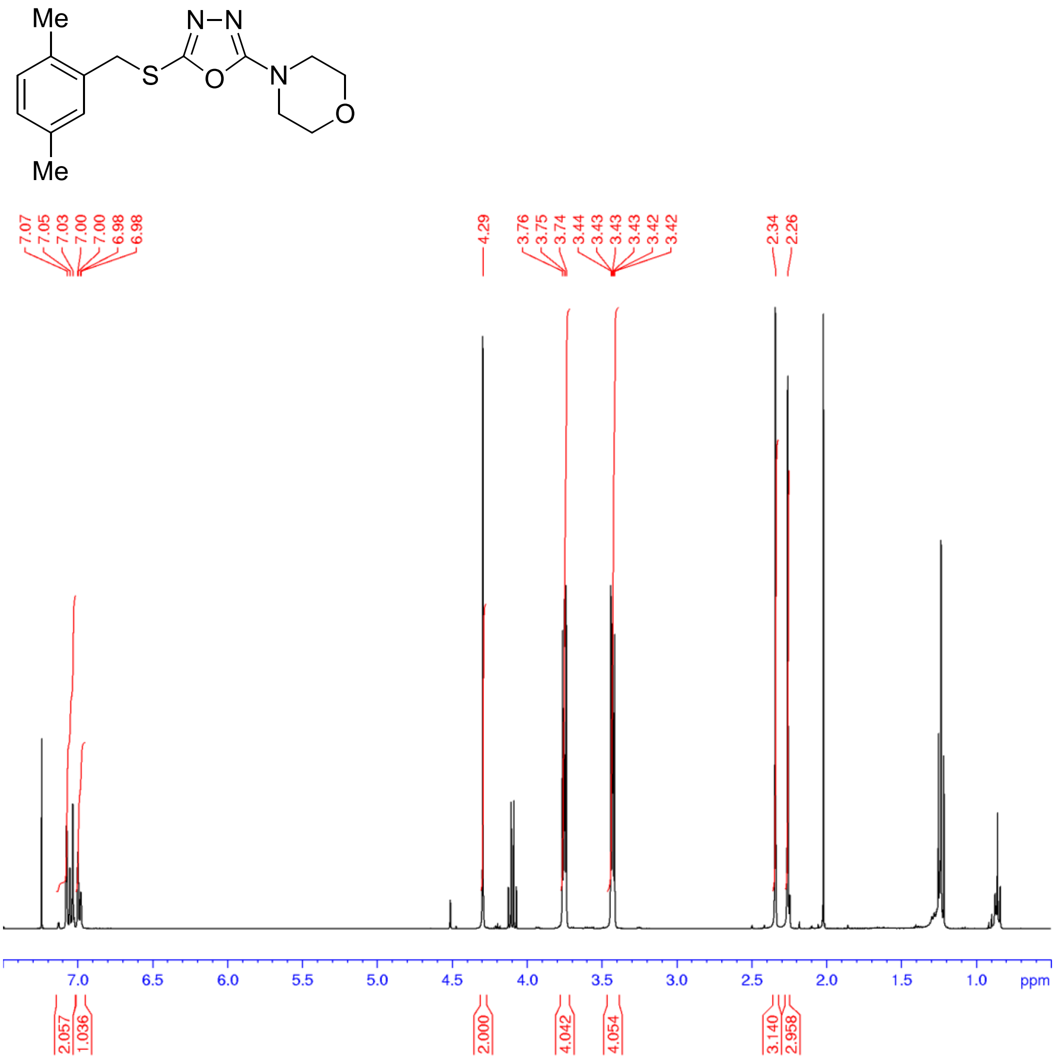

**Figure S16**. $^{13}C$ NMR spectrum (125 MHz, $CDCl_3$) of **4**

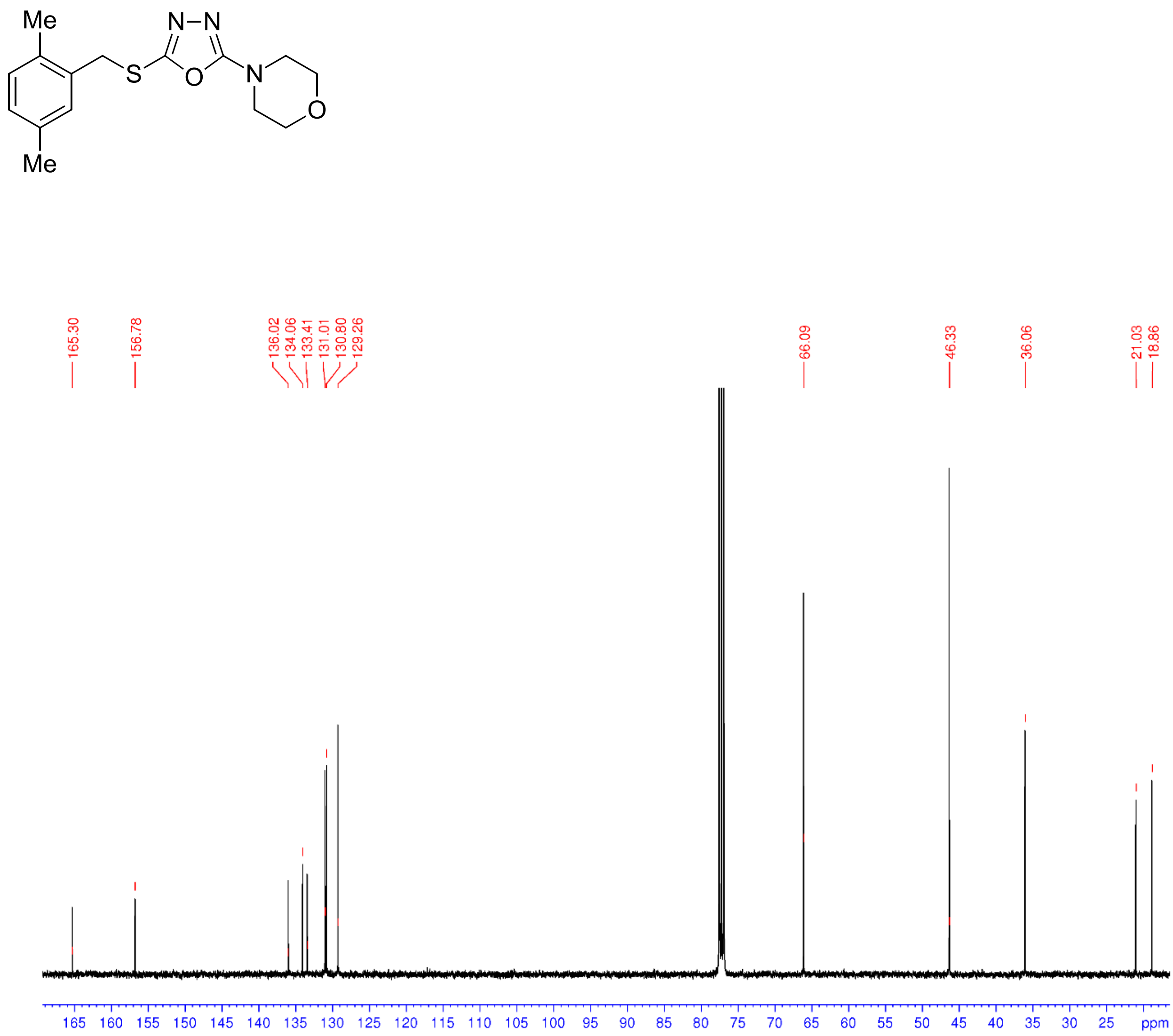

**Figure S17**. $^{1}$H NMR spectrum (400 MHz, $CDCl_3$) of **5**

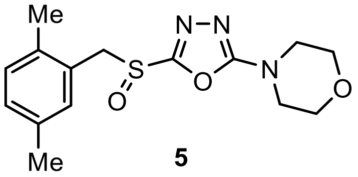


**5**

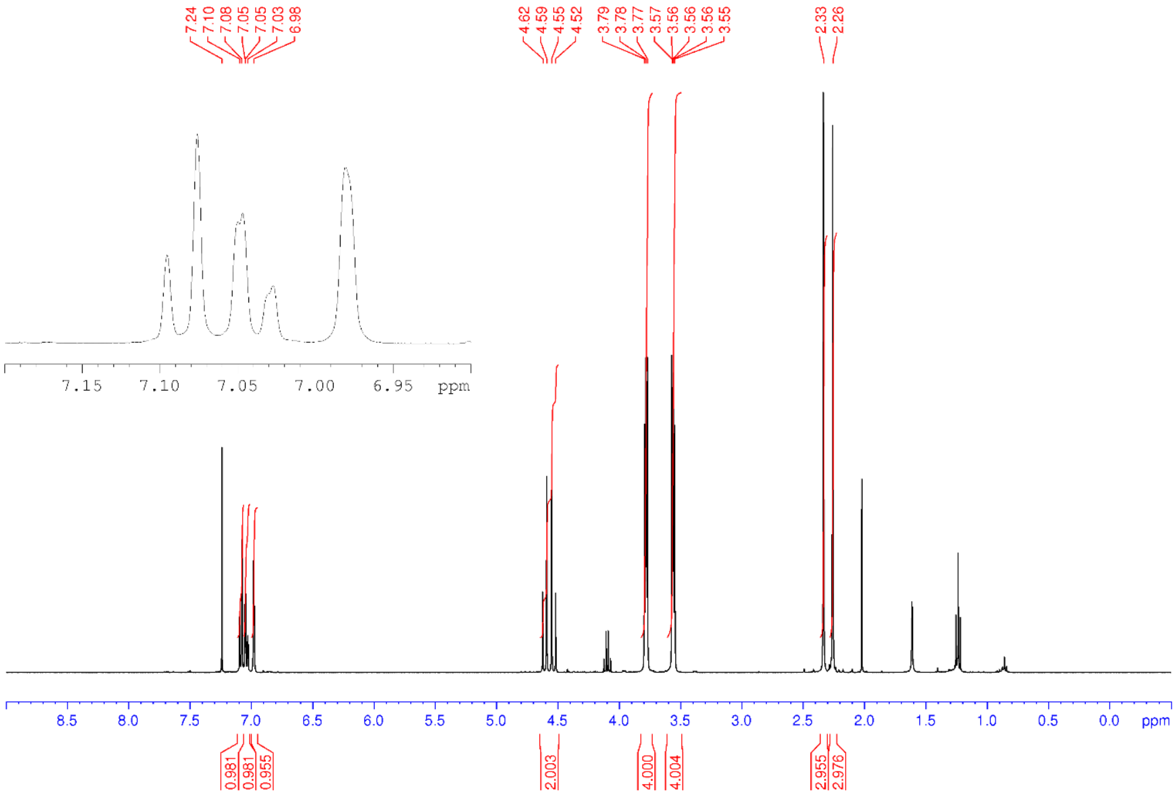

**Figure S18**. $^{13}C$ NMR spectrum (100 MHz, $CDCl_3$) of **5**

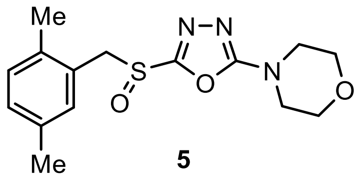


**5**

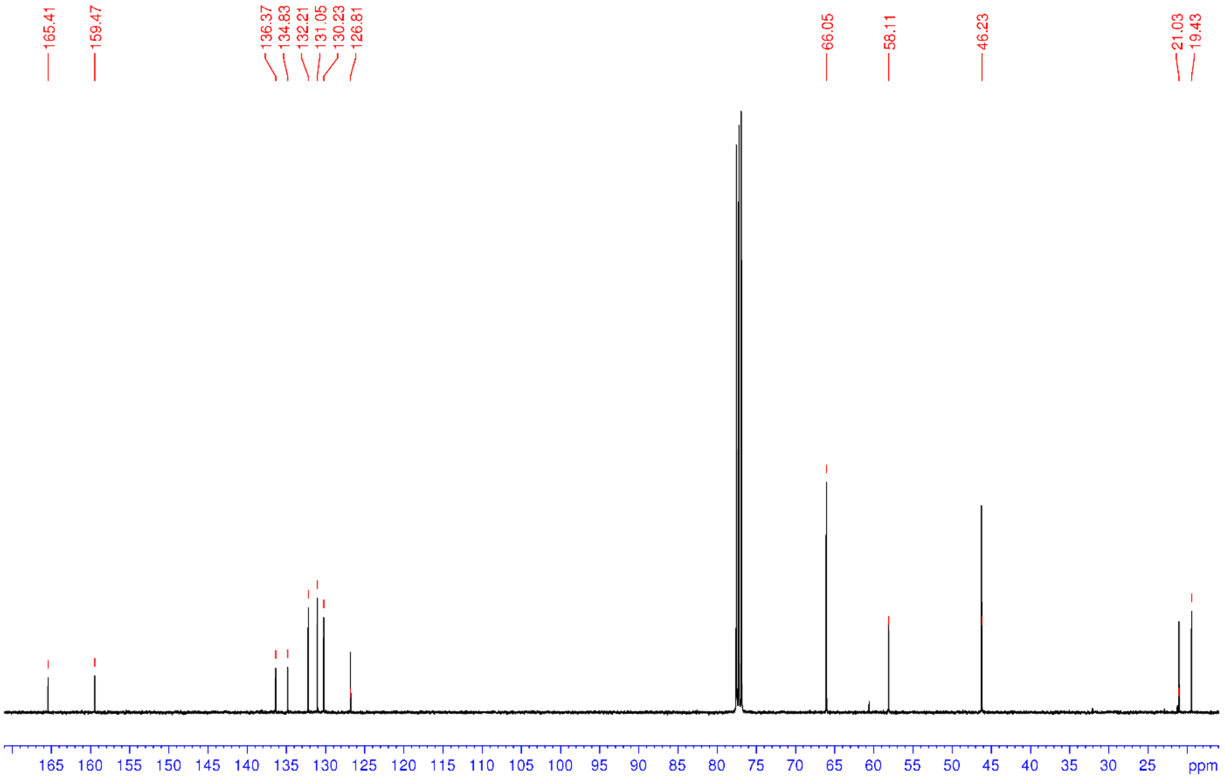

**Figure S19**. [1]H NMR spectrum (400 MHz, $CDCl_3$) of **6**

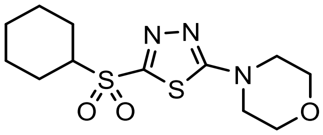

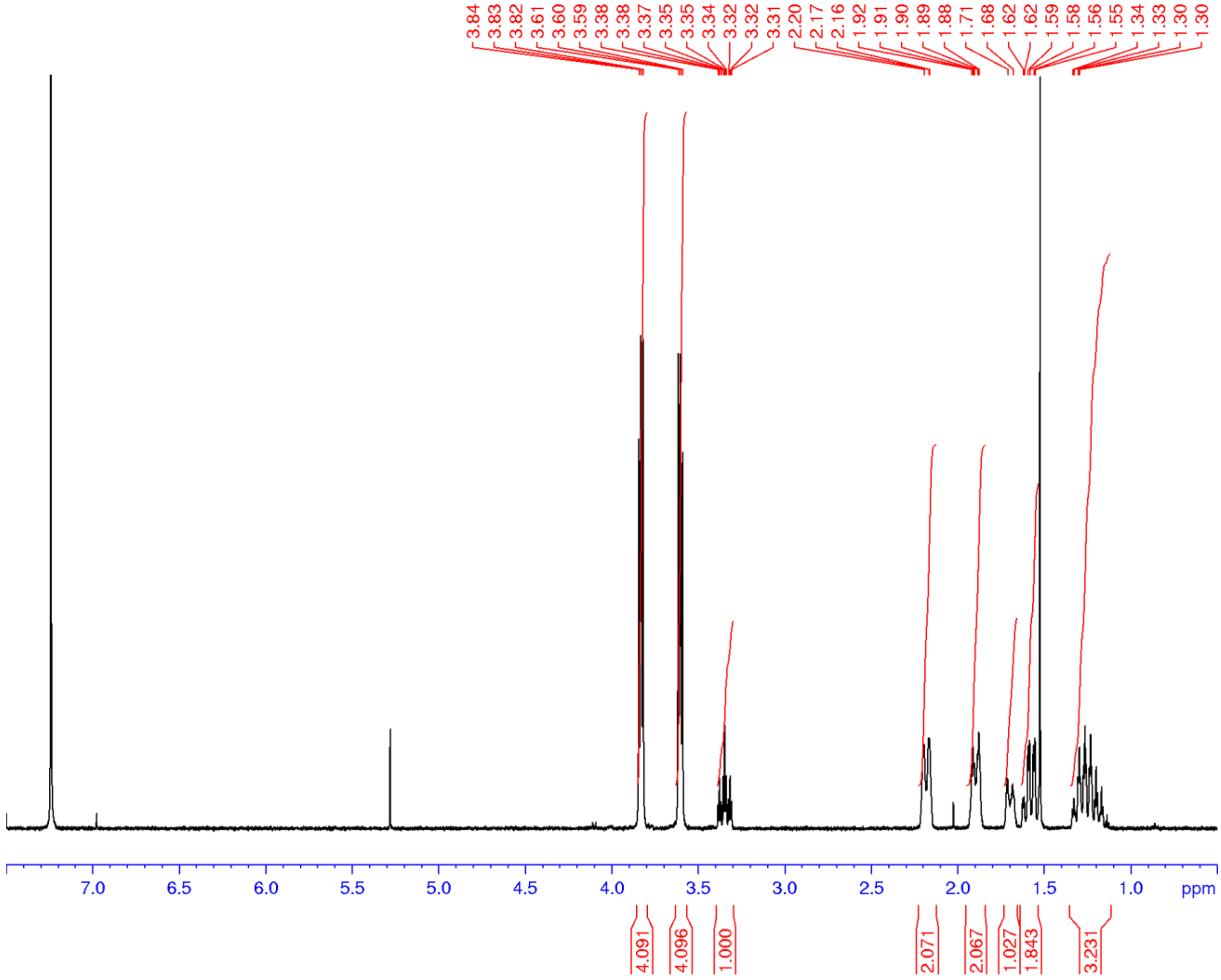

**Figure S20**. $^{13}C$ NMR spectrum (100 MHz, $CDCl_3$) of **6**

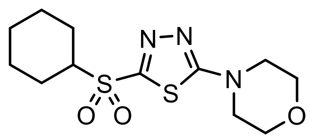

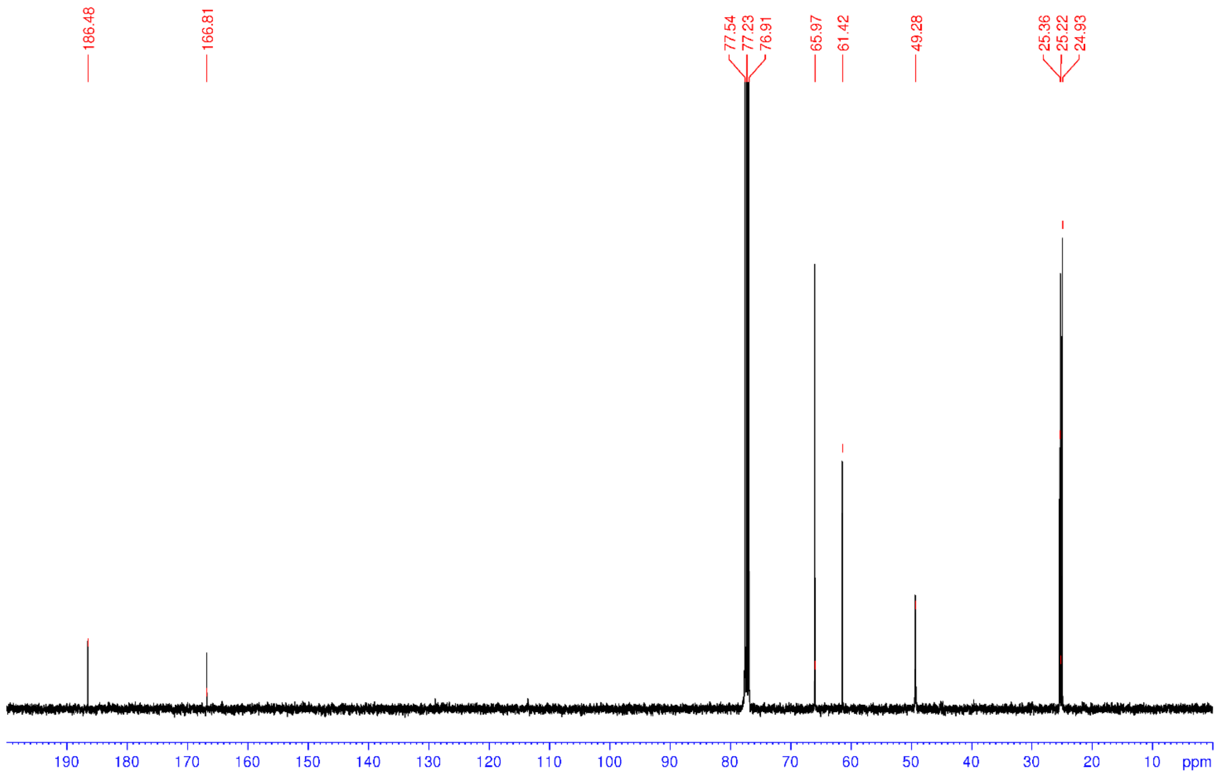

**Figure S21**. [1]H NMR spectrum (400 MHz, $CDCl_3$) of **7**

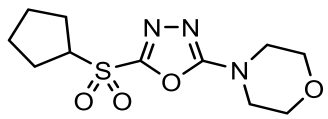

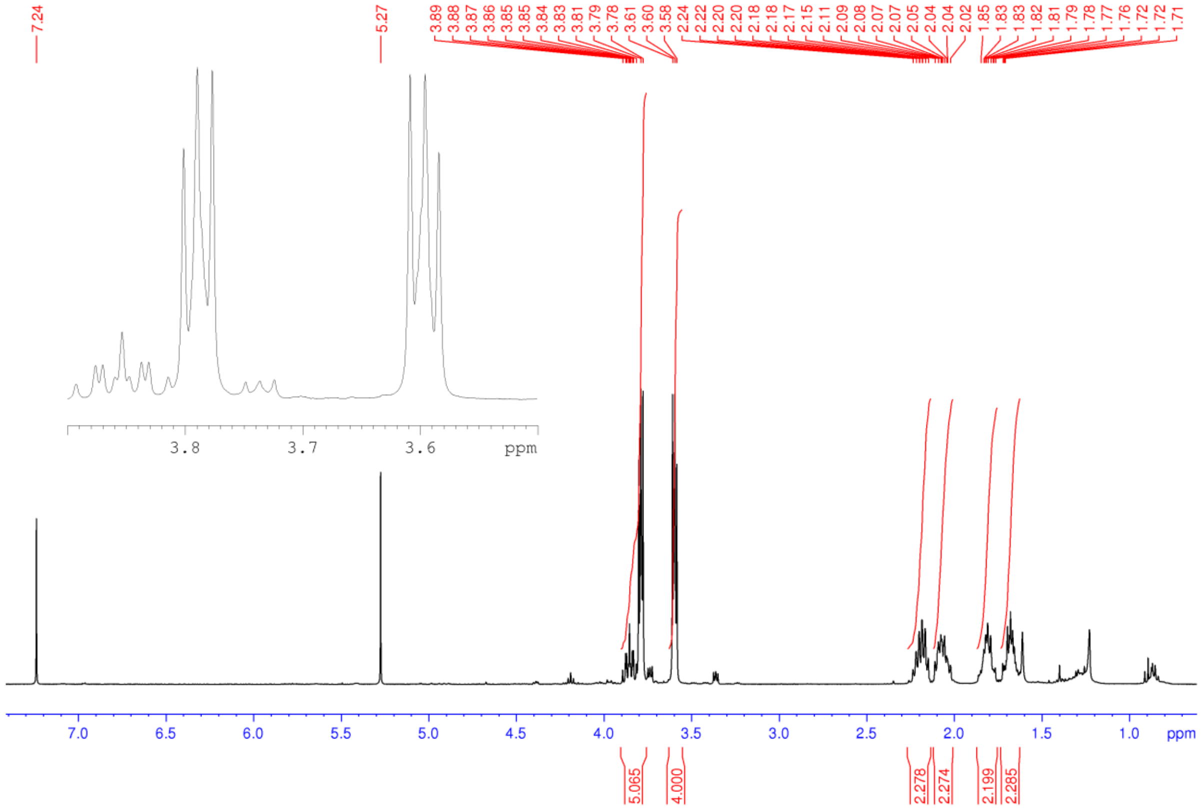

**Figure S22**. $^{1}H$ NMR spectrum (500 MHz, $CDCl_3$) of **8**

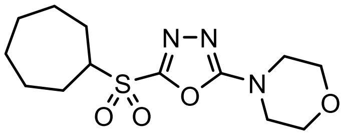

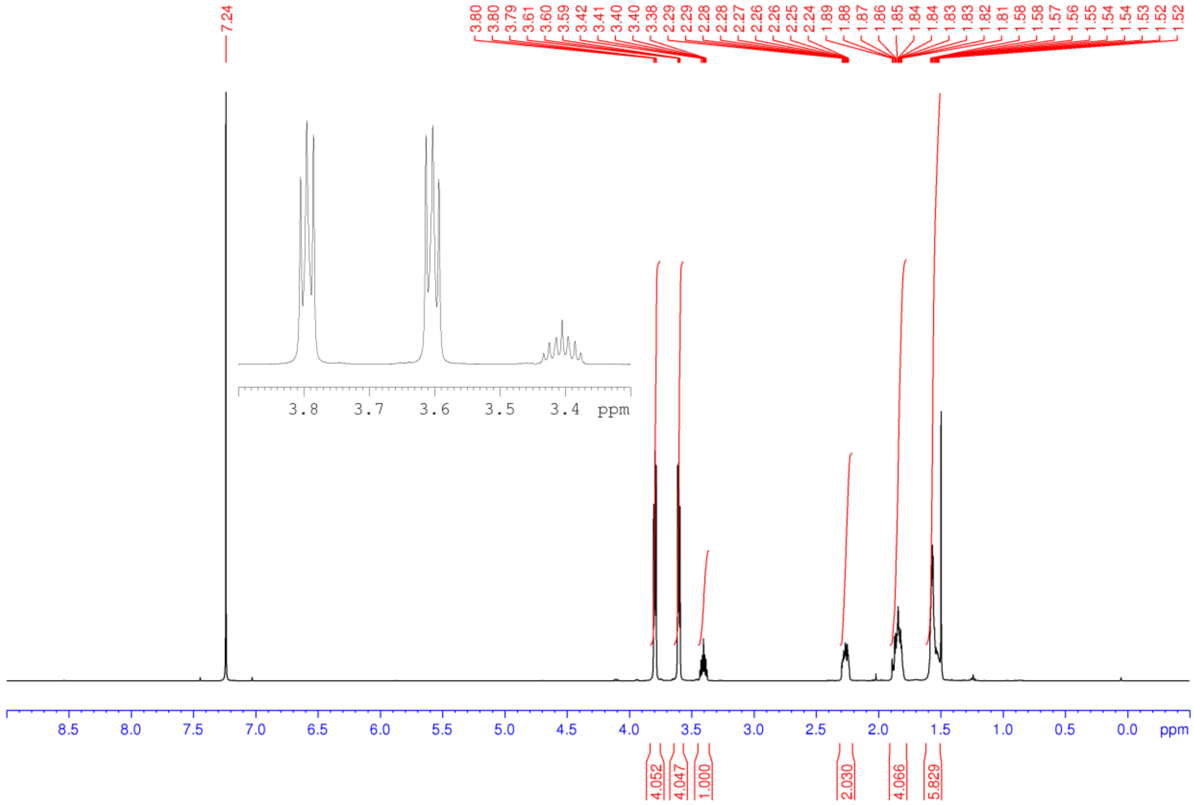

**Figure S23**. [1]H NMR spectrum (400 MHz, $CDCl_3$) of **9**

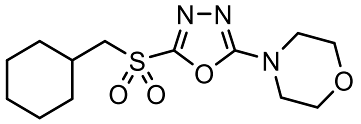

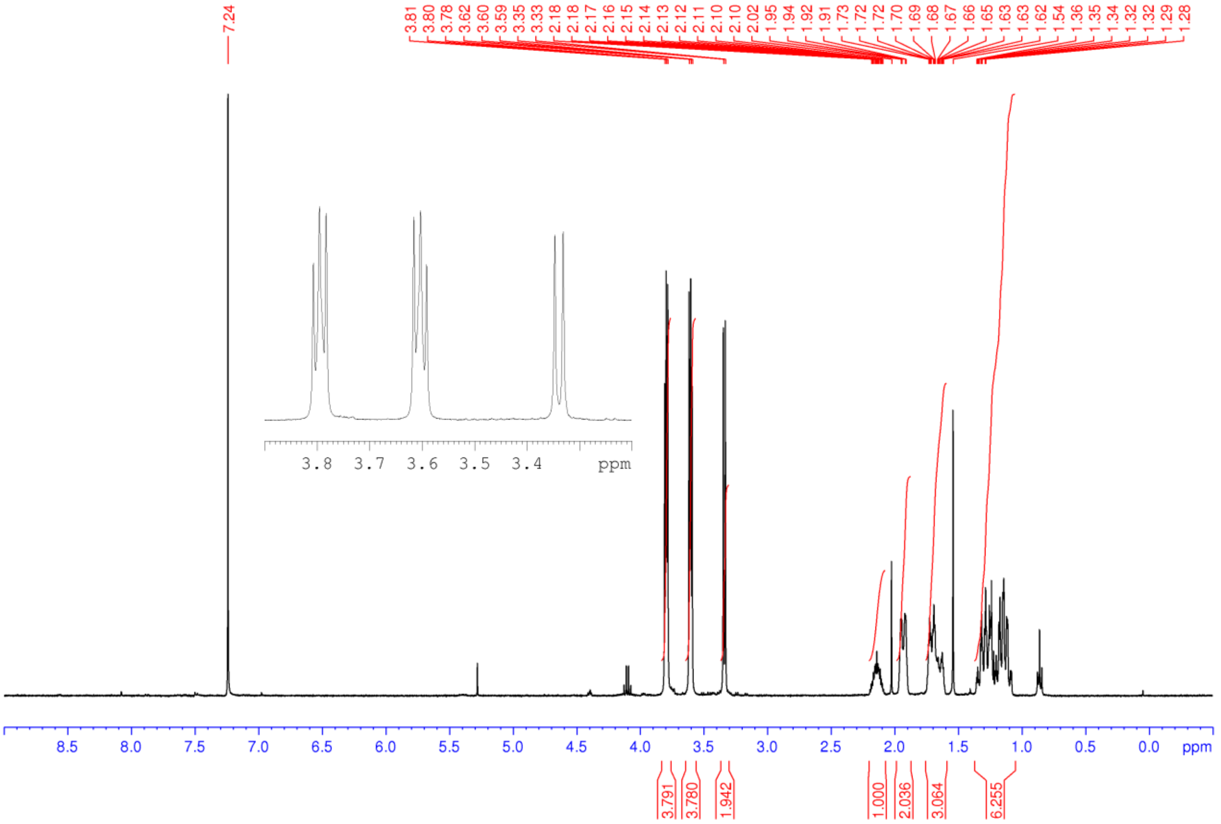

**Figure S24**. $^{1}H$ NMR spectrum (400 MHz, $CDCl_3$) of **10**

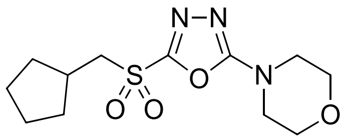

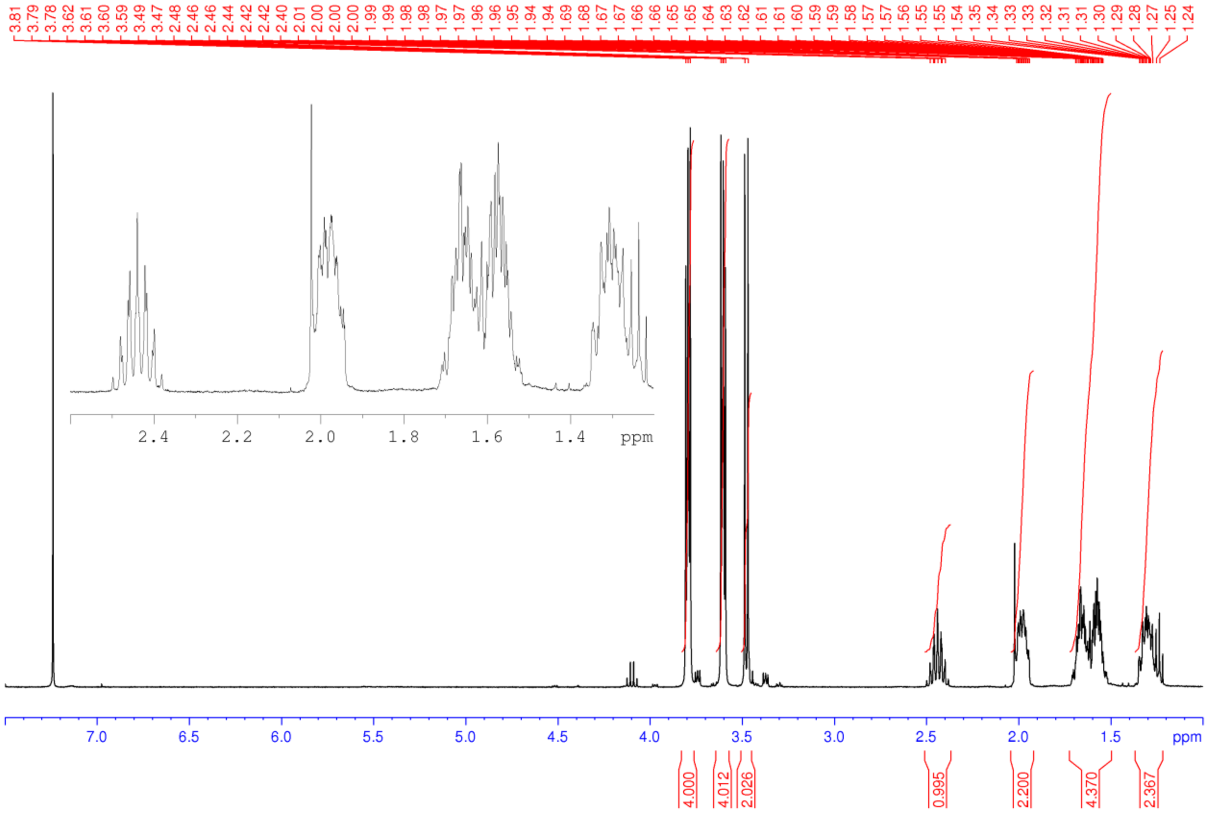

**Figure S25**. [1]H NMR spectrum (400 MHz, $CDCl_3$) of **11**

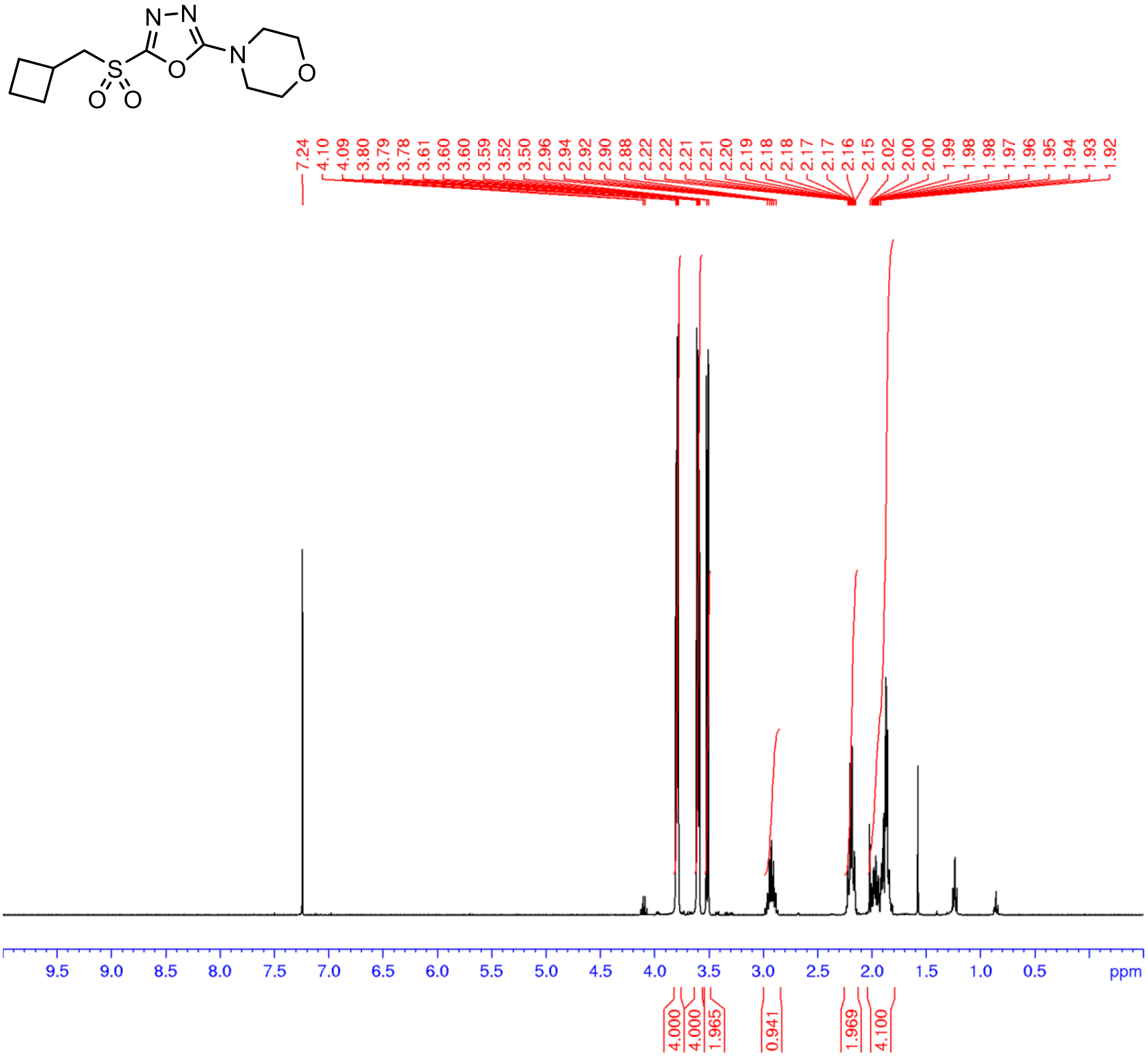


**Figure S26.** [13]C NMR spectrum (100 MHz, $CDCl_3$) of **11**

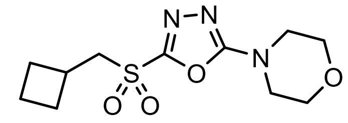
N
N
S
O
O
O
N
O

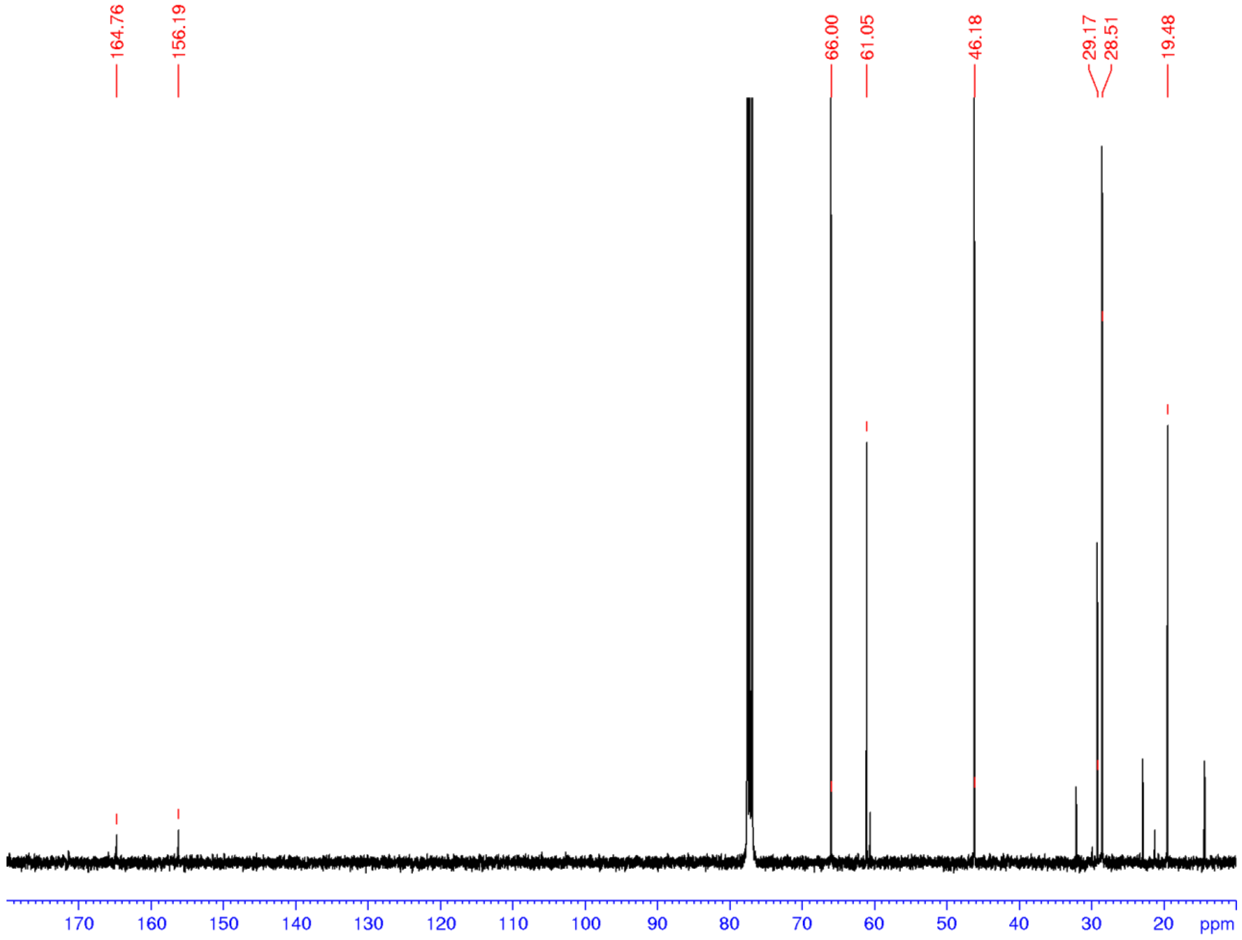
164.76
156.19
66.00
61.05
46.18
29.17
28.51
19.48
170
160
150
140
130
120
110
100
90
80
70
60
50
40
30
20
ppm

**Figure S27.** $^1$H NMR spectrum (400 MHz, $CDCl_3$) of **12**

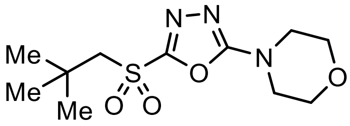


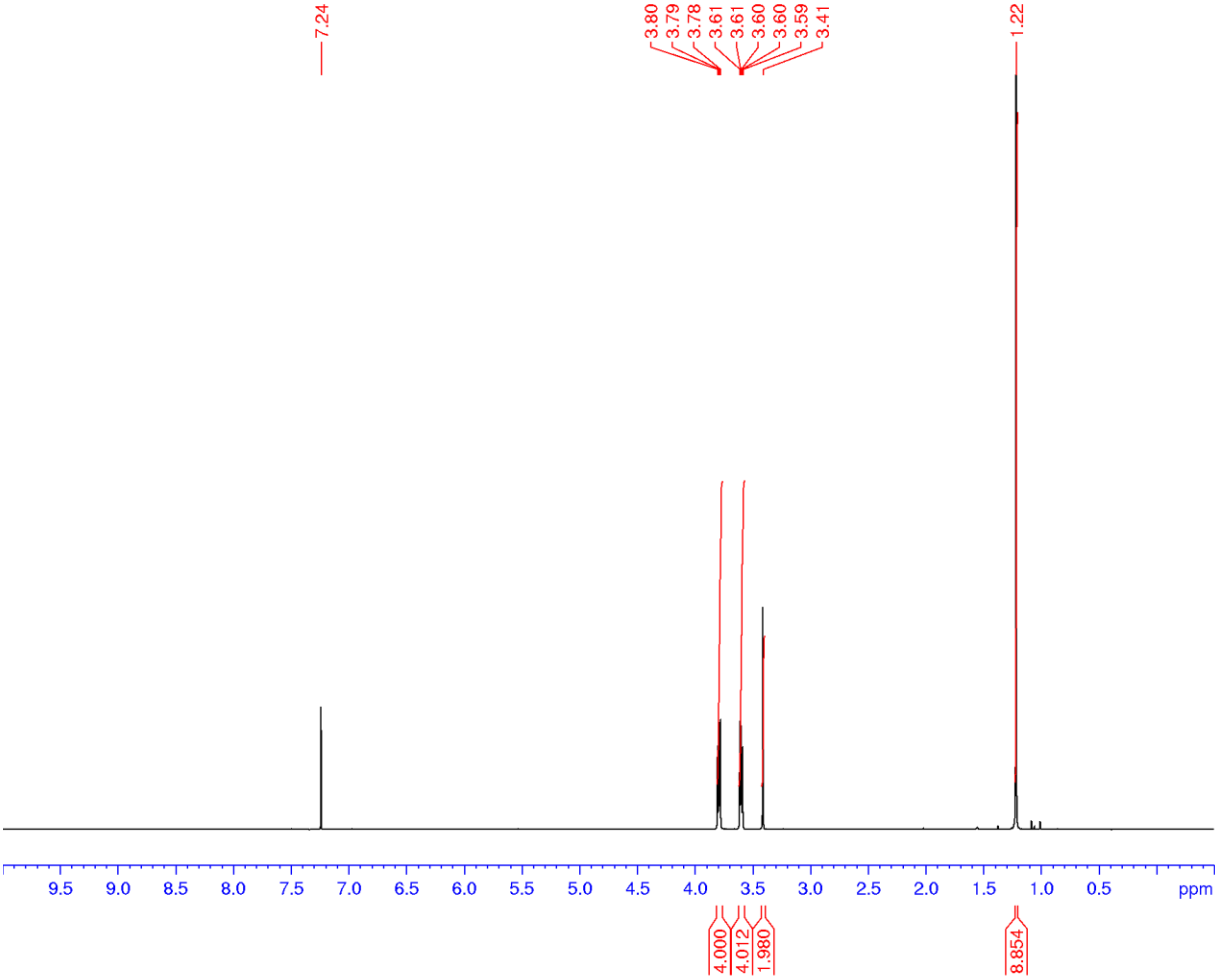

**Figure S28.** $^{1}$H NMR spectrum (400 MHz, $CDCl_3$) of **13**

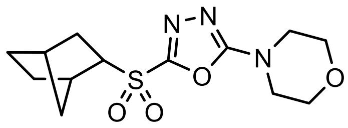

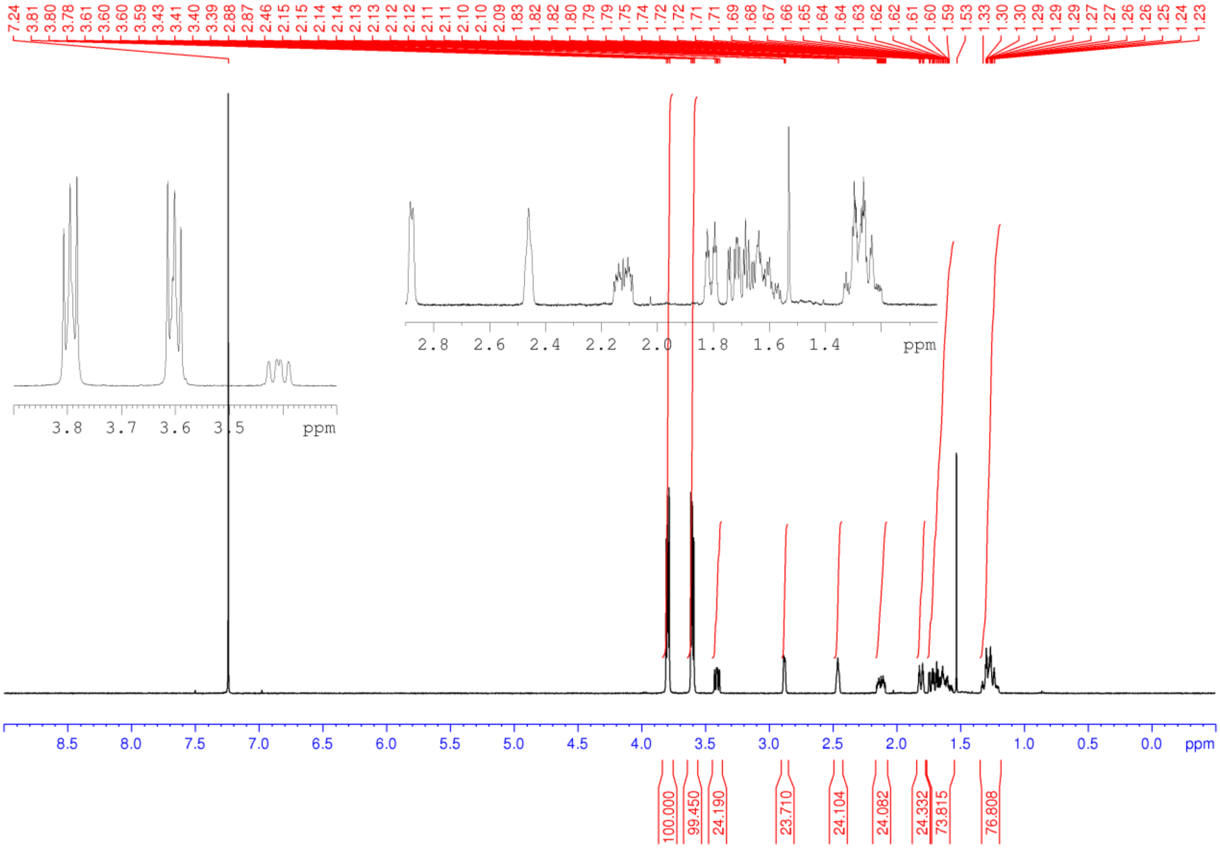

**Figure S29.** $^{1}H$ NMR spectrum (500 MHz, $CDCl_3$) of **14**

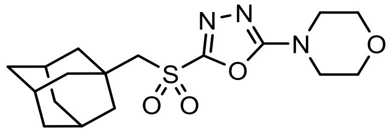

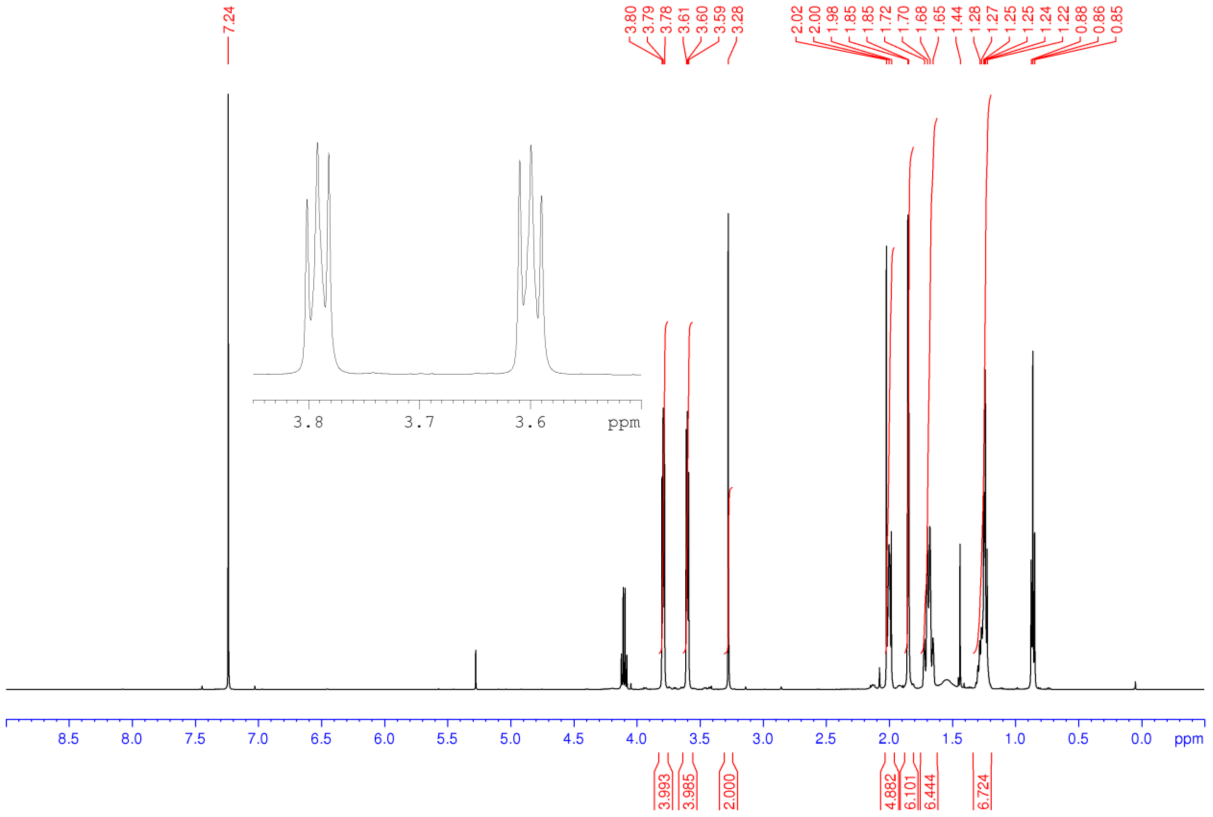

**Figure S30.** $^{1}H$ NMR spectrum (400 MHz, $CDCl_3$) of **15**

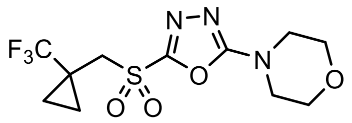

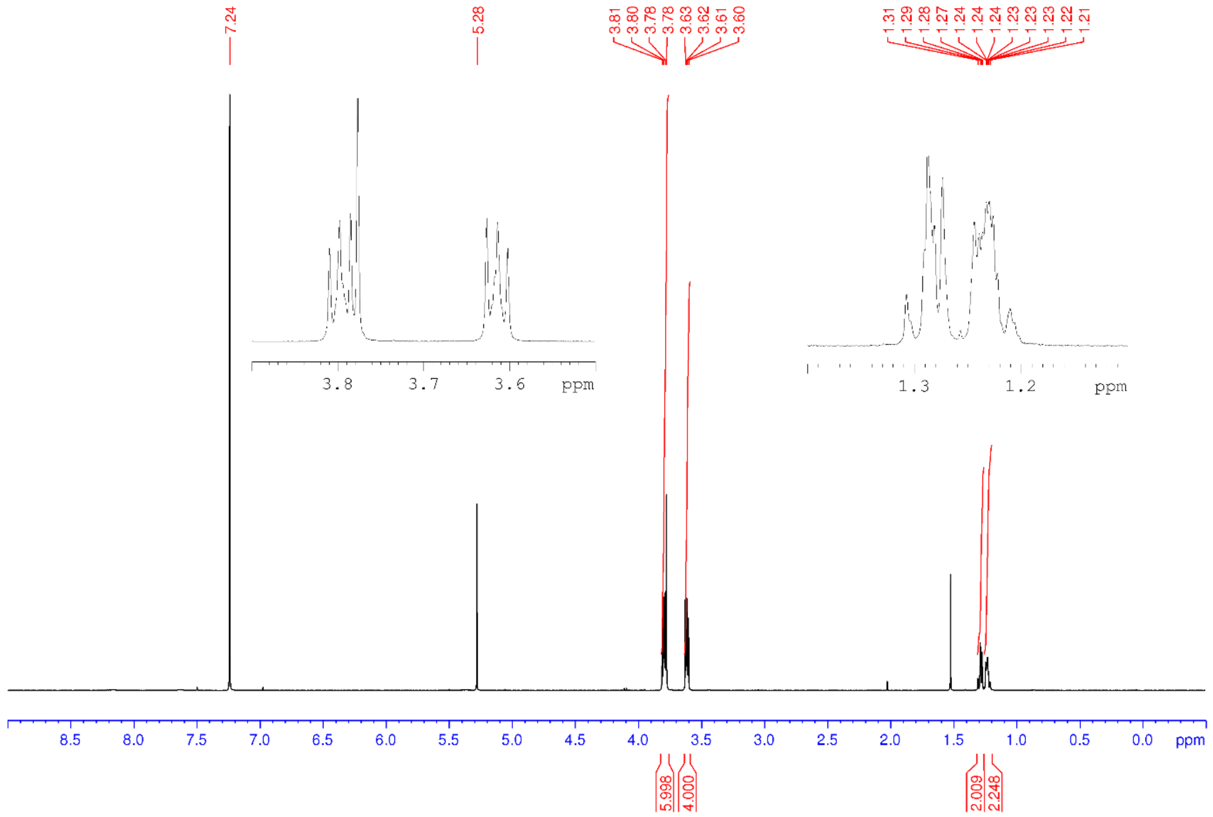

**Figure S31.** $^{1}H$ NMR spectrum (400 MHz, $CDCl_3$) of **16**

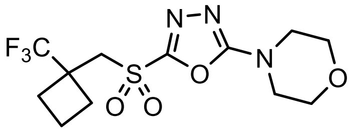


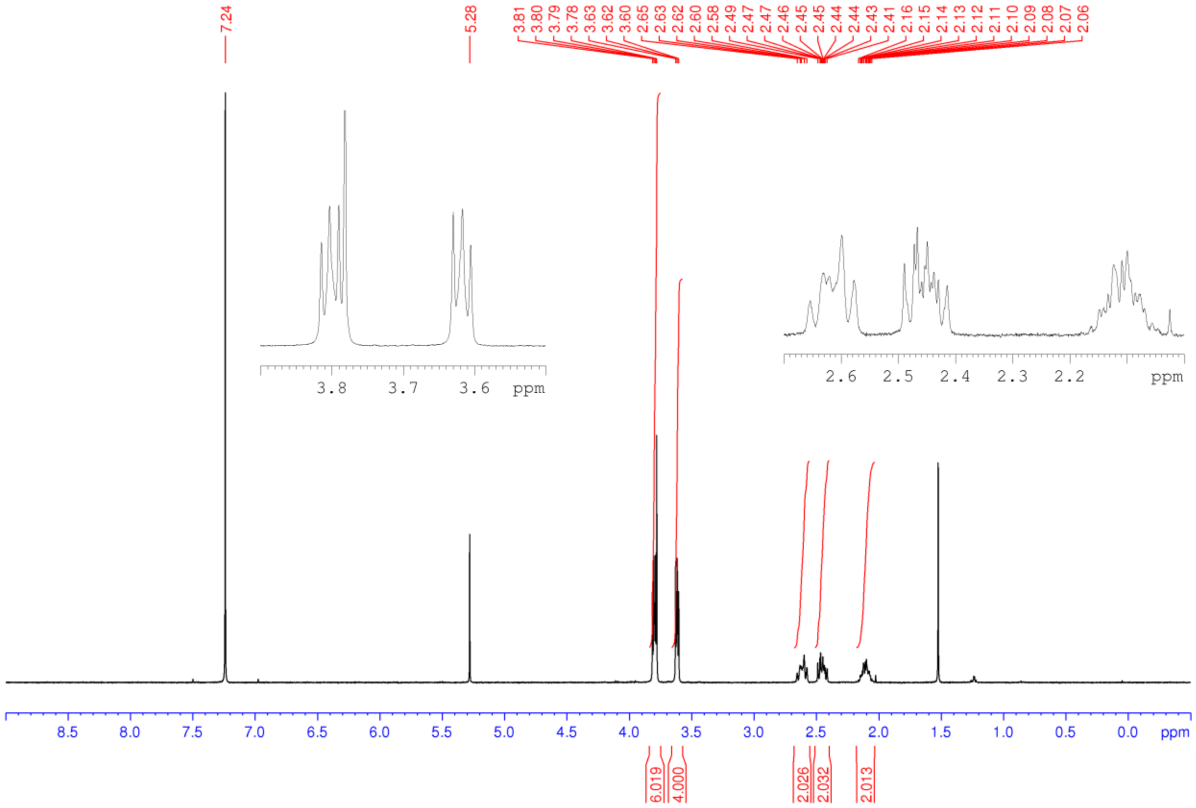

**Figure S32.** $^1$H NMR spectrum (400 MHz, $CDCl_3$) of **17**

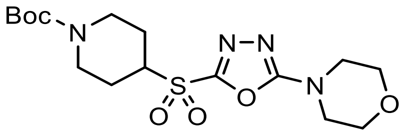


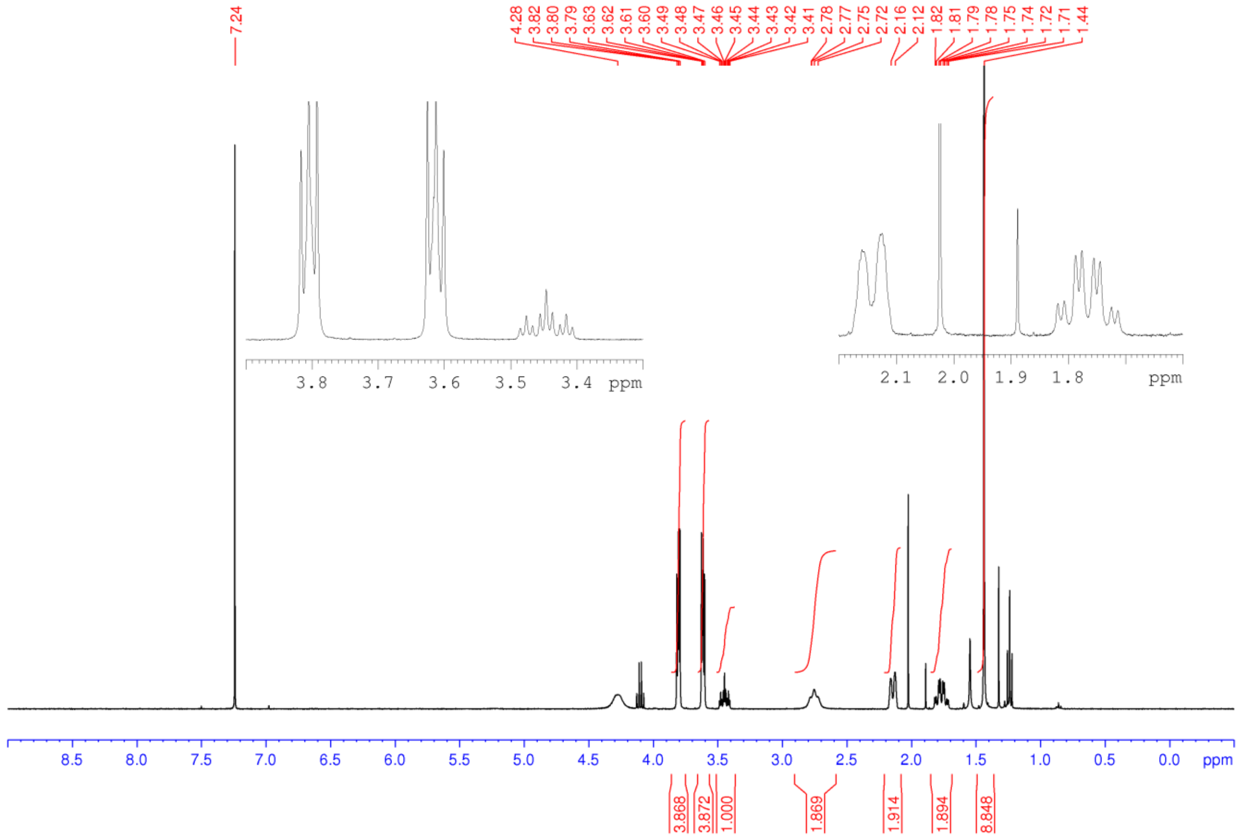

**Figure S33.** $^1$H NMR spectrum (400 MHz, $CD_3OD$) of **18**

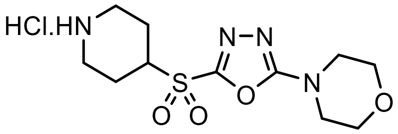


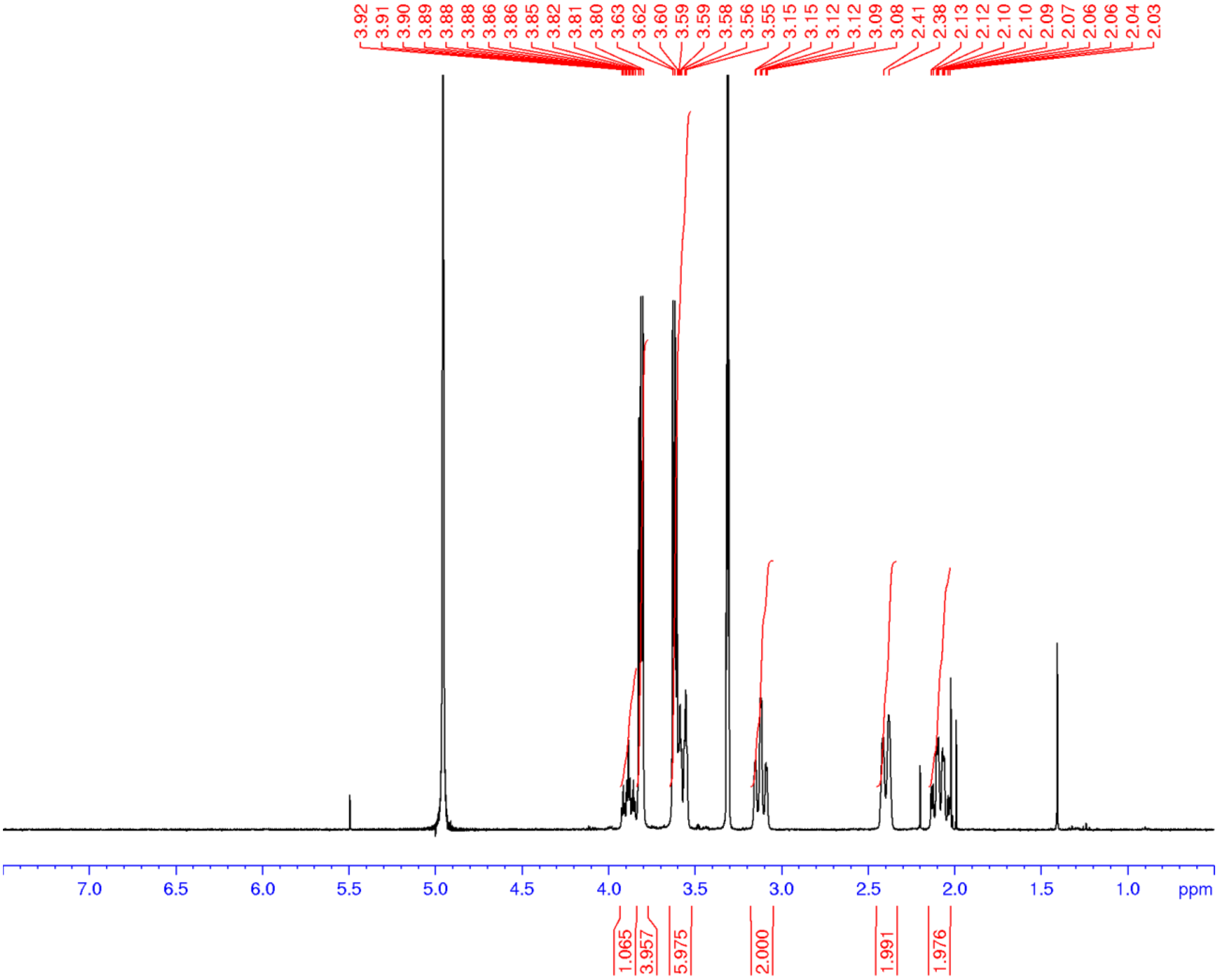

**Figure S34.** $^1$H NMR spectrum (400 MHz, $CDCl_3$) of **19**

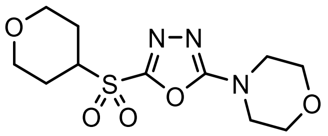

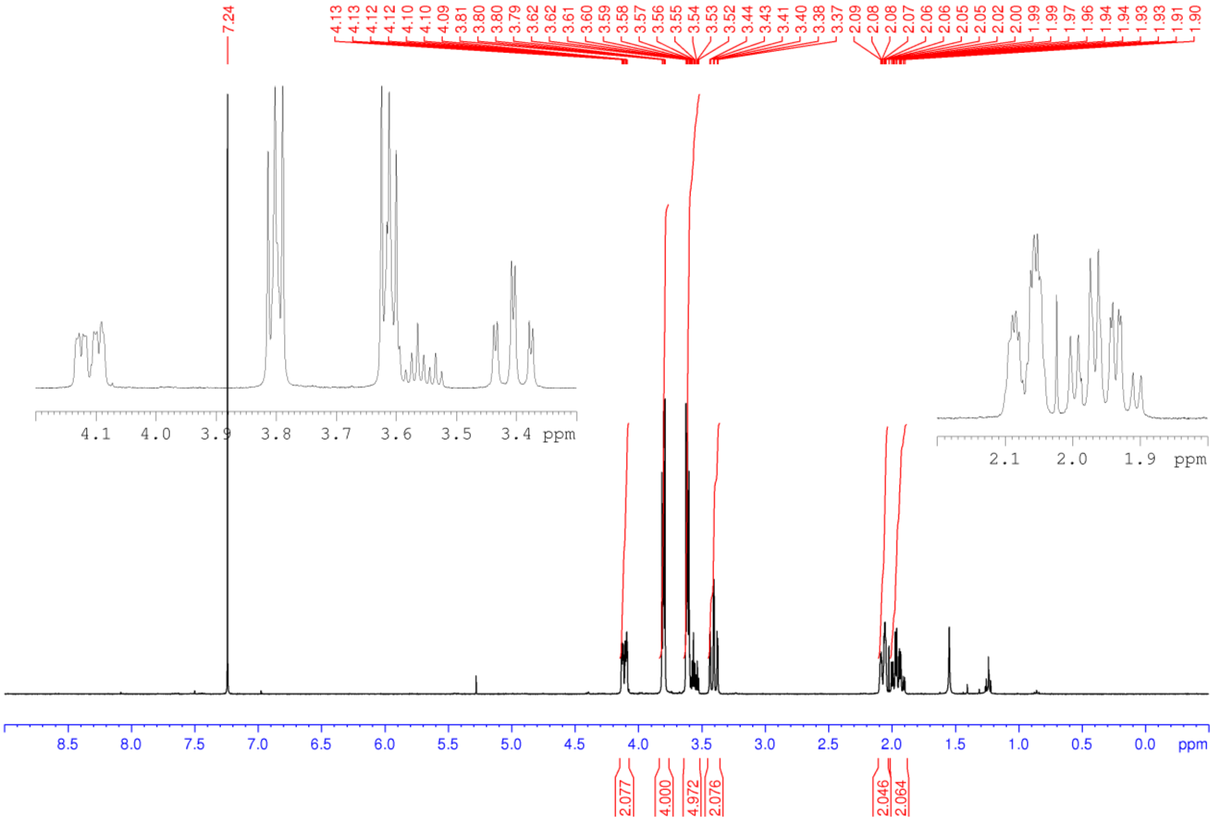

**Figure S35.** $^{1}H$ NMR spectrum (400 MHz, $CDCl_3$) of **20**

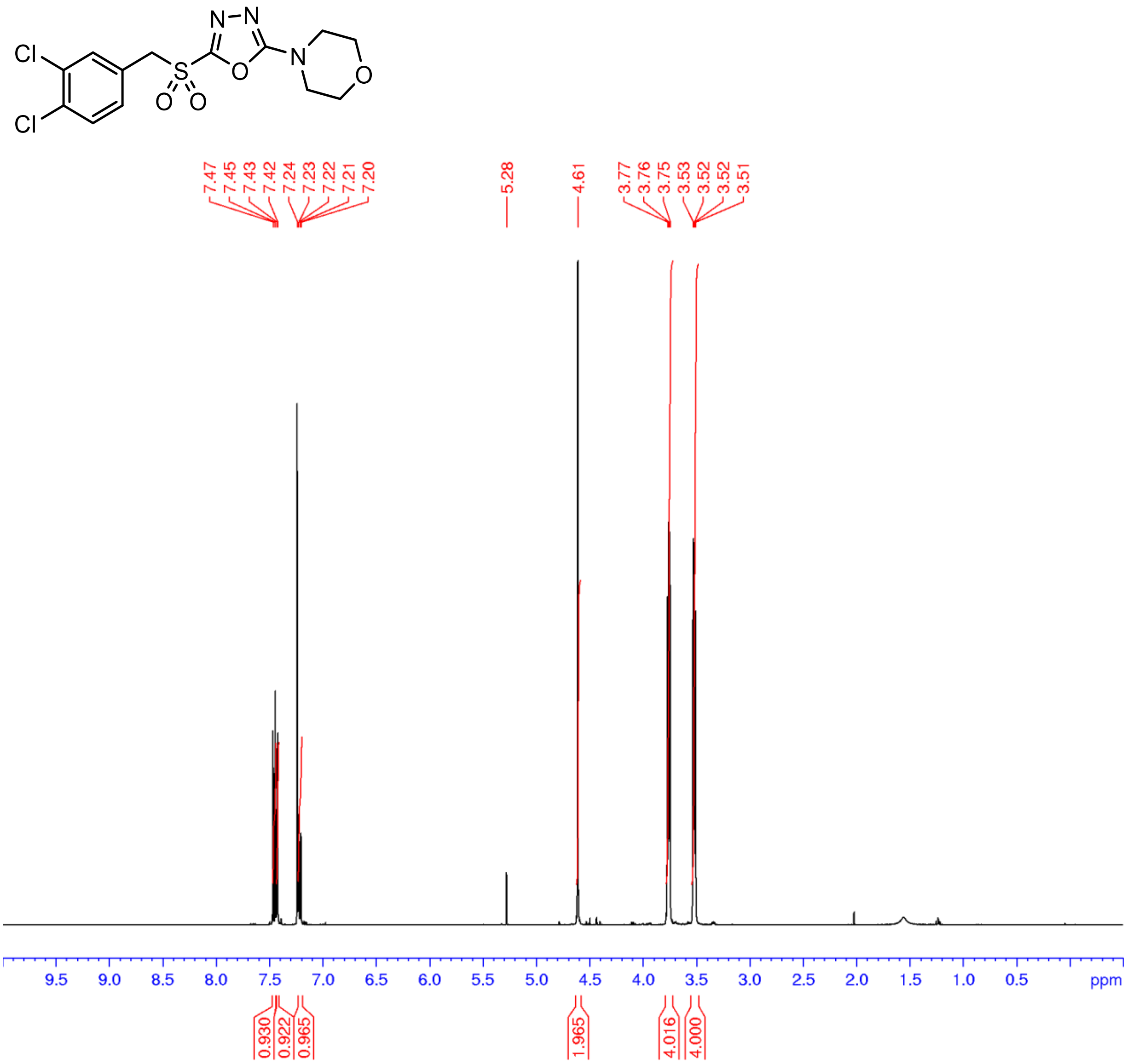

**Figure S36.** $^{1}$H NMR spectrum (400 MHz, $CDCl_3$) of **21**

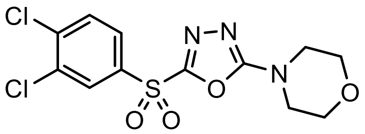

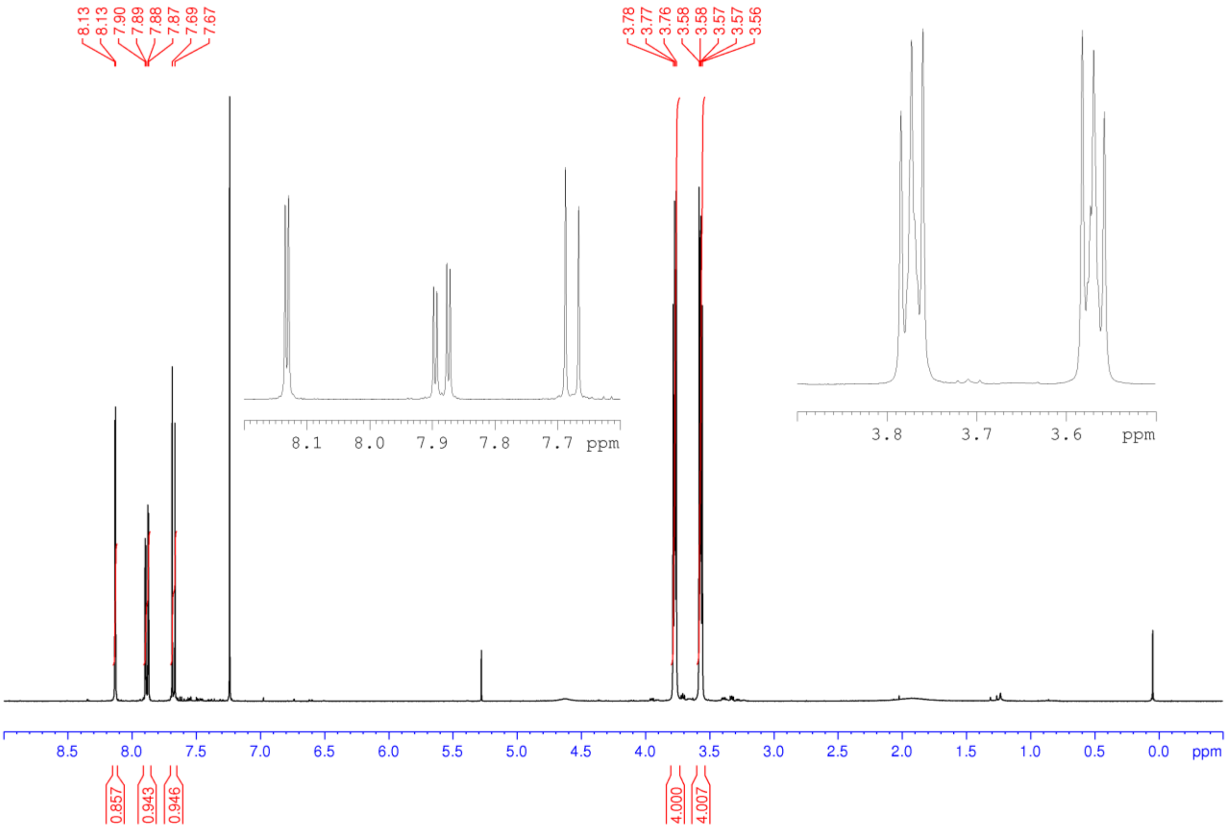

**Figure S37.** $^{13}C$ NMR spectrum (100 MHz, $CDCl_3$) of **21**

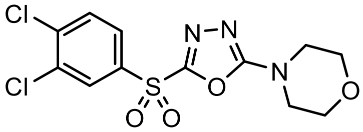


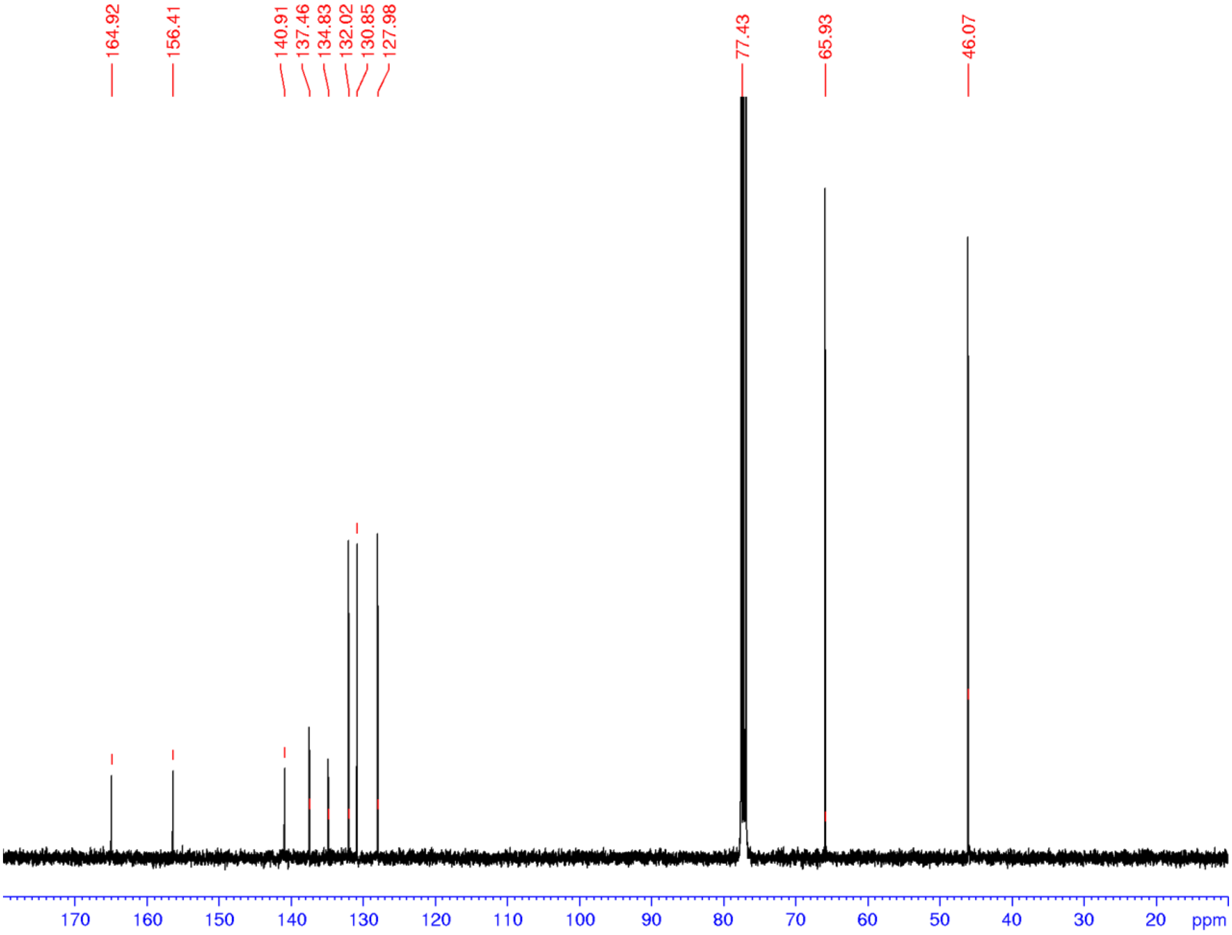

**Figure S38.** $^{1}$H NMR spectrum (400 MHz, $CDCl_3$) of **22**

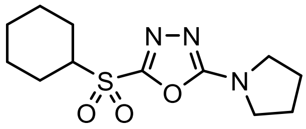

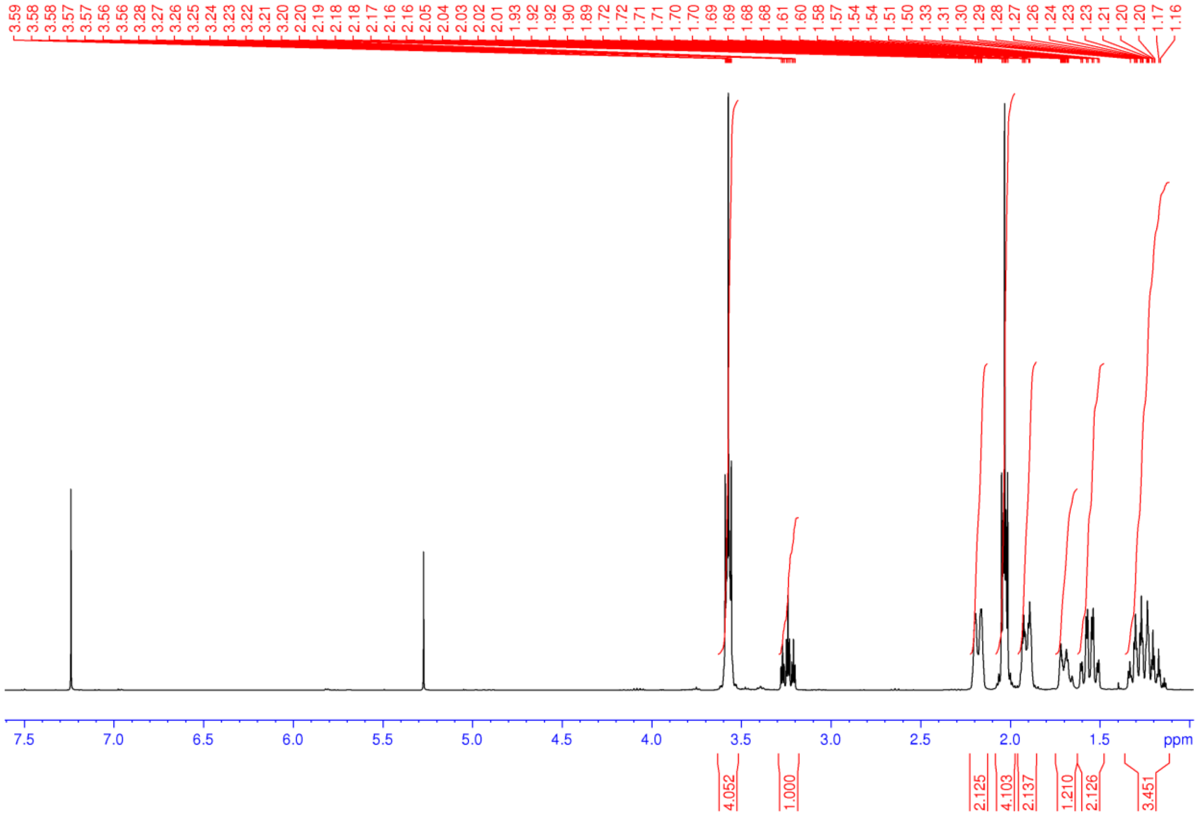

**Figure S39.** $^1$H NMR spectrum (500 MHz, $CDCl_3$) **23**

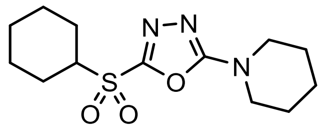

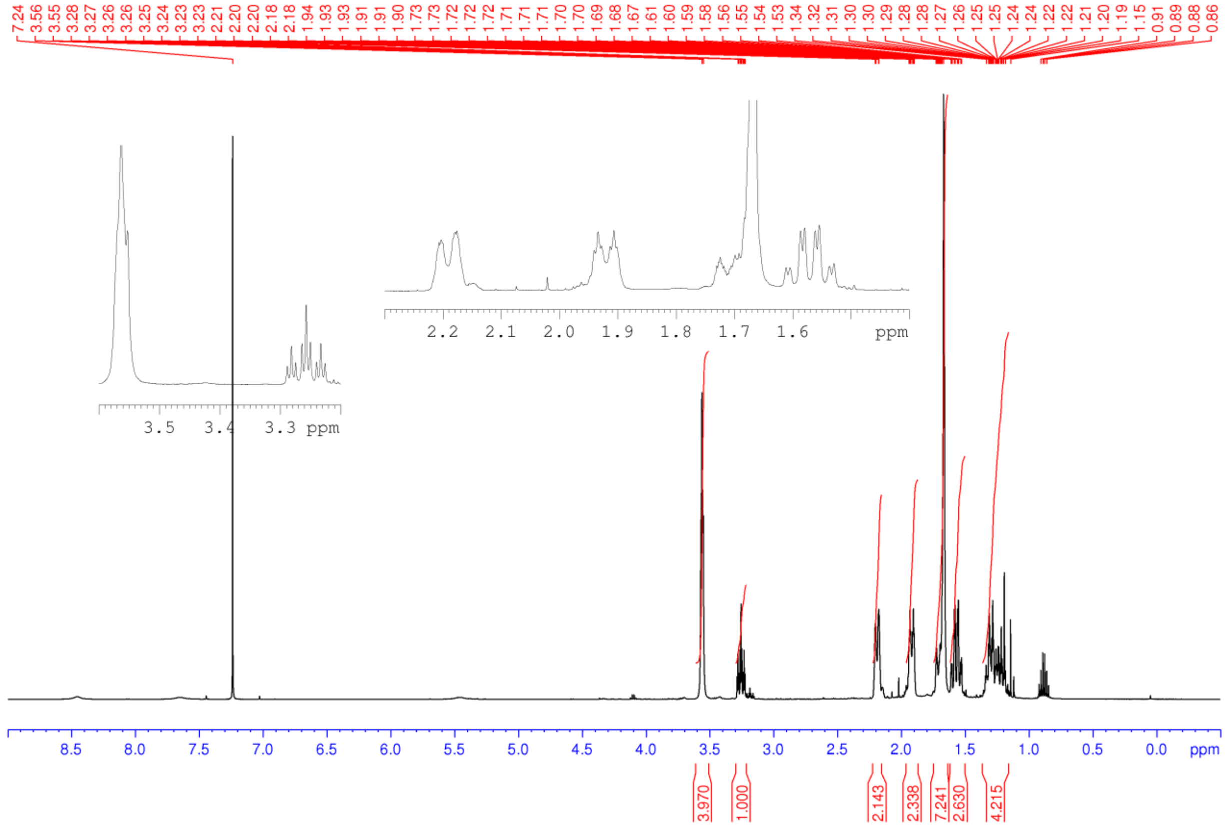

**Figure S40.** $^1$H NMR spectrum (400 MHz, $CDCl_3$) of **24**

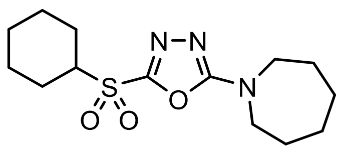


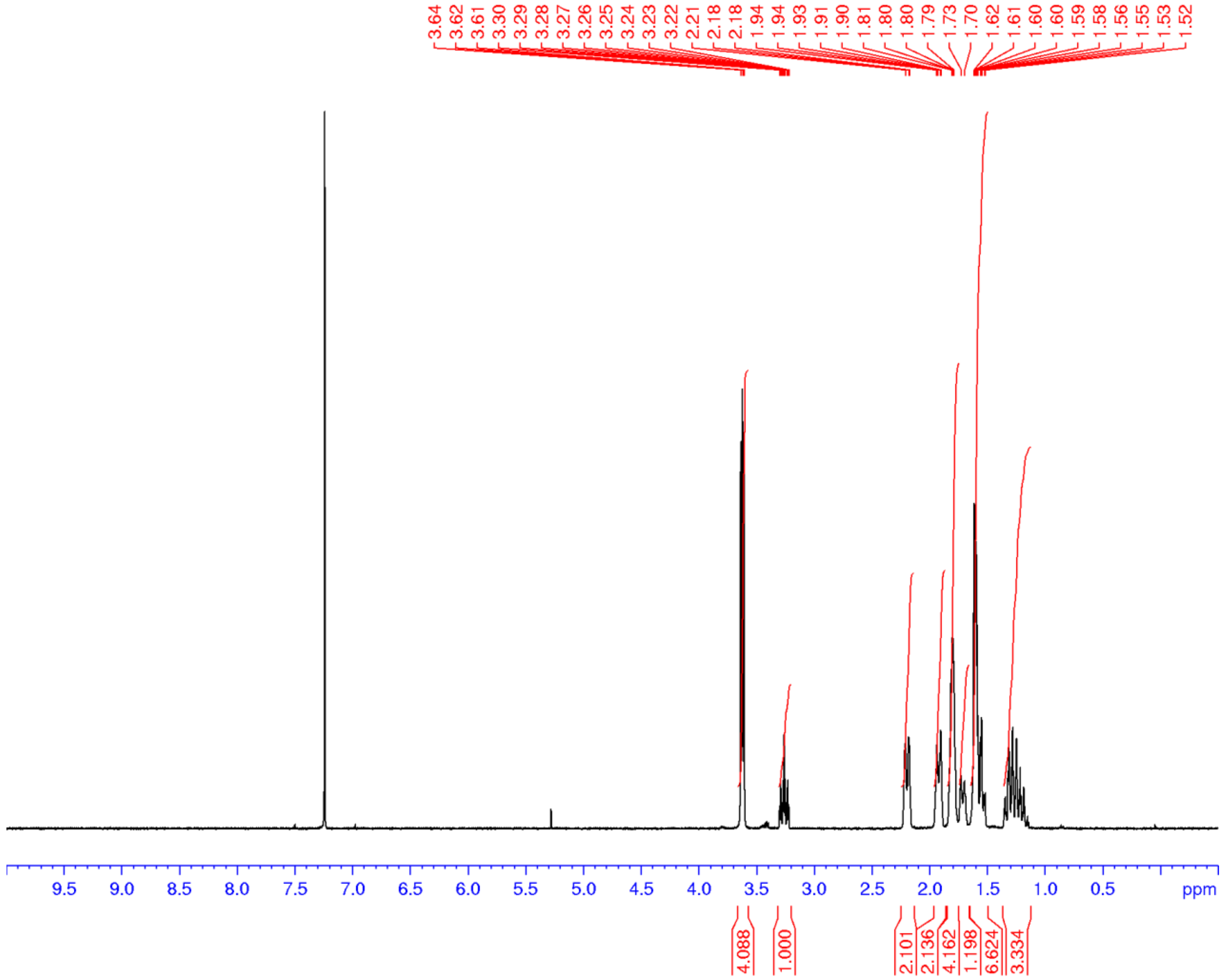

**Figure S41.** $^{13}C$ NMR spectrum (100 MHz, $CDCl_3$) of **24**

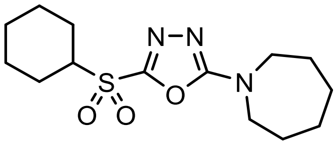

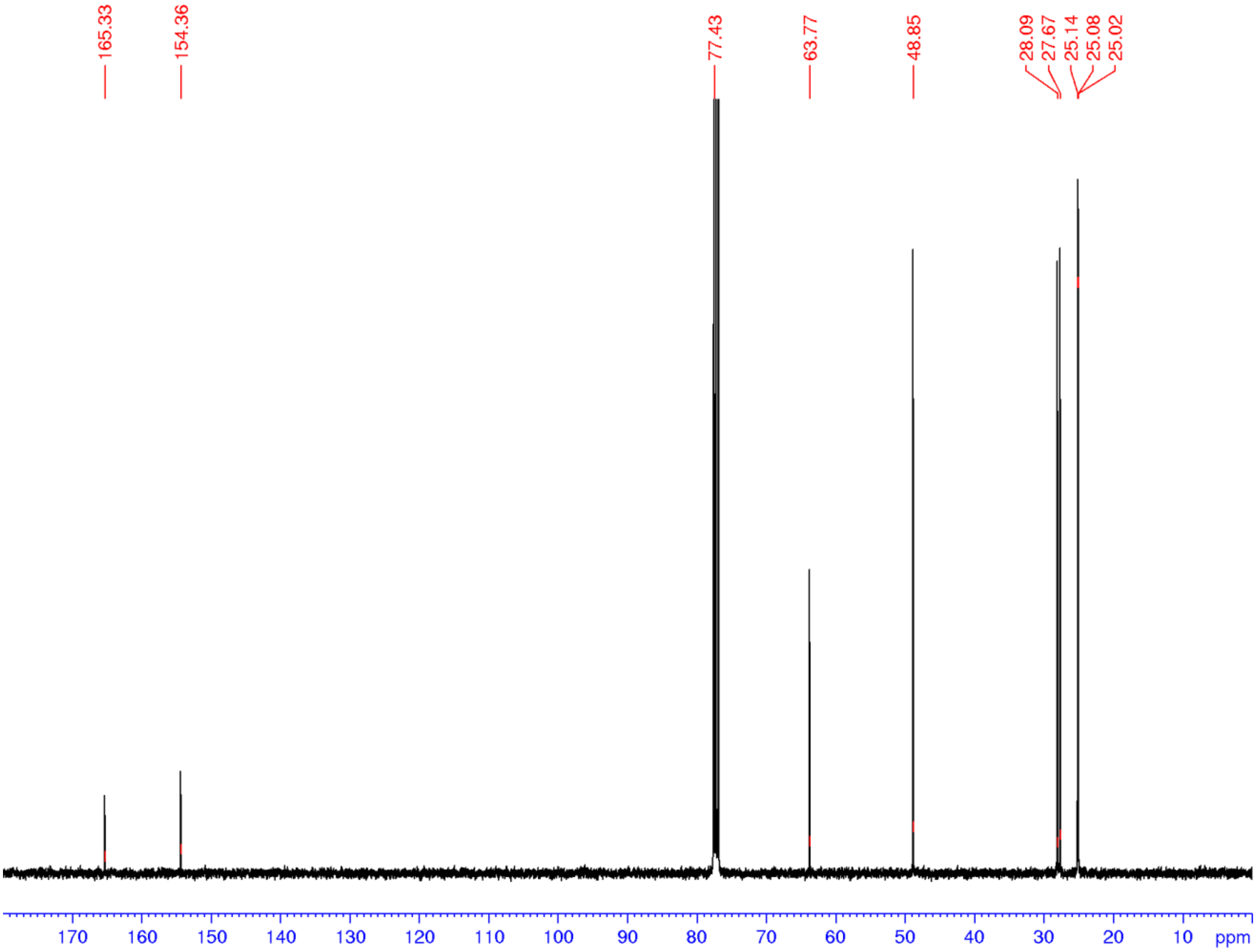

**Figure S42.** $^{1}$H NMR spectrum (400 MHz, $CDCl_3$) for **25**

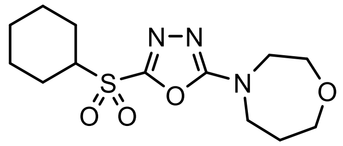

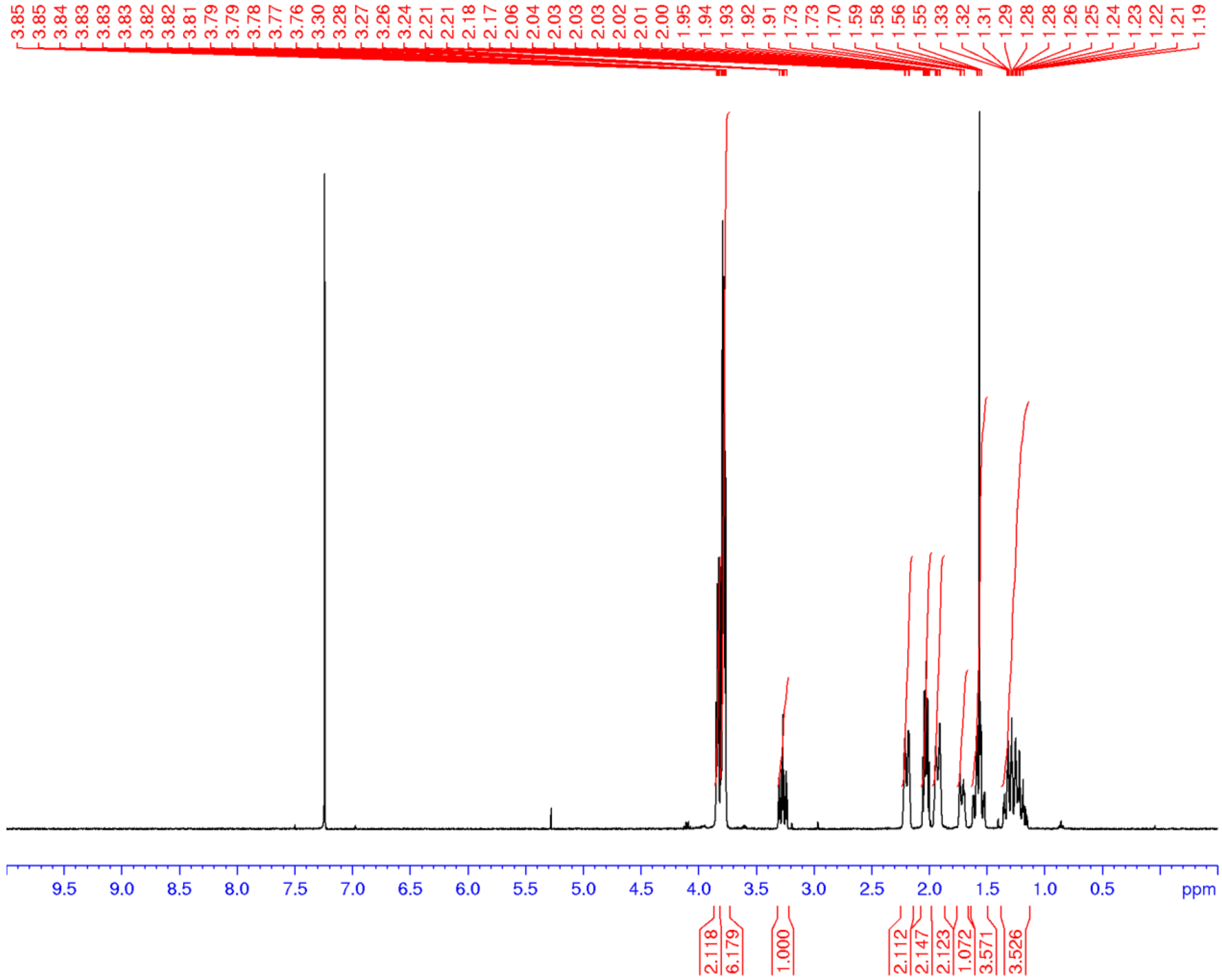

**Figure S43.** $^{1}$H NMR spectrum (400 MHz, $CDCl_3$) of **26**

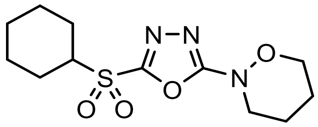

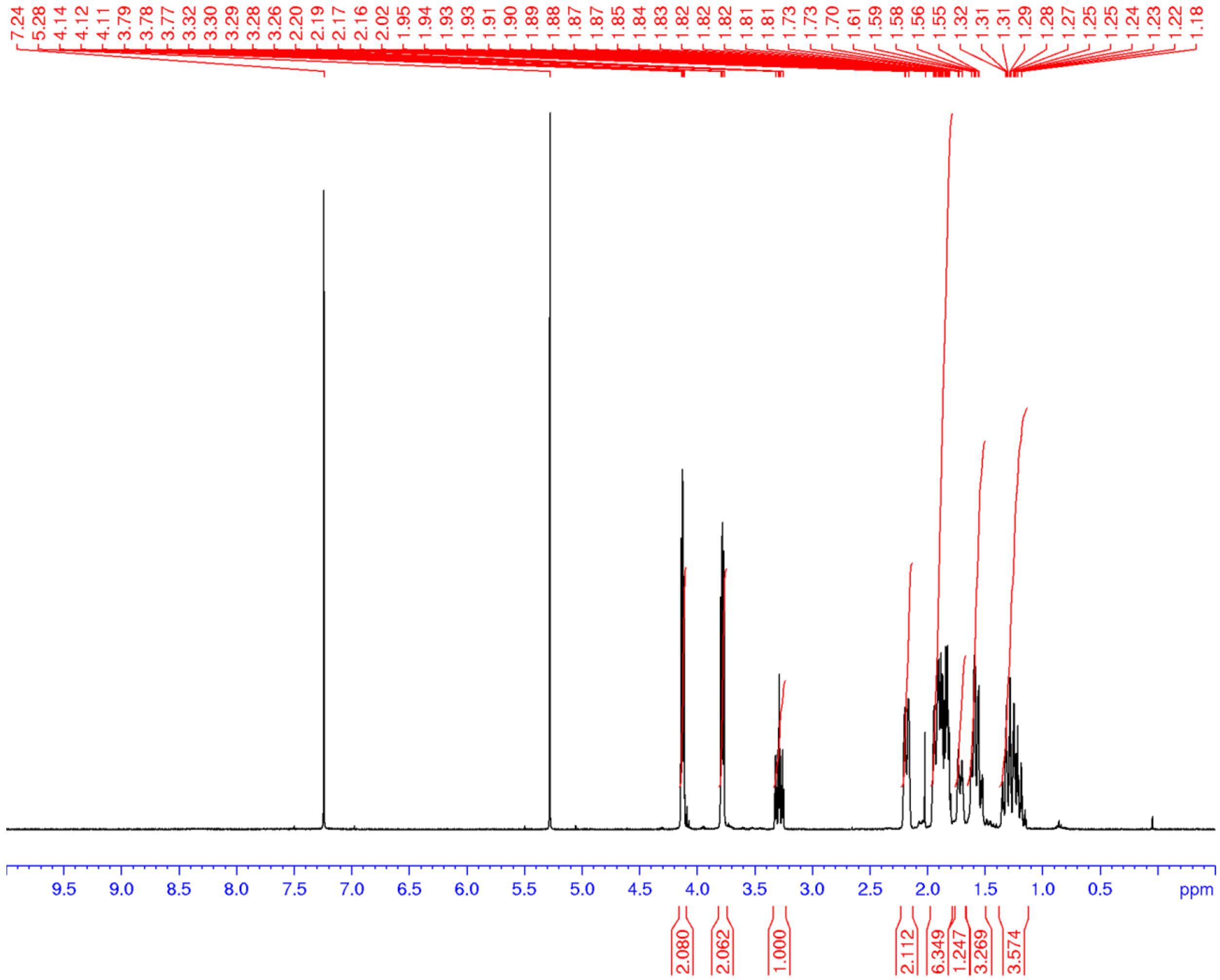

**Figure S44.** $^{1}H$ NMR spectrum (400 MHz, $CDCl_3$) of **27**

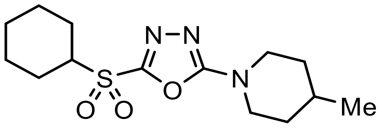


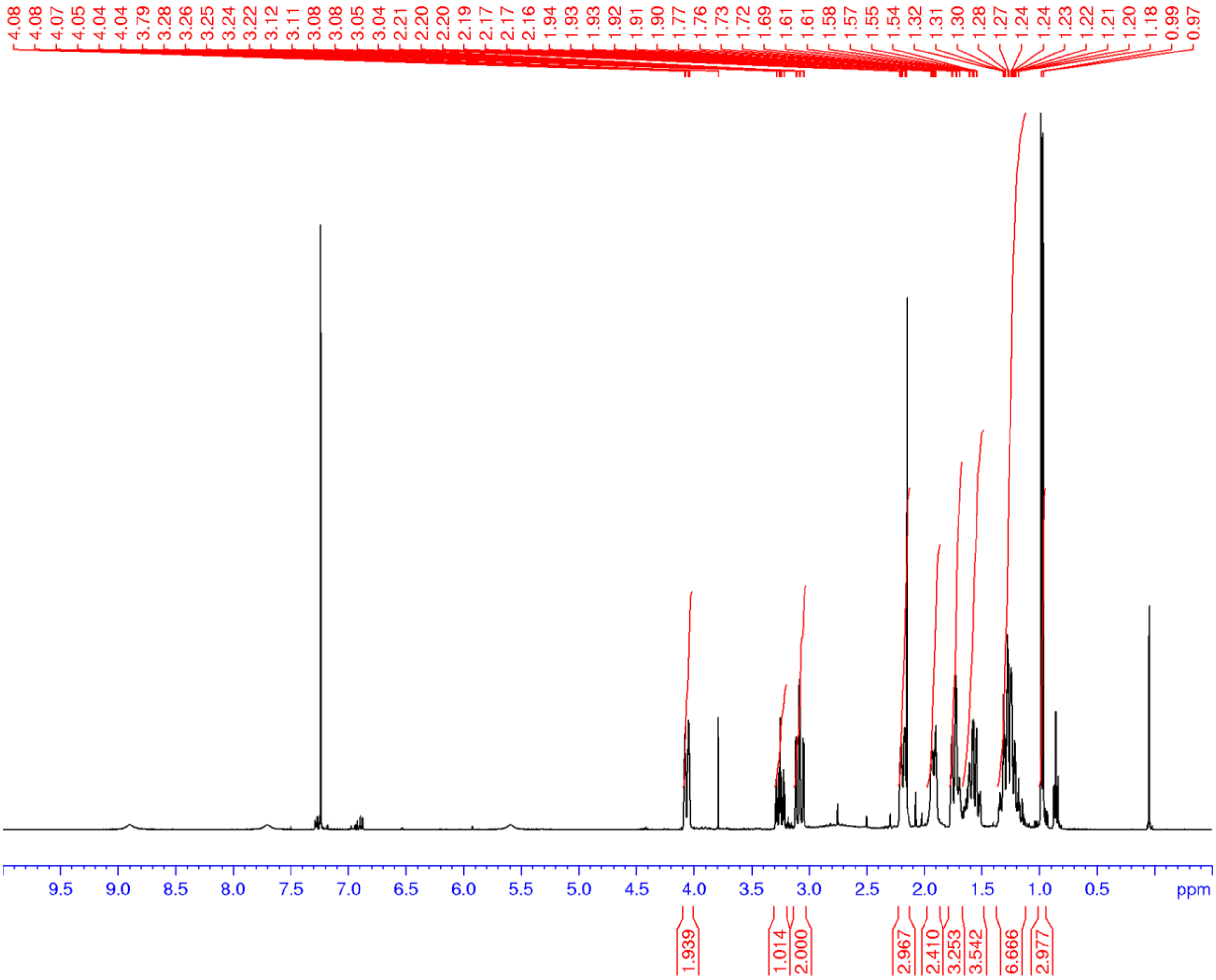

**Figure S45.** $^{1}H$ NMR spectrum ($CDCl_3$, 400 MHz) of **28**

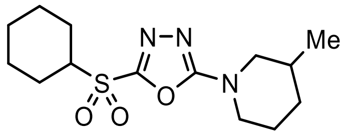


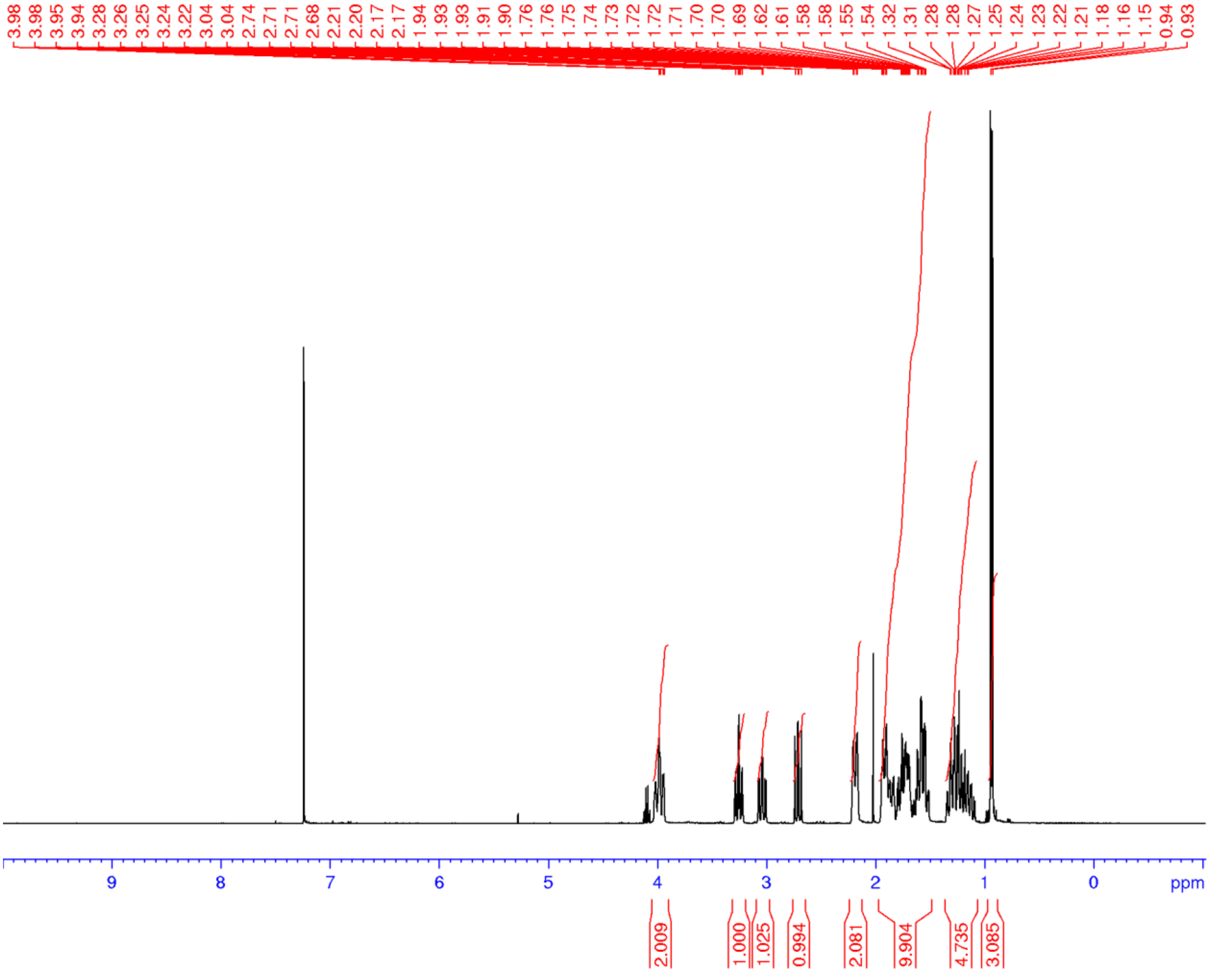

**Figure S46.** $^{1}H$ NMR spectrum (500 MHz, $CDCl_3$) of **29**

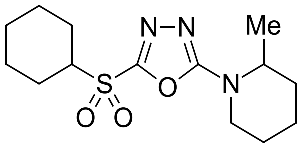


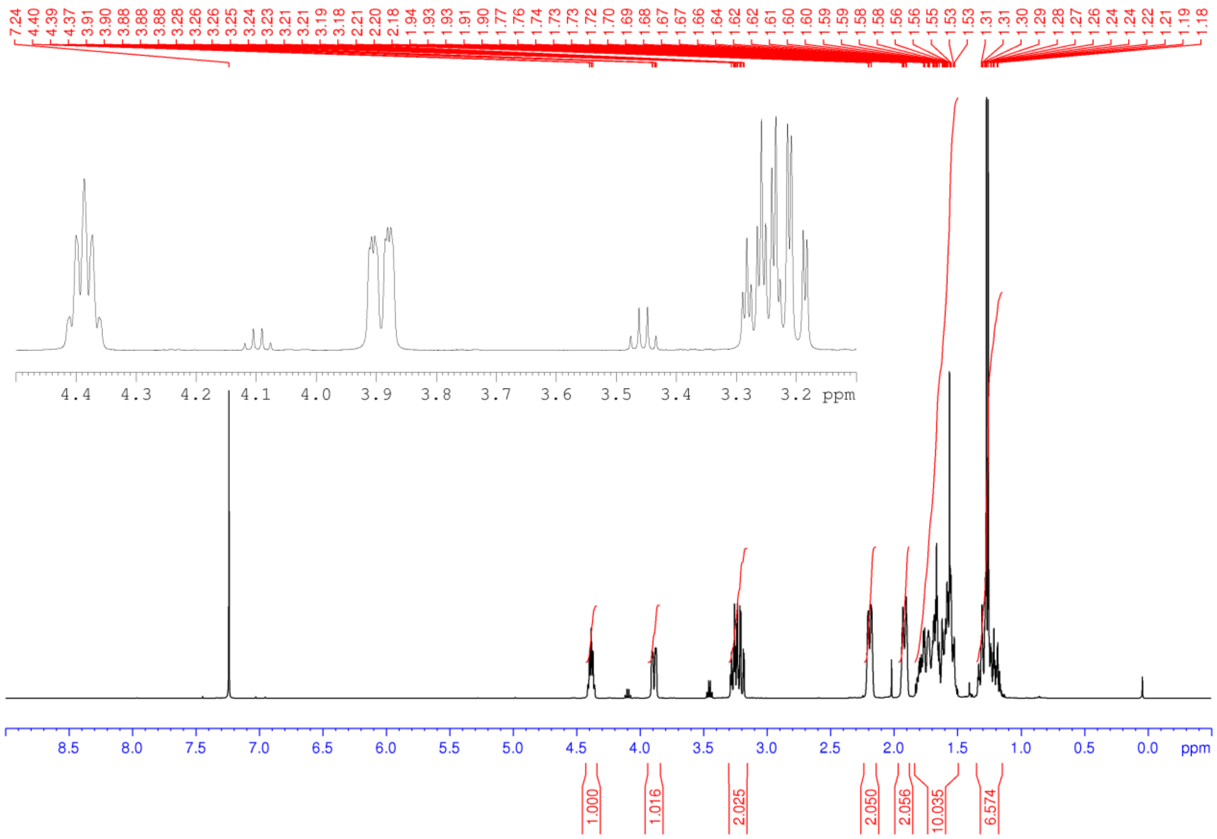

**Figure S47.** $^{1}$H NMR spectrum (400 MHz, $CDCl_3$) of **30**

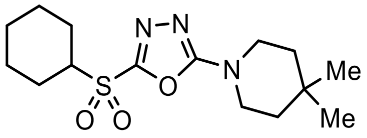

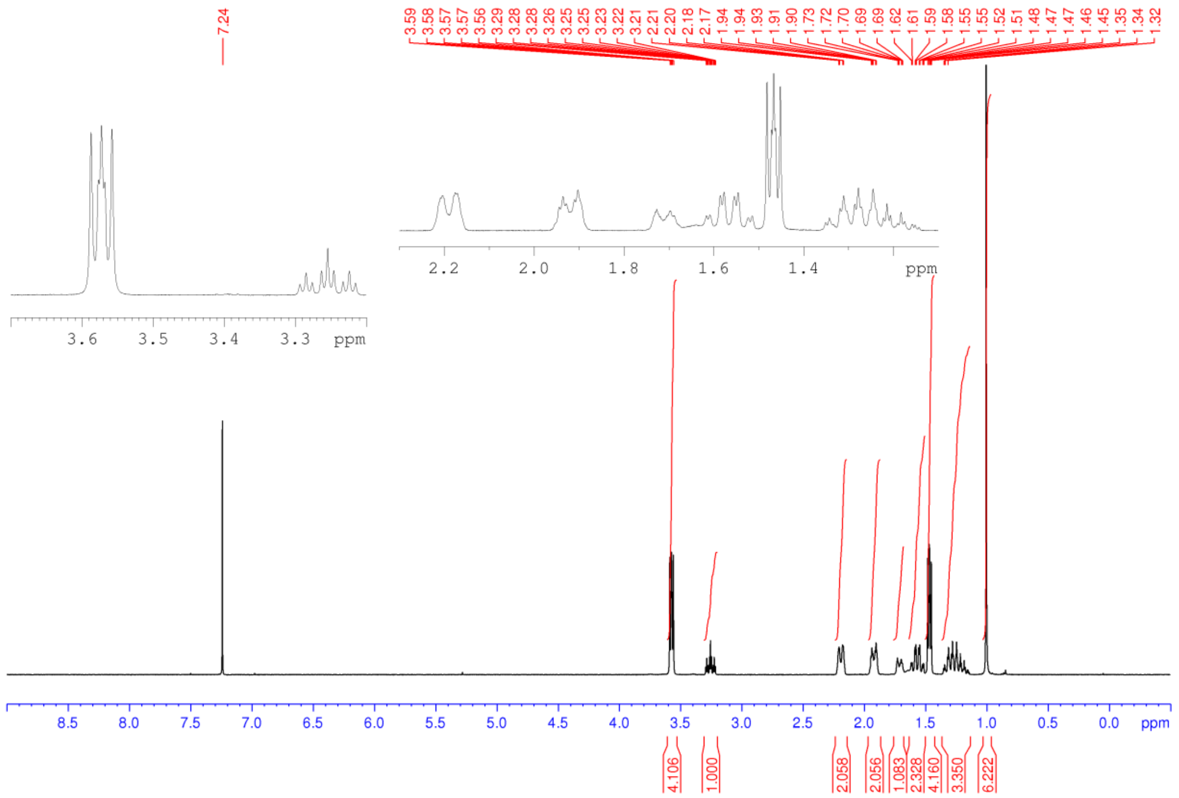

**Figure S48.** $^{1}H$ NMR spectrum (500 MHz, $CDCl_3$) of **31**

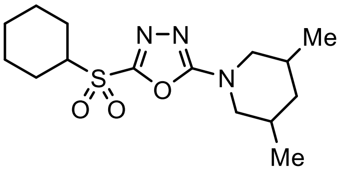


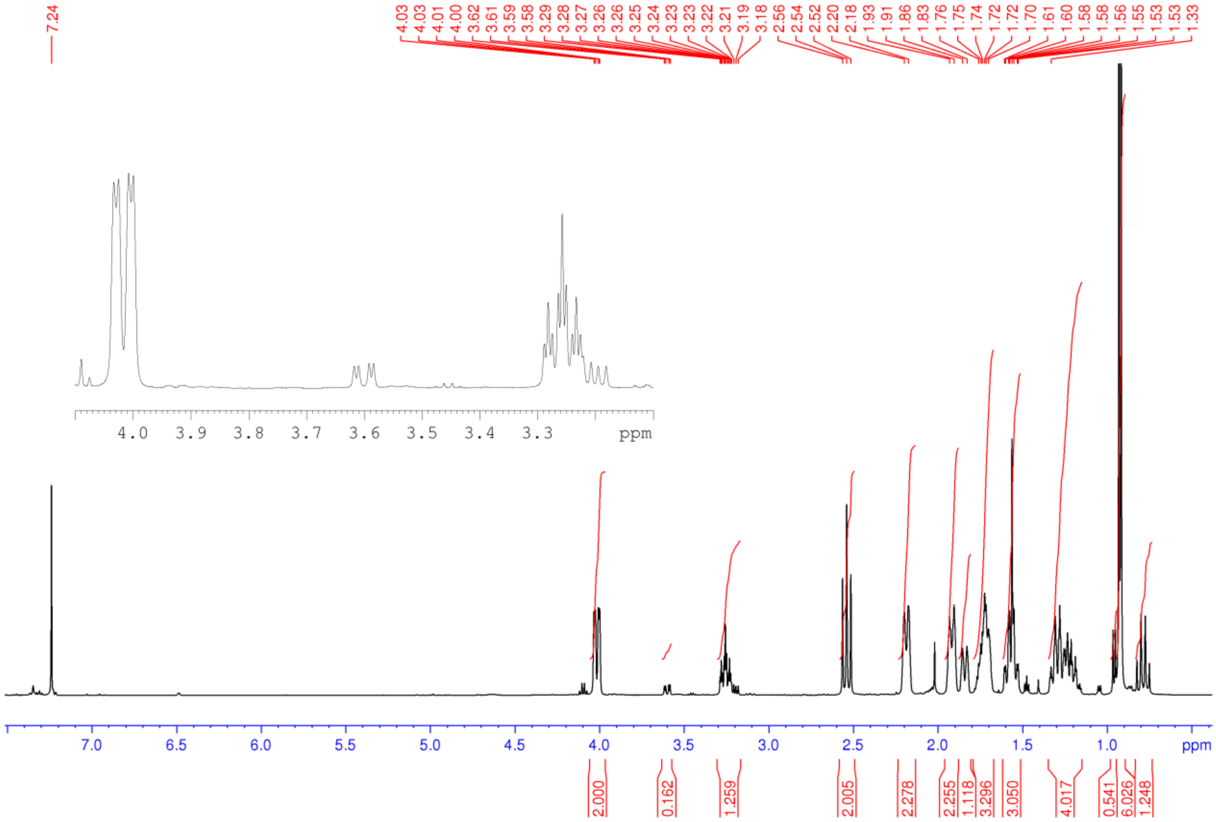

**Figure S49.** $^{1}$H NMR spectrum (400 MHz, $CDCl_3$) of **32**

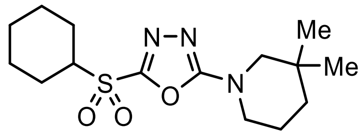

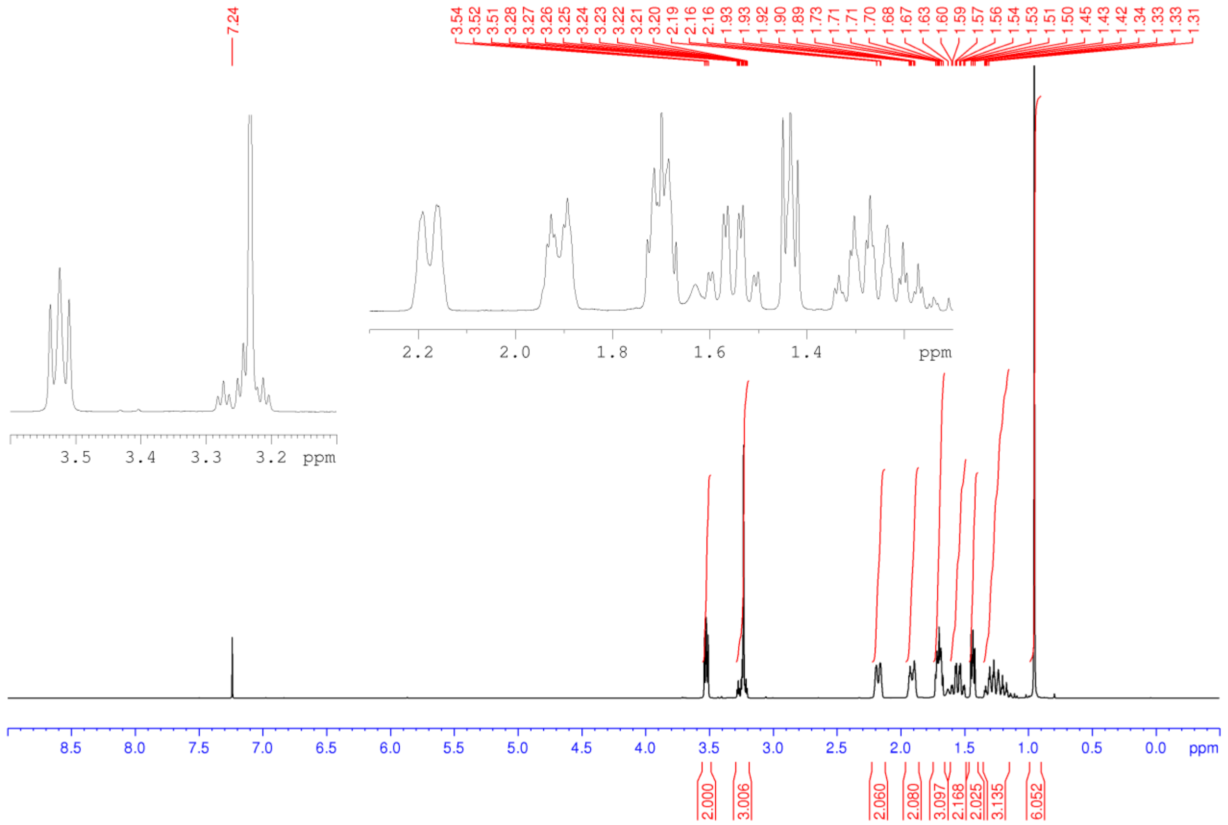

**Figure S50.** $^{1}$H NMR spectrum (400 MHz, $CDCl_3$) of **33**

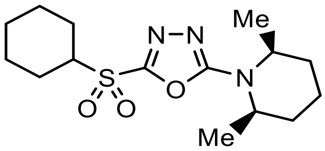


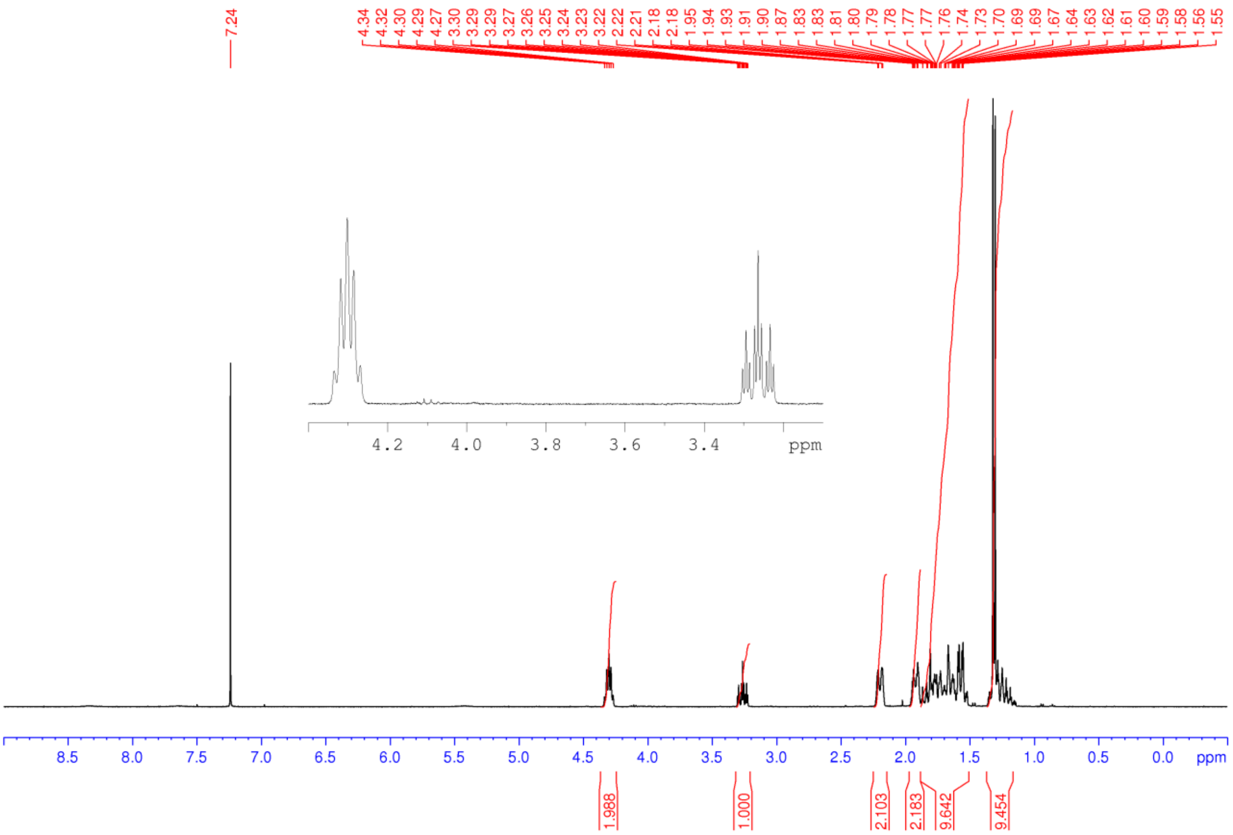

**Figure S51.** $^{13}$C NMR spectrum (100 MHz, $CDCl_3$) of **33**

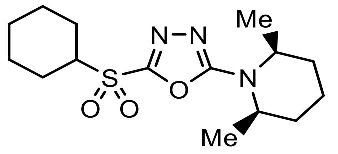


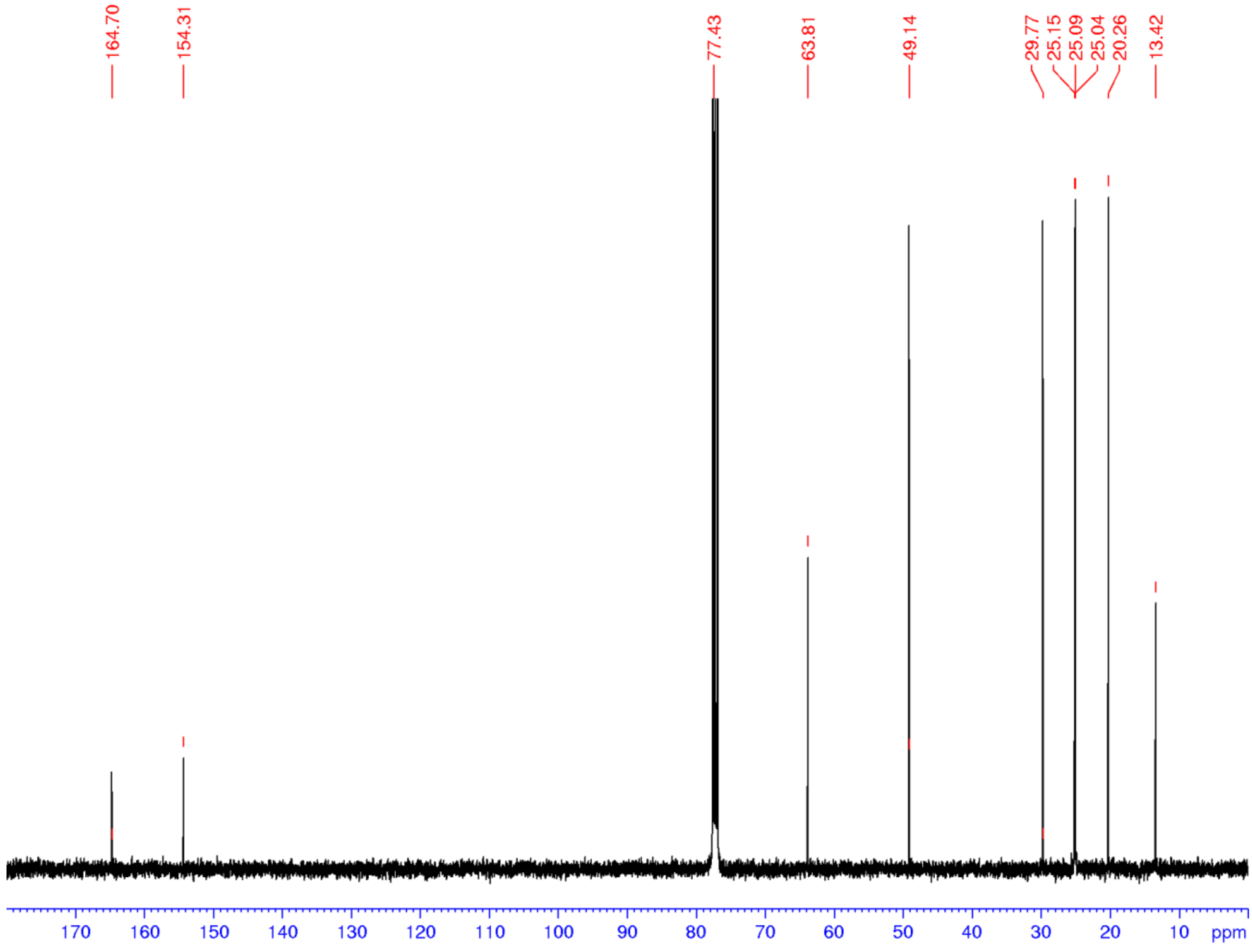

**Figure S52.** $^{1}$H NMR spectrum (400 MHz, $CDCl_3$) of **34**

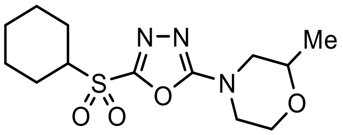


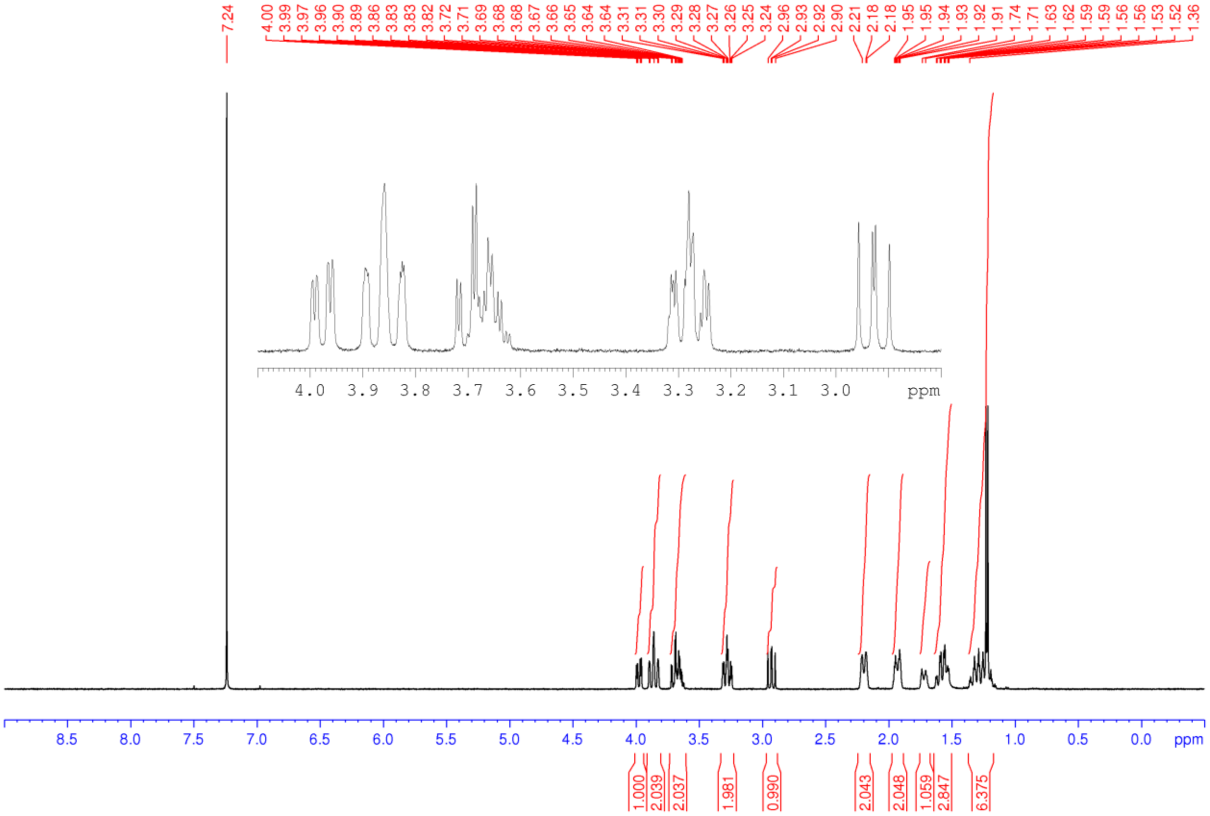

**Figure S53.** $^{1}H$ NMR spectrum (400 MHz, $CDCl_3$) of **35**

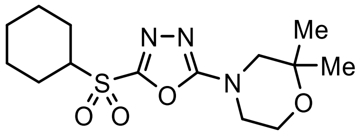


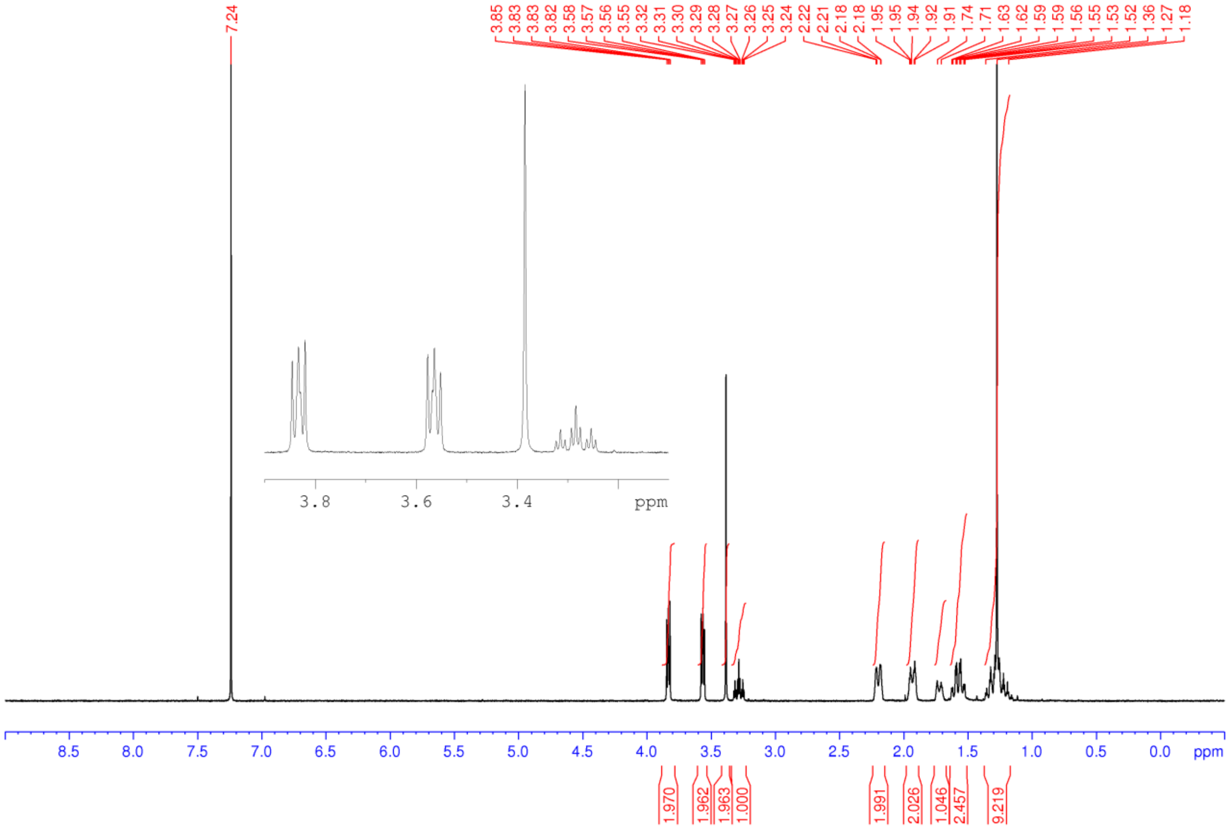

**Figure S54.** $^{1}$H NMR spectrum (400 MHz, $CDCl_3$) of **36**

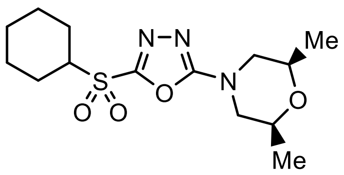


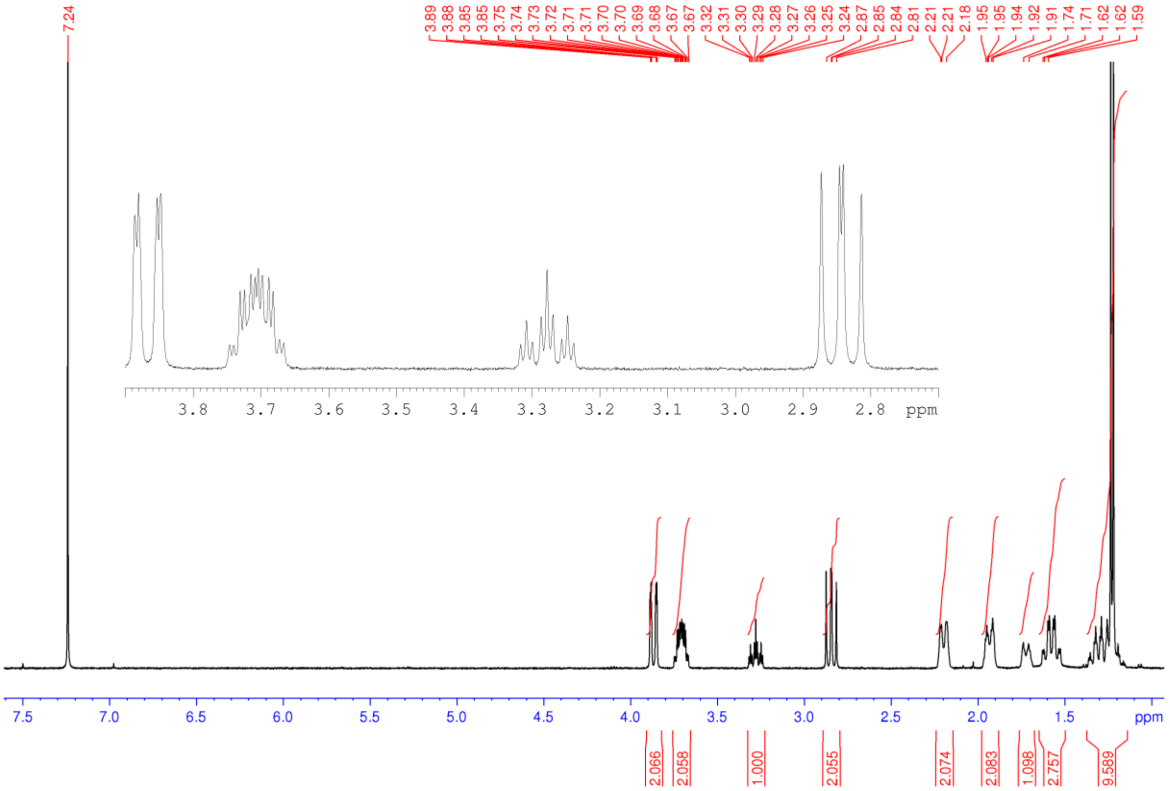

**Figure S55.** $^1$H NMR spectrum (400 MHz, $CDCl_3$) of **37**

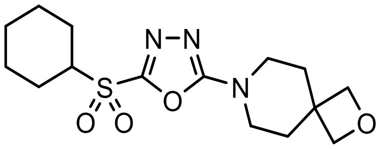

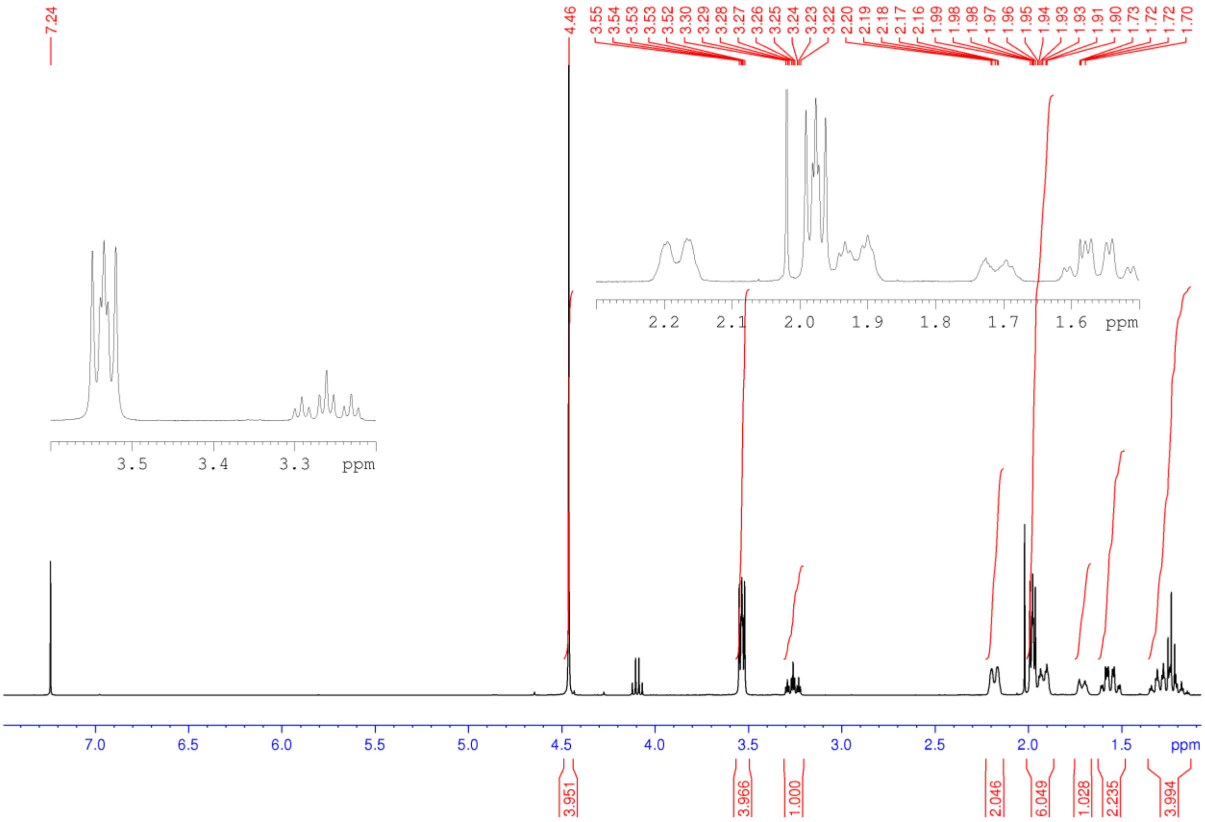

**Figure S56.** $^{1}H$ NMR spectrum (400 MHz, $CDCl_3$) of **38**

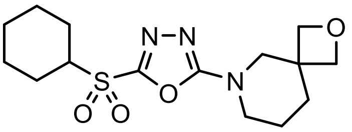


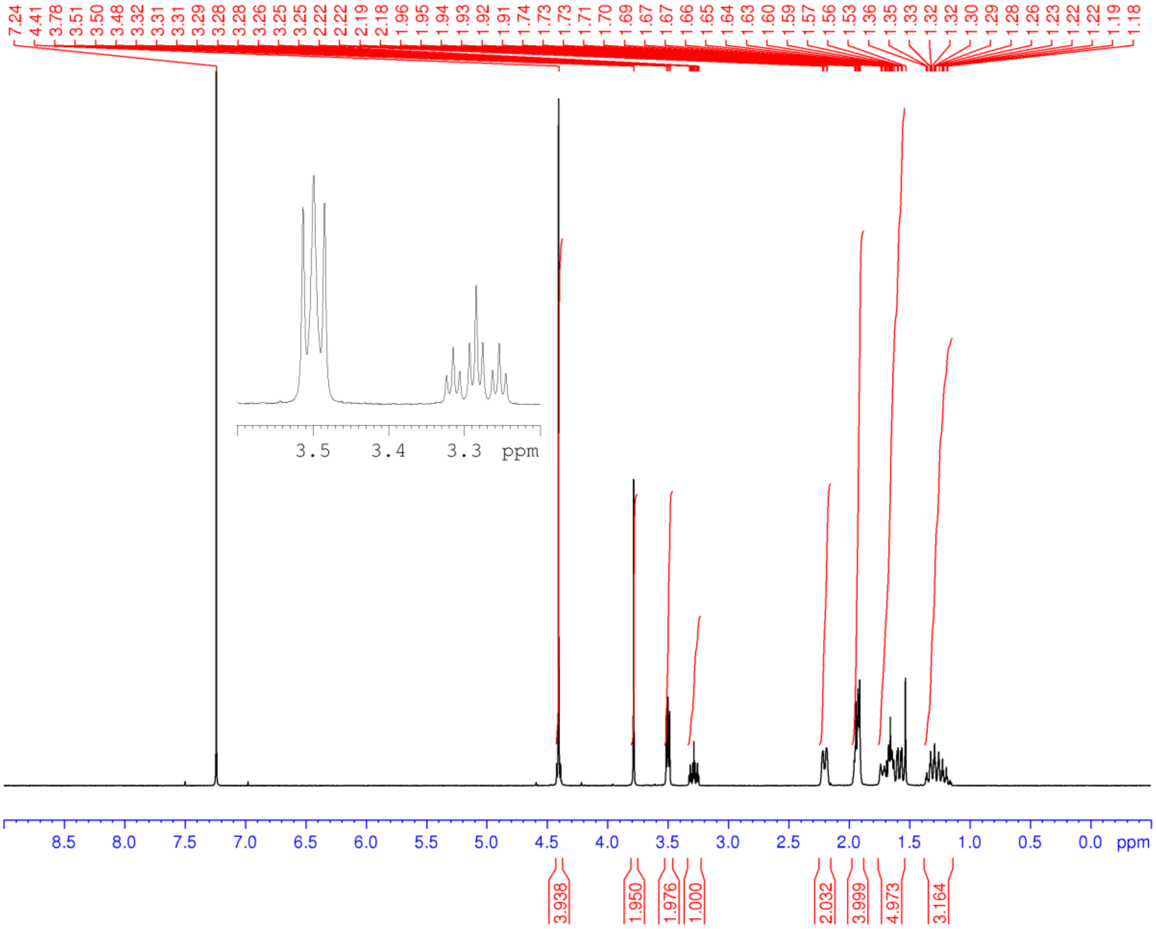

**Figure S57.** $^1$H NMR spectrum (400 MHz, $CDCl_3$) of **39**

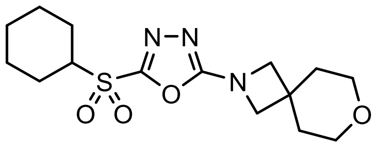


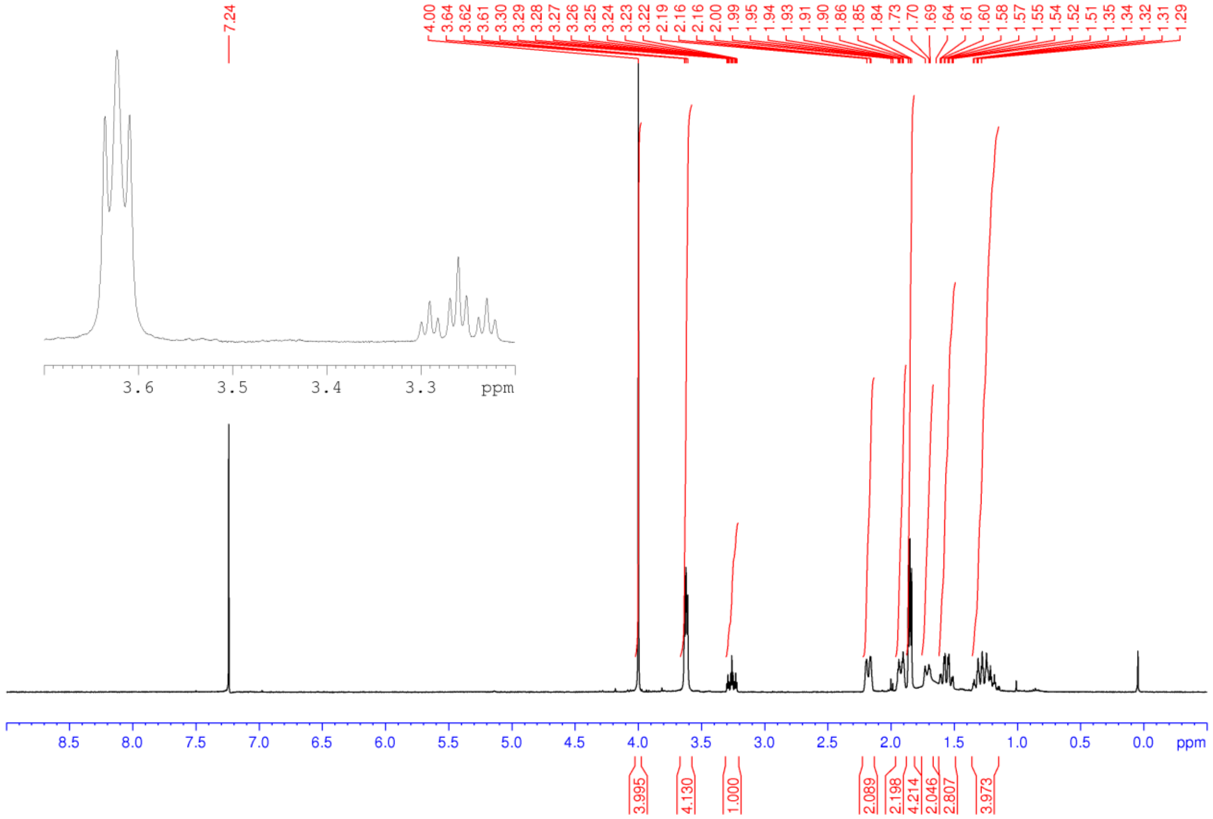

**Figure S58.** $^1$H NMR spectrum (500 MHz, $CDCl_3$) of **40**

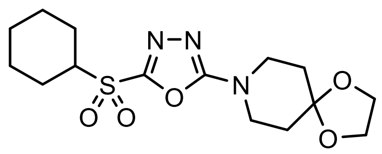

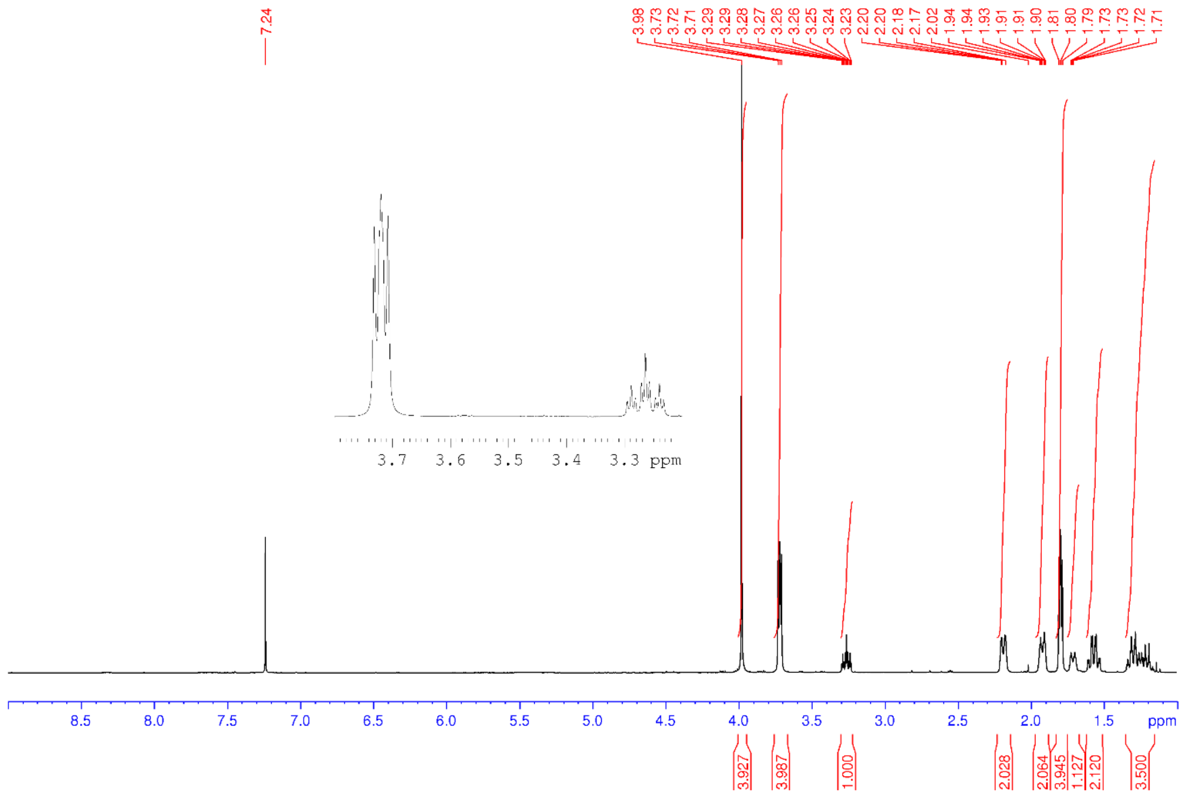

**Figure S59.** $^{1}$H NMR spectrum (400 MHz, $CDCl_3$) of **41**

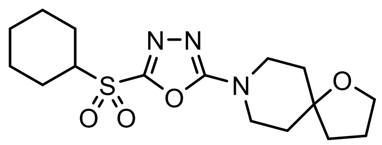

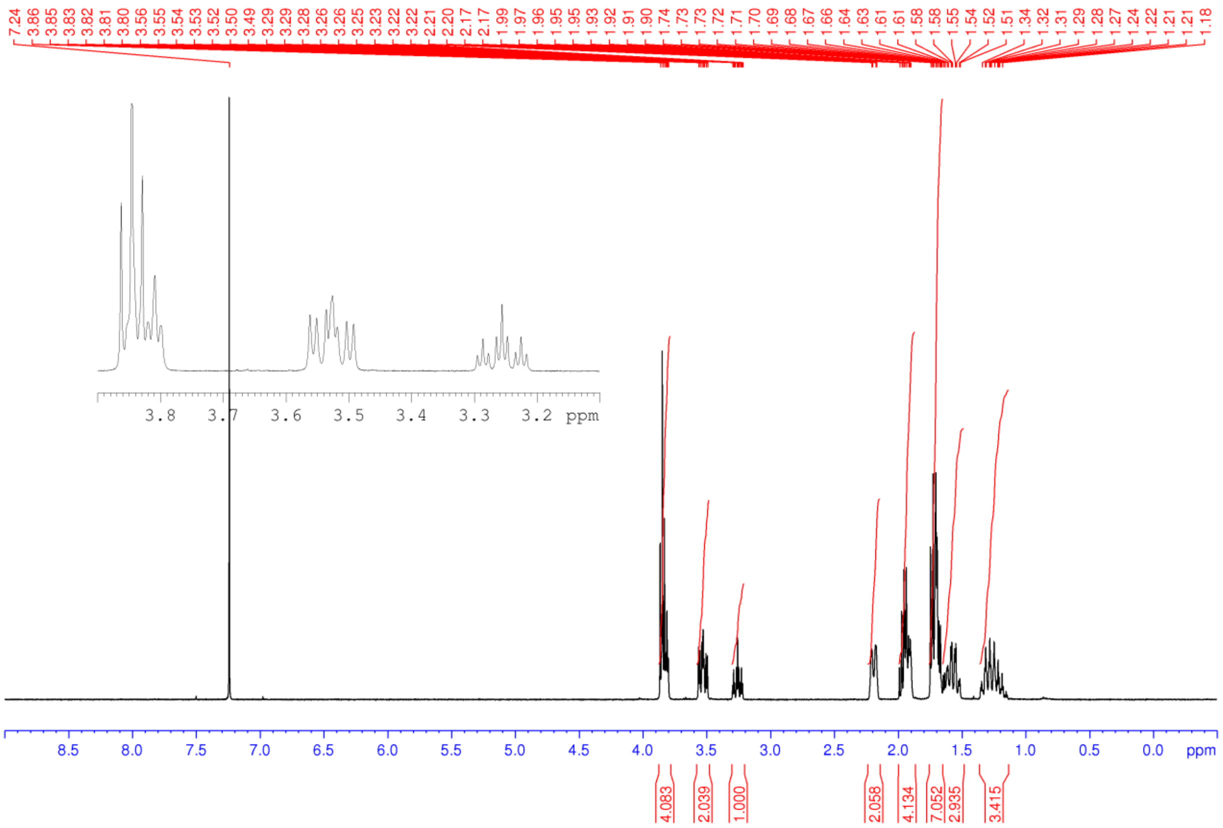

**Figure S60.** $^1$H NMR spectrum (400 MHz, $CDCl_3$) for **42**

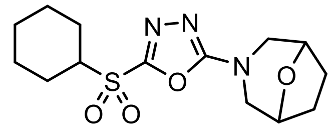

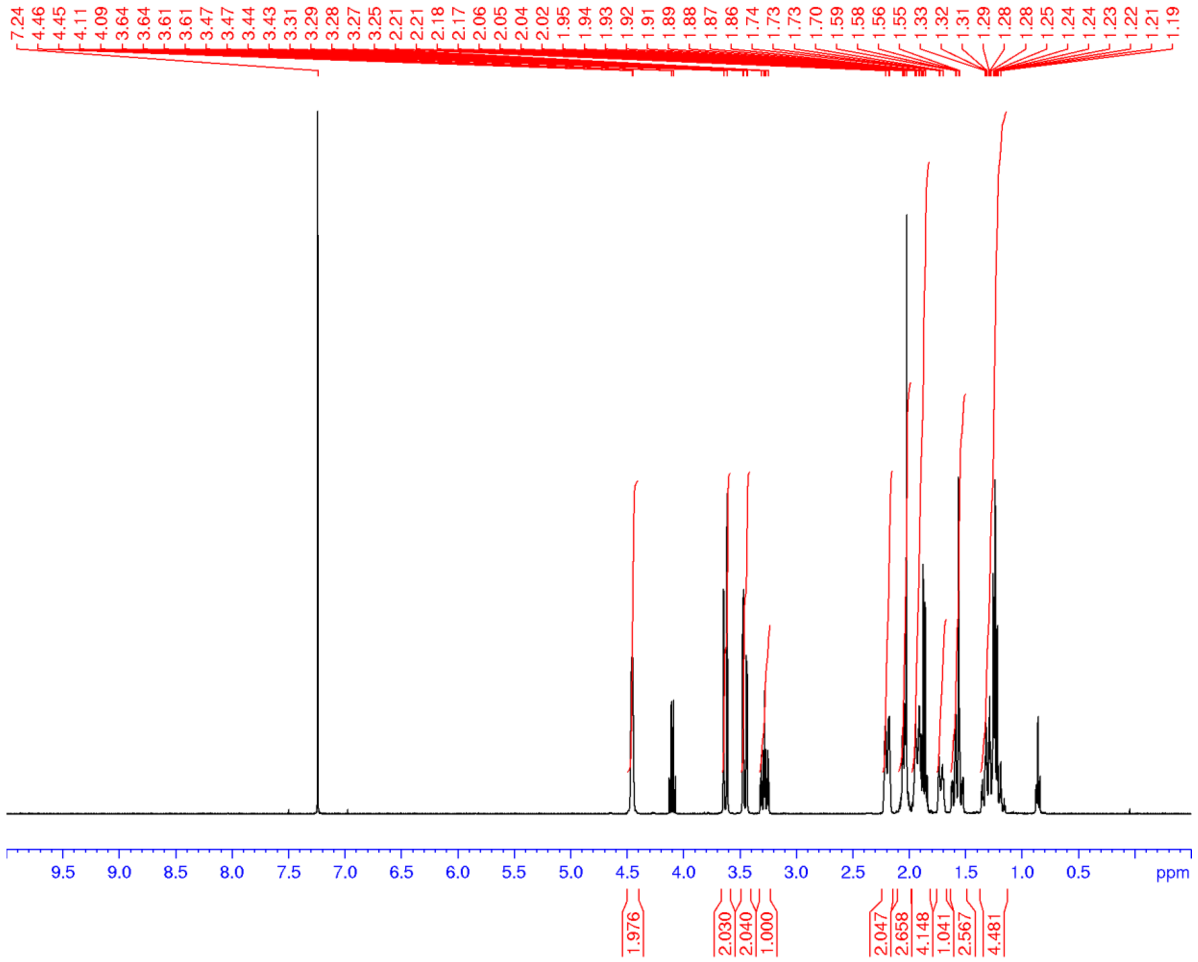

**Figure S61.** $^1$H NMR spectrum (400 MHz, $CDCl_3$) of **43**

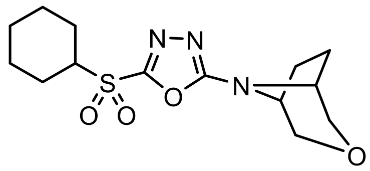

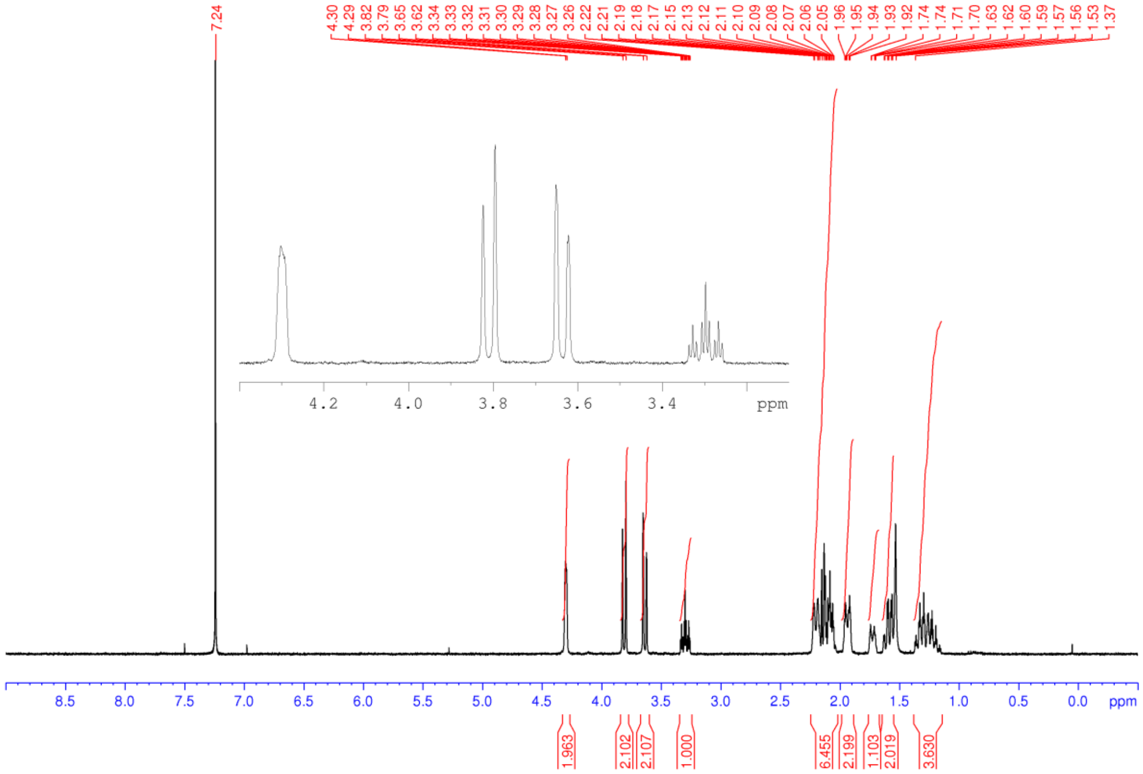

**Figure S62.** $^{1}$H NMR spectrum (400 MHz, $D_3COD$) of **44**

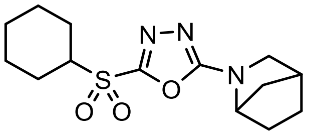

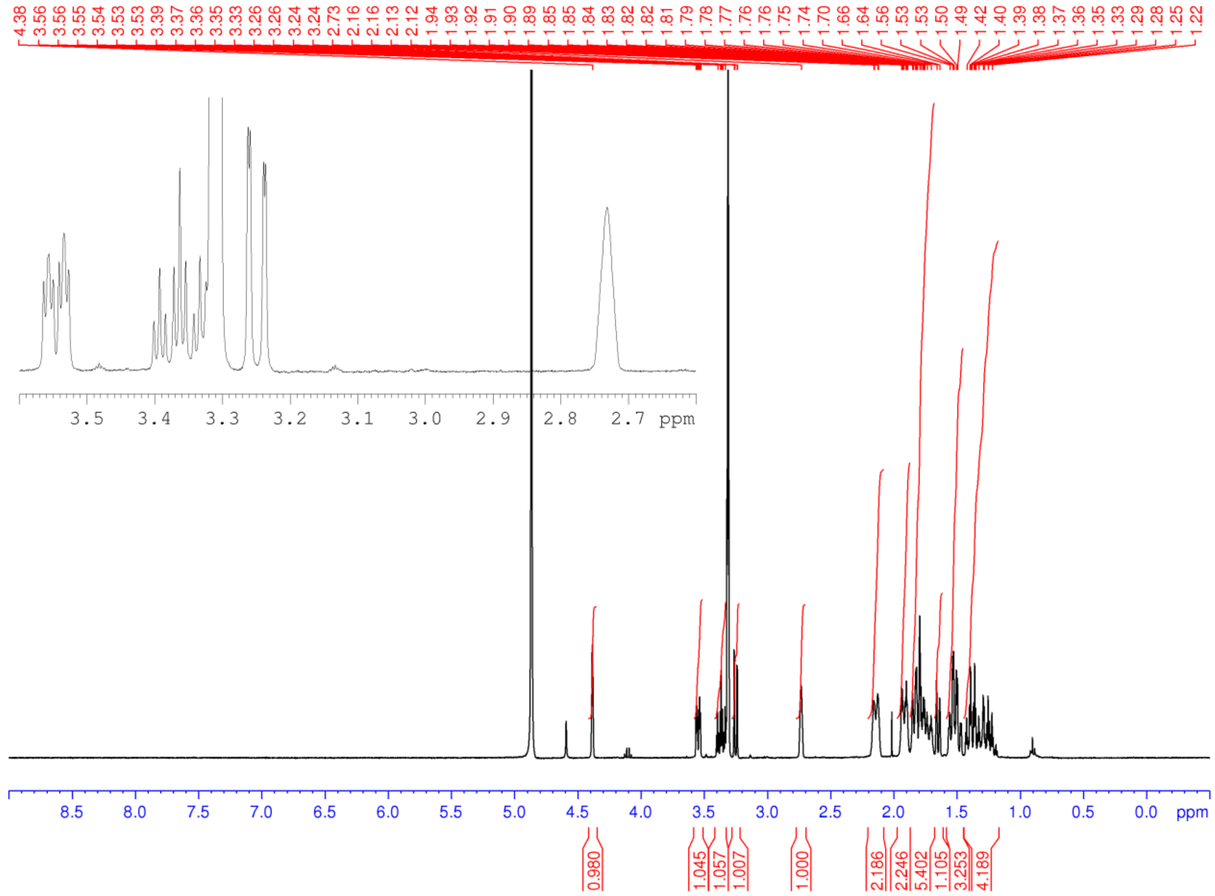

**Figure S63.** $^{1}$H NMR spectrum (400 MHz, $CD_3OD$) of **45**

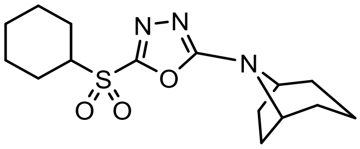


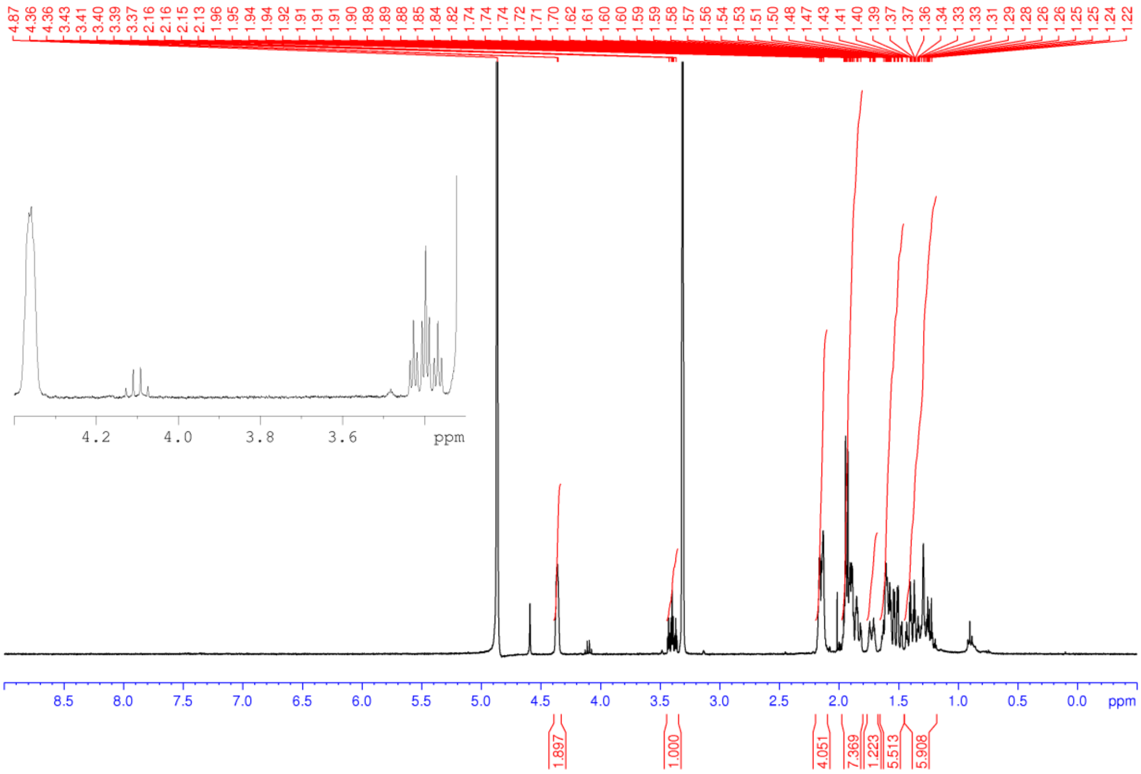

**Figure S64.** $^{1}$H NMR spectrum (400 MHz, $D_3COD$) of **46**

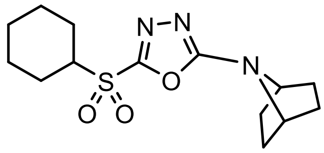

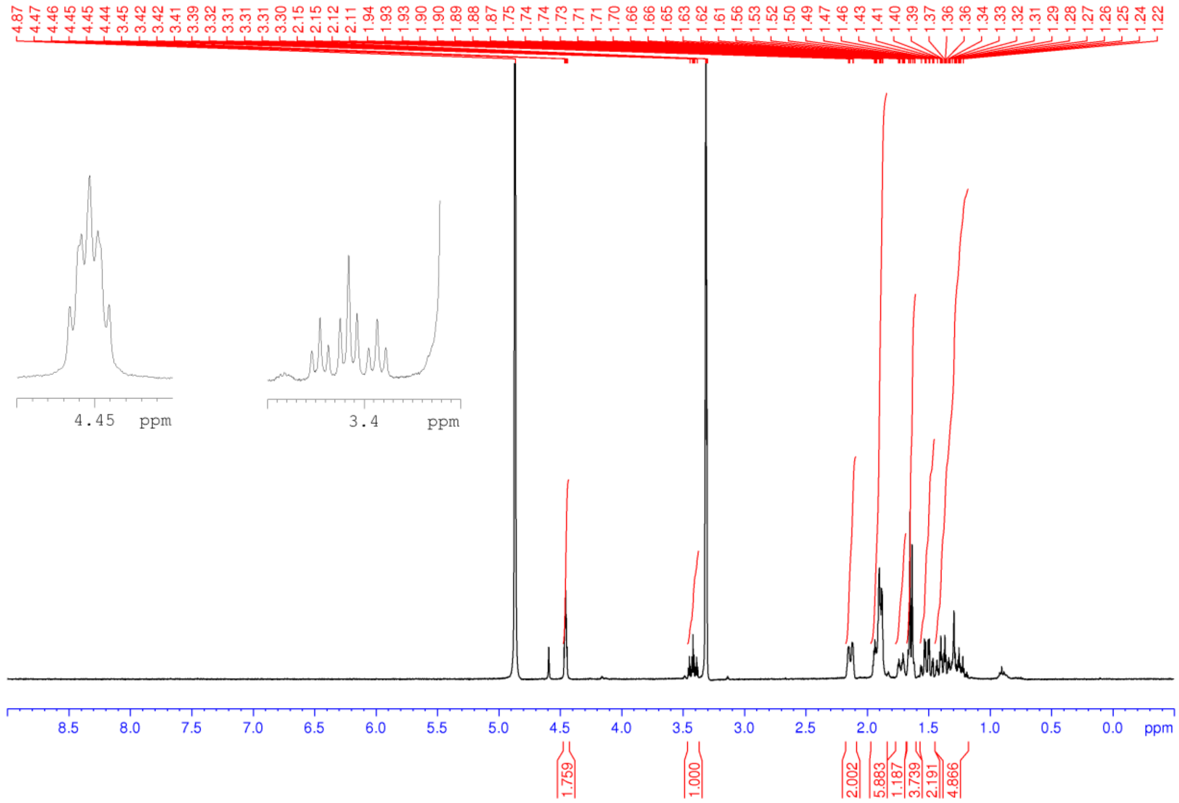

**Figure S65.** $^{1}$NMR spectrum (400 MHz, $D_3COD$) of **47**

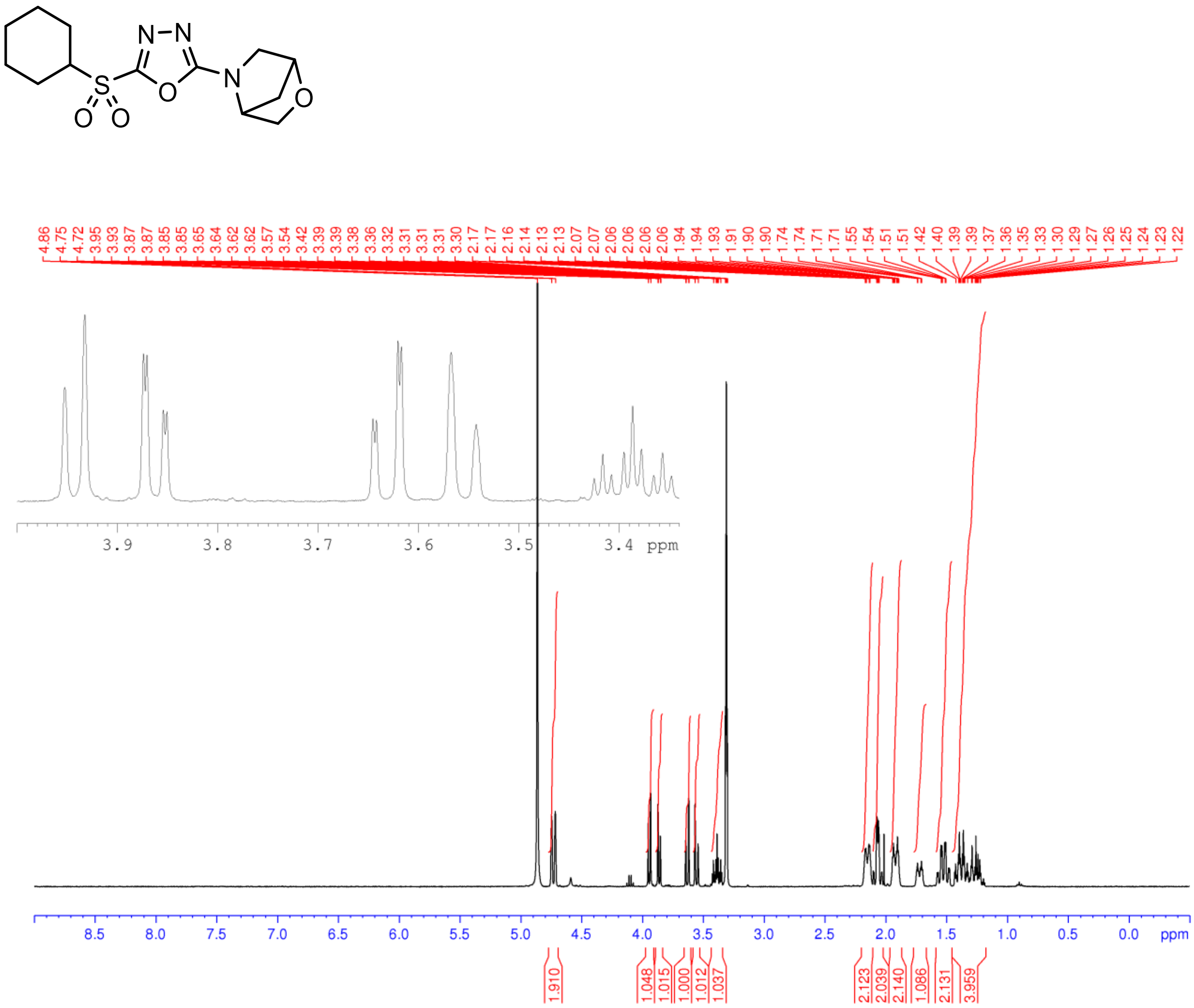

**Figure S66.** $^{1}$H NMR spectrum (400 MHz, $CDCl_3$) for **48**

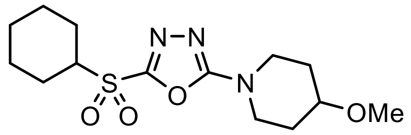


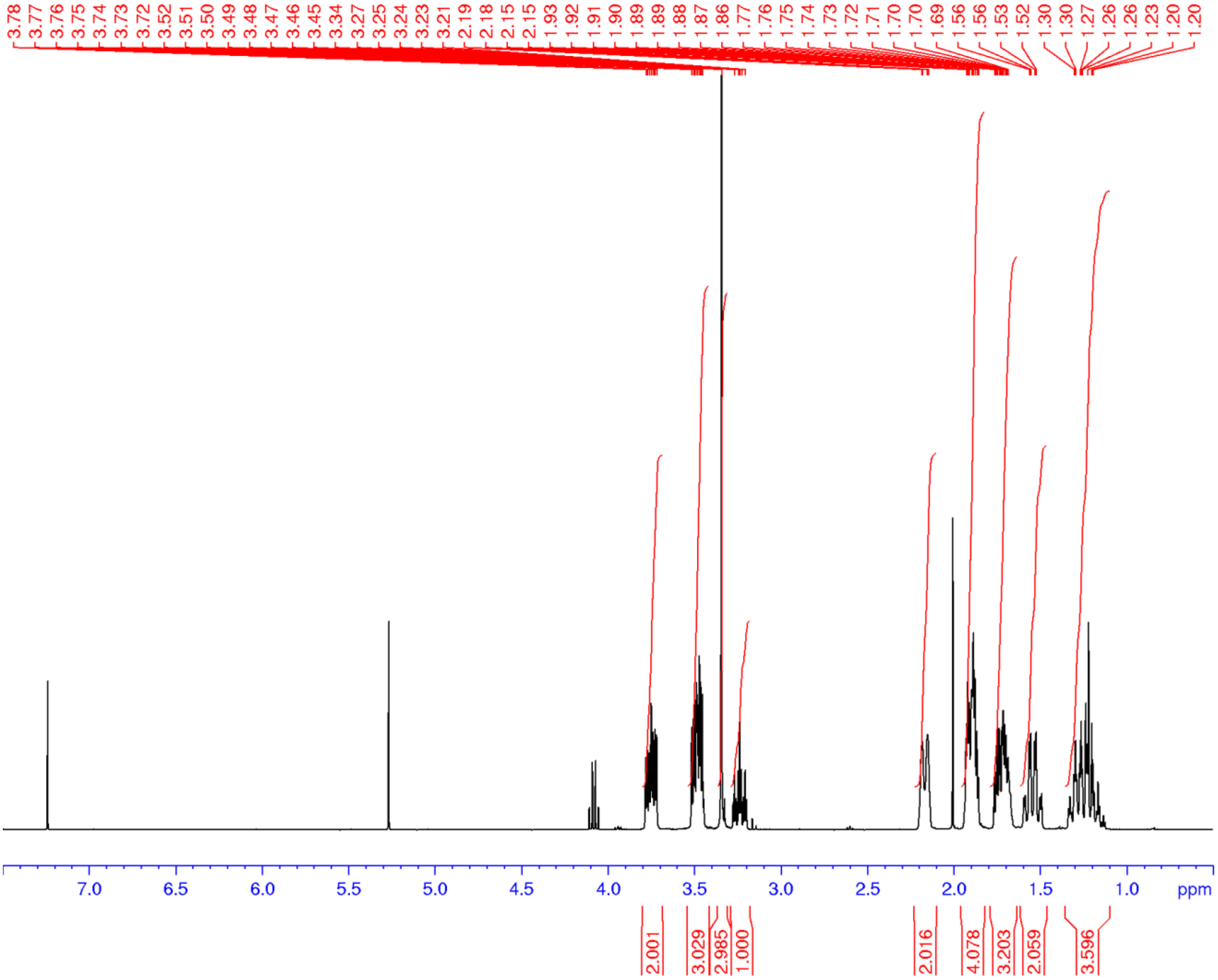

**Figure S67.** $^1$H NMR spectrum (400 MHz, $CDCl_3$) of **49**

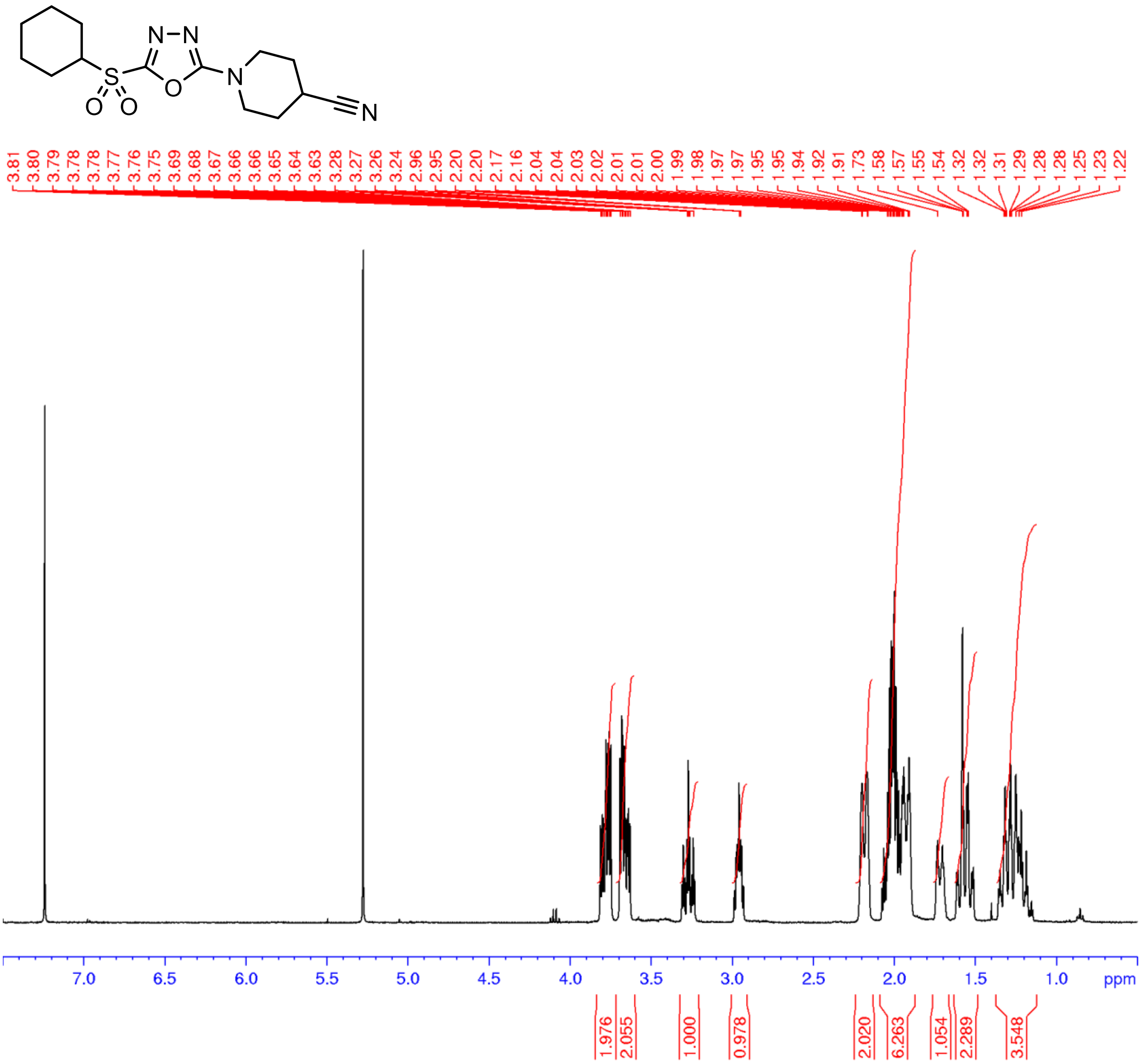

**Figure S68.** $^{1}H$ NMR spectrum (400 MHz, $CDCl_3$) of **50**

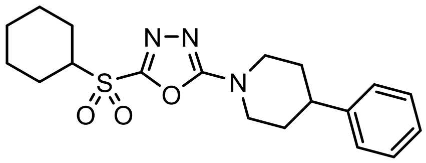

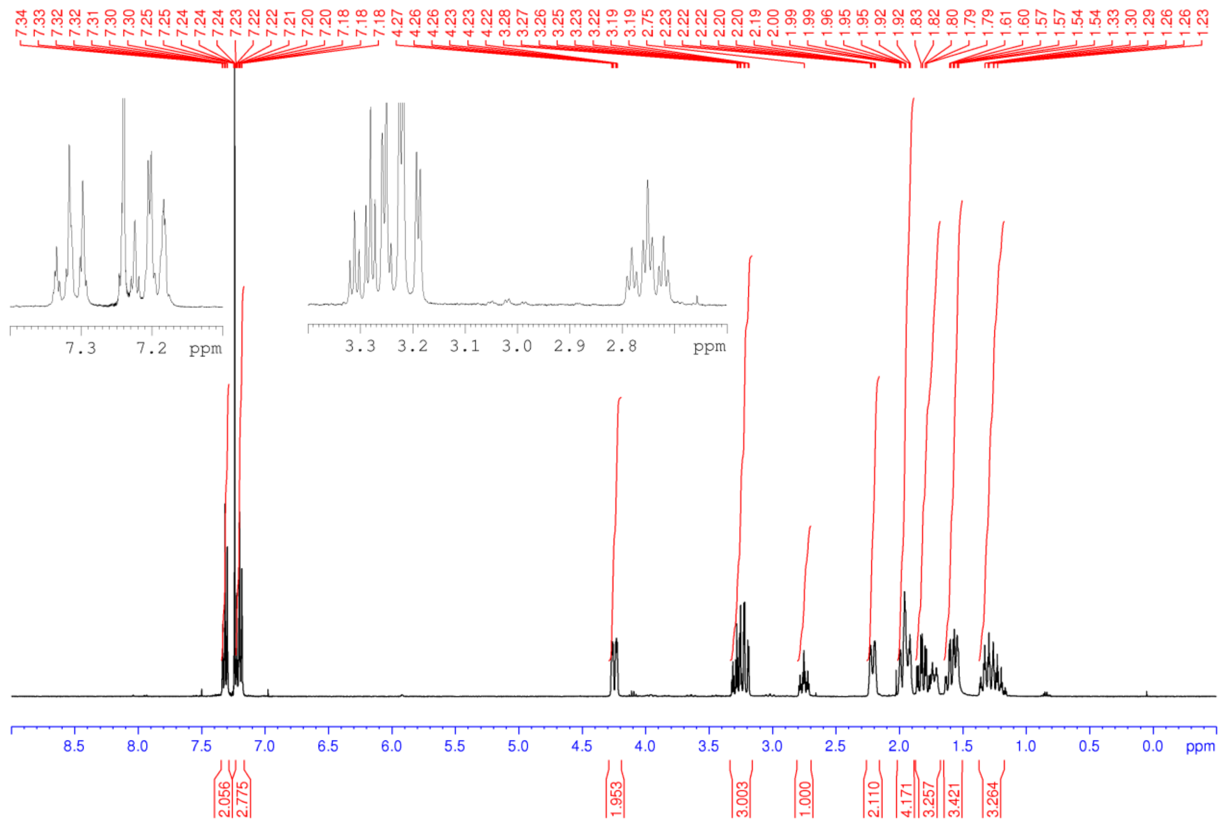

**Figure S69.** $^{1}$H NMR spectrum (400 MHz, $CDCl_3$) of **51**

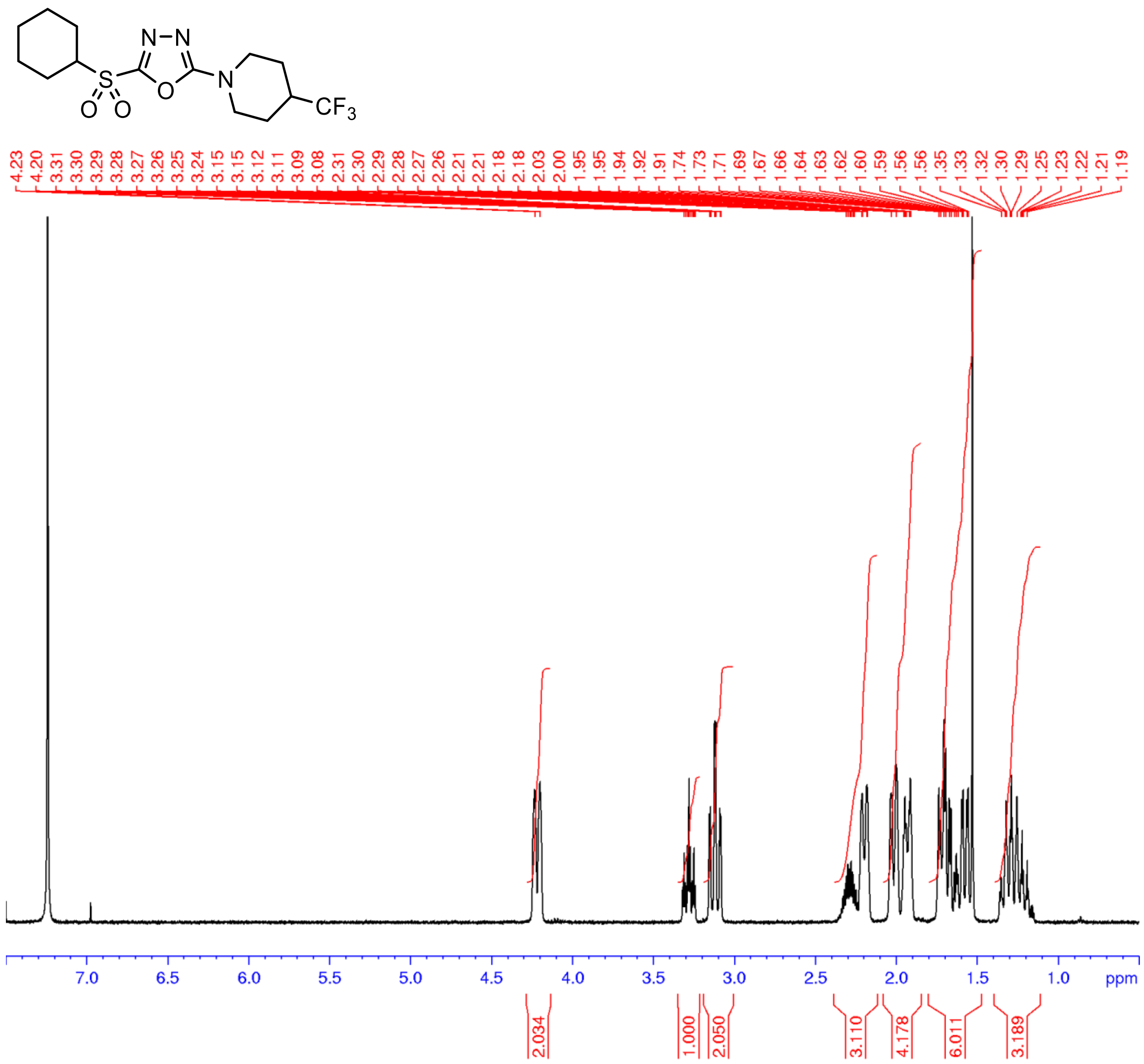

**Figure S70.** $^1$H NMR spectrum (500 MHz, $CDCl_3$) of **52**

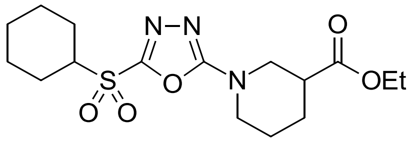

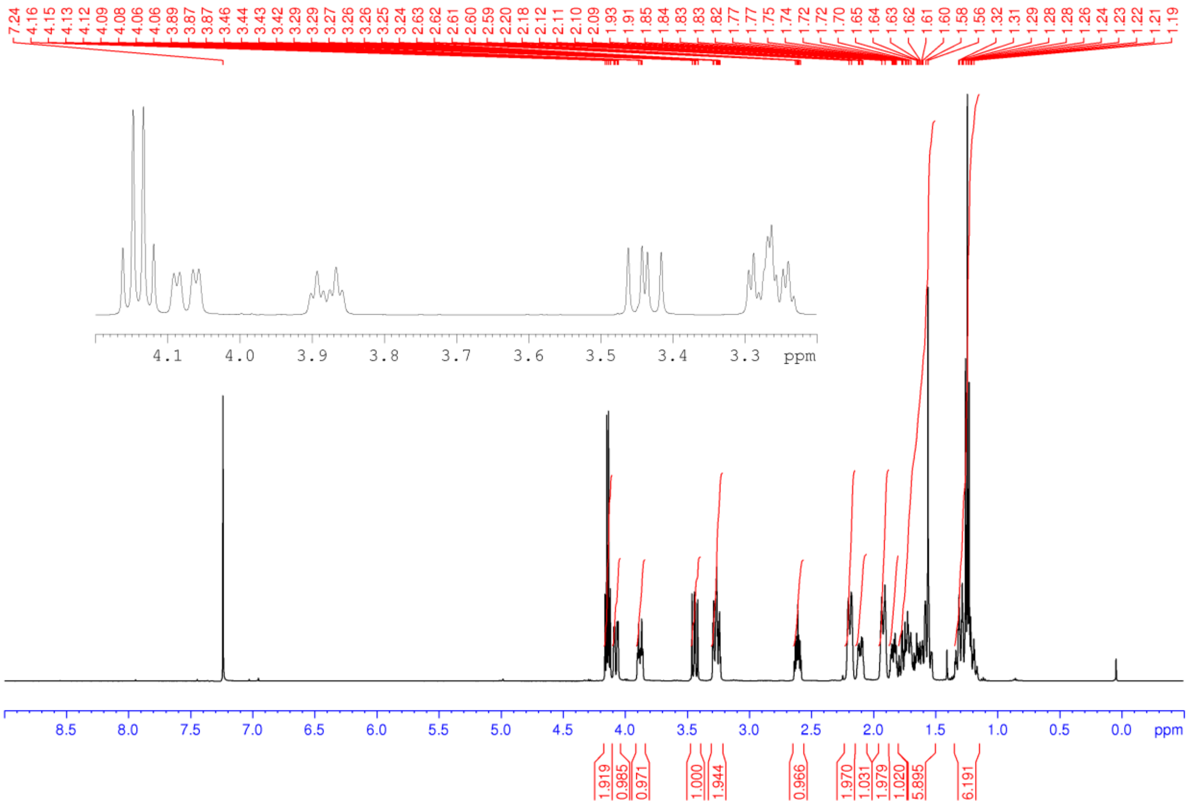

**Figure S71.** [1]H NMR spectrum (400 MHz, $CDCl_3$) of **53**

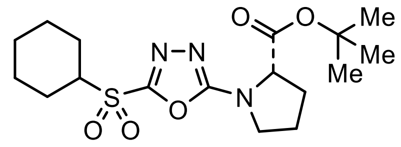


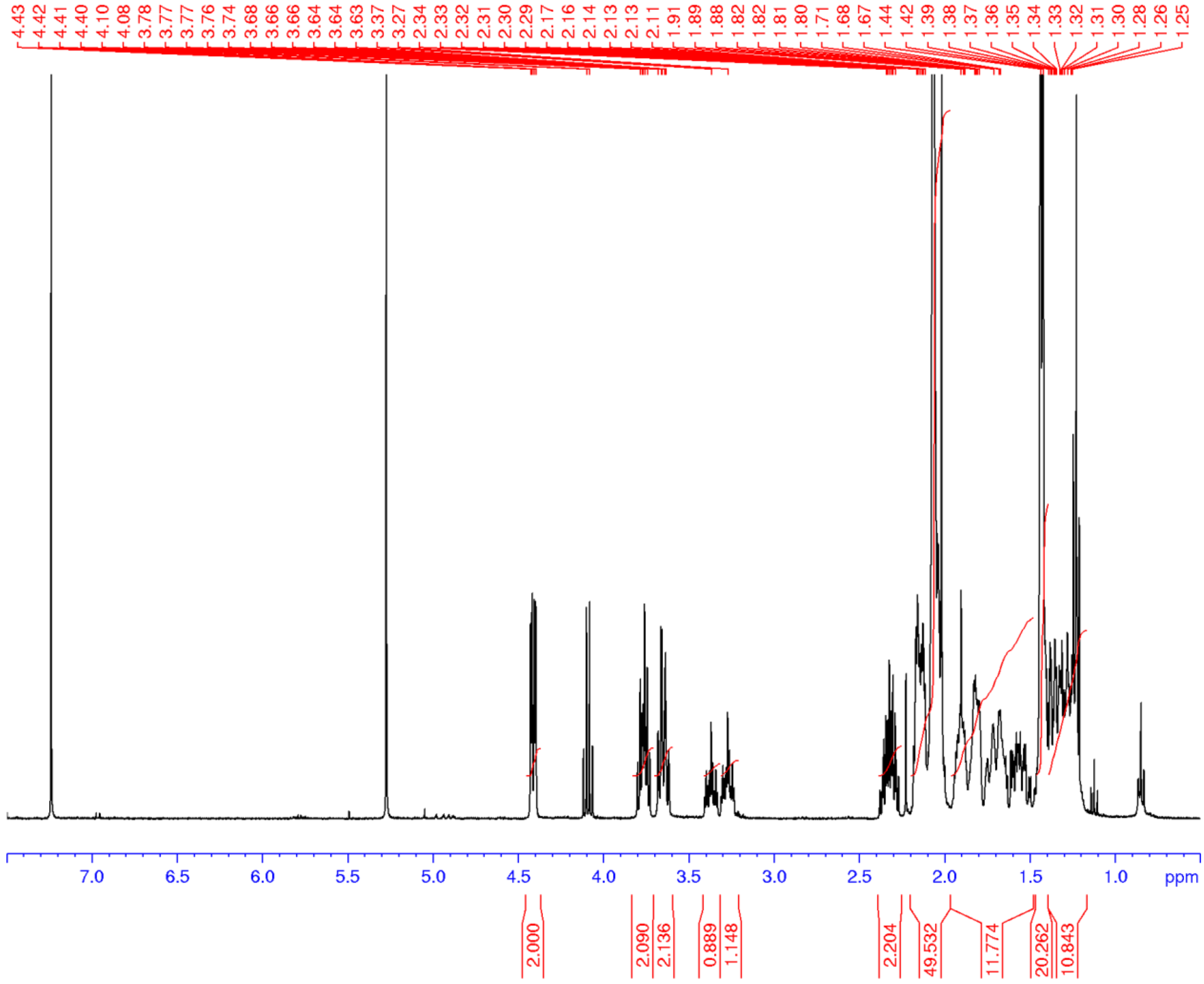

**Figure S72.** $^{1}$H NMR spectrum (400 MHz, $CDCl_3$) of **54**

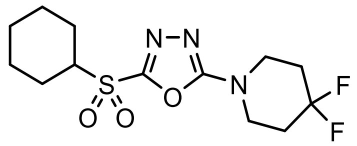

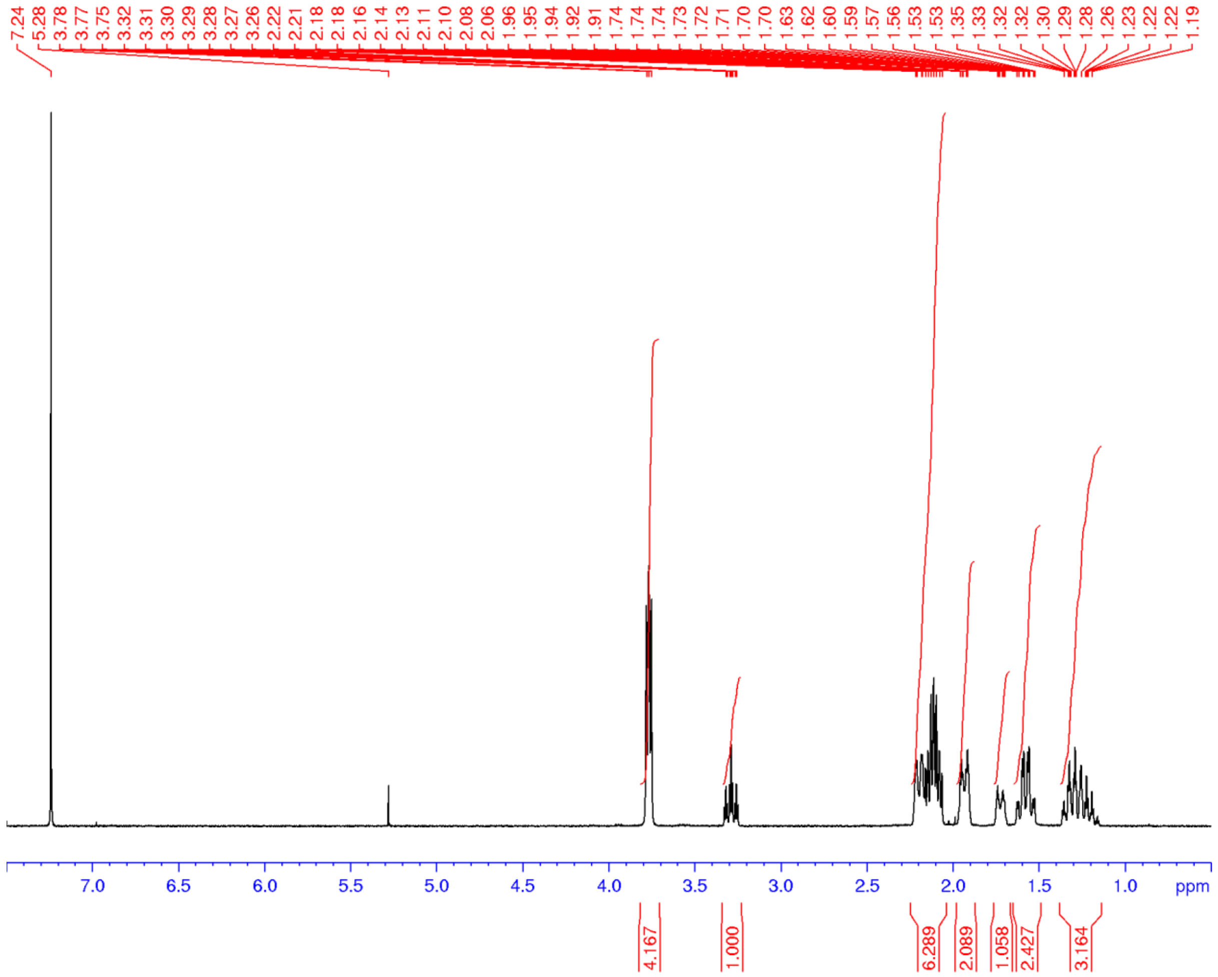

**Figure S73.** $^{1}$H NMR spectrum (400 MHz, $CDCl_3$) of **55**

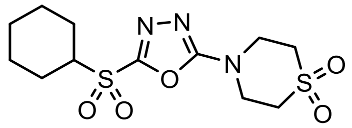

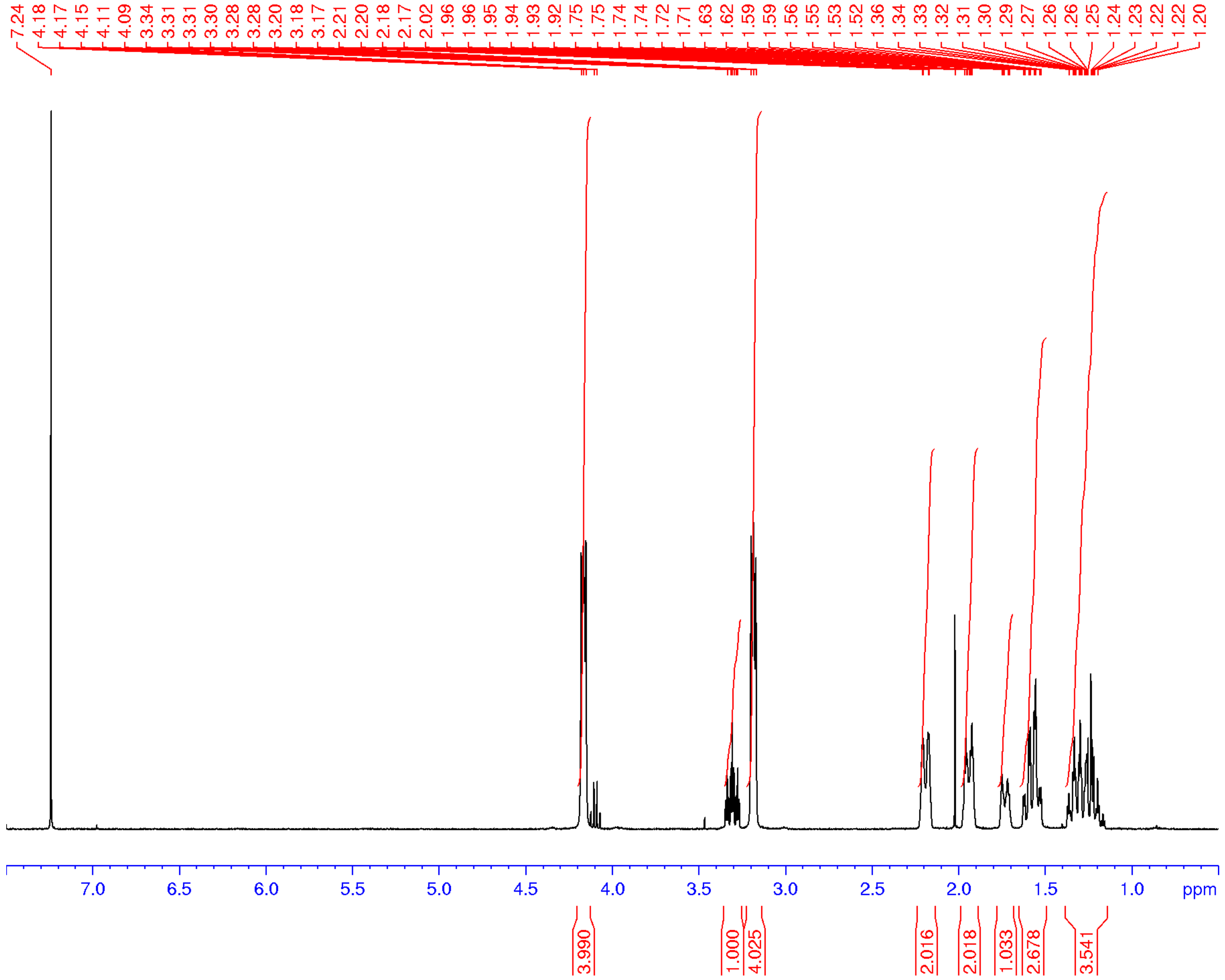

**Figure S74.** $^{1}$H NMR spectrum (400 MHz, $CDCl_3$) of **56**

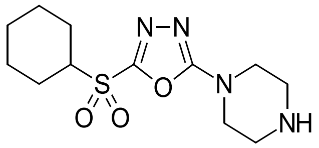

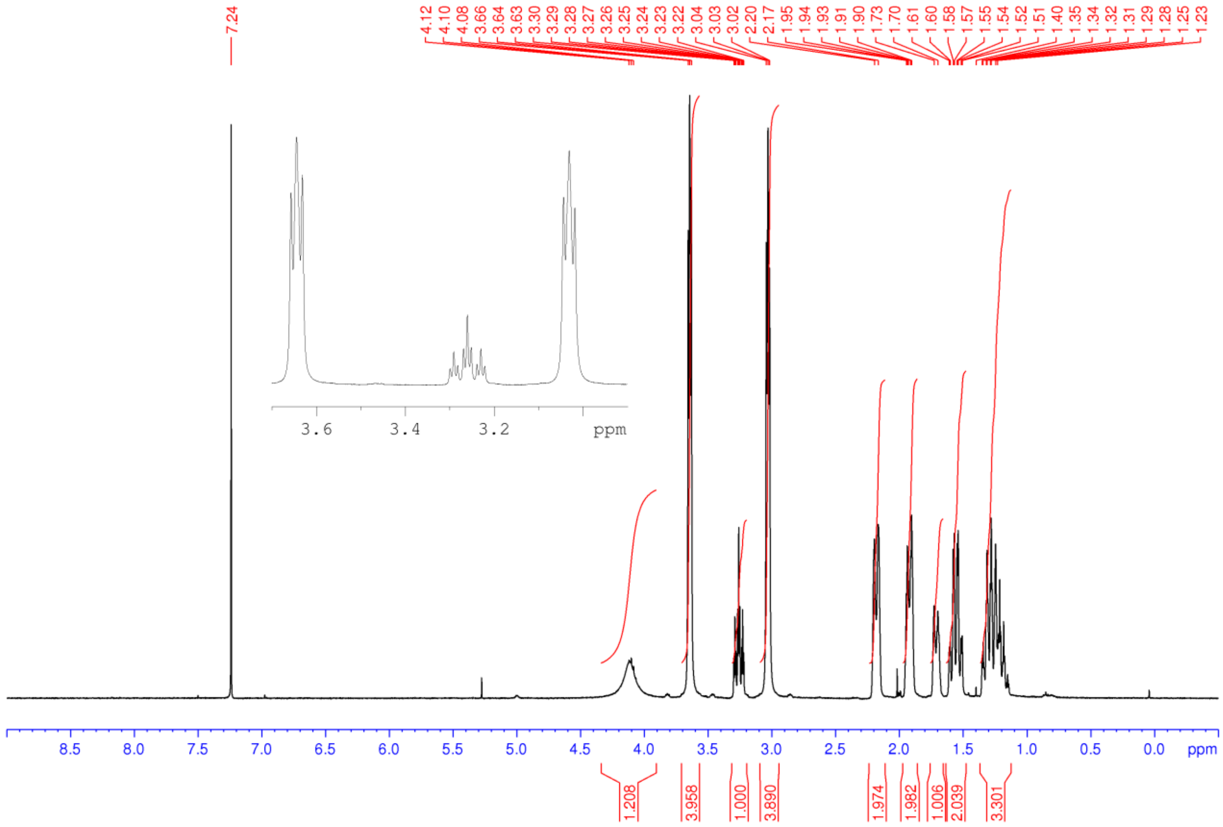

**Figure S75.** $^1$H NMR spectrum (400 MHz, CDCl$_3$) of **57**

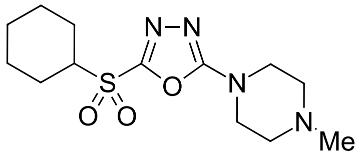


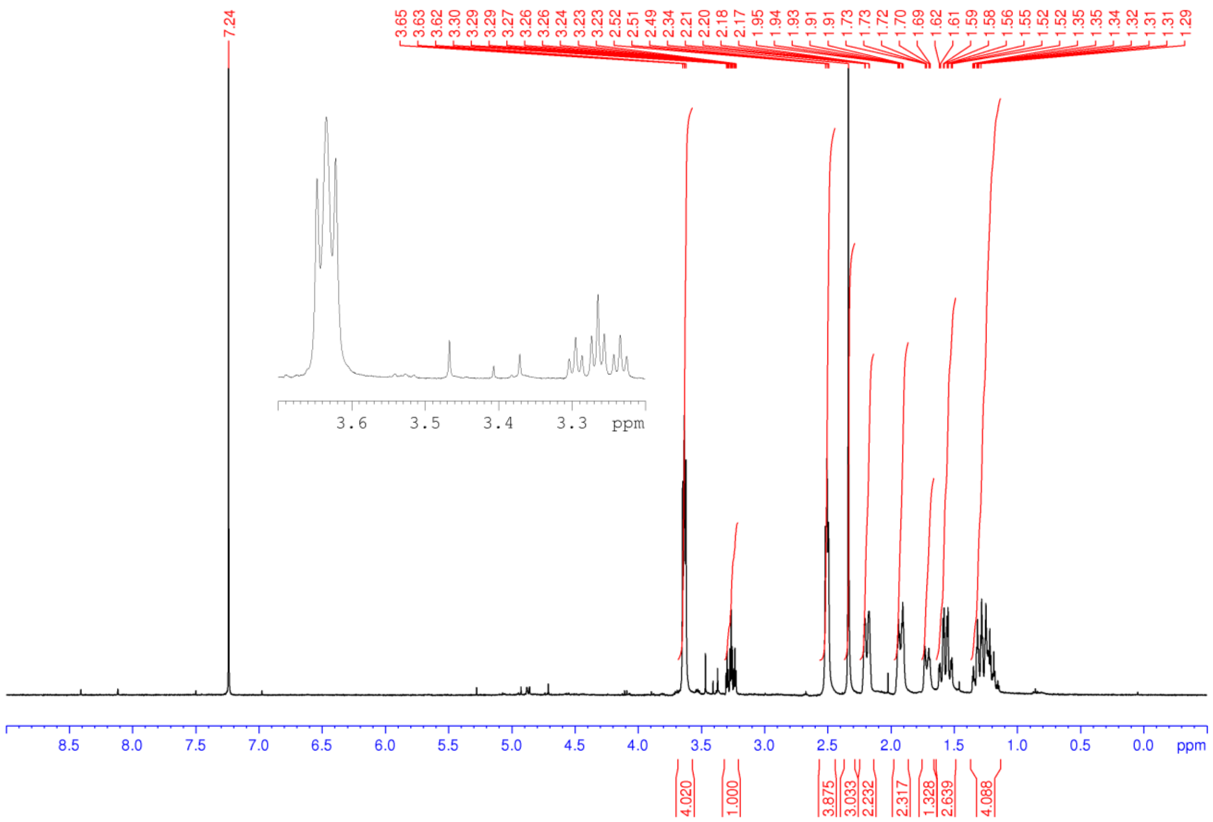

**Figure S76.** $^{1}H$ NMR spectrum (400 MHz, $CDCl_3$) of **58**

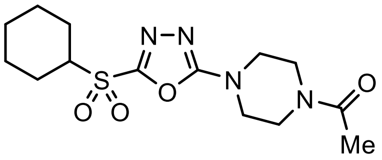


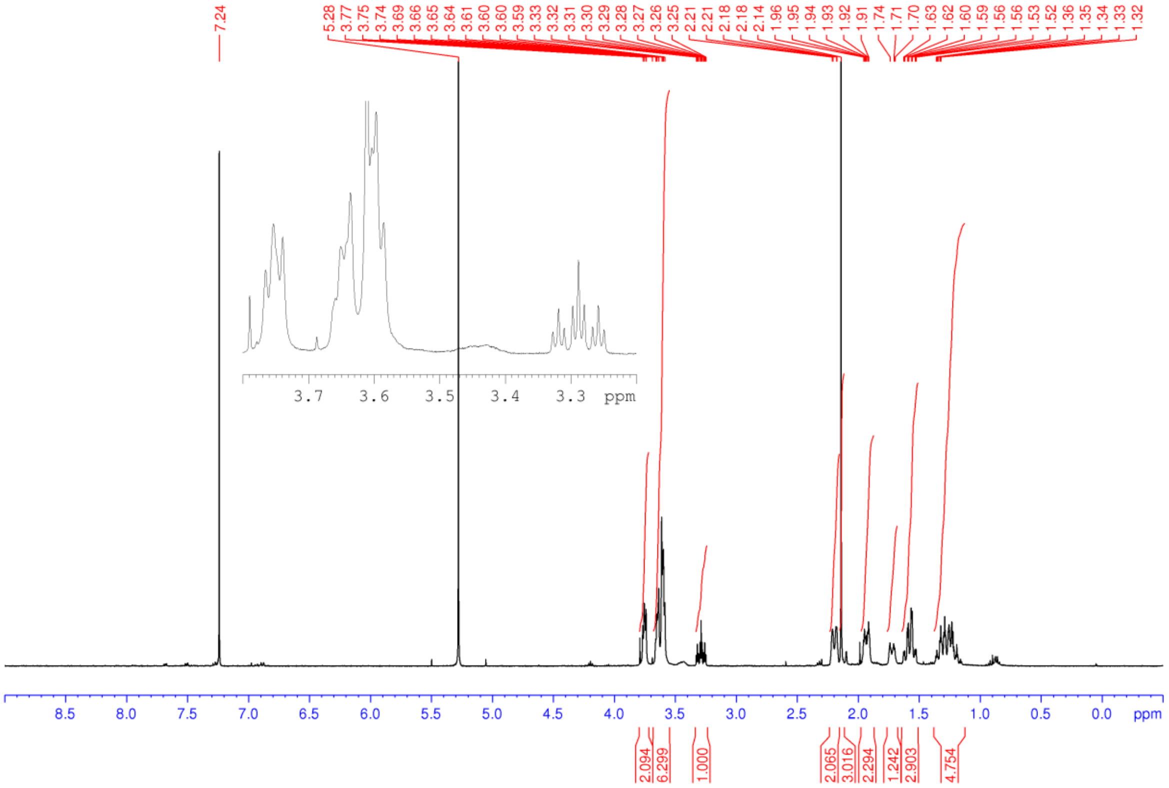

**Figure S77.** $^{1}H$ NMR spectrum (400 MHz, $CDCl_3$) of **59**

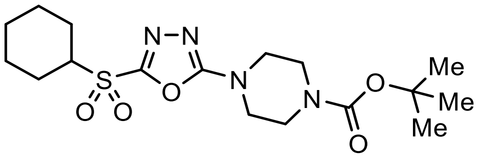


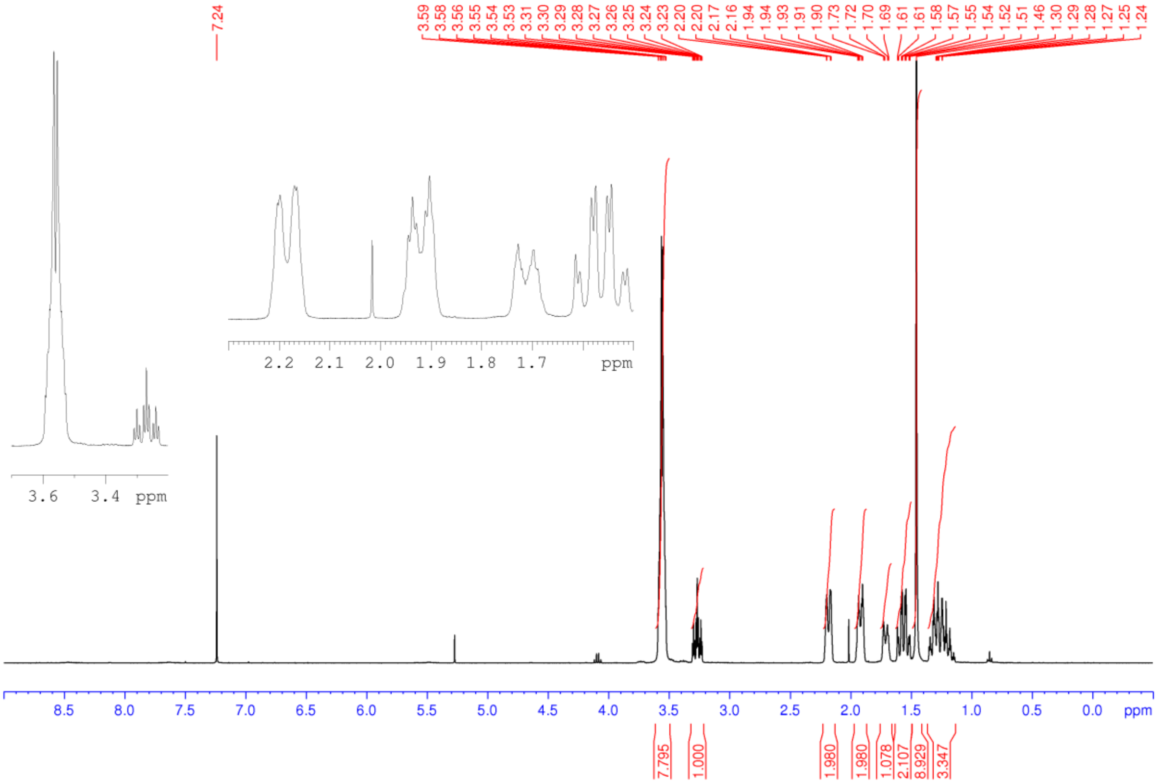

**Figure S78.** $^1$H NMR spectrum (400 MHz, $CDCl_3$) of **60**

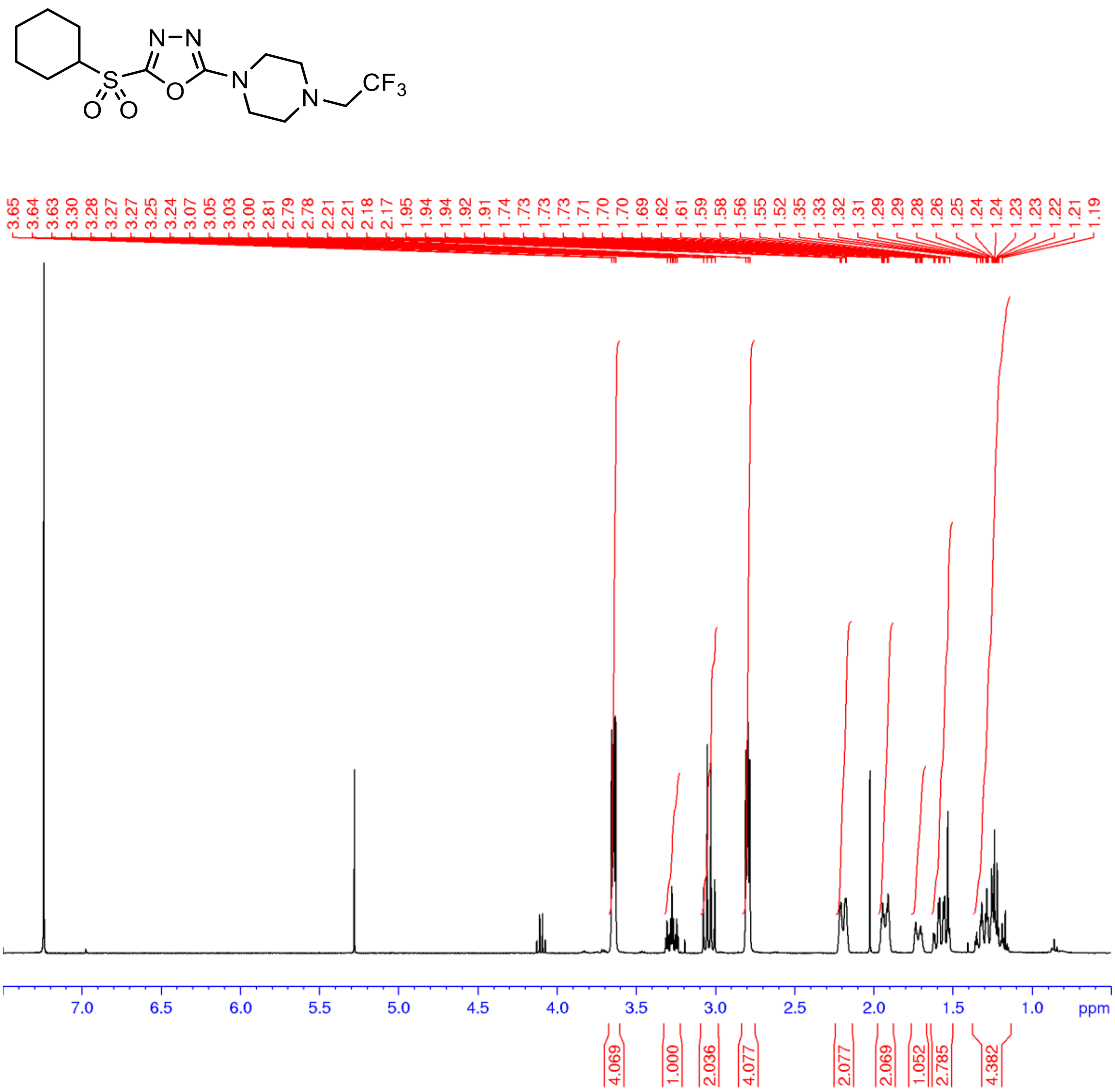

**Figure S79.** $^{1}$H NMR spectrum (500 MHz, $CDCl_3$) of **61**

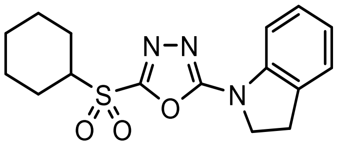

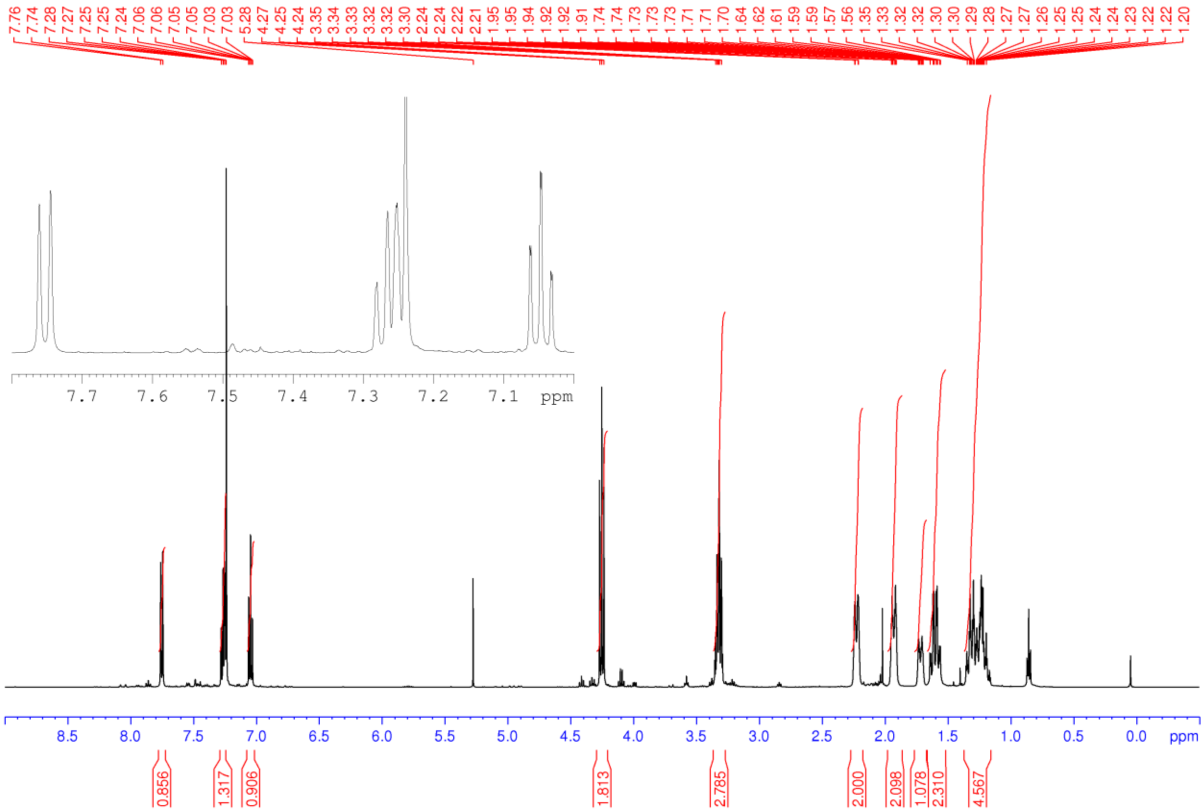

**Figure S80.** $^1$H NMR spectrum (400 MHz, $CDCl_3$) of **62**

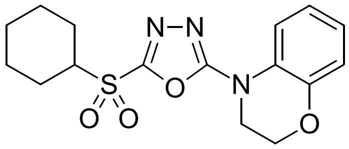

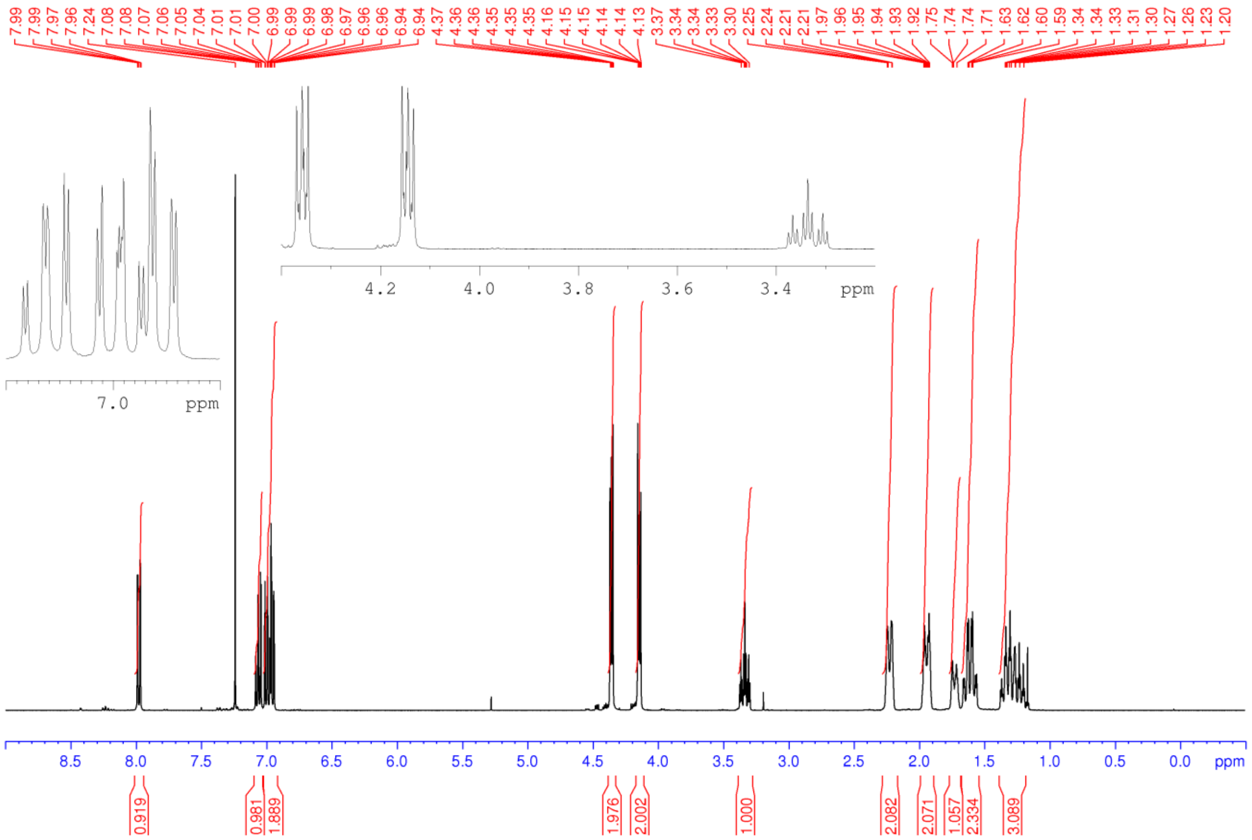
